# Rational Chemical Design of Molecular Glue Degraders

**DOI:** 10.1101/2022.11.04.512693

**Authors:** Ethan S. Toriki, James W. Papatzimas, Kaila Nishikawa, Dustin Dovala, Lynn M. McGregor, Matthew J. Hesse, Jeffrey M. McKenna, John A. Tallarico, Markus Schirle, Daniel K. Nomura

## Abstract

Targeted protein degradation with molecular glue degraders has arisen as a powerful therapeutic modality for eliminating classically undruggable disease-causing proteins through proteasome-mediated degradation. However, we currently lack rational chemical design principles for converting protein-targeting ligands into molecular glue degraders. To overcome this challenge, we sought to identify a transposable chemical handle that would convert protein-targeting ligands into molecular degraders of their corresponding targets. Using the CDK4/6 inhibitor Ribociclib as a prototype, we identified a covalent handle that, when appended to the exit vector of Ribociclib, induced the proteasome-mediated degradation of CDK4 in cancer cells. Covalent chemoproteomic profiling of this CDK4 degrader revealed covalent interactions with cysteine 32 of the RING family E3 ubiquitin ligase RNF126. Structural modification of our initial covalent scaffold led to an improved CDK4 degrader with the development of a but-2-ene, 1,4-dione (**“**fumarate**”**) handle that showed improved interactions with RNF126. Thereafter, we worked to identify the minimum covalent motif required for interaction with RNF126, which we then transplanted onto chemically related and un-related protein-targeting ligands. This strategy successfully produced molecules which induced the degradation of several proteins across diverse protein classes, including BRD4, BCR-ABL and c-ABL, PDE5, AR and AR-V7, BTK, LRRK2, and SMARCA2. Our study undercovers a design strategy for converting protein-targeting ligands into covalent molecular glue degraders.

## Introduction

Targeted protein degradation (TPD) has arisen as a powerful approach for destroying classically undruggable disease-causing proteins through ubiquitination and proteasome-mediated degradation. Two major strategic medicinal chemistry approaches exist for TPD: heterobifunctional Proteolysis Targeting Chimeras (PROTACs) and monovalent molecular glue degraders. Both induce the proximity of an E3 ubiquitin ligase with a target protein leading to the ubiquitination and degradation of the protein in a proteasome-dependent manner ^1–4^. While PROTAC design is more modular wherein protein-targeting ligands can be connected via a linker to an E3 ligase recruiter, the discovery of novel molecular glue degraders has mostly been either fortuitous from phenotypic screens or via specific well-characterized E3 ligase-targeting recruiters (e.g. for cereblon) ^5–11^. Given this landscape, a rational chemical design principle for converting protein-targeting ligands into molecular glue degraders would be highly desired thereby facilitating a modular target-based design strategy of molecular glue degraders that is currently lacking.

An example of a fortuitously discovered molecular glue degrader is thalidomide, which was originally developed as a morning sickness medicine but found to cause phocomelia birth defects; this effect was subsequently found to be due to ternary complex formation between the E3 ubiquitin ligase substrate receptor cereblon and the transcription factor SALL4, leading to SALL4 ubiquitination and proteasome-mediated degradation ^7,12,13^. Immunomodulatory Drug (IMiD) thalidomide analogs, such as pomalidomide and lenalidomide, have also been developed as anti-cancer therapeutics that similarly act through ternary complex formation between cereblon and oncogenic Ikaros transcription factors, leading to Ikaros degradation ^5^. Subsequent medicinal chemistry efforts with IMiD thalidomide analogs, both as molecular glues and in the context of PROTACs have led to the development of many additional neo-substrate degradation events exploiting cereblon, including many PROTACs exploiting cereblon recruiters for targeted protein degradation ^5,9,10^. Another example of a fortuitously discovered molecular glue degrader is Indisulam, an arylsulfonamide drug with selective anticancer activity, that was later discovered to exert its therapeutic activity through ternary complex formation between CUL4-DCAF15 and RBM39 leading to RBM39 degradation ^14^.

Moving beyond fortuitous findings, cell-based phenotypic screening has also arisen as a powerful approach for discovering novel molecular glue degraders. Chemical screening for anti-cancer phenotypes coupled with counter-screening in hyponeddylated lines impaired in Cullin E3 ligase activity with subsequent multi-omic mechanistic deconvolution has led to the discovery of novel molecular glue degraders that act through interactions of CDK12-cyclin K with a CRL4B ligase complex to induce cyclin K degradation ^15^. Phenotypic screening of covalent ligands for NEDDylation- and proteasome-dependent anti-cancer activity coupled with chemoproteomic approaches were also used to discover a covalent molecular glue between the E2 ubiquitin conjugating enzyme UBE2D and the transcription factor NFKB1 to ubiquitinate and degrade NFKB1 leading to impaired cancer cell viability ^16^. These cell-based phenotypic screening approaches can also be deployed to perform target-based molecular glue degrader screens using cell lines expressing target proteins bearing high-throughput screening-detectable tags (e.g. nanoluciferase, HiBiT) ^11^.

Several recent independent studies with specific protein targets and compounds have also revealed the possibility for converting protein-targeting ligands into molecular glue degraders. Correlative analysis between the cytotoxicity of clinical and pre-clinical small-molecules and the expression of E3 ligase components across hundreds of human cancer cell lines identified CR8, a CDK12 inhibitor analog of *(R)-*roscovitine, that bears a 2-pyridyl moiety on the solvent exposed terminus of the molecule that engages in a ternary complex between CDK12-cyclin K and the CUL4 adaptor protein DDB1, leading to cyclin K ubiquitination and degradation ^17^. Roscovitine itself does not induce cyclin degradation, but the mere addition of a 2-pyridyl moiety converted this compound into a molecular glue degrader. In another study, a BCL6 inhibitor, BI-3802, was discovered to be a degrader of BCL6 through BI-3802-mediated polymerization of BCL6 leading to recognition of BCL6 filaments by the SIAH1 E3 ligase and subsequent ubiquitination and proteasomal degradation ^18^. While the mechanism of degradation is quite unique from a traditional molecular glue degrader, this study highlighted that a minor chemical modification was sufficient to move from the non-degradative BCL6 inhibitor BI-3812 to the BCL6 degrader BI-3802. In another striking example, Genentech scientists discovered a propargyl amine chemical handle that when appended onto the non-degradative BET family bromodomain inhibitor JQ1 led to BRD4 degradation ^19^.

There are thus many emerging innovative strategies for discovering novel molecular glue degraders, but their discoveries have largely been either through cell-based phenotypic screens or several independent one-off isolated examples of minor structural modifications to non-degradative inhibitors that have converted these compounds into degraders of their targets or associated protein complexes. The latter examples indicate that rational chemical design principles may exist to convert non-degradative protein-targeting ligands more systematically and modularly into molecular glue degraders of those targets, but such design principles are still poorly understood. To address this challenge, we sought to identify a chemical handle that could be modularly appended onto protein-targeting ligands across different chemical and protein classes to convert these ligands into molecular glue degraders of their targets. Through these efforts, we discovered a minimal covalent chemical moiety that can be appended onto various protein targeting ligands to induce the proteasome-mediated degradation of their target proteins.

## Results

### Synthesis and Testing of Analogs with Chemical Handles Appended to the Exit Vector of the CDK4/6 Inhibitor Ribociclib

To identify potential chemical handles that could convert protein-targeting ligands into molecular glue degraders of their targets, we appended various moieties onto the solvent-exposed end of the CDK4/6 inhibitor Ribociclib **(Figure 1a)** ^20^. Among the 9 initial Ribociclib analogs tested, we found one compound, EST1027, a trifluoromethylphenyl cinnamamide, that led to >50 % reduction in CDK4/6 levels in C33A cervical cancer cells treated for 24 hours at 3 μM **(Figure 1b-1d)**. This EST1027-mediated reduction in CDK4 occurred via proteasome-mediated degradation, since pre-treatment of C33A cells with the proteasome inhibitor bortezomib attenuated EST1027-mediated CDK4 degradation **(Figure 1e-1f)**. The cinnamamide motif is a possible covalent substrate that can undergo 1,4-addition with a cysteine thiolate anion. Confirming the necessity of this covalent functionality for the degradation of CDK4 by EST1027, a non-reactive trifluoromethylphenyl propionamide analog EST1036 did not induce the degradation of CDK4 in C33A cells **(Figure 1g-1i)**.

**Figure 1.**
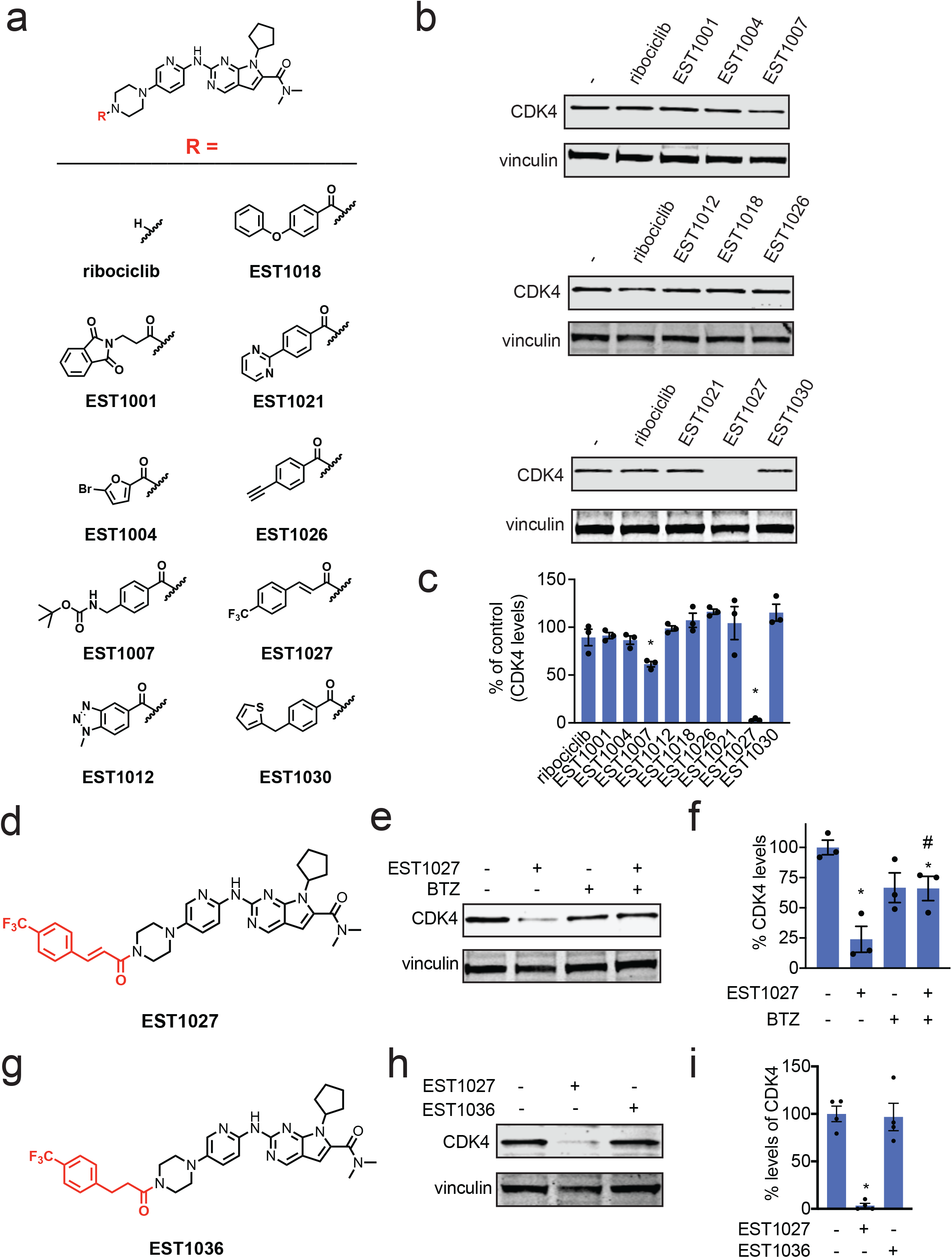
Synthesis and Testing of Analogs with Chemical Handles Appended to the Exit Vector of the CDK4/6 Inhibitor Ribociclib. **(a)** Structures of Ribociclib analogs wherein various chemical handles were appended onto the solvent-exposed end of Ribociclib. **(b)** Testing Ribociclib analogs in C33A cervical cancer cells to identify compounds that reduce CDK4 levels. C33A cells were treated with DMSO vehicle or compounds (3 μM) for 24 h. CDK4 and loading control vinculin levels were assessed by Western blotting. **(c)** Quantification of the data shown in **(b). (d)** Full structure of hit compound EST1027 that showed >50 % loss of CDK4 in **(b-c)** with the appended chemical handle shown in red. **(e)** Proteasome-dependent degradation of CDK4 by EST1027. C33A cells were pre-treated with DMSO vehicle or the proteasome inhibitor bortezomib (BTZ) (10 μM) 1 h prior to treatment of cells with DMSO vehicle or EST1027 (5 μM) and CDK4 and loading control vinculin levels were assessed by Western blotting. **(f)** Quantification of the experiment described in **(e). (g)** Structure of EST1036, a non-reactive derivative of EST1027. **(h)** EST1036 does not degrade CDK4. C33A cells were treated with DMSO vehicle or compounds for (5 μM) for 24 h and CDK4 and loading control vinculin levels were assessed by Western blotting. **(i)** Quantification of experiment in **(h)**. Blots shown in **(b, e, h**) are representative of n=3 biologically independent replicates/group. Bar graphs in **(c, f, i)** show individual replicate values and average ± sem. Statistical significance is calculated as *p<0.05 compared to DMSO vehicle in **(c, f, i)** and #p<0.05 compared to EST1027-treated group in **(f)**.

### Characterizing the Mechanism of the CDK4 Degrader

Based on our data indicating that the covalent functionality on EST1027 was necessary to induce the degradation of CDK4, we postulated that this motif was leading to the specific and covalent recognition of particular E3 ubiquitin ligases, leading to molecular glue interactions between this E3 ligase and CDK4 and subsequent CDK4 ubiquitination and proteasome-mediated degradation. To identify the E3 ligase covalently targeted by EST1027, we performed isotopic tandem orthogonal activity-based protein profiling (isoTOP-ABPP) in which we treated C33A cells *in situ* with vehicle or EST1027, and subsequently labeled resulting cell lysates with an alkyne-functionalized iodoacetamide probe to identify cysteines that were highly engaged by EST1027 across the proteome ^21–24^. Out of 2726 quantified probe-modified cysteines that we observed in at least 2 out of 3 biological replicates, we identified 6 targets that showed control/EST1027 ratios of >2 with an adjusted p-value of p<0.05. Among these six targets, we identified cysteine 32 (C32) on the RING-family E3 ubiquitin ligase RNF126 as a target of EST1027 that showed a control/EST1027 ratio of 2.5 **(Figure 2a; Table S1)**. RNF126 is an important E3 ligase involved in cellular protein quality control and is necessary to ubiquitinate and degrade mislocalized proteins in the cytosol through its association with Bag6 and ubiquitination of BAG6-associated client proteins ^25^. RNF126 is also necessary to degrade p97-extracted membrane proteins from the endoplasmic reticulum through association with BAG6 and secondary ubiquitination on target proteins ^26^. Interestingly, C32 is a known zinc-coordinating cysteine within RNF126 ^27^. We previously also identified a covalent RNF4 E3 ligase recruiter that could be used in PROTAC applications that also targeted a zinc-coordinating cysteine, while maintaining RNF4 function ^28^. Analogous to our previous study with RNF4, we posited that labeling by EST1027 of only one of the zinc-coordinating cysteines still allows functional zinc coordination with the other cysteines C13, C16, and C29 and maintaining RNF126 function. We confirmed the interaction of EST1027 with RNF126 by gel-based ABPP methods by showing dose-responsive competition of EST1027 against rhodamine-functionalized iodoacetamide cysteine labeling of recombinant RNF126 without causing any protein precipitation **(Figure 2b)**. We also observed the mass adduct of EST1027 on pure RNF126 protein by mass spectrometry analysis of EST1027 labeled RNF126 tryptic digests **(Figure 2c-2d)**. RNF126 knockdown completely attenuated EST1027-mediated CDK4 degradation in C33A cells, demonstrating that RNF126 was at least required for the degradation of CDK4 **(Figure 2e-2f)**. Interestingly, we also observed significant reduction in RNF126 levels with EST1027 treatment, which was potentially through the proteasomal degradation of the CDK4-EST027-RNF126 ternary complex **(Figure 2e-2f)**.

**Figure 2.**
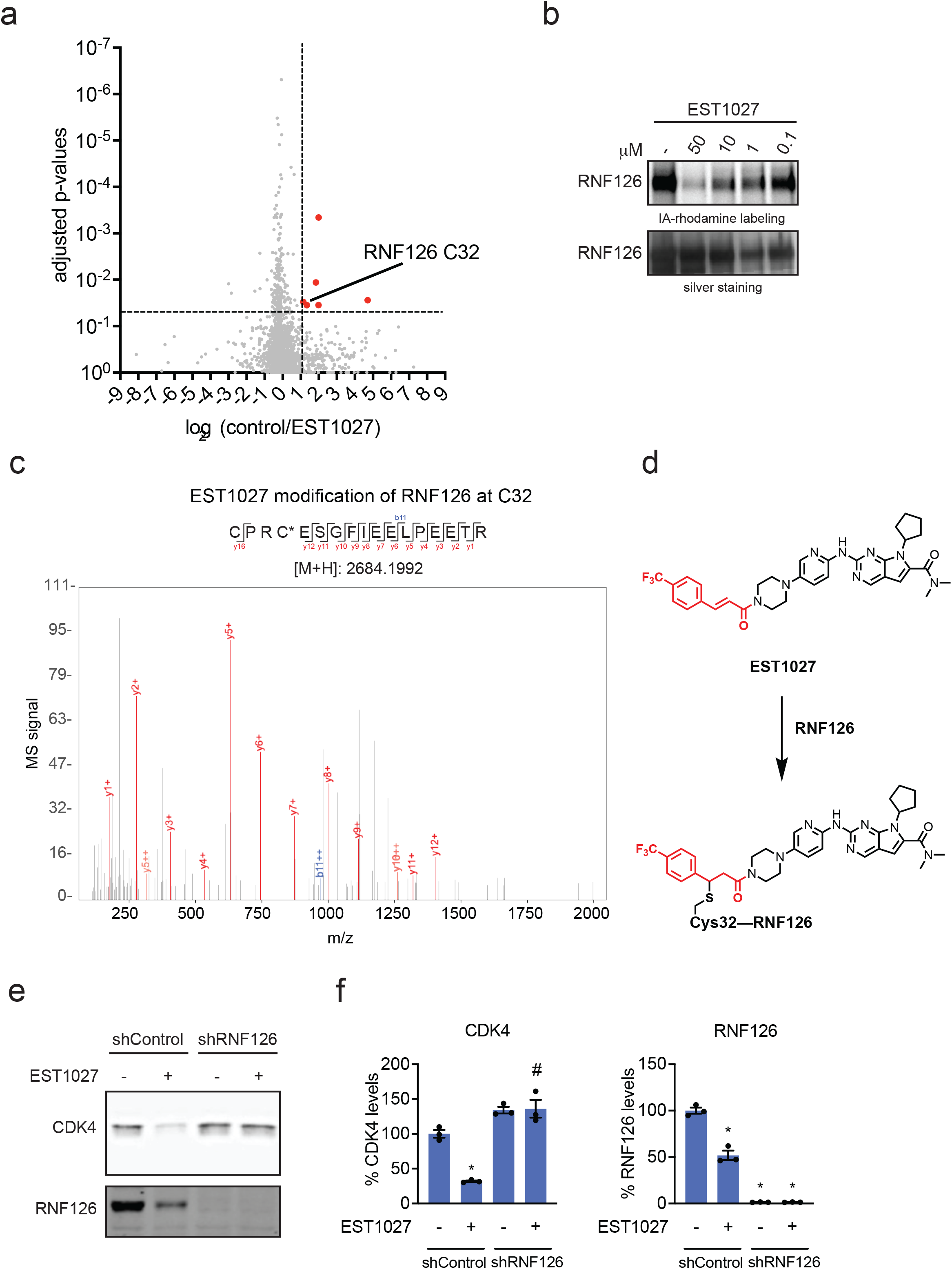
Characterizing the Mechanism of the CDK4 Degrader. **(a)** cysteine chemoproteomic profiling of EST1027 in C33A cells using isoTOP-ABPP. C33A cells were treated with DMSO vehicle or EST1027 (10 μM) for 2 h. Resulting lysates were labeled with an alkyne-functionalized iodoacetamide probe (IA-alkyne) (200 μM) for 1 h, after which isotopically light (for control) or heavy (for treated) biotin-azide tags bearing a TEV cleavable site were appended onto probe-labeled proteins by copper-catalyzed azide-alkyne cycloaddition. Control and treated groups were combined in a 1:1 ratio, avidin-enriched, tryptically digested, and probe-modified tryptic peptides were eluted by TEV protease and analyzed by LC/MS/MS and probe-modified peptides were quantified. Shown in red are probe-modified cysteines that showed control/EST1027 ratios >2 with adjusted p<0.05 from n=3 biologically-independent replicates. **(b)** gel-based ABPP of EST1027 against RNF126. Recombinant RNF126 was pre-incubated with DMSO vehicle or EST1027 for 30 min prior to labeling of RNF126 with IA-rhodamine (50 nM) for 1 h. Gels were visualized by in-gel fluorescence and protein loading was assessed by silver staining. **(c)** Site of modification analysis of EST1027 with pure RNF126 protein. RNF126 protein (24 μg) was incubated with EST1027 (50 μM) for 30 min and tryptic digests were analyzed by LC-MS/MS. **(d)** Structure of presumed adduct of EST1027 with C32 of RNF126. **(e)** RNF126 knockdown attenuates EST1027-mediated CDK4 degradation. RNF126 was stably knocked down in C33A cells using short hairpin oligonucleotides (shRNF126) compared to non-targeting shControl oligonucleotides. C33A shControl and shRNF126 cells were treated with DMSO vehicle or EST1027 (5 μM) for 24 h. CDK4, RNF126, and loading control vinculin levels were assessed by Western blotting. **(f)** Quantification of the experiment in **(e)**. Gels and blots in **(b, e)** are representative images from n=3 biologically independent replicates/group. Bar graphs in **(f)** show individual replicate values and average ± sem. Statistical significance is calculated as *p<0.05 compared to DMSO vehicle-treated shControl cells and #p<0.05 compared to EST1027-treated shControl group.

### Structure-Activity Relationship of CDK4 Degrader

Encouraged by these data, we postulated that this trifluoromethylphenyl cinnamamide moiety was conferring binding and reactivity with RNF126 and that this motif could be appended onto other protein-targeting ligands to induce their degradation. Thus, we next appended this motif onto another structurally similar CDK4/6 inhibitor, Palbociclib, to generate EST1090 **(Figure S1a)**. Disappointingly, EST1090 did not show any binding to RNF126 and did not degrade CDK4 in C33A cells **(Figure S1b-S1c)**, indicating that this chemical motif was not generalizable and could not be simply transplanted onto other even very similar protein-targeting ligands to induce targeted degradation.

We therefore sought to explore structure-activity relationships of the cinnamamide motif that we had identified to induce the degradation of CDK4. Still using Ribociclib as our testbed, we generated seven additional analogs in an effort to identify a better covalent chemical module **(Figure 3a)**. Interestingly, moving the trifluoromethyl moiety from the *para*-to *ortho-* and/or *meta*-positions with EST1051, EST1054, and KN1002 abrogated CDK4 degradation in C33A cells, giving further support to specific interactions of these degrader compounds with an E3 ligase rather than non-specific mechanisms **(Figure 3a-3b)**. Merely appending an acrylamide handle with EST1057 also did not cause CDK4 degradation **(Figure 3a-3b)**. Among additional derivatives tested, we observed improved dose-responsive CDK4 degradation with EST1060, with a methoxyphenyl but-2-ene-1,4-dione (**“**fumarate derivative**”**), in C33A cells **(Figure 3a-3d)**. EST1060 also showed significantly improved potency against RNF126 compared to our original EST1027, showing strong displacement of cysteine probe labeling of recombinant RNF126 down to 1 μM **(Figure 3e)**. We confirmed that EST1060 still reacts with C32 of RNF126 by mass spectrometry analysis of EST1060 labeled RNF126 tryptic digests **(Figure 3f-3g)**. A non-reactive derivative of EST1060, JP-2-230, interestingly still showed binding to RNF126, albeit weaker than EST1060, indicating the fumarate motif may possess reversible binding affinity to RNF126 beyond its inherent reactivity **(Figure S2a-S2b)**. Nonetheless, this non-reactive analog did not degrade CDK4 in C33A cells **(Figure S2c)**.

**Figure 3.**
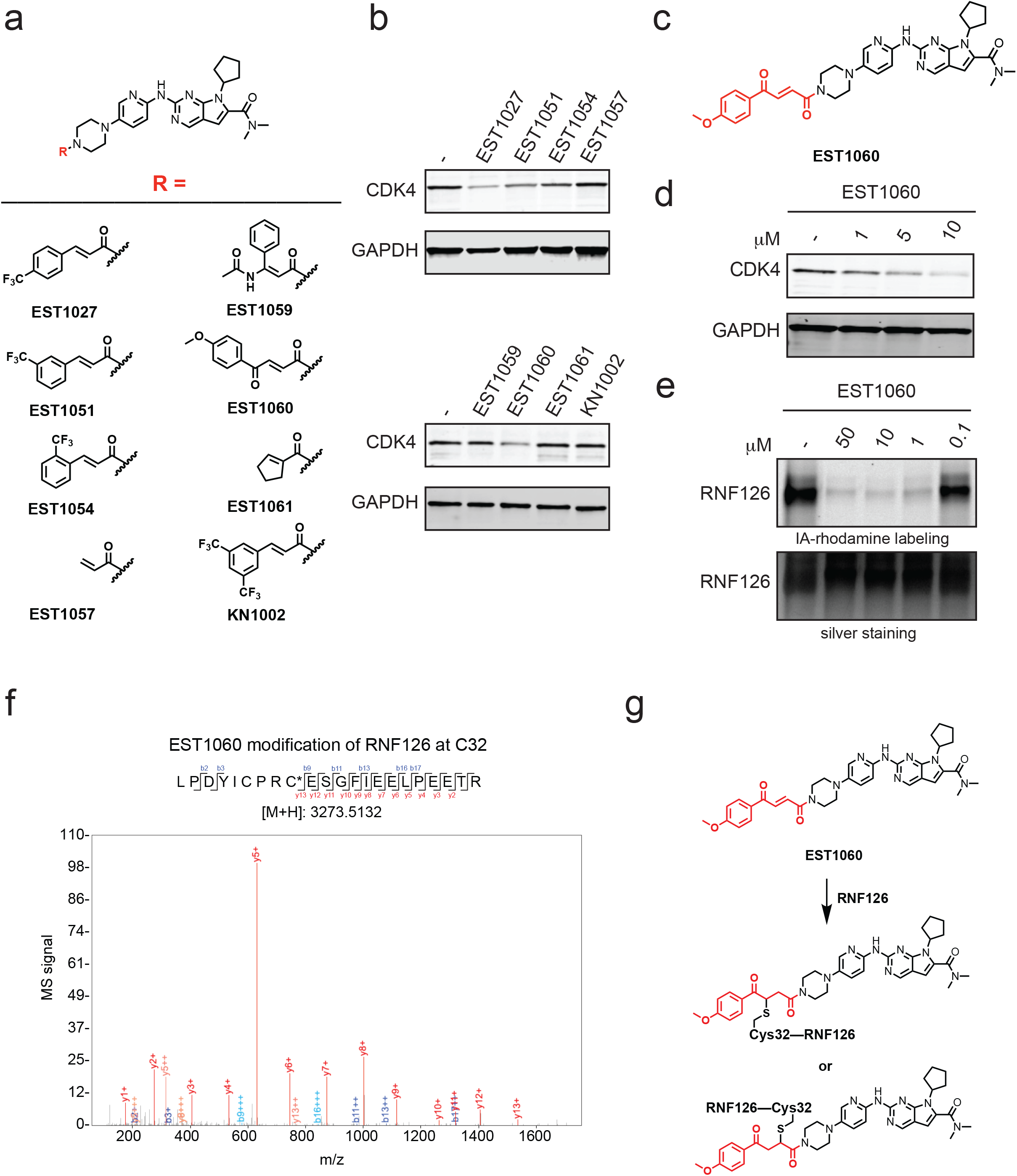
Structure-Activity Relationship of CDK4 Degrader. **(a)** Structures of EST1027 analogs assessing structure-activity relationships. **(b)** Testing EST1027 analogs in C33A cervical cancer cells to identify compounds that reduce CDK4 levels. C33A cells were treated with DMSO vehicle or compounds (5 μM) for 24h. CDK4 and loading control GAPDH levels were assessed by Western blotting. **(c)** Full structure of hit compound EST1060. **(d)** Dose-response of EST1060 CDK4 degradation. **(d)** C33A cells were treated with DMSO vehicle or EST1060 (5 μM) for 24 h. CDK4 and loading control GAPDH levels were assessed by Western blotting. **(e)** gel-based ABPP of EST1060 against RNF126. Recombinant RNF126 was pre-incubated with DMSO vehicle or EST1060 for 30 min prior to labeling of RNF126 with IA-rhodamine (50 nM) for 1 h. Gels were visualized by in-gel fluorescence and protein loading was assessed by silver staining. **(f)** Site of modification analysis of EST1060 with pure RNF126 protein. RNF126 protein (24 μg) was incubated with EST1060 (50 μM) for 30 min and tryptic digests were analyzed by LC-MS/MS. **(g)** Structure of presumed adduct of EST1060 with C32 of RNF126. Gels and blots in **(b, d, e)** are representative images from n=3 biologically independent replicates/group.

We next appended this fumarate module onto Palbociclib to generate EST1089 **(Figure S3a)**. EST1089 maintained binding to RNF126 and was able capable of degrading CDK4 **(Figure S3b-S3c)**, suggesting that this moiety may be a more versatile chemical handle when compared to our original one.

### Identifying the Minimal Covalent Chemical Handle Required for RNF126 Interactions

Given the substantially improved labeling of RNF126 by EST1060 compared to EST1027, we next sought to understand the minimal chemical functionality and pharmacophore necessary to covalently interact with RNF126. To achieve this, we reverse engineered the EST1027 and EST1060 structures by taking their respective covalent handles and iteratively appending increasing portions of the Ribociclib scaffold and assessing their labeling of RNF126 by gel-based ABPP **(Figure 4a)**. With trifluoromethylphenyl cinnamic acid appended to the minimal diethylamine moiety, KN1026, no RNF126 labeling was observed. In contrast, the fumarate handle linked to the diethylamine moiety, KN1025, showed labeling of RNF126, albeit weaker than EST1060 **(Figure 4a)**. While appending a piperazine moiety onto trifluoromethylphenyl cinnamic acid to form EST1102 still did not confer labeling of RNF126, appending this substituent to the methoxyphenyl fumarate handle yielded JP-2-196 which showed significant potency comparable to EST1060 **(Figure 4a)**. With the trifluoromethylphenyl cinnamamide handle, installing phenylpyridine or pyridinylpiperazine substituents, KN1023 or KN1017, respectively, started to show binding to RNF126 with comparable potency to that observed with EST1027. However, this required linking a substantial portion of the Ribociclib structure as exemplified by JP-2-200 **(Figure 4a)**. In contrast, growing substituents on the fumarate handle with equivalent moieties, KN1021, KN1018, and JP-2-199, did not substantially improve potency against RNF126 beyond that that was observed with the piperazine substituent alone of JP-2-196 **(Figure 4a)**. These data collectively showed that the covalent fumarate-derived motif is a better ligand for RNF126 and the *p*-methoxyphenylpiperazinyl fumarate JP-2-196 handle is the best minimal unit identified so far for covalently engaging RNF126.

**Figure 4.**
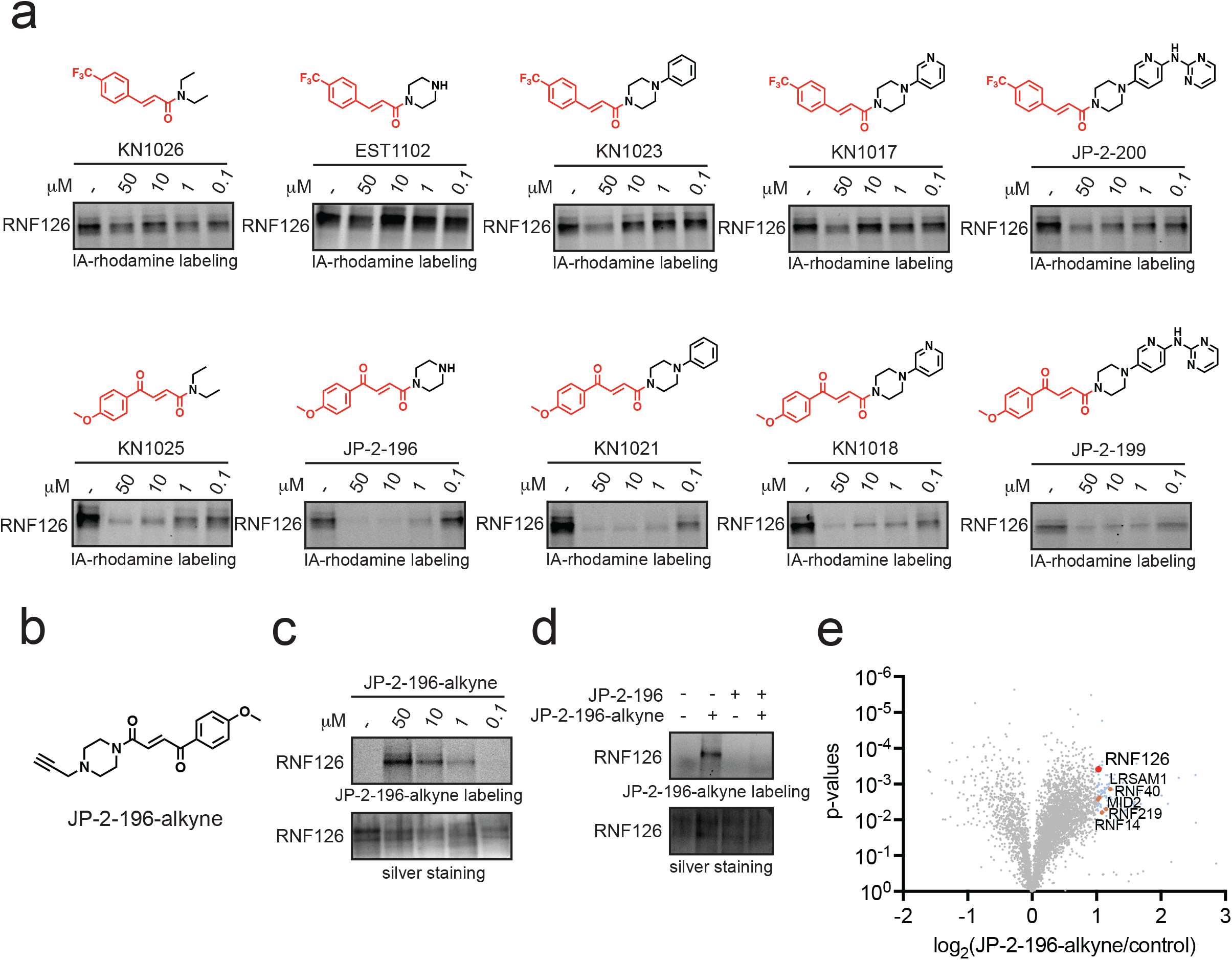
Identifying the Minimal Covalent Chemical Handle Required for RNF126 Interactions. **(a)** Gel-based ABPP of covalent chemical handles against RNF126. Recombinant RNF126 was pre-incubated with DMSO vehicle or compounds for 30 min prior to labeling of RNF126 with IA-rhodamine (50 nM) for 1 h. Gels were visualized by in-gel fluorescence. Gels are representative images from n=3 biologically independent replicates/group. **(b)** Structure of alkyne-functionalized probe of JP-2-196, JP-2-196-alkyne. **(c)** JP-2-196 labeling of pure RNF126 protein. RNF126 was labeled with DMSO vehicle or JP-2-196-alkyne for 30 min. Probe-modified RNF126 was subjected to CuAAC with a rhodamine-functionalized azide handle and visualized by SDS/PAGE and in-gel fluorescence. **(d)** Competition of JP-2-196-alkyne labeling of RNF126 by JP-2-196. RNF126 pure protein was pre-incubated with JP-2-196 (50 μM) for 30 min at 37 °C prior to JP-2-196 labeling (50 μM) for 30 min at room temperature. Probe-modified RNF126 was subjected to CuAAC with a rhodamine-functionalized azide handle and visualized by SDS/PAGE and in-gel fluorescence. **(e)** JP-2-196 pulldown at proteomics showing significant and moderately selective engagement of RNF126 with less significant engagement of 5 additional E3 ubiquitin ligases LRSAM1, RNF40, MID2, RNF219, and RNF14. HEK293T cells were treated with DMSO vehicle or JP-2-196-alkyne (10 μM) for 6 h. Subsequent lysates were subjected to copper-catalyzed azide-alkyne cycloaddition (CuAAC) with an azide-functionalized biotin handle, after which probe-modified proteins were avidin-enriched, eluted, and digested, and analyzed by TMT-based quantitative proteomics. Data shown are ratio of JP-2-196-alkyne vs DMSO control enriched proteins and p-values from n=3 biologically independent replicates/group.

To assess whether this JP-2-196 scaffold still covalently engaged RNF126 both *in vitro* and in cells and whether additional E3 ligases might be engaged by this chemical handle, we synthesized an alkyne-functionalized probe of JP-2-196—JP-2-196-alkyne **(Figure 4b)**. We first demonstrated covalent and dose-responsive labeling of pure RNF126 protein with JP-2-196 by gel-based ABPP and that this labeling was attenuated upon pre-treatment with JP-2-196 **(Figure 4c, 4d)**. To assess cellular target engagement and overall selectivity of this JP-2-196-alkyne probe, we next treated HEK293T cells with either the JP-2-196-alkyne probe or vehicle and subsequently appended an azide-functionalized biotin enrichment handle through copper-catalyzed **“**click-chemistry**”** followed by avidin-enrichment of probe modified peptides to assess probe-enriched proteins by quantitative proteomics **(Figure 4e; Table S2)**. We identified 23 distinct protein targets that were significantly (p<0.001) enriched by the JP-2-196-alkyne probe over vehicle control by >2-fold of which RNF126 was the only E3 ligase among these 23 targets **(Figure 4e)**. Using a less stringent filter, an additional 87 proteins were significantly (p<0.01) enriched by the JP-2-196-alkyne probe by >2-fold, which included 5 additional E3 ligases—RNF40, MID2, RNF219, and RNF14 **(Figure 4e)**. Thus, RNF126 is the most significantly enriched E3 ligase by the JP-2-196-alkyne probe, but there may be additional E3 ligases engaged by this chemical handle.

### Transplanting Fumarate-Based Covalent Chemical Handle onto Protein-Targeting Ligands that Already Possess Piperazines or Morpholines at the Exit Vector

Having identified a minimal motif required to potently engage RNF126, we next sought to transplant this handle onto other protein-targeting ligands to test if this module could be widely used to degrade their respective protein targets. We focused on ligands that like ribociclib already possessed piperazine moieties at the solvent exposed exit vector. First, we incorporated the fumarate motif onto another clinically approved kinase inhibitor dasatinib that inhibits the fusion oncogene BCR-ABL and the parent kinase c-ABL in leukemia, synthesizing JP-2-227 **(Figure 5a)**. This degrader showed potent labeling of pure RNF126 protein and robustly degraded both BCR-ABL and c-ABL in K562 leukemia cancer cells in a dose-responsive manner **(Figure 5b-5d)**. Previous BCR-ABL and c-ABL PROTACs bearing cereblon, VHL, or RNF114 recruiters demonstrated either preferential degradation of c-ABL compared to BCR-ABL, or showed incomplete degradation of BCR-ABL ^29–31^. Here, we showed robust degradation of both BCR-ABL and c-ABL with near-complete loss of BCR-ABL **(Figure 5c-5d)**.

**Figure 5.**
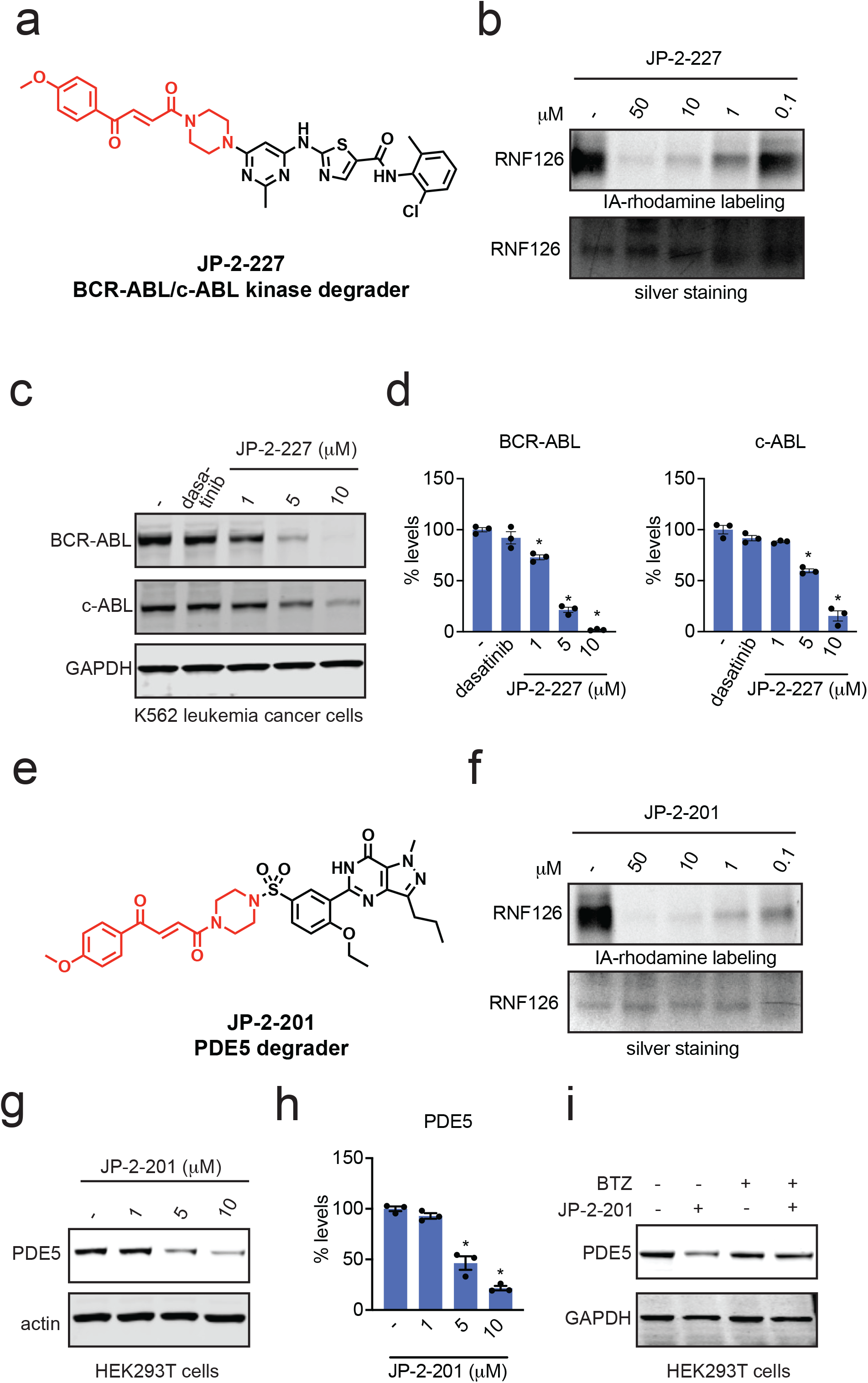
Transplanting Covalent Chemical Handle onto Protein-Targeting Ligands that Already Possess Piperazines at the Exit Vector. **(a)** Structure of JP-2-227 with the optimized covalent handle shown in red that was appended onto the BCR-ABL and c-ABL kinase inhibitor Dasatinib. **(b)** Gel-based ABPP analysis of JP-2-227 against RNF126. Recombinant RNF126 was pre-incubated with DMSO vehicle or JP-2-227 for 30 min prior to labeling of RNF126 with IA-rhodamine (50 nM) for 1 h. Gels were visualized by in-gel fluorescence and protein loading was assessed by silver staining. **(c)** JP-2-227 degrades BCR-ABL and c-ABL in K562 leukemia cancer cells. K562 cells were treated with DMSO vehicle or JP-2-227 for 24 h and BCR-ABL, c-ABL, and loading control GAPDH levels were assessed by Western blotting. **(d)** Quantification of experiment in **(c). (e)** Structure of JP-2-201 with the optimized covalent handle shown in red that was appended onto the PDE5 inhibitor Sildenafil. **(f)** Gel-based ABPP analysis of JP-2-201 against RNF126. Recombinant RNF126 was pre-incubated with DMSO vehicle or JP-2-201 for 30 min prior to labeling of RNF126 with IA-rhodamine (50 nM) for 1 h. Gels were visualized by in-gel fluorescence and protein loading was assessed by silver staining. **(g)** JP-2-201 degrades PDE5 in HEK293T cells. HEK293T cells were treated with DMSO vehicle or JP-2-201 for 24 h and PDE5 and loading control actin levels were assessed by Western blotting. **(h)** Quantification of experiment in **(g). (i)** JP-2-201-mediated degradation of PDE5 is proteasome-dependent. HEK293T cells were pre-treated with DMSO vehicle or bortezomib (1 μM) for 1 h prior to treating cells with DMSO or JP-2-201 (10 μM). PDE5 and loading control GAPDH levels were assessed by Western blotting. Blots and gels shown in **(b, c, f, g, i**) are representative images from n=3 biological replicates. Bar graphs in **(d, h)** show individual replicate values and average ± sem. Statistical significance is calculated as *p<0.05 compared to DMSO vehicle in **(d, h)**.

Next, we incorporated the fumarate handle onto a ligand from a target outside of the kinase family with the clinically approved phosphodiesterase 5 (PDE5) inhibitor Sildenafil, that already possessed a methylpiperazine within its core chemical scaffold, to generate JP-2-201 **(Figure 5e)**. JP-2-201 showed labeling of pure RNF126 protein and indeed demonstrated degradation of PDE5 in HEK293T cells in a dose-responsive and proteasome-dependent manner **(Figure 5f-5i)**. We next appended our fumarate handle onto the SMARCA2-bromodain ligand 1 used in a previously developed SMARCA2 PROTAC ABCI1 developed by the Ciulli group that also bears a piperazine at the exit vector, to generate JP-2-249 **(Figure 6a)** ^32^. Again, this degrader showed potent labeling of RNF126 and degraded SMARCA2 with near complete loss of protein at 10 μM and still significant loss at 1 μM in MV-4-11 leukemia cancer cells. **(Figure 6b-6d)**. Finally, we appended the fumarate handle onto the LRRK2 inhibitor HG-10-102-01, which is currently under evaluation for Parkinson**’**s disease that bears a morpholine exit vector, to generate JP-2-244 **(Figure 6e)**. This compound also potently labeled RNF126 and led to LRRK2 loss in A549 lung cancer cells **(Figure 6f-6h)**.

**Figure 6.**
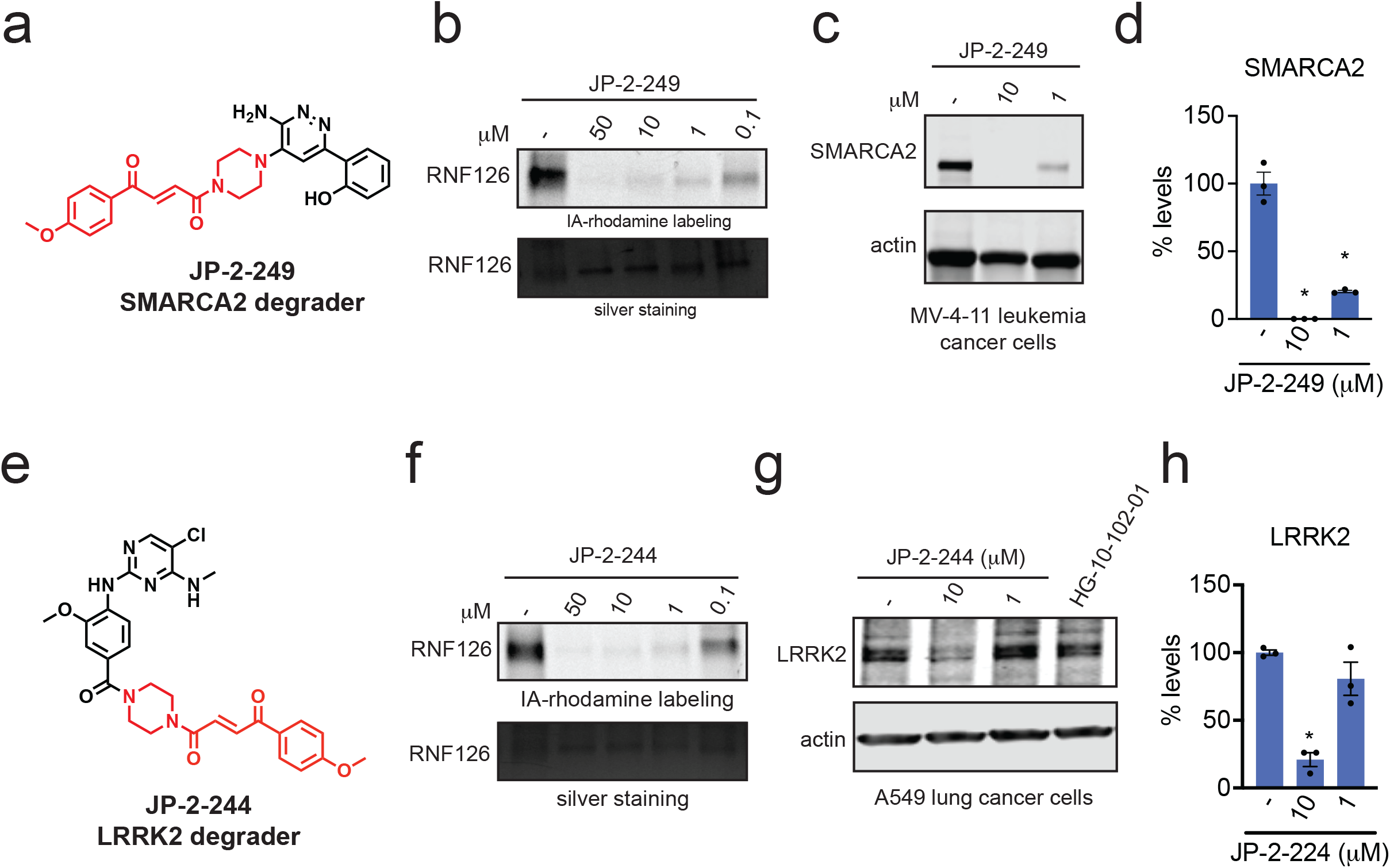
Transplanting Covalent Chemical Handle onto Additional Protein-Targeting Ligands that Already Possess Piperazines or Morpholines at the Exit Vector. **(a)** Structure of SMARCA2 degrader JP-2-249 consisting of the optimized covalent handle incorporated into a previously reported protein-targeting ligand for SMARCA2. **(b)** Gel-based ABPP analysis of JP2-249 against RNF126. Recombinant RNF126 was pre-incubated with DMSO vehicle or JP-2-249 for 30 min prior to labeling of RNF126 with IA-rhodamine (50 nM) for 1 h. Gels were visualized by in-gel fluorescence and protein loading was assessed by silver staining. **(c)** MV-4-11 leukemia cancer cells were treated with DMSO vehicle or JP2-2-49 for 24 h and SMARCA2 and actin loading control levels were assessed by Western blotting. **(d)** Quantitation of experiment from **(c). (e)** Structure of LRRK2 degrader JP-2-244 consisting of the optimized covalent handle incorporated into a previously reported LRRK2 inhibitor. **(f)** Gel-based ABPP analysis of JP2-244 against RNF126. Recombinant RNF126 was pre-incubated with DMSO vehicle or JP-2-244 for 30 min prior to labeling of RNF126 with IA-rhodamine (50 nM) for 1 h. Gels were visualized by in-gel fluorescence and protein loading was assessed by silver staining. **(g)** A549 lung cancer cells were treated with DMSO vehicle or JP2-2-44 for 24 h and LRRK2 and actin loading control levels were assessed by Western blotting. **(h)** Quantitation of experiment from **(g)**. Blots and gels shown in **(b, c, f, g**) are representative images from n=3 biologically independent replicates. Bar graphs in **(d, h)** show individual replicate values and average ± sem. Statistical significance is calculated as *p<0.05 compared to DMSO vehicle in **(d, h)**.

Collectively, our data demonstrated that this fumarate chemical handle could be used to extend several protein-targeting ligands already bearing piperazine exit vectors to degrade their respective targets across several protein classes from kinases, phosphodiesterases, and bromodomains.

### Transplanting Covalent Chemical Handle onto Protein-Targeting Ligands from Unrelated Chemical and Protein Classes

Having shown degradation of additional kinase and non-kinase targets with our fumarate chemical module when appended onto protein-targeting ligands already bearing a piperazine moiety at the exit vector, we next sought to transplant this handle onto ligands from unrelated chemical and protein classes. We first incorporated the JP-2-196 handle onto the BET family bromodomain inhibitor JQ1 which does not already have a piperazine moiety within its core structure, forming JP-2-197 **(Figure 7a)**. This compound still potently labeled pure RNF126 protein and led to highly potent mid-nanomolar degradation of both long and short isoforms of BRD4 in HEK293T cells in a dose-responsive and time-responsive manner **(Figure 7b-7d; Figure S4a)**. We also observed modest attenuation of BRD4 degradation at higher concentrations, indicating potential **“**hook effects.**”** Like our observation of RNF126 loss with EST1027, we also observed RNF126 loss at the highest concentration of JP-2-197 **(Figure 7c-7d)**. We also further confirmed that this BRD4 degradation by JP-2-197 was proteasome-dependent **(Figure 7e-7f)**. Pretreatment of cells with excess JQ1 also completely attenuated JP-2-197-mediated BRD4 degradation **(Figure S4b)**. Quantitative proteomic profiling of JP-2-197 in HEK293T cells also demonstrated relatively selective degradation of BRD4 with 3 other targets—DDX5, RBM28, and UTP20—also showing reduced protein levels **(Figure 7g; Table S3)**. A non-reactive derivative of JP-2-197, JP-2-232, still showed binding to RNF126, albeit less strongly and potently compared to JP-2-197 **(Figure S4c-S4d)**, once again indicating modest non-covalent binding affinity of the fumarate motif to RNF126. However, as observed with CDK4, this non-reactive analog did not degrade BRD4 in HEK293T cells **(Figure S4e)**. We also demonstrated that the chemical handle JP-2-196 itself does not alter BRD4 levels compared to the BRD4 degrader JP-2-197 **(Figure S4f)**. We also generated a derivative of JP-2-197 that replaced its piperazine moiety with an ethyl diamine linker, JP-2-219 **(Figure S5)**. JP-2-219 was still able to degrade BRD4, but substantially less potently compared to JP-2-197, again demonstrating tunable SAR for these covalent glue degraders **(Figure S5)**.

**Figure 7.**
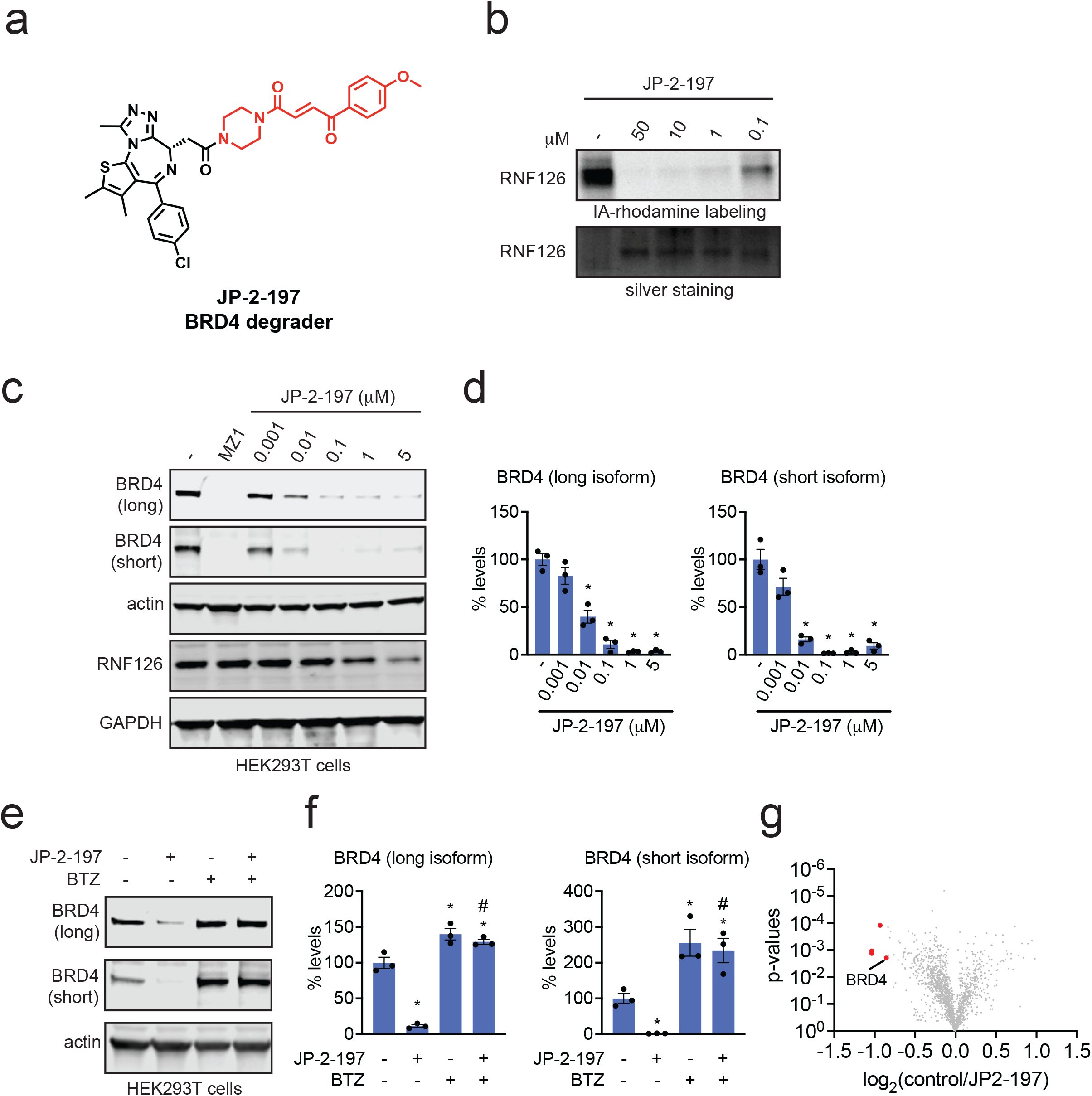
Transplanting Covalent Chemical Handle onto BET Family Inhibitor JQ1 to Degrade BRD4. **(a)** Structure of JP-2-197 with the optimized covalent handle shown in red that was appended onto the BET family bromodomain inhibitor JQ1. **(b)** Gel-based ABPP analysis of JP-2-197 against RNF126. Recombinant RNF126 was pre-incubated with DMSO vehicle or JP-2-197 for 30 min prior to labeling of RNF126 with IA-rhodamine (50 nM) for 1 h. Gels were visualized by in-gel fluorescence and protein loading was assessed by silver staining. **(c)** JP-2-197 degrades BRD4 in HEK293T cells. HEK293T cells were treated with DMSO vehicle, positive control BRD4 PROTAC MZ1 (1 μM), or JP-2-197 for 24 h. BRD4 long and short isoforms, RNF126, and loading controls actin and GAPDH were assessed by Western blotting. **(d)** Quantification of experiment described in **(c). (e)** Proteasome-dependent degradation of BRD4 by JP-2-197. HEK293T cells were pre-treated with DMSO vehicle or the proteasome inhibitor BTZ (10 μM) for 1 h prior to treatment of cells with DMSO vehicle or JP-2-197 (1 μM) and BRD4 and loading control actin levels were assessed by Western blotting. **(f)** Quantification of the experiment described in **(e). (g)** TMT-based quantitative proteomic profiling of JP-2-197 in HEK293T cells. HEK293T cells were treated with DMSO vehicle or JP-2-197 (1 μM) for 24 h. Data are from n=3 biological replicates per group. Blots and gels shown in **(b, c, e**) are representative images from n=3 biologically independent replicates. Bar graphs in **(d, f)** show individual replicate values and average ± sem. Statistical significance is calculated as *p<0.05 compared to DMSO vehicle in **(d, f)** and #p<0.05 compared to JP-2-197-treated group in **(f)**.

BRD4 is a protein which is relatively easy to degrade, and has been used as a test case for many different types of degraders ^23,28,33,34^. Thus, we next sought to expand the scope of ligand and target classes to understand the diversity of targets that our chemical handle could access for targeted protein degradation applications. First, we incorporated our fumarate handle into the BTK inhibitor Ibrutinib, replacing the BTK C481-targeting cysteine-reactive acrylamide warhead to generate JP-2-247 **(Figure 8a)**. Interestingly, this molecule still potently labeled RNF126 but also showed BTK degradation in MINO lymphoma cancer cells **(Figure 8b-8d)**.

**Figure 8.**
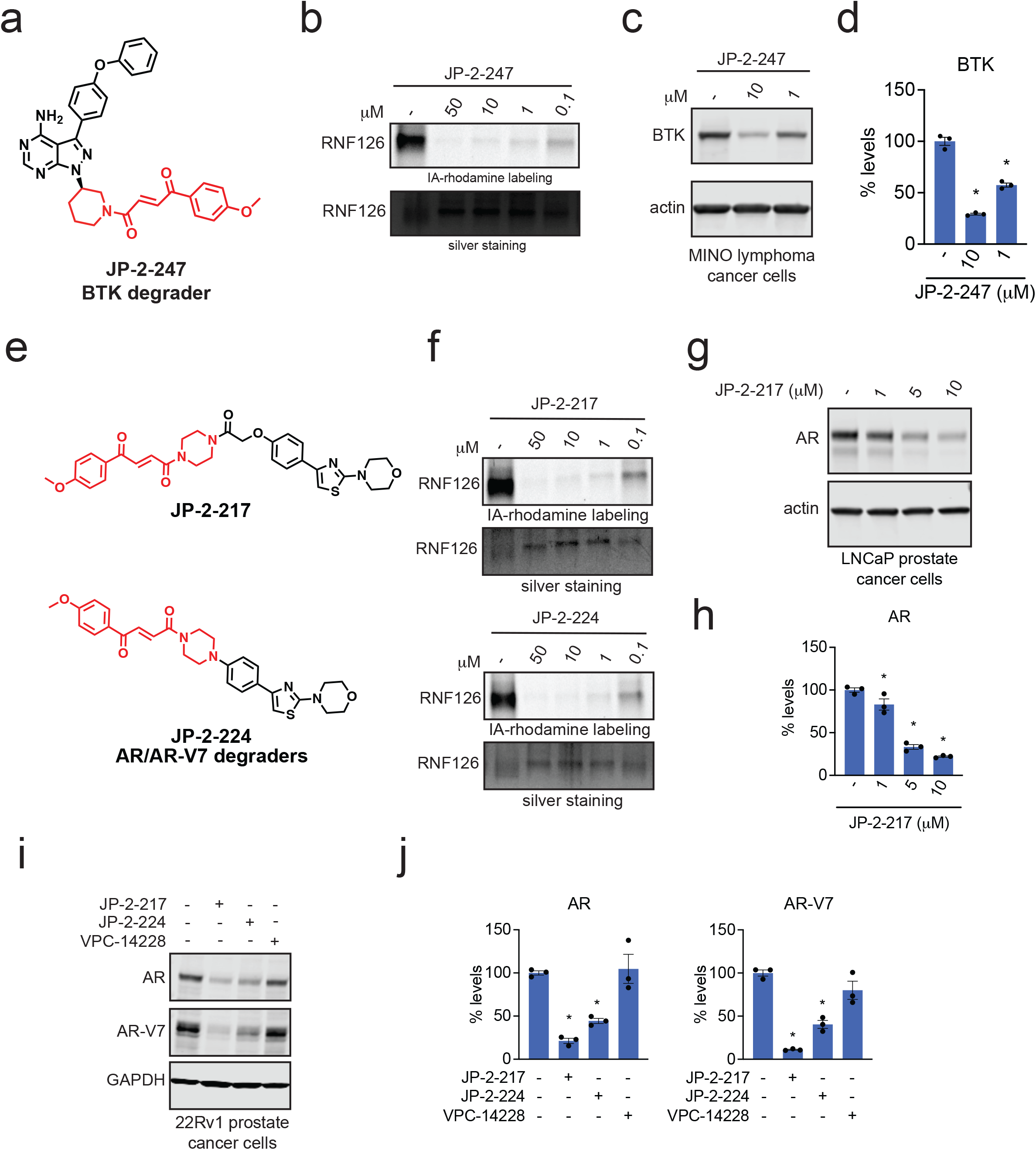
Transplanting Covalent Chemical Handle onto BTK Inhibitor and Androgen Receptor DNA Binding Domain-Targeting Ligand to Degrade BTK and AR and AR-V7. **(a)** Structure of a BTK degrader JP-2-247 consisting of the fumarate handle incorporated into the BTK inhibitor ibrutinib. **(b)** Gel-based ABPP analysis of JP-2-247 against RNF126. Recombinant RNF126 was pre-incubated with DMSO vehicle or JP-2-247 for 30 min prior to labeling of RNF126 with IA-rhodamine (50 nM) for 1 h. Gels were visualized by in-gel fluorescence and protein loading was assessed by silver staining. **(c)** MINO lymphoma cancer cells were treated with JP-2-247 for 24 h and BTK and loading control actin levels were detected by Western blotting. **(d)** Quantitation of the data shown in **(c). (e)** Structures of two AR-V7 degraders consisting of the fumarate handle linked to an AR DNA-binding domain ligand VPC-14228—JP-2-217 and JP-2-224. **(f)** Gel-based ABPP analysis of JP-2-217 and JP-2-224 against RNF126. Recombinant RNF126 was pre-incubated with DMSO vehicle or JP-2-217 or JP-2-224 for 30 min prior to labeling of RNF126 with IA-rhodamine (50 nM) for 1 h. Gels were visualized by in-gel fluorescence and protein loading was assessed by silver staining. **(g)** LNCaP prostate cancer cells were treated with JP-2-217 for 24 h and AR and loading control actin levels were detected by Western blotting. **(h)** Quantitation of the data shown in **(g). (i)** 22Rv1 prostate cancer cells were treated with DMSO vehicle, JP-2-217, JP-2-224, or VPC-14228 (10 μM) for 24 h and AR and AR-V7 and GAPDH loading control levels were assessed by Western blotting. **(j)** Quantitation of experiment from **(i)**. Blots and gels shown in **(b, c, f, g, i**) are representative images from n=3 biologically independent replicates. Bar graphs in **(d, h, j)** show individual replicate values and average ± sem. Statistical significance is calculated as *p<0.05 compared to DMSO vehicle in **(d, h, j)**.

We next tackled a more challenging target, the truncated and constitutively active mutant of androgen receptor (AR), AR-V7, that drives the pathogenesis of androgen-resistant prostate cancers ^35^. AR-V7 is considered to be a relatively undruggable target, given that the ligand binding domain that is the target of most AR-targeting drugs is missing from AR-V7. We linked our fumarate derivative JP-2-196 onto a previously discovered DNA-binding domain ligand, VPC-14228, for the androgen receptor that had recently been used in several VHL-based PROTACs to degrade AR-V7, through two different types of linkages to yield JP-2-217 and JP-2-224 **(Figure 8e)** ^36–38^. Both of these compounds showed potent binding to pure RNF126 protein **(Figure 8f)**. Given that VPC-14228 binds to the DNA binding domain shared between wild-type full-length AR as well as its truncation mutants, we first tested this degrader for wild-type AR degradation in AR-sensitive LNCaP prostate cancer cells **(Figure 8g)**. JP-2-217 degraded AR in LNCaP cells in a dose-responsive manner **(Figure 8g-8h)**. We next tested both degraders in the androgen-resistant prostate cancer cell line 22Rv1 that expresses wild-type AR and AR-V7 and demonstrated that both JP-2-217 and JP-2-224, but not VPC-14228 or a previously reported VHL-based AR-V7 PROTAC (compound 6)^38^, degraded both wild-type AR and AR-V7 **(Figure 8i-8j; Figure S6a-S6d)**. Overall, we demonstrated that the minimal covalent handle JP-2-196 could be used to convert protein targeting ligands into molecular glue degraders of several proteins from different protein classes.

Overall, we demonstrate the utility and versatility of JP-2-196, a low molecular weight covalent fumarate derivative, in enabling the degradation of a wide range of proteins using a diverse class of protein-targeting ligands.

## Discussion

While our chemical handle likely still requires significant optimization to improve potency, selectivity, and its versatility as a general motif for degrader design, we demonstrated proof-of-concept to convert protein-targeting ligands in a more rational manner into monovalent degraders of their targets without the need for long linkers and resultant high molecular weight PROTACs. We demonstrate the utility of our simple covalent derivative appended to ligands against several targets, including CDK4, BCR-ABL and c-ABL, PDE5, BRD4, AR and AR-V7, BTK, SMARCA2, and LRRK2 to induce degradation. We note that the molecular weights of all our degraders are lower than traditional PROTAC molecules which may ease the burden of future medicinal chemistry efforts to generate compounds that show optimal pharmacokinetic parameters.

In our study, we initially used the CDK4/6 inhibitor Ribociclib as a testbed, where we discovered a covalent chemical handle that, when appended to the solvent exposed end of Ribociclib, induced CDK4 degradation. Using chemoproteomic platforms, we discovered that this molecule covalently targeted C32 in the E3 ligase RNF126. We demonstrated that this CDK4 degradation was proteasome-dependent and was at least in-part driven through RNF126. Structure-activity relationship studies of our initial hit structure led to further optimization of both the chemical handle and CDK4 degradation. Through further analysis and optimization of the covalent chemical handle for RNF126 recognition, we found that the piperazine moiety that was part of the Ribociclib core structure was also necessary for RNF126 recognition, yielding the minimal covalent fumarate derivative JP-2-196. We demonstrated that this covalent motif still engaged RNF126 in cells with a moderate degree of selectivity, with potentially additional E3 ligases that were less significantly engaged. We showed that this fumarate-based covalent module could be transplanted onto several other protein-targeting ligands across diverse chemical and protein classes to convert these ligands into degraders of their targets.

There are still many open questions that we hope to address in future studies. These questions include further understanding the mechanism underlying the versatile degradation observed with our chemical motif once appended to numerous ligands across so many different protein and ligand classes and the contribution of RNF126 and the role of BAG6 in RNF126-mediated responses, and the contribution of potentially additional components of the ubiquitin-proteasome system beyond RNF126. We further hope to understand the structural and biochemical underpinnings and flexibility of the ternary complex formed between RNF126, our molecular glue degraders, and the diversity of targets degraded in this study. We would also like to better understand how targeting a zinc-coordinating cysteine within RNF126 enables RNF126 recruitment to neo-substrates to ubiquitinate and degrade their respective targets without disrupting RNF126 structure and function, and whether targeting this site may influence RNF126 interactions with BAG6. We would also like to determine in future studies whether we are disrupting RNF126 endogenous function and whether this may have any toxicity. While our current hypothesis is the direct recruitment of RNF126 to the target, there are other scenarios that would explain our data such as a fumarate-induced unfolding and destabilization of target proteins that are then resolved in a RNF126-dependent manner as part of its role in protein quality control for hydrophobic substrates ^39–41^. Additional areas of further investigation will be to further optimize the potency and selectivity and ultimately further reduce the molecular weight footprint of the chemical handle that can be transplanted onto various protein-targeting ligands.

Another major question is whether we can further expand the strategies that we have employed here to identify chemical handles against other E3 ligases beyond RNF126 that can be used to rationally develop molecular glue degraders. Given previous successes in using covalent ligands and chemoproteomic platforms to develop novel E3 ligase recruiters for PROTAC applications, including RNF114, RNF4, DCAF16, DCAF11, FEM1B, and DCAF1 and the high degree of E3 ligase ligandability using covalent strategies, it would be of future interest to further expand on the efforts started here with an expanded library of covalent moieties that could potentially access additional E3 ligases ^23,28,34,42–46^.

Overall, our study identifies a potential starting point for developing chemical rational design principles for converting protein-targeting ligands into monovalent molecular glue degraders through appending a minimal linker-less covalent handle that can recruit RNF126, and potentially additional E3 ligases. We recognize that with the optimization of the RNF126-recognizing ligand, our resulting covalent handle linked to protein-targeting ligands could also be considered linker-less PROTACs, but we anticipate that as molecular glue and PROTAC design evolves, the definition between PROTACs and molecular glue degraders will likely start to merge, and that our study represents a step towards that direction.

## Acknowledgement

We thank the members of the Nomura Research Group and Novartis Institutes for BioMedical Research for critical reading of the manuscript. This work was supported by Novartis Institutes for BioMedical Research and the Novartis-Berkeley Translational Chemical Biology Institute (NB-TCBI) for all listed authors. This work was also supported by the Nomura Research Group and the Mark Foundation for Cancer Research ASPIRE Award for DKN, EST, JP, and KN. JP was also supported by the Mark Foundation for Cancer Research Momentum Postdoctoral Award and the Canadian NSERC CRSNG Postdoctoral Fellowship. This work was also supported by grants from the National Institutes of Health (R01CA240981 and R35CA263814 for DKN) and the National Science Foundation Molecular Foundations for Biotechnology Award (2127788). We also thank Drs. Hasan Celik, Alicia Lund, and UC Berkeley**’**s NMR facility in the College of Chemistry (CoC-NMR) for spectroscopic assistance. Instruments in the College of Chemistry NMR facility are supported in part by NIH S10OD024998.

## Author Contributions

EST, JP, DKN conceived of the project idea, designed experiments, performed experiments, analyzed and interpreted the data, and wrote the paper. EST, JP, DKN, KN performed experiments, analyzed and interpreted data, and provided intellectual contributions. DD, LM, MJH, JAT, JMK, MS provided intellectual contributions to the project and overall design of the project.

## Online Methods

### Cell Culture

C33A cells were purchased from American Type Culture Collection (ATCC) and were cultured in Dulbecco**’**s Modified Eagle Medium (DMEM) containing 10% (v/v) fetal bovine serum (FBS) and maintained at 37 °C with 5% CO_2_. 22RV1 cells were purchased from the ATCC and were cultured in RPMI-1640 Medium containing 10% (v/v) FBS and maintained at 37 °C with 5% CO_2_. HEK293T cells were obtained from the UC Berkeley Cell Culture Facility and were cultured in DMEM containing 10% (v/v) FBS and maintained at 37 °C with 5% CO_2_. K562 cells were obtained from the UC Berkeley Cell Culture Facility and were cultured in Iscove**’**s Modified Dulbecco**’**s Medium (IMDM) containing 10% (v/v) FBS and maintained at 37 °C with 5% CO_2_. MV-4-11 cells were obtained from the ATCC and were cultured in IMDM containing 10% (v/v) FBS and maintained at 37 °C with 5% CO_2_. A549 cells were obtained from the ATCC and were cultured in F-12K Medium containing 10% (v/v) FBS and maintained at 37 °C with 5% CO_2_. Mino cells were obtained from the ATCC and were cultured in RPMI-1640 Medium containing 10% (v/v) FBS and maintained at 37 °C with 5% CO_2_. LNCaP cells were obtained from the UC Berkeley Cell Culture Facility and were cultured in RPMI-1640 Medium containing 10% (v/v) FBS and maintained at 37 °C with 5% CO2. Unless otherwise specified, all cell culture materials were purchased from Gibco. It is not known whether HEK293T cells are from male or female origin.

### Preparation of Cell Lysates

Cells were washed twice with cold PBS, scraped, and pelleted by centrifugation (700 g, 5 min, 4 °C). Pellets were resuspended in PBS, sonicated, clarified by centrifugation (12,000 g, 10 min, 4 °C), and lysate was transferred to new low-adhesion microcentrifuge tubes. Proteome concentrations were determined using BCA assay and lysate was diluted to appropriate working concentrations.

### Western Blotting

Proteins were resolved by SDS/PAGE and transferred to nitrocellulose membranes using the Trans-Blot Turbo transfer system (Bio-Rad). Membranes were blocked with 5% BSA in Tris-buffered saline containing Tween 20 (TBS-T) solution for 30 min at RT, washed in TBS-T, and probed with primary antibody diluted in recommended diluent per manufacturer overnight at 4 °C. After 3 washes with TBS-T, the membranes were incubated in the dark with IR680- or IR800-conjugated secondary antibodies at 1:10,000 dilution in 5 % BSA in TBS-T at RT for 1 h. After 3 additional washes with TBST, blots were visualized using an Odyssey Li-Cor fluorescent scanner. The membranes were stripped using ReBlot Plus Strong Antibody Stripping Solution (EMD Millipore) when additional primary antibody incubations were performed. Antibodies used in this study were CDK4 (Abcam ab108357), Vinculin (Abcam ab129002), GAPDH (Cell Signaling Technology 14C10), RNF126 (Santa Cruz Biotechnology sc-376005), BRD4 (Abcam ab128874), Beta Actin (Cell Signaling Technology 13E5), PDE5 (Abcam ab259945), AR-V7 (Abcam ab273500), c-Abl (Santa Cruz Biotechnology sc-23), SMARCA2 (Abcam ab240648), LRRK2 (Abcam ab133474), BTK (Cell Signaling Technology D3H5), Androgen Receptor (Cell Signaling Technology D6F11).

### Expression and purification of recombinant RNF126 protein

RNF126 mammalian expression plasmid with a C-terminal FLAG tag was purchased from Origene (Origene Technologies Inc., RC204986). The plasmid was transformed into NEB 5-alpha Competent E. coli (DH5**α**) cells (NEB product no. C2987H). The following day, a single transformed colony was used to inoculate 50 ml of nutrient rich LB medium containing kanamycin (50 μg/ml) and was incubated at 37 °C overnight, with agitation (250 rpm). A Miniprep (Qiagen) kit was used to isolate the plasmid before sequence verification with appropriate primers.

HEK293T cells were grown to 30-50% confluency in DMEM supplemented with 10% FBS (Corning) and maintained at 37 °C with 5% CO2. Immediately before transfection, media was replaced with DMEM containing 5% FBS. Each plate was transfected with 24 μg of overexpression plasmid with 24 μL Lipofectamine 3000 (Invitrogen) in Opti-MEM. After 48 h cells were collected in PBS, lysed by sonication, and batch bound with anti-DYKDDDDK resin (GenScript, L00432) for 2 hours. Lysate and resin were washed with PBS and eluted with 133.33 μg/ml 3XFLAG peptide (ApexBio, A6001) in PBS. Five elutions were performed for 15 minutes each. Elutions were concentrated and the protein was stored in PBS. Concentration and purity was determined using the BCA assay and western blotting.

### IsoTOP-ABPP Chemoproteomic Experiments

IsoTOP-ABPP studies were done as previously reported ^21,47,48^. All of the isoTOP-ABPP datasets were prepared as described below using the IA-alkyne probe. Cells were lysed by probe sonication in PBS and protein concentrations were measured by BCA assay. Cells were treated for 2 h with either DMSO vehicle or compound before cell collection and lysis. Proteomes were subsequently labeled with IA-alkyne labeling (200 μM) for 1 h at room temperature. CuAAC was used by sequential addition of tris(2-carboxyethyl)phosphine (1 mM, Strem, 15-7400), tris[(1-benzyl-1H-1,2,3-triazol-4-yl)methyl]amine (34 μM, Sigma, 678937), copper(II) sulfate (1 mM, Sigma, 451657) and biotin-linker-azide—the linker functionalized with a tobacco etch virus(TEV) protease recognition sequence as well as an isotopically light or heavy valine for treatment of control or treated proteome, respectively. After CuAAC, proteomes were precipitated by centrifugation at 6,500*g*, washed in ice-cold methanol, combined in a 1:1 control:treated ratio, washed again, then denatured and resolubilized by heating in 1.2% SDS–PBS to 90 °C for 5 min. Insoluble components were precipitated by centrifugation at 6,500*g* and soluble proteome was diluted in 5 ml 0.2% SDS–PBS. Labeled proteins were bound to streptavidin-agarose beads (170 μl resuspended beads per sample, Thermo Fisher, 20349) while rotating overnight at 4 °C. Bead-linked proteins were enriched by washing three times each in PBS and water, then resuspended in 6 M urea/PBS, reduced in DTT (9.26 mM, ThermoFisher, R0861), and alkylated with iodoacetamide (18 mM, Sigma, I6125), before being washed and resuspended in 2 M urea/PBS and trypsinized overnight with 0.5 μg /μL sequencing grade trypsin (Promega, V5111). Tryptic peptides were eluted off. Beads were washed three times each in PBS and water, washed in TEV buffer solution (water, TEV buffer, 100 μM dithiothreitol) and resuspended in buffer with Ac-TEV protease (Invitrogen, 12575-015) and incubated overnight. Peptides were diluted in water and acidified with formic acid (1.2 M, Fisher, A117-50) and prepared for analysis.

### IsoTOP-ABPP Mass Spectrometry Analysis

Peptides from all chemoproteomic experiments were pressure-loaded onto a 250 μm inner diameter fused silica capillary tubing packed with 4 cm of Aqua C18 reverse-phase resin (Phenomenex, 04A-4299), which was previously equilibrated on an Agilent 600 series high-performance liquid chromatograph using the gradient from 100% buffer A to 100% buffer B over 10 min, followed by a 5 min wash with 100% buffer B and a 5 min wash with 100% buffer A. The samples were then attached using a MicroTee PEEK 360 μm fitting (ThermoFisher Scientific p-888) to a 13 cm laser pulled column packed with 10 cm Aqua C18 reverse-phase resin and 3 cm of strong-cation exchange resin for isoTOP-ABPP studies. Samples were analyzed using an Q Exactive Plus mass spectrometer (Thermo Fisher Scientific) using a five-step Multidimensional Protein Identification Technology (MudPIT) program, using 0, 25, 50, 80 and 100% salt bumps of 500 mM aqueous ammonium acetate and using a gradient of 5–55% buffer B in buffer A (buffer A: 95:5 water:acetonitrile, 0.1% formic acid; buffer B 80:20 acetonitrile:water, 0.1% formic acid). Data were collected in data-dependent acquisition mode with dynamic exclusion enabled (60 s). One full mass spectrometry (MS1) scan (400–1,800 mass-to-charge ratio (*m/z*)) was followed by 15 MS2 scans of the *n*th most abundant ions. Heated capillary temperature was set to 200 °C and the nanospray voltage was set to 2.75 kV.

Data were extracted in the form of MS1 and MS2 files using Raw Extractor v.1.9.9.2 (Scripps Research Institute) and searched against the Uniprot human database using ProLuCID search methodology in IP2 v.3-v.5 (Integrated Proteomics Applications, Inc.) ^49^. Cysteine residues were searched with a static modification for carboxyaminomethylation (+57.02146) and up to two differential modifications for methionine oxidation and either the light or heavy TEV tags (+464.28596 or +470.29977, respectively). Peptides were required to be fully tryptic peptides and to contain the TEV modification. ProLUCID data were filtered through DTASelect to achieve a peptide false-positive rate below 5%. Only those probe-modified peptides that were evident across two out of three biological replicates were interpreted for their isotopic light to heavy ratios. For those probe-modified peptides that showed ratios greater than two, we only interpreted those targets that were present across all three biological replicates, were statistically significant and showed good quality MS1 peak shapes across all biological replicates. Light versus heavy isotopic probe-modified peptide ratios are calculated by taking the mean of the ratios of each replicate paired light versus heavy precursor abundance for all peptide-spectral matches associated with a peptide. The paired abundances were also used to calculate a paired sample *t*-test *P* value in an effort to estimate constancy in paired abundances and significance in change between treatment and control. *P* values were corrected using the Benjamini–Hochberg method.

### Gel-Based ABPP

Recombinant RNF126 (0.1μg/sample) was pre-treated with either DMSO vehicle or covalent ligand at 37 °C for 30 min in 25 μL of PBS, and subsequently treated with of IA-Rhodamine (concentrations designated in figure legends) (Setareh Biotech) at room temperature for 1 h in the dark. The reaction was stopped by addition of 4×reducing Laemmli SDS sample loading buffer (Alfa Aesar). After boiling at 95 °C for 5 min, the samples were separated on precast 4 −20% Criterion TGX gels (Bio-Rad). Probe-labeled proteins were analyzed by in-gel fluorescence using a ChemiDoc MP (Bio-Rad).

### JP-2-196-Alkyne Pulldown Quantitative Proteomics

Cells were treated with either DMSO vehicle or compound (JP-2-196-alkyne 10 μM) for 6 h. Cells were harvested and lysed by probe sonication in PBS and protein concentrations were measured by BCA assay. CuAAC was used by sequential addition of tris(2-carboxyethyl)phosphine (893 **μ**M, Strem, 15-7400), tris[(1-benzyl-1H-1,2,3-triazol-4-yl)methyl]amine (91 μM, Sigma, 678937), copper(II) sulfate (893 μM, Sigma, 451657) and biotin picolyl azide (179 μM, Sigma, 900912). After 1 h, proteomes were precipitated by centrifugation at 6,500*g*, washed in ice-cold methanol, combined to attain 10 mg per sample, washed again, then denatured and resolubilized by heating in 1.2% SDS–PBS to 90 °C for 5 min. The soluble proteome was diluted with 4 mL of PBS and labeled proteins were bound to streptavidin-agarose beads (170 μl resuspended beads per sample, Thermo Fisher, 20349) while rotating overnight at 4 °C. Bead-linked proteins were enriched by washing three times each in PBS and water, then resuspended in 6 M urea/PBS, reduced in DTT (9.26 mM, ThermoFisher, R0861), and alkylated with iodoacetamide (18 mM, Sigma, I6125), before being washed and resuspended in 50 mM Triethylammonium bicarbonate (TEAB) and trypsinized overnight with 0.5 μg /μL sequencing grade trypsin (Promega, V5111). Tryptic peptides were eluted off. Individual samples were then labeled with isobaric tags using commercially available TMTsixplex (Thermo Fisher Scientific, P/N 90061) kits, in accordance with the manufacturer**’**s protocols. Tagged samples (20 μg per sample) were combined, dried with SpeedVac, resuspended with 300 μL 0.1% TFA in H2O, and fractionated using high pH reversed-phase peptide fractionation kits (Thermo Scientific, P/N 84868) according to manufacturer**’**s protocol. Fractions were dried with SpeedVac, resuspended with 50 μL 0.1% FA in H2O, and analyzed by LC-MS/MS as described below.

Quantitative TMT-based proteomic analysis was performed as previously described using a Thermo Eclipse with FAIMS LC-MS/MS ^16^. Acquired MS data was processed using ProLuCID search methodology in IP2 v.3-v.5 (Integrated Proteomics Applications, Inc.) ^49^. Trypsin cleavage specificity (cleavage at K, R except if followed by P) allowed for up to 2 missed cleavages. Carbamidomethylation of cysteine was set as a fixed modification, methionine oxidation, and TMT-modification of N-termini and lysine residues were set as variable modifications. Reporter ion ratio calculations were performed using summed abundances with most confident centroid selected from 20 ppm window. Only peptide-to-spectrum matches that are unique assignments to a given identified protein within the total dataset are considered for protein quantitation. High confidence protein identifications were reported with a <1% false discovery rate (FDR) cut-off. Differential abundance significance was estimated using ANOVA with Benjamini-Hochberg correction to determine p-values.

### Quantitative TMT Proteomics Analysis

Cells were treated with either DMSO vehicle or compound (JP-2-197 1 μM) for 24 h and lysate was prepared as described above. Briefly, 25-100 μg protein from each sample was reduced, alkylated and tryptically digested overnight. Individual samples were then labeled with isobaric tags using commercially available TMTsixplex (Thermo Fisher Scientific, P/N 90061) kits, in accordance with the manufacturer**’**s protocols. Tagged samples (20 μg per sample) were combined, dried with SpeedVac, resuspended with 300 μL 0.1% TFA in H2O, and fractionated using high pH reversed-phase peptide fractionation kits (Thermo Scientific, P/N 84868) according to manufacturer**’**s protocol. Fractions were dried with SpeedVac, resuspended with 50 μL 0.1% FA in H2O, and analyzed by LC-MS/MS as described below.

Quantitative TMT-based proteomic analysis was performed as previously described using a Thermo Eclipse with FAIMS LC-MS/MS ^16^. Acquired MS data was processed using ProLuCID search methodology in IP2 v.3-v.5 (Integrated Proteomics Applications, Inc.) ^49^. Trypsin cleavage specificity (cleavage at K, R except if followed by P) allowed for up to 2 missed cleavages. Carbamidomethylation of cysteine was set as a fixed modification, methionine oxidation, and TMT-modification of N-termini and lysine residues were set as variable modifications. Reporter ion ratio calculations were performed using summed abundances with most confident centroid selected from 20 ppm window. Only peptide-to-spectrum matches that are unique assignments to a given identified protein within the total dataset are considered for protein quantitation. High confidence protein identifications were reported with a <1% false discovery rate (FDR) cut-off. Differential abundance significance was estimated using ANOVA with Benjamini-Hochberg correction to determine p-values.

### Knockdown studies

Short-hairpin oligonucleotides were used to knock down the expression of RNF126 in C33A cells. For lentivirus production, lentiviral plasmids and packaging plasmids (pMD2.5G, Addgene catalog no. 12259 and psPAX2, Addgene catalog no. 12260) were transfected into HEK293T cells using Lipofectamine 2000 (Invitrogen). Lentivirus was collected from filtered cultured medium and used to infect the target cell line with 1:1000 dilution of polybrene. Target cells were selected over 3 d with 1 μg/ml of puromycin for C33A cells and 7.5 μg ml-1 for HEK293T cells. The short-hairpin sequences which were used for generation of the knockdown lines were: RNF126: TGCCATCATCACACAGCTCCT (Sigma RNF126 MISSION shRNA Bacterial Glycerol Stock, TRCN0000368954).

MISSION TRC1.5 pLKO.1- or TRC2 pLKO.5-puro Non-Mammalian shRNA Control (Sigma) was used as a control shRNA

## Data Availability Statement

The datasets generated during and/or analyzed during the current study are available from the corresponding author on reasonable request.

## Code Availability Statement

Data processing and statistical analysis algorithms from our lab can be found on our lab**’**s Github site: https://github.com/NomuraRG, and we can make any further code from this study available at reasonable request.

## References

1. Bond, M. J. & Crews, C. M. Proteolysis targeting chimeras (PROTACs) come of age: entering the third decade of targeted protein degradation. RSC Chem. Biol. 2, 725–742 (2021).

2. Hughes, S. J. & Ciulli, A. Molecular recognition of ternary complexes: a new dimension in the structure-guided design of chemical degraders. Essays Biochem. 61, 505–516 (2017).

3. Schreiber, S. L. The Rise of Molecular Glues. Cell 184, 3–9 (2021).

4. Burslem, G. M. & Crews, C. M. Small-Molecule Modulation of Protein Homeostasis. Chem. Rev. 117, 11269–11301 (2017).

5. Chamberlain, P. P. & Hamann, L. G. Development of targeted protein degradation therapeutics. Nat. Chem. Biol. 15, 937–944 (2019).

6. Chamberlain, P. P. et al. Structure of the human Cereblon-DDB1-lenalidomide complex reveals basis for responsiveness to thalidomide analogs. Nat. Struct. Mol. Biol. 21, 803–809 (2014).

7. Donovan, K. A. et al. Thalidomide promotes degradation of SALL4, a transcription factor implicated in Duane Radial Ray Syndrome. eLife 7, (2018).

8. Mayor-Ruiz, C. et al. Rational discovery of molecular glue degraders via scalable chemical profiling. Nat. Chem. Biol. 16, 1199–1207 (2020).

9. Powell, C. E. et al. Selective Degradation of GSPT1 by Cereblon Modulators Identified via a Focused Combinatorial Library. ACS Chem. Biol. 15, 2722–2730 (2020).

10. Nishiguchi, G. et al. Identification of Potent, Selective, and Orally Bioavailable Small-Molecule GSPT1/2 Degraders from a Focused Library of Cereblon Modulators. J. Med. Chem. 64, 7296–7311 (2021).

11. Scholes, N. S., Mayor-Ruiz, C. & Winter, G. E. Identification and selectivity profiling of small-molecule degraders via multi-omics approaches. Cell Chem. Biol. 28, 1048–1060 (2021).

12. Ito, T. et al. Identification of a primary target of thalidomide teratogenicity. Science 327, 1345–1350 (2010).

13. Matyskiela, M. E. et al. SALL4 mediates teratogenicity as a thalidomide-dependent cereblon substrate. Nat. Chem. Biol. 14, 981–987 (2018).

14. Han, T. et al. Anticancer sulfonamides target splicing by inducing RBM39 degradation via recruitment to DCAF15. Science 356, eaal3755 (2017).

15. Mayor-Ruiz, C. et al. Rational discovery of molecular glue degraders via scalable chemical profiling. Nat. Chem. Biol. 16, 1199–1207 (2020).

16. King, E. A. et al. Chemoproteomics-Enabled Discovery of a Covalent Molecular Glue Degrader Targeting NF-κB. 2022.05.18.492542 Preprint at https://doi.org/10.1101/2022.05.18.492542 (2022).

17. Slabicki, M. et al. The CDK inhibitor CR8 acts as a molecular glue degrader that depletes cyclin K. Nature 585, 293–297 (2020).

18. Slabicki, M. et al. Small-molecule-induced polymerization triggers degradation of BCL6. Nature 588, 164– 168 (2020).

19. Petretich, M., Demont, E. H. & Grandi, P. Domain-selective targeting of BET proteins in cancer and immunological diseases. Curr. Opin. Chem. Biol. 57, 184–193 (2020).

20. Poratti, M. & Marzaro, G. Third-generation CDK inhibitors: A review on the synthesis and binding modes of Palbociclib, Ribociclib and Abemaciclib. Eur. J. Med. Chem. 172, 143–153 (2019).

21. Weerapana, E. et al. Quantitative reactivity profiling predicts functional cysteines in proteomes. Nature 468, 790–795 (2010).

22. Backus, K. M. et al. Proteome-wide covalent ligand discovery in native biological systems. Nature 534, 570–574 (2016).

23. Spradlin, J. N. et al. Harnessing the anti-cancer natural product nimbolide for targeted protein degradation. Nat. Chem. Biol. 15, 747–755 (2019).

24. Spradlin, J. N., Zhang, E. & Nomura, D. K. Reimagining Druggability Using Chemoproteomic Platforms. Acc. Chem. Res. 54, 1801–1813 (2021).

25. Rodrigo-Brenni, M. C., Gutierrez, E. & Hegde, R. S. Cytosolic quality control of mislocalized proteins requires RNF126 recruitment to Bag6. Mol. Cell 55, 227–237 (2014).

26. Hu, X. et al. RNF126-Mediated Reubiquitination Is Required for Proteasomal Degradation of p97-Extracted Membrane Proteins. Mol. Cell 79, 320-331.e9 (2020).

27. Krysztofinska, E. M. et al. Structural and functional insights into the E3 ligase, RNF126. Sci. Rep. 6, 26433 (2016).

28. Ward, C. C. et al. Covalent Ligand Screening Uncovers a RNF4 E3 Ligase Recruiter for Targeted Protein Degradation Applications. ACS Chem. Biol. 14, 2430–2440 (2019).

29. Burslem, G. M. et al. Targeting BCR-ABL1 in Chronic Myeloid Leukemia by PROTAC-Mediated Targeted Protein Degradation. Cancer Res. 79, 4744–4753 (2019).

30. Lai, A. C. et al. Modular PROTAC Design for the Degradation of Oncogenic BCR-ABL. Angew. Chem. Int.Ed Eng l. 55, 807–810 (2016).

31. Tong, B. et al. A Nimbolide-Based Kinase Degrader Preferentially Degrades Oncogenic BCR-ABL. ACS Chem. Biol. 15, 1788–1794 (2020).

32. Farnaby, W. et al. BAF complex vulnerabilities in cancer demonstrated via structure-based PROTAC design. Nat. Chem. Biol. 15, 672–680 (2019).

33. Zengerle, M., Chan, K.-H. & Ciulli, A. Selective Small Molecule Induced Degradation of the BET Bromodomain Protein BRD4. ACS Chem. Biol. 10, 1770–1777 (2015).

34. Henning, N. J. et al. Discovery of a Covalent FEM1B Recruiter for Targeted Protein Degradation Applications. J. Am. Chem. Soc. 144, 701–708 (2022).

35. Uo, T., Plymate, S. R. & Sprenger, C. C. The potential of AR-V7 as a therapeutic target. Expert Opin. Ther. Targets 22, 201–216 (2018).

36. Dalal, K. et al. Selectively targeting the DNA-binding domain of the androgen receptor as a prospective therapy for prostate cancer. J. Biol. Chem. 289, 26417–26429 (2014).

37. Lee, G. T. et al. Effects of MTX-23, a Novel PROTAC of Androgen Receptor Splice Variant-7 and Androgen Receptor, on CRPC Resistant to Second-Line Antiandrogen Therapy. Mol. Cancer Ther. 20, 490–499 (2021).

38. Bhumireddy, A. et al. Design, synthesis, and biological evaluation of phenyl thiazole-based AR-V7 degraders. Bioorg. Med. Chem. Lett. 55, 128448 (2022).

39. Park, J. S. et al. Abstract LB-108: A potent and selective small molecule degrader of STAT5 for the treatment of hematological malignancies. Cancer Res. 80, LB–108 (2020).

40. Desroses, M. et al. STAT3 differential scanning fluorimetry and differential scanning light scattering assays: Addressing a missing link in the characterization of STAT3 inhibitor interactions. J. Pharm. Biomed. Anal. 160, 80–88 (2018).

41. Yang, J. et al. Covalent modification of Cys-239 in β-tubulin by small molecules as a strategy to promote tubulin heterodimer degradation. J. Biol. Chem. 294, 8161–8170 (2019).

42. Luo, M. et al. Chemoproteomics-enabled discovery of covalent RNF114-based degraders that mimic natural product function. Cell Chem. Biol. 28, 559-566.e15 (2021).

43. Zhang, X., Crowley, V. M., Wucherpfennig, T. G., Dix, M. M. & Cravatt, B. F. Electrophilic PROTACs that degrade nuclear proteins by engaging DCAF16. Nat. Chem. Biol. 15, 737–746 (2019).

44. Zhang, X. et al. DCAF11 Supports Targeted Protein Degradation by Electrophilic Proteolysis-Targeting Chimeras. J. Am. Chem. Soc. 143, 5141–5149 (2021).

45. Belcher, B. P., Ward, C. C. & Nomura, D. K. Ligandability of E3 Ligases for Targeted Protein Degradation Applications. Biochemistry (2021) doi:10.1021/acs.biochem.1c00464.

46. Tao, Y. et al. Targeted Protein Degradation by Electrophilic PROTACs that Stereoselectively and Site-Specifically Engage DCAF1. J. Am. Chem. Soc. (2022) doi:10.1021/jacs.2c08964.

47. Spradlin, J. N. et al. Harnessing the anti-cancer natural product nimbolide for targeted protein degradation. Nat. Chem. Biol. 15, 747–755 (2019).

48. Grossman, E. A. et al. Covalent Ligand Discovery against Druggable Hotspots Targeted by Anti-cancer Natural Products. Cell Chem. Biol. 24, 1368-1376.e4 (2017).

49. Xu, T. et al. ProLuCID: An improved SEQUEST-like algorithm with enhanced sensitivity and specificity. J. Proteomics 129, 16–24 (2015).

