## Supporting Information for "Rational Chemical Design of Molecular Glue Degraders"

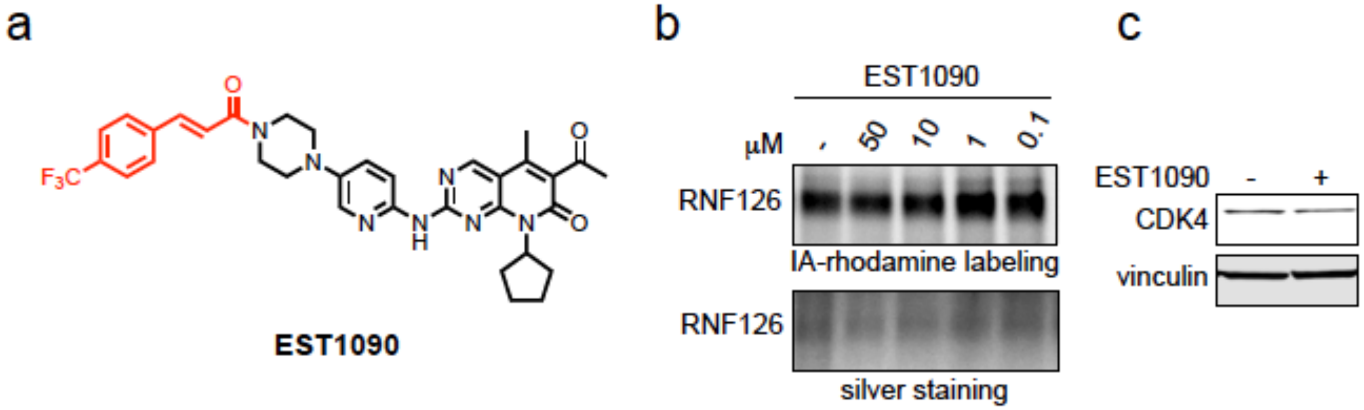


**Figure S1.** Palbociclib derivatives and their ability to bind to RNF126 and degrade CDK4. **(a)** Structure of EST1090 with the trifluoromethyl propenamide handle linked to the CDK4/6 inhibitor Palbociclib. **(b)** Gel-based ABPP of EST1090 against RNF126. Recombinant RNF126 was pre-incubated with DMSO vehicle or EST1090 for 30 min prior to labeling of RNF126 with IA-rhodamine (50 nM) for 1 h. Gels were visualized by in-gel fluorescence and protein loading was assessed by silver staining. **(c)** CDK4 levels in C33A cells. C33A cells were treated with DMSO vehicle or EST1090 (10 μM) for 24 h and CDK4 and vinculin levels were detected by Western blotting. Gels and blots in **(b, c)** are representative of n=3 biologically independent replicates/group.

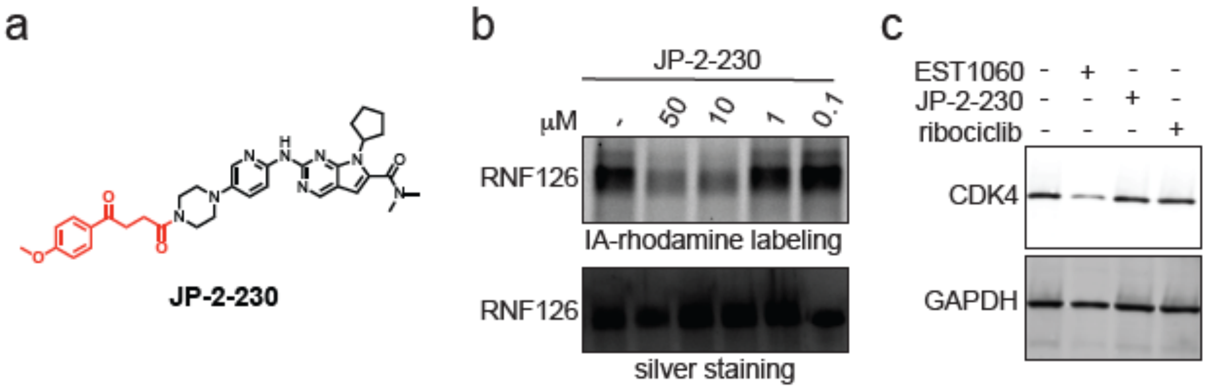


**Figure S2.** Ribociclib derivatives and their ability to bind to RNF126 and degrade CDK4. **(a)** Structure of JP-2-230 with non-reactive handle linked to the CDK4/6 inhibitor Ribociclib. **(b)** Gel-based ABPP of JP-2-230 against RNF126. Recombinant RNF126 was pre-incubated with DMSO vehicle or JP-2-230 for 30 min prior to labeling of RNF126 with IA-rhodamine (50 nM) for 1 h. Gels were visualized by in-gel fluorescence and protein loading was assessed by silver staining. **(c)** CDK4 levels in C33A cells. C33A cells were treated with DMSO vehicle, EST1060, or JP-2-230 (10 μM) for 24 h and CDK4 and GAPDH levels were detected by Western blotting. Gels and blots in **(b, c)** are representative of n=3 biologically independent replicates/group.

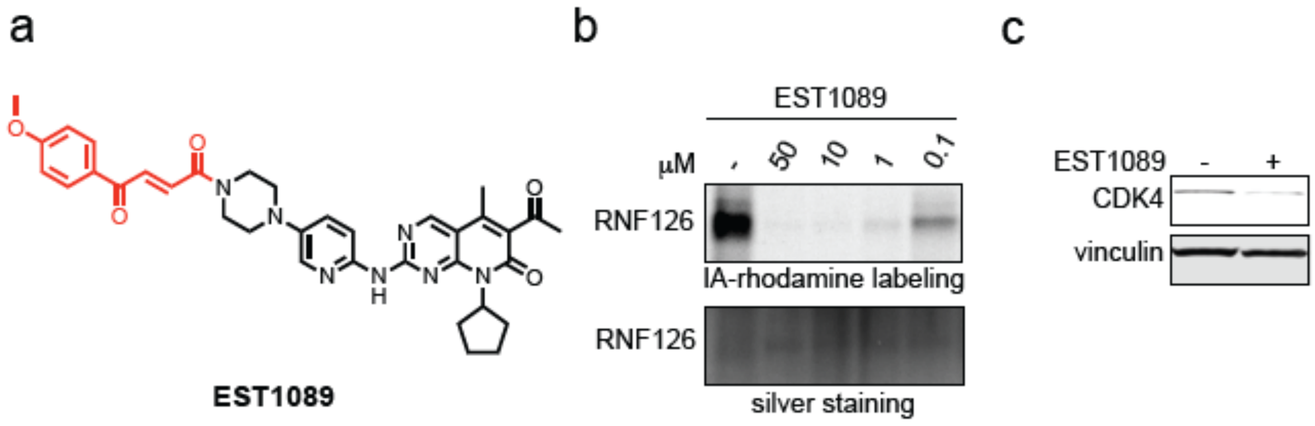


**Figure S3.** Palbociclib derivatives and their ability to bind to RNF126 and degrade CDK4. **(a)** Structure of EST1089 with fumarate handle linked to the CDK4/6 inhibitor Palbociclib. **(b)** Gel-based ABPP of EST0189 against RNF126. Recombinant RNF126 was pre-incubated with DMSO vehicle or EST1089 for 30 min prior to labeling of RNF126 with IA-rhodamine (50 nM) for 1 h. Gels were visualized by in-gel fluorescence and protein loading was assessed by silver staining. **(c)** CDK4 levels in C33A cells. C33A cells were treated with DMSO vehicle, EST1089 (10 μM) for 24 h and CDK4 and vinculin levels were detected by Western blotting. Gels and blots in **(b, c)** are representative of n=3 biologically independent replicates/group.

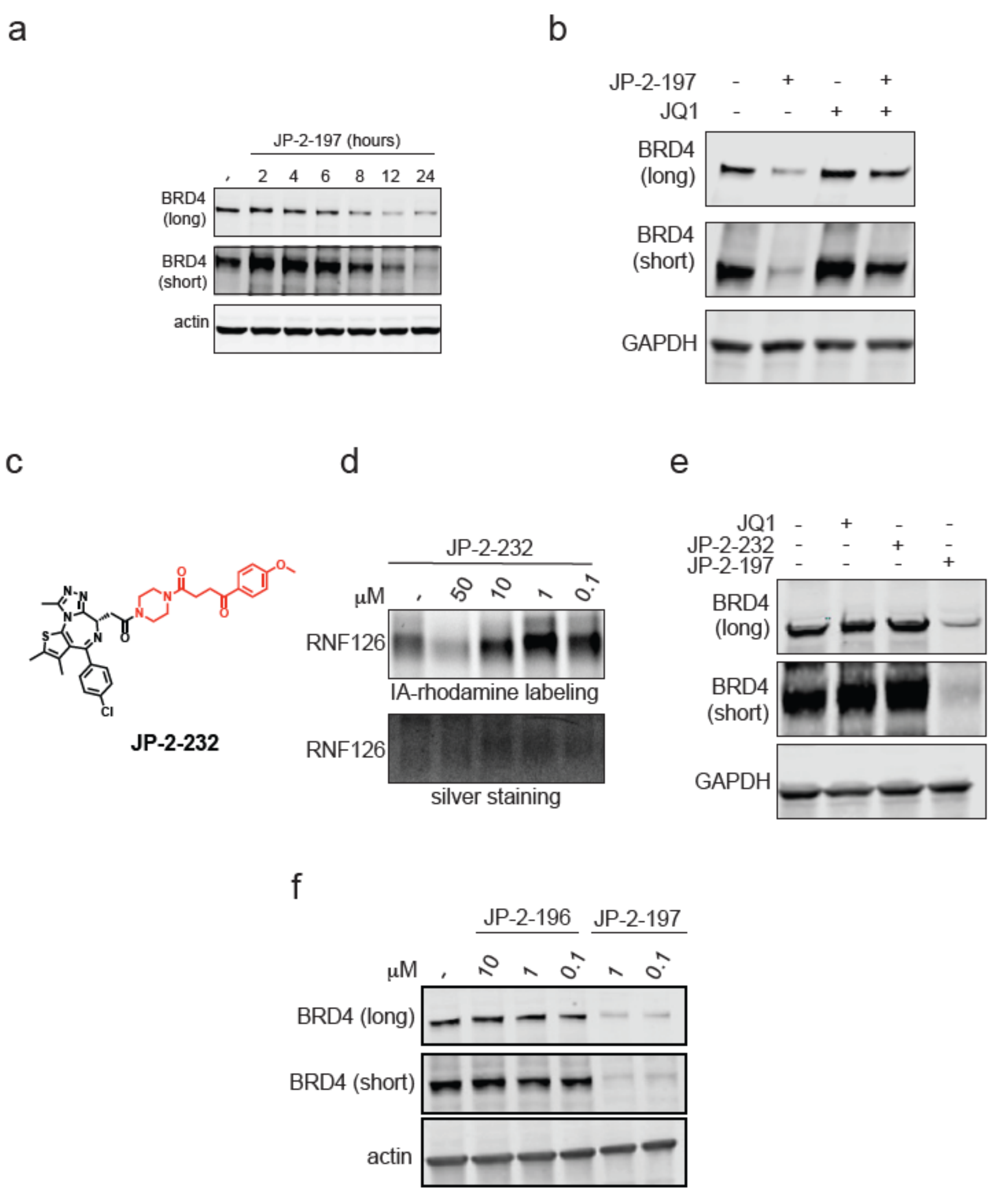


**Figure S4. Characterization of BRD4 degraders and their derivatives. (a)** Time-course of BRD4 degradation with JP-2-197. HEK293T cells were treated with DMSO vehicle or JP-2-197 (1 μM) and BRD4 and loading control actin levels were assessed by Western blotting. **(b)** HEK293T cells were pre-treated with either DMSO vehicle or JQ1 (1 μM) for 1 h prior to treating cells with DMSO or JP-2-197 (0.1 μM) for 24 h and BRD4 and loading control GAPDH levels were assessed by Western blotting. **(c)** Structure of non-reactive derivative of JP-2-197, JP-2-232. **(d)** Gel-based ABPP of JP-2-232 against RNF126. Recombinant RNF126 was pre-incubated with DMSO vehicle or JP-2-232 for 30 min prior to labeling of RNF126 with IA-rhodamine (50 nM) for 1 h. Gels were visualized by in-gel fluorescence and protein loading was assessed by silver staining. **(e)** BRD4 levels in HEK293T cells. HEK293T cells were treated with DMSO vehicle, JP-2-197, or JP-2-232 (1 μM) for 24 h and BRD4 and GAPDH levels were detected by Western blotting. **(f)** HEK293T cells were treated with DMSO vehicle, JP-2-196, or JP-2-197 for 24 h and BRD4 and loading control acting levels were assessed by Western blotting. Gels and blots in **(a, b, d, e, f)** are representative of n=3 biologically independent replicates/group.

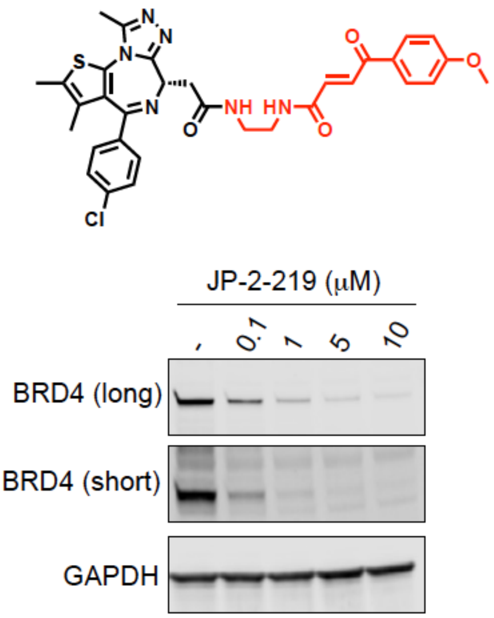


**Figure S5. Characterization of BRD4 degraders and their derivatives.** Shown above is a structure of a BRD4 degrader with an alkyl linker rather than a piperazine linking the fumarate handle to the BRD4 inhibitor JQ1. Shown below are BRD4 levels in HEK293T cells. HEK293T cells were treated with DMSO vehicle, JP-2-197, or JP-2-232 (1 μM) for 24 h and BRD4 and loading control GAPDH levels were detected by Western blotting. Gels and blots are representative of n=3 biologically independent replicates/group.

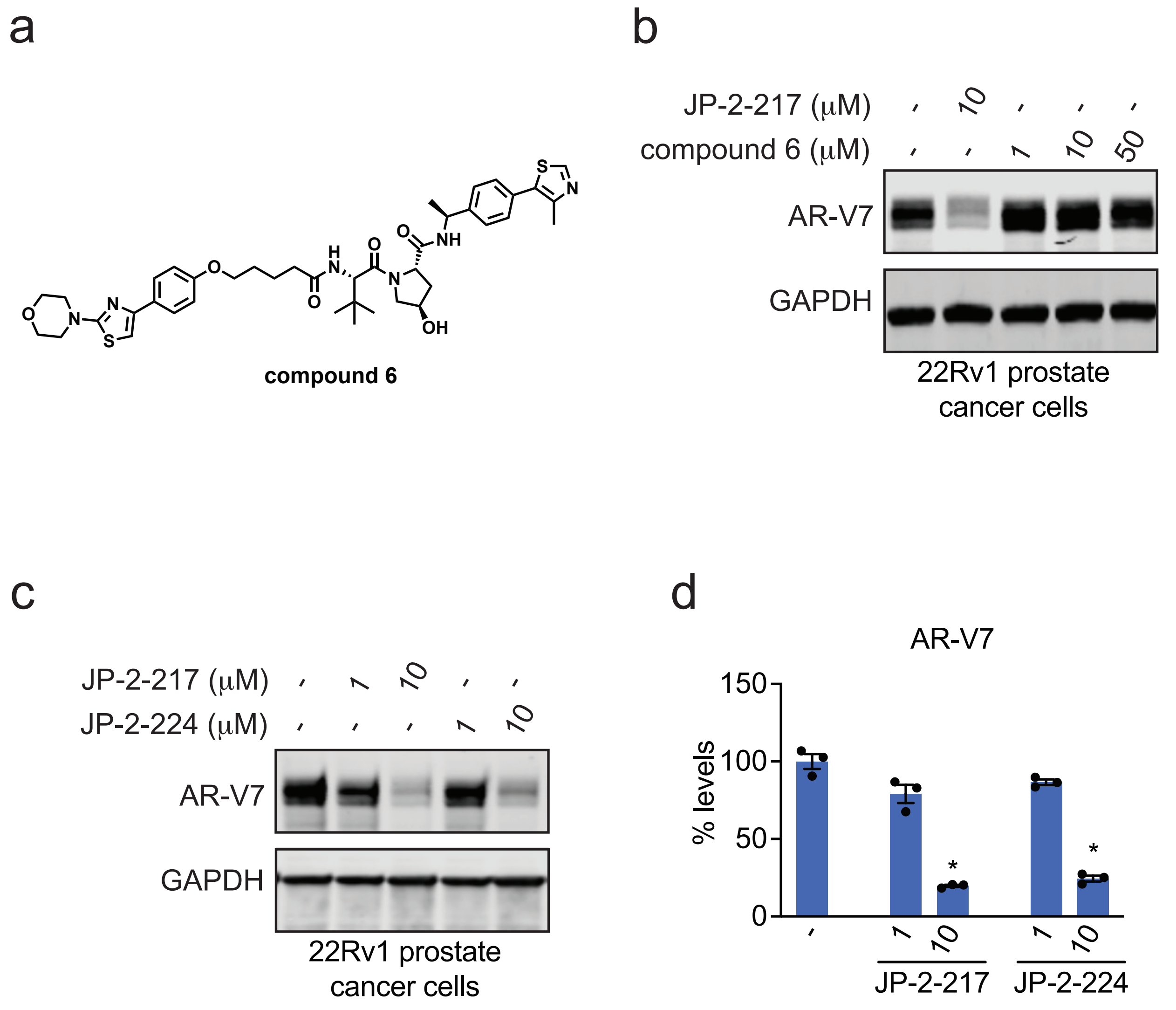


**Figure S6. Characterization of AR-V7 Degrader. (a)** Structure of a previously reported AR-V7 degrader linking the AR DNA binding domain ligand VPC-14228 to a VHL recruiter, compound 6. **(b)** AR-V7 levels in 22Rv1 cells. 22Rv1 cells were treated with DMSO vehicle, JP-2-217, or compound 6 for 24 h and AR-V7 and GAPDH loading control levels were detected by Western blotting. **(c)** 22Rv1 cells were treated with DMSO vehicle, JP-2-217, or JP-2-224 for 24 h and AR-V7 and loading control GAPDH levels were detected by Western blotting. **(d)** Quantified AR-V7 levels from experiment described in **(c).** Blots are representative of n=3 biologically independent replicates/group with individual replicates and average/sem values plotted in the bar graph. Statistical significance is calculated as *$p<0.05$ compared to DMSO vehicle.

**Supporting Table Legends**

**Table S1. Cysteine chemoproteomic profiling of EST1027 in C33A cells using isoTOP-ABPP.** C33A cells were treated with DMSO vehicle or EST1027 (10 μM) for 4 h**.** Resulting lysates were labeled with an alkyne-functionalized iodoacetamide probe (IA-alkyne) (200 μM) for 1 h, after which isotopically light (for control) or heavy (for treated) biotin-azide tags bearing a TEV cleavable site were appended onto probe-labeled proteins by copper-catalyzed azide-alkyne cycloaddition. Control and treated groups were combined in a 1:1 ratio, avidin-enriched, tryptically digested, and probe-modified tryptic peptides were eluted by TEV protease and analyzed by LC/MS/MS and probe-modified peptides were quantified. Shown in the first tab are the raw data, and shown in the second tab are only those probe-modified peptides that appeared in at least two out of the three biological replicates.

**Table S2. Quantitative Proteomics of JP-2-196-Alkyne Pulldown in HEK293T cells.** JP-2-196 pulldown proteomics showing significant and moderately selective engagement of RNF126 with less significant engagement of 5 additional E3 ubiquitin ligases LRSAM1, RNF40, MID2, RNF219, and RNF14. HEK293T cells were treated with DMSO vehicle or JP-2-196-alkyne (10 mM) for 4 h. Subsequent lysates were subjected to copper-catalyzed azide-alkyne cycloaddition (CuAAC) with an azide-functionalized biotin handle, after which probe-modified proteins were avidin-enriched, eluted, and digested, and analyzed by TMT-based quantitative proteomics. Data shown are ratio of JP-2-196-alkyne vs DMSO control enriched proteins and p-values from n=3 biologically indepdent replicates/group.

**Table S3. TMT-based quantitative proteomic profiling of JP-2-197 in HEK293T cells.** HEK293T cells were treated with DMSO vehicle or JP-2-197 (1 μM) for 24 h. Data are from n=3 biological replicates per group.

**Synthetic Methods and Characterization**

All chemical reactions were carried out under a nitrogen atmosphere with dry solvents under anhydrous conditions, unless otherwise noted. Reagents were purchased at the highest commercial quality and used without further purification, unless otherwise stated. Reactions were stirred magnetically and monitored by thin layer chromatography (TLC) carried out on Merck glass silica gel plates (60 F254) using UV light as a visualizing agent and iodine and/or phosphomolybdic acid stain as developing agents. Solvents were removed *in vacuo* using either a Buchi R-300 Rotavapor (equipped with an I-300 Pro Interface, B-300 Base Heating Bath, Welch 2037B-01 DryFast pump, and VWR AD15R-40-V11B Circulating Bath). Solvents for silica gel chromatography were used as supplied by Sigma-Aldrich. Automated flash chromatography was performed on a Biotage Isolera instrument, equipped with a UV detector. Chromatograms were recorded at 254 and 280 nm. Low-resolution mass spectra were obtained using Agilent 6460 Triple Quad LC/MS instrument. High-resolution mass spectra (HRMS) were obtained at the Catalysis Center at the College of Chemistry, University of California, Berkeley. $^{1}H$ and $^{13}C$ nuclear magnetic resonance (NMR) spectra were recorded on Nuclear Magnetic Resonance (NMR) spectra were recorded on BRUKER AV (600 MHz and 700 MHz), AVB (400 MHz), AVQ (400 MHz) and NEO (500 MHz) spectrometers. Measurements were carried out at ambient temperature. Chemical shifts (δ) are reported in ppm with the residual solvent signal as internal standard (chloroform at 7.26 and 77.00 ppm for $^{1}H$ NMR and $^{13}C$ NMR spectroscopy, respectively). The data is reported as (s = singlet, d = doublet, t = triplet, q = quartet, p = quintet, m = multiplet or unresolved, br = broad signal, coupling constant(s) in Hz, integration). 13C NMR spectra were recorded with broadband 1H decoupling.

**General Procedures**

**Amide Couplings**

**General Procedure A**

A mixture of the corresponding carboxylic acid (1.1 equiv.) and HATU (1.2 equiv.) was purged with $N_2$ for 5 minutes. The mixture was dissolved in *N,N*-dimethylformamide (DMF) (0.1 M), *N,N*-diisopropylethylamine (DIPEA) (3 equiv.) was added and the reaction mixture was allowed to stir at ambient temperature for 30 minutes. The corresponding amine (1 equiv.) was dissolved in DMF (0.1 M) then added dropwise and the reaction mixture was stirred at ambient temperature overnight. The reaction was quenched with 5 times the reaction volume of 5% $LiCl_{(aq)}$ and extracted 3 times with dichloromethane (DCM) or ethyl acetate (EtOAc). The organic extracts were dried over $Na_2SO_4$, vacuum filtered, and concentrated *in vacuo*. The resultant residue was purified by silica gel flash chromatography to afford the title compound.

**General Procedure B**

The corresponding carboxylic acid (1.1 equiv.) was added to a vessel and purged with $N_2$ for 5 minutes. The acid was then dissolved in DCM (0.1 M) and cooled to 0 °C in an ice bath. Oxalyl chloride (1.2 equiv.) was added dropwise at 0 °C. A few drops of DMF were added and the reaction mixture was allowed to stir and come to ambient temperature over 2 hours. The volatiles were removed *in vacuo* and the resultant residue was redissolved in DCM (0.1 M) and cooled to 0 °C in an ice bath. The corresponding amine (1.1 equiv.) was dissolved in DCM (0.1M). DIPEA (3 equiv.) was added, and the mixture was stirred for 5 minutes at ambient temperature before being added to the acyl chloride dropwise at 0 °C. The reaction mixture was allowed to stir and come to ambient temperature overnight. The reaction was quenched with water and extracted 3 times with EtOAc. The organic extracts were dried over $Na_2SO_4$, vacuum filtered, and concentrated *in vacuo*. The resultant residue was purified by silica gel flash chromatography to afford the title compound.

**General Procedure C**

The corresponding carboxylic acid (1.1 equiv.) was added to a vessel and purged with $N_2$ for 5 minutes. The acid was dissolved in acetonitrile (0.1 M) and DIPEA (2 equiv.) was added. Pentafluoropyridine (1.1 equiv.) was added dropwise, and the reaction mixture was stirred at ambient temperature for 30 minutes. The corresponding amine was dissolved in acetonitrile (0.1 M) and added to acyl fluoride. The reaction mixture was stirred at 100 0 °C overnight. The volatiles were removed *in vacuo*, and the resultant residue was purified by silica gel flash chromatography to afford the title compound.

**General Procedure D**

The corresponding carboxylic acid (1.0 equiv.) was added to a vessel and purged with $N_2$ for 5 minutes. The acid was dissolved in DMF (0.1 M) and DIPEA (3 equiv.) was added. A >50% wt. solution of propylphosphonic anhydride (T3P) in EtOAc (1.5 equiv.) was added dropwise, and the reaction mixture was stirred at ambient temperature for 30 minutes. The corresponding amine (1.2 equiv.) was dissolved in DMF (0.1 M) then added dropwise and the reaction mixture was stirred at ambient temperature overnight. The reaction was quenched with 5 times the reaction volume of 5% $LiCl_{(aq)}$ and extracted 3 times with EtOAc. The organic extracts were washed once with 5% $LiCl_{(aq)}$, dried over $Na_2SO_4$, vacuum filtered, and concentrated *in vacuo*. The resultant residue was purified by silica gel flash chromatography to afford the title compound.

**Tert-butyloxycarbonyl Deprotection**

**General Procedure E**

The corresponding tert-butyloxycarbonyl protected amine (1 equiv.) was dissolved in DCM (0.1 M). Trifluoroacetic acid (32 equiv.) was added, and the reaction mixture was stirred at ambient temperature for 30 minutes to overnight. The volatiles were removed *in vacuo* and the crude residue was used without further purification, unless otherwise noted.

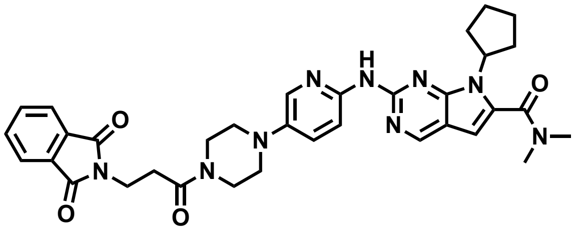

**EST1001**

**General Procedure A** was followed with 3-phthalimidopropionic acid (21.6 mg, 0.06 mmol), HATU (43.8 mg, 0.12 mmol), DIPEA (0.04 mL, 0.23 mmol), and ribociclib (25.0 mg, 0.06 mmol). The crude residue was purified by silica gel chromatography (0-10% MeOH in DCM) to afford 14.9 mg (41%) of the title compound as a yellow film.

**$^1$H NMR** (500 MHz, $CDCl_3$) δ 8.70 (s, 1H), 8.37 (d, *J* = 9.0 Hz, 1H), 8.03 – 7.95 (m, 2H), 7.89 – 7.81 (m, 2H), 7.75 – 7.68 (m, 2H), 7.32 (dd, *J* = 9.1, 3.0 Hz, 1H), 6.44 (s, 1H), 4.79 (p, *J* = 9.0 Hz, 1H), 4.09 – 4.03 (m, 2H), 3.80 (t, *J* = 5.2 Hz, 2H), 3.64 (t, *J* = 5.1 Hz, 2H), 3.15 (s, 6H), 3.14 – 3.08 (m, 4H), 2.85 – 2.78 (m, 2H), 2.65 – 2.52 (m, 2H), 2.06 (m 4H), 1.77 – 1.70 (m, 2H).

**$^{13}$C NMR** (151 MHz, $CDCl_3$) δ 168.4, 168.2, 164.1, 154.5, 151.9, 151.8, 147.6, 142.1, 137.6, 134.0, 132.1, 132.1, 127.4, 123.3, 112.7, 112.4, 101.0, 57.9, 50.6, 50.3, 45.4, 41.5, 34.3, 31.6, 30.2, 24.7.

**LRMS (ESI)** *m/z* calcd for $[C_{34}H_{37}N_9O_4 + H]^+$ = 636.3, found 636.4.

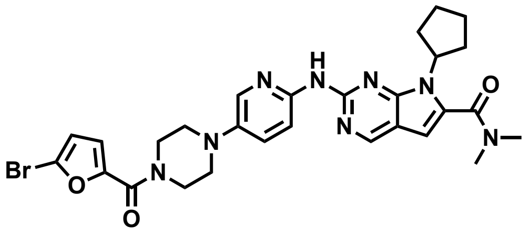

**EST1004**

**General Procedure A** was followed with 5-Bromo-2-furoic acid (11.0 mg, 0.06 mmol), HATU (43.8 mg, 0.12 mmol), DIPEA (0.04 mL, 0.23 mmol), and ribociclib (25.0mg, 0.06 mmol). The crude residue was purified by silica gel chromatography (0-15% MeOH in DCM) to afford 23.9 mg (63%) of the title compound as a yellow film.

$^1$**H NMR** (500 MHz, $CDCl_3$) δ 8.72 (s, 1H), 8.39 (d, *J* = 9.0 Hz, 1H), 8.25 (s, 1H), 8.06 (d, *J* = 3.0 Hz, 1H), 7.34 (dd, *J* = 9.1, 3.0 Hz, 1H), 7.03 (d, *J* = 3.5 Hz, 1H), 6.48 – 6.42 (m, 2H), 4.79 (p, *J* = 8.9 Hz, 1H), 3.98 (s, 4H), 3.24 – 3.18 (m, 4H), 3.15 (s, 6H), 2.58 (m, 2H), 2.11 – 2.01 (m, 4H), 1.76 – 1.68 (m, 2H).

$^{13}$**C NMR** (151 MHz, $CDCl_3$) δ 164.1, 157.8, 154.6, 151.9, 151.9, 149.5, 147.7, 142.0, 137.5, 132.0, 127.3, 124.4, 119.3, 113.5, 112.6, 112.5, 101.0, 57.9, 50.6, 30.2, 24.7.

**LRMS (ESI)** *m/z* calcd for $[C_{28}H_{31}BrN_8O_3 + H]^+$ = 607.3, found 607.2.

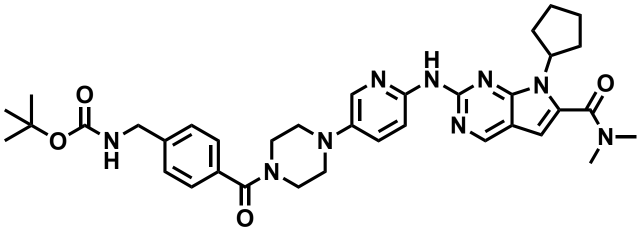

**EST1007**

**General Procedure A** was followed with 4-(Boc-aminomethyl)benzoic acid (14.5 mg, 0.06 mmol), HATU (43.8 mg, 0.12 mmol), DIPEA (0.04 mL, 0.23 mmol), and ribociclib (25.0 mg, 0.06 mmol). The crude residue was purified by silica gel chromatography (0-15% MeOH in DCM) to afford 29.5 mg (77%) of the title compound as a yellow film.

$^1$**H NMR** (500 MHz, $CDCl_3$) δ 8.72 (s, 1H), 8.37 (d, *J* = 9.1 Hz, 1H), 8.26 (s, 1H), 8.04 (d, *J* = 2.9 Hz, 1H), 7.41 (d, *J* = 7.9 Hz, 2H), 7.36 – 7.30 (m, 3H), 6.44 (s, 1H), 4.99 (d, *J* = 6.5 Hz, 1H), 4.79 (p, *J* = 9.0 Hz, 1H), 4.35 (d, *J* = 6.1 Hz, 2H), 3.94 (s, 2H), 3.62 (s, 2H), 3.20 (br s, 2H), 3.15 (s, 6H), 3.09 (br s, 2H), 2.58 (m, 2H), 2.11 – 2.00 (m, 4H), 1.76 – 1.67 (m, 2H), 1.46 (s, 9H).

**$^{13}$C NMR** (151 MHz, $CDCl_3$) δ 170.2, 164.1, 156.0, 154.6, 151.9, 151.9, 147.7, 142.0, 141.1, 137.6, 134.4, 132.0, 127.5, 127.4, 112.7, 112.5, 101.0, 79.7, 57.9, 50.6, 44.3, 39.4, 35.2, 30.2, 28.4, 24.7.

**LRMS (ESI)** *m/z* calcd for $[C_{36}H_{45}N_9O_4 + H]^+$ += 668.4, found 668.5.

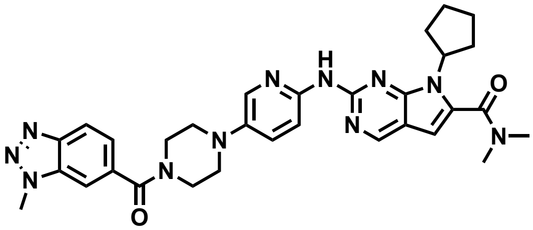


**EST1012**

**General Procedure A** was followed with 1-Methyl-1h-1,2,3-benzotriazole-6-carboxylic acid (10.2 mg, 0.06 mmol), HATU (43.8 mg, 0.12 mmol), DIPEA (0.04 mL, 0.23 mmol), and ribociclib (25.0 mg, 0.06 mmol). The crude residue was purified by silica gel chromatography (0-15% MeOH in DCM) to afford 23.7 mg (69%) of the title compound as a yellow film.

**$^1$H NMR** (500 MHz, $CDCl_3$) δ 8.72 (s, 1H), 8.34 (s, 1H), 8.21 (s, 1H), 8.15 (d, *J* = 1.1 Hz, 1H), 8.04 (d, *J* = 2.9 Hz, 1H), 7.68 – 7.58 (m, 2H), 7.35 (dd, *J* = 9.0, 2.9 Hz, 1H), 6.44 (s, 1H), 4.79 (p, *J* = 8.9 Hz, 1H), 4.34 (s, 3H), 3.94 (br s, 2H), 3.70 (br s, 2H), 3.15 (m, 10H), 2.57 (m, 2H), 2.12 – 1.99 (m, 4H), 1.71 (d, *J* = 4.1 Hz, 2H).

**$^{13}$C NMR** (151 MHz, $CDCl_3$) δ 169.7, 164.0, 154.4, 151.9, 145.3, 142.0, 134.1, 131.3, 127.0, 119.2, 112.8, 112.6, 109.9, 101.0, 57.9, 50.6, 34.4, 31.9, 30.2, 29.4, 24.7, 22.7, 14.2, 14.1.

**LRMS (ESI)** *m/z* calcd for $[C_{31}H_{35}N_{11}O_2 + H]^+$ = 594.3, found 594.3 .

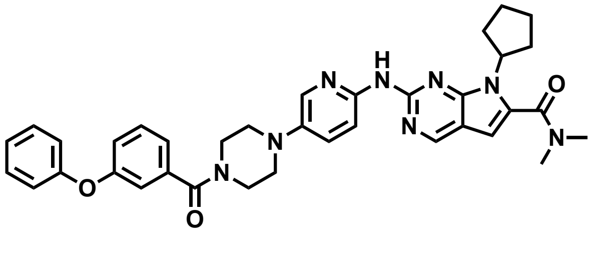


**EST1018**

**General Procedure A** was followed with 3-phenoxybenzoic acid (12.3 mg, 0.06 mmol), HATU (43.8 mg, 0.12 mmol), DIPEA (0.04 mL, 0.23 mmol), and ribociclib (25.0 mg, 0.06 mmol). The crude residue was purified by silica gel chromatography (0-15% MeOH in DCM) to afford 30.1 mg (83%) of the title compound as a yellow film.

**$^1$H NMR** (500 MHz, $CDCl_3$) δ 8.72 (s, 1H), 8.39 (d, *J* = 9.0 Hz, 1H), 8.30 (s, 1H), 8.05 (d, *J* = 2.9 Hz, 1H), 7.47 – 7.42 (m, 2H), 7.42 – 7.31 (m, 3H), 7.19 – 7.12 (m, 1H), 7.04 (ddd, *J* = 13.1, 7.6, 1.6 Hz, 4H), 6.44 (s, 1H),

4.79 (p, $J$ = 8.9 Hz, 1H), 3.96 – 3.62 (m, 4H), 3.15 (m, 10H), 2.64 – 2.53 (m, 2H), 2.11 – 2.00 (m, 4H), 1.76 – 1.65 (m, 2H).

**$^{13}$C NMR** (151 MHz, $CDCl_3$) δ 170.1, 164.1, 159.1, 156.2, 154.6, 151.9, 151.9, 147.7, 142.1, 137.6, 132.0, 130.0, 129.8, 129.3, 127.4, 124.1, 119.6, 118.1, 112.6, 112.5, 101.1, 57.9, 50.7, 39.4, 35.2, 33.7, 31.9, 30.2, 29.4, 26.7, 24.7, 23.2, 22.7, 14.2, 14.1. (Mixture of rotamers)

**LRMS (ESI)** *m/z* calcd for $[C_{36}H_{38}N_8O_3 + H]^+$ = 631.3, found 631.3.

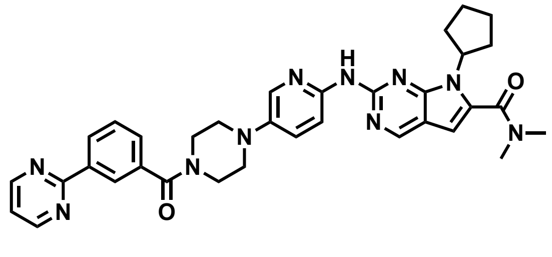

**EST1021**

**General Procedure A** was followed with 3-(2-pyrimidinyl)benzoic acid (11.5 mg, 0.06 mmol), HATU (43.8 mg, 0.12 mmol), DIPEA (0.04 mL, 0.23 mmol), and ribociclib (25.0 mg, 0.06 mmol). The crude residue was purified by silica gel chromatography (0-15% MeOH in DCM) to afford 12.2 mg (34%) of the title compound as a yellow film.

**$^1$H NMR** (500 MHz, $CDCl_3$) δ 8.83 (d, $J$ = 4.8 Hz, 2H), 8.69 (s, 1H), 8.54 (dt, $J$ = 8.9, 1.7 Hz, 2H), 8.38 (d, $J$ = 9.1 Hz, 1H), 8.01 (d, $J$ = 2.9 Hz, 1H), 7.92 (s, 1H), 7.63 – 7.54 (m, 2H), 7.35 (dd, $J$ = 9.1, 2.9 Hz, 1H), 7.23 (t, $J$ = 4.8 Hz, 1H), 6.44 (s, 1H), 4.79 (p, $J$ = 8.9 Hz, 1H), 4.01 (br s, 2H), 3.69 (br s, 2H), 3.25 (br s, 2H), 3.15 (s, 6H), 3.13 (br s, 2H), 2.64 – 2.52 (m, 2H), 2.12 – 1.99 (m, 4H), 1.77 – 1.69 (m, 2H).

**$^{13}$C NMR** (151 MHz, $CDCl_3$) δ 170.1, 164.1, 163.9, 157.4, 154.5, 151.9, 151.8, 147.6, 142.2, 137.9, 135.9, 132.1, 129.5, 129.5, 129.1, 127.4, 126.8, 119.5, 112.7, 112.5, 101.0, 57.9, 31.9, 30.2, 29.4, 24.7, 22.7, 14.1.

**LRMS (ESI)** *m/z* calcd for $[C_{34}H_{36}N_{10}O_2 + H]^+$ = 617.3, found 617.3.

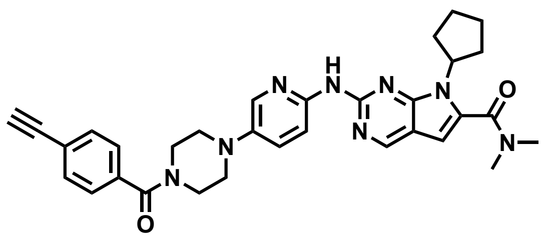

**EST1026**

**General Procedure A** was followed with 4-Ethynylbenzoic acid (8.4 mg, 0.06 mmol), HATU (43.8 mg, 0.12 mmol), DIPEA (0.04 mL, 0.23 mmol), and ribociclib (25.0 mg, 0.06 mmol). The crude residue was purified by silica gel chromatography (0-15% MeOH in DCM) to afford 13.6 mg (42%) of the title compound as a yellow film.

**$^1$H NMR** (500 MHz, CDCl3) δ 8.70 (s, 1H), 8.35 (d, J = 9.1 Hz, 1H), 8.16 (s, 1H), 7.99 (d, J = 2.9 Hz, 1H), 7.58 - 7.53 (m, 2H), 7.44 - 7.39 (m, 2H), 7.35 (dd, J = 9.1, 3.0 Hz, 1H), 6.45 (s, 1H), 4.79 (p, J = 9.0, 9.0, 9.0, 9.0 Hz, 1H), 3.93 (s, 2H), 3.61 (s, 2H), 3.16 (d, J = 3.9 Hz, 10H), 2.61 (s, 1H), 2.56 (dt, J = 16.4, 8.3, 8.3 Hz, 2H), 2.11 - 2.01 (m, 4H), 1.74 - 1.68 (m, 2H).

**$^{13}$C NMR** (151 MHz, $CDCl_3$) δ 169.7, 164.0, 154.3, 151.9, 151.6, 142.0, 135.6, 132.3, 127.2, 123.9, 113.9, 112.7, 101.0, 82.8, 78.9, 57.9, 41.0, 39.4, 31.9, 30.2, 29.4, 24.7, 22.7, 14.1.

**LRMS (ESI)** *m/z* calcd for $[C_{32}H_{34}N_8O_2 + H]^+$ = 563.3, found 563.3.

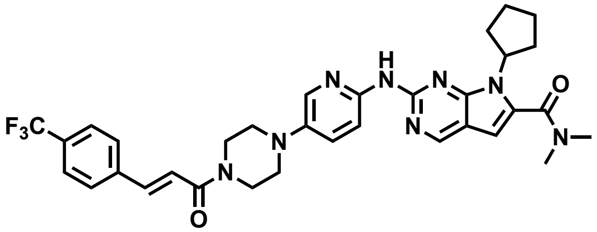


**EST1027**

**General Procedure A** was followed with *trans*-4-(trifluoromethyl)cinnamic acid (12.4 mg, 0.06 mmol), HATU (43.8 mg, 0.12 mmol), DIPEA (0.04 mL, 0.23 mmol), and ribociclib (25.0 mg, 0.06 mmol). The crude residue was purified by silica gel chromatography (0-15% MeOH in DCM) to afford 24.7 mg (68%) of the title compound as a yellow film.

**$^1$H NMR** (500 MHz, $CDCl_3$) δ 8.73 (s, 1H), 8.17 (s, 1H), 8.00 (d, *J* = 2.9 Hz, 1H), 7.71 (d, *J* = 15.4 Hz, 1H), 7.64 (s, 4H), 7.44 – 7.37 (m, 1H), 7.01 (d, *J* = 15.5 Hz, 1H), 6.46 (s, 1H), 4.77 (q, *J* = 8.9 Hz, 1H), 3.99 – 3.78 (m, 4H), 3.19 (br s, 4H), 3.15 (s, 6H), 2.53 (m, 2H), 2.10 – 1.97 (m, 4H), 1.69 (m, 2H).

**$^{13}$C NMR** (151 MHz, $CDCl_3$) δ 165.0, 163.8, 151.9, 142.1, 141.5, 138.5, 131.3 (q, $^4J_{CF}$ = 32.5 Hz), 129.5, 128.0, 125.8 (q, $^3J_{CF}$ = 3.7 Hz), 123.9 (q, $^1J_{CF}$ = 272.2 Hz), 119.3, 113.2, 101.1, 58.0, 45.7, 42.1, 39.4, 35.2, 33.7, 31.9, 30.3, 30.2, 29.4, 24.7, 22.7.

**HRMS (ESI)** *m/z* calcd for $[C_{33}H_{35}F_3N_8O_2 + H]^+$ = 633.2835, found 633.2906.

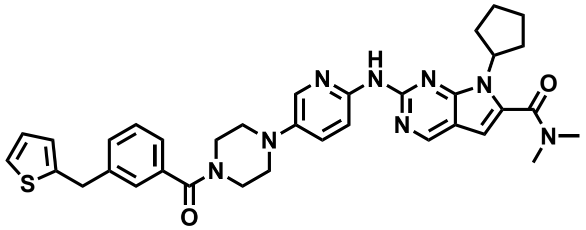

**EST1030**

**General Procedure A** was followed with 4-(2-thienylmethyl)benzoic acid (12.6 mg, 0.06 mmol), HATU (43.8 mg, 0.12 mmol), DIPEA (0.04 mL, 0.23 mmol), and ribociclib (25.0 mg, 0.06 mmol). The crude residue was purified by silica gel chromatography (0-15% MeOH in DCM) to afford 32.3 mg (88%) of the title compound as a yellow film.

**$^1$H NMR** (500 MHz, $CDCl_3$) δ 8.65 (s, 1H), 8.35 (d, *J* = 9.1 Hz, 1H), 7.89 (d, *J* = 2.9 Hz, 1H), 7.35 – 7.28 (m, 3H), 7.27 – 7.21 (m, 2H), 7.10 (dd, *J* = 5.2, 1.2 Hz, 1H), 6.87 (dd, *J* = 5.2, 3.4 Hz, 1H), 6.75 (dd, *J* = 3.3, 1.2 Hz, 1H), 6.38 (s, 1H), 4.73 (p, *J* = 8.8 Hz, 1H), 4.12 (s, 2H), 3.94 – 3.50 (m, 4H), 3.08 (s, 10H), 2.54 – 2.43 (m, 2H), 2.06 – 1.91 (m, 4H), 1.70 – 1.60 (m, 2H).

**$^{13}$C NMR** (151 MHz, $CDCl_3$) δ 170.4, 163.9, 154.0, 151.9, 151.3, 147.1, 143.0, 142.5, 142.1, 133.6, 132.6, 128.8, 128.4, 127.5, 126.9, 125.5, 124.2, 113.1, 113.0, 100.9, 57.9, 50.4, 39.4, 35.8, 35.2, 31.9, 30.3, 30.2, 29.4, 24.8, 23.2, 22.7.

**LRMS (ESI)** *m/z* calcd for $[C_{35}H_{38}N_8O_2S + H]^+$ = 635.3, found 635.4.

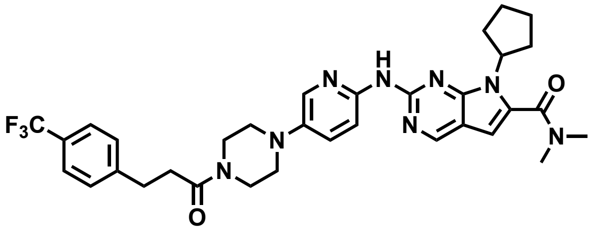

**EST1036**

**General Procedure A** was followed with 4-(trifluoromethyl)hydrocinnamic acid (99,5 mg, 0.46 mmol), HATU (350.2 mg, 0.92 mmol), DIPEA (0.32 mL, 1.4 mmol), and ribociclib (200.1 mg, 0.46 mmol). The crude residue was purified by reverse phase silica gel chromatography (5-95% MeCN in $H_2O$) to afford 133.0 mg (46%) of the title compound as a yellow film.

**$^1$H NMR** (500 MHz, $CDCl_3$) δ 9.93 (s, 1H), 8.76 (s, 1H), 7.79 (s, 1H), 7.54 (d, *J* = 8.2 Hz, 2H), 7.47 (d, *J* = 7.9 Hz, 2H), 7.32 (d, *J* = 7.9 Hz, 2H), 6.48 (s, 1H), 4.73 (p, *J* = 8.8 Hz, 1H), 3.74 (t, *J* = 5.0 Hz, 2H), 3.58 (t, *J* = 4.6

Hz, 2H), 3.12 (d, $J$ = 13.6 Hz, 6H), 3.08 – 3.00 (m, 6H), 2.71 (t, $J$ = 7.5 Hz, 2H), 2.38 (m, 2H), 2.05 – 1.91 (m, 4H), 1.64 – 1.56 (m, 2H).

**$^{13}$C NMR** (151 MHz, $CDCl_3$) δ 170.5, 163.2, 152.4, 151.8, 145.3, 142.1, 128.9, 128.4 (q, $^2J_{CF}$ = 32.2 Hz), 125.4 (q, $^3J_{CF}$ = 3.7 Hz), 124.3 (q, $^1J_{CF}$ = 271.8 Hz), 114.7, 114.3, 101.1, 67.1, 58.1, 48.8, 45.0, 41.2, 39.3, 35.1, 34.1, 30.9, 30.5, 24.7.

**HRMS (ESI)** *m/z* calcd for $[C_{33}H_{37}F_3N_8O_2 + H]^+$ = 635.2992, found 635.3061.

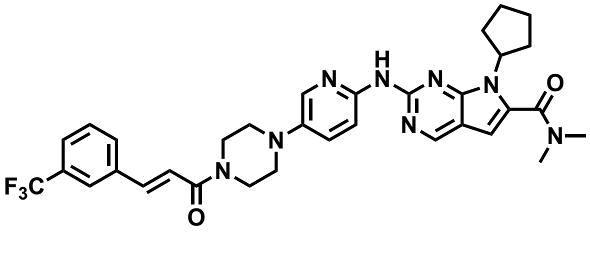

**EST1051**

**General Procedure A** was followed with 3-(trifluoromethyl)cinnamic acid (12.4 mg, 0.06 mmol), HATU (43.8 mg, 0.12 mmol), DIPEA (0.04 mL, 0.23 mmol), and ribociclib (25.0 mg, 0.06 mmol). The crude residue was purified by silica gel chromatography (0-10% MeOH in DCM) to afford 30.3 mg (83%) of the title compound as a yellow film.

**$^1$H NMR** (500 MHz, CDCl3) δ 8.65 (s, 1H), 8.32 (d, J = 9.0 Hz, 1H), 8.12 (s, 1H), 7.97 (d, J = 2.9 Hz, 1H), 7.72 (d, J = 2.1 Hz, 1H), 7.66 (d, J = 15.4 Hz, 1H), 7.62 (d, J = 7.7 Hz, 1H), 7.55 (d, J = 7.8 Hz, 1H), 7.45 (t, J = 7.8, 7.8 Hz, 1H), 7.29 (dd, J = 9.1, 3.0 Hz, 1H), 6.92 (d, J = 15.4 Hz, 1H), 6.38 (s, 1H), 4.73 (p, J = 9.0, 9.0, 8.9, 8.9 Hz, 1H), 3.91 - 3.76 (m, 4H), 3.13 (t, J = 5.0, 5.0 Hz, 4H), 3.09 (s, 6H), 2.57 - 2.46 (m, 2H), 2.05 - 1.95 (m, 4H), 1.69 - 1.62 (m, 2H).

**$^{13}$C NMR** (151 MHz, $CDCl_3$) δ 164.9, 164.1, 154.5, 151.9, 151.8, 147.6, 142.0, 141.5, 136.0, 132.1, 131.4 (q, $^2J_{CF}$ = 31.7 Hz), 131.3, 129.4, 127.5, 126.1 (q, $^3J_{CF}$ = 3.7 Hz), 123.9 (q, $^3J_{CF}$ = 3.8 Hz), 123.9 (q, $^1J_{CF}$ = 272.6 Hz), 118.7, 112.7, 112.5, 101.0, 57.9, 50.8, 50.4, 45.9, 42.2, 39.5, 35.2, 31.9, 30.2, 29.4, 24.7, 22.7, 14.1.

**HRMS (ESI)** *m/z* calcd for $[C_{33}H_{35}F_3N_8O_2 + H]^+$ = 633.2835, found 633.2906.

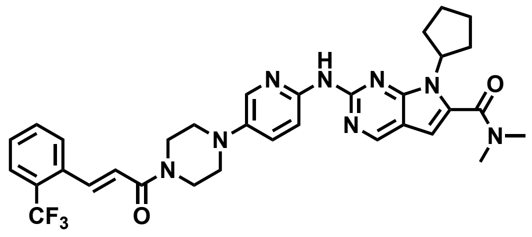

**EST1054**

**General Procedure A** was followed with *trans*-2-(trifluoromethyl)cinnamic acid (12.4 mg, 0.06 mmol), HATU (43.8 mg, 0.12 mmol), DIPEA (0.04 mL, 0.23 mmol), and ribociclib (25.0 mg, 0.06 mmol). The crude residue was purified by silica gel chromatography (0-10% MeOH in DCM) to afford 26.1 mg (72%) of the title compound as a yellow film.

**$^1$H NMR** (500 MHz, $CDCl_3$) δ 8.65 (s, 1H), 8.36 (d, *J* = 9.2 Hz, 1H), 7.96 – 7.87 (m, 2H), 7.66 – 7.60 (m, 2H), 7.50 (t, *J* = 7.6 Hz, 1H), 7.43 – 7.32 (m, 2H), 6.76 (d, *J* = 15.3 Hz, 1H), 6.39 (s, 1H), 4.73 (p, *J* = 8.7 Hz, 1H), 3.92 – 3.71 (m, 4H), 3.12 (t, *J* = 5.1 Hz, 4H), 3.08 (s, 6H), 2.48 (m, 2H), 2.06 – 1.95 (m, 4H), 1.71 – 1.59 (m, 2H).

**$^{13}$C NMR** (151 MHz, $CDCl_3$) δ 165.0, 163.9, 154.0, 151.9, 151.3, 147.1, 142.0, 138.4, 134.4, 132.6, 132.0, 129.1, 128.7 (q, $^2J_{CF}$ = 30.3 Hz), 127.9, 126.2 (q, $^3J_{CF}$ = 5.5 Hz), 124.0 (d, $^1J_{CF}$ = 274.0 Hz), 121.9, 113.2, 113.0, 100.9, 58.0, 31.9, 30.3, 24.8, 22.7, 14.1.

**HRMS (ESI)** *m/z* calcd for $[C_{33}H_{35}F_3N_8O_2 + H]^+$ = 633.2835, found 633.2906.

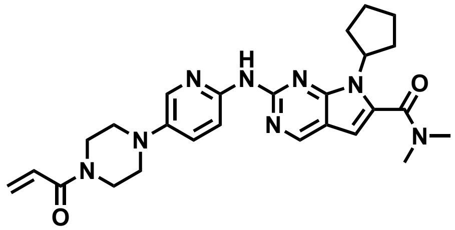

**EST1057**

**General Procedure A** was followed with acrylic acid (0.004 mL, 0.06 mmol), HATU (43.8 mg, 0.12 mmol), DIPEA (0.04 mL, 0.23 mmol), and ribociclib (25.0 mg, 0.06 mmol). The crude residue was purified by silica gel chromatography (0-10% MeOH in DCM) to afford 15.2 mg (54%) of the title compound as a yellow film.

**$^1$H NMR** (700 MHz, $CDCl_3$) δ 8.7 (s, 1H), 7.9 (br s, 2H), 7.6 – 7.4 (m, 1H), 6.6 (dd, *J* = 16.8, 10.6 Hz, 1H), 6.5 (s, 1H), 6.2 (d, *J* = 16.8 Hz, 1H), 5.7 (d, *J* = 10.6 Hz, 1H), 4.7 (p, *J* = 9.0 Hz, 1H), 3.8 (t, *J* = 5.2 Hz, 2H), 3.7 (q, *J* = 9.4 Hz, 2H), 3.1 – 3.1 (m, 8H), 3.0 – 2.9 (m, 2H), 2.4 (p, *J* = 8.3 Hz, 2H), 2.0 – 2.0 (m, 2H), 2.0 (q, *J* = 7.7 Hz, 2H), 1.6 (h, *J* = 8.0 Hz, 2H).

**$^{13}$C NMR** (126 MHz, $CDCl_3$) δ 165.9, 163.8, 151.9, 149.9, 145.9, 142.0, 128.7, 127.0, 113.8, 101.2, 58.1, 49.7, 49.3, 45.5, 41.7, 39.4, 35.2, 30.3, 29.6, 24.6.

**LRMS (ESI)** *m/z* calcd for $[C_{26}H_{32}N_8O_2 + H]^+$ = 489.3, found 489.4.

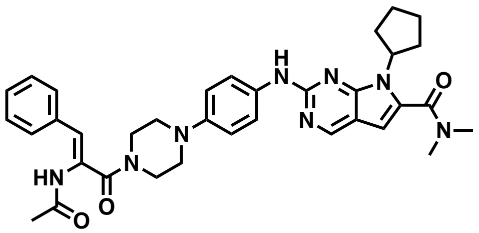

**EST1059**

**General Procedure A** was followed with 2-(acetylamino)-3-phenyl-2-propenoic acid (12.7 mg, 0.06 mmol), HATU (43.8 mg, 0.12 mmol), DIPEA (0.04 mL, 0.23 mmol), and ribociclib (25.0 mg, 0.06 mmol). The crude residue was purified by silica gel chromatography (0-10% MeOH in DCM) to afford 34.2 mg (96%) of the title compound as a yellow film.

**$^{1}$H NMR** (700 MHz, DMSO) δ 10.84 (s, 1H), 10.11 (s, 1H), 8.93 (s, 1H), 7.95 (d, *J* = 2.6 Hz, 2H), 7.75 (d, *J* = 8.9 Hz, 1H), 7.68 (d, *J* = 8.4 Hz, 2H), 7.63 (d, *J* = 8.3 Hz, 2H), 7.49 (d, *J* = 15.2 Hz, 1H), 7.23 (d, *J* = 15.3 Hz, 1H), 6.76 (s, 1H), 4.80 (p, *J* = 8.9 Hz, 1H), 3.91 (s, 2H), 3.77 (s, 2H), 3.22 (s, 4H), 3.06 (s, 6H), 2.40 – 2.31 (m, 2H), 2.07 (s, 3H), 2.05 – 2.00 (m, 2H), 1.99 – 1.93 (m, 2H), 1.66 (m, 2H).

**$^{13}$C NMR** (151 MHz, DMSO) δ 169.0, 165.2, 162.9, 153.0, 152.0, 149.7, 145.1, 142.4, 142.1, 141.1, 134.9, 130.2, 129.3, 119.3, 116.5, 115.0, 113.9, 101.4, 79.6, 57.5, 49.5, 49.1, 48.8, 45.1, 41.7, 39.2, 35.0, 30.5, 24.7, 24.5.

**LRMS (ESI)** *m/z* calcd for $[C_{35}H_{40}N_8O_3 + H]^+$ = 621.8, found 622.3.

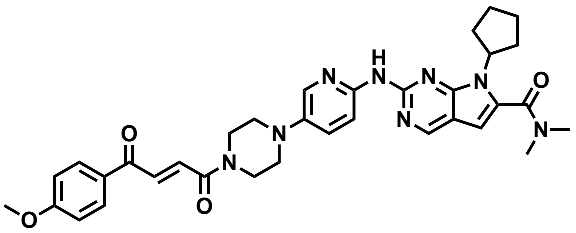

**EST1060**

**General Procedure D** was followed with *trans*-3-(4-methoxybenzoyl)acrylic acid (99.6 mg, 0.48 mmol), T3P (0.46 mL, 0.72 mmol), DIPEA (0.25 mL, 1.5 mmol), and ribociclib (216.0 mg, 0.57 mmol). The crude residue was purified by silica gel chromatography (0-10% MeOH in DCM). The resultant dark yellow oil was triturated in diethyl ether to afford 199.3 mg (83%) of the title compound as an orange powder.

**$^{1}$H NMR** (500 MHz, $CDCl_3$) δ 8.75 (s, 2H), 8.28 (d, *J* = 8.1 Hz, 1H), 8.06 – 8.01 (m, 3H), 7.98 (d, *J* = 14.9 Hz, 1H), 7.50 (d, *J* = 15.0 Hz, 1H), 7.36 (dd, *J* = 9.1, 2.9 Hz, 1H), 7.00 – 6.93 (m, 2H), 6.45 (s, 1H), 4.78 (p, *J* = 8.9 Hz, 1H), 3.92 (t, *J* = 5.2 Hz, 2H), 3.88 (s, 3H), 3.82 (t, *J* = 5.1 Hz, 2H), 3.17 (t, *J* = 5.2 Hz, 4H), 3.14 (s, 6H), 2.60 – 2.49 (m, 2H), 2.11 – 1.97 (m, 4H), 1.73 – 1.63 (m, 2H).

**$^{13}$C NMR** (126 MHz, $CDCl_3$) δ 187.6, 164.2, 164.1, 164.1, 154.7, 152.0, 151.9, 148.0, 141.8, 137.7, 134.7, 131.9, 131.3, 129.9, 127.4, 114.1, 112.6, 112.5, 101.1, 457.9, 55.6, 50.9, 50.3, 46.0, 42.2, 30.1, 24.7.

**HRMS (ESI)** *m/z* calcd for $[C_{34}H_{38}N_8O_4 + H]^+$ = 623.3016, found 623.3086.

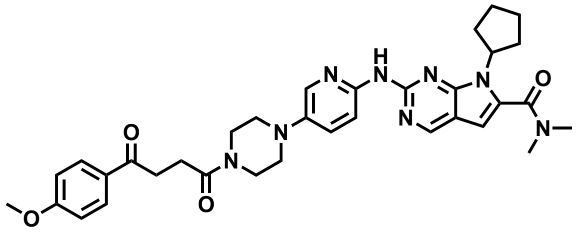

**JP-2-230**

A modified **General Procedure A** was followed with 3-(4-Methoxybenzoyl)propionic acid (15.9 mg, 0.08 mmol), HATU (25.5 mg, 0.07 mmol), DIPEA (0.04 mL, 0.23 mmol), and ribociclib (27.6 mg, 0.06 mmol). A precipitate had formed during the reaction which was vacuum filtered and washed with hexanes. The yellow precipitate was dissolved in DCM and dry loaded onto silica gel and was purified by silica gel chromatography (0-7% MeOH in DCM) to afford 28.9 mg (73%) of the title compound as a yellow-white powder.

**$^{1}$H NMR** (700 MHz, $CDCl_3$) δ 8.74 (s, 1H), 8.43 (s, 1H), 8.38 (d, *J* = 9.0 Hz, 1H), 8.07 (d, *J* = 2.9 Hz, 1H), 8.02 – 7.98 (m, 2H), 7.33 (dd, *J* = 9.1, 2.9 Hz, 1H), 6.95 – 6.91 (m, 2H), 6.44 (s, 1H), 4.79 (p, *J* = 8.9 Hz, 1H), 3.86 (s, 3H), 3.81 (t, *J* = 5.1 Hz, 2H), 3.76 (t, *J* = 5.1 Hz, 2H), 3.34 (t, *J* = 6.6 Hz, 2H), 3.18 (t, *J* = 5.1 Hz, 2H), 3.15 (s, 6H), 3.11 (t, *J* = 5.2 Hz, 2H), 2.82 (t, *J* = 6.6 Hz, 2H), 2.62 – 2.54 (m, 2H), 2.11 – 2.00 (m, 4H), 1.75 – 1.67 (m, 2H).

**$^{13}$C NMR** (151 MHz, $CDCl_3$) δ 197.6, 170.4, 164.1, 163.5, 154.6, 152.0, 151.9, 147.6, 142.1, 137.5, 132.0, 130.4, 129.9, 127.3, 113.7, 112.6, 112.4, 101.0, 57.9, 55.5, 50.6, 50.3, 45.4, 41.7, 33.2, 30.2, 27.1, 24.7.

**HRMS (ESI)** *m/z* calcd for $[C_{34}H_{40}N_8O_4 + H]^+$ = 625.3173, found 625.3242.

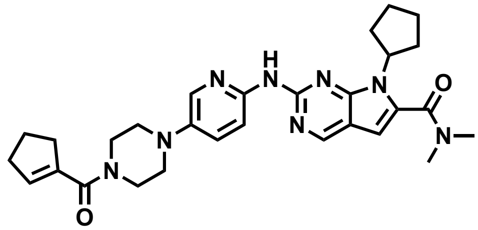

**EST1061**

**General Procedure A** was followed with 1-cyclopentenecarboxylic acid (6.5 mg, 0.06 mmol), HATU (43.8 mg, 0.12 mmol), DIPEA (0.04 mL, 0.23 mmol), and ribociclib (25.0 mg, 0.06 mmol). The crude residue was purified by silica gel chromatography (0-10% MeOH in DCM) to afford 23.3 mg (77%) of the title compound as a yellow powder.

**$^1$H NMR** (500 MHz, $CDCl_3$) δ 8.75 (s, 1H), 8.02 (s, 1H), 7.94 (d, *J* = 2.9 Hz, 1H), 7.50 – 7.42 (m, 1H), 6.47 (s, 1H), 5.93 (p, *J* = 2.2 Hz, 1H), 4.77 (p, *J* = 8.9 Hz, 1H), 3.78 (s, 4H), 3.21 – 3.08 (m, 10H), 2.63 (tq, *J* = 7.5, 2.4 Hz, 2H), 2.54 – 2.42 (m, 4H), 2.12 – 1.99 (m, 4H), 1.96 (p, *J* = 7.6 Hz, 2H), 1.67 (h, *J* = 8.4 Hz, 2H).

**$^{13}$C NMR** (151 MHz, $CDCl_3$) δ 168.6, 163.6, 151.9, 142.2, 137.9, 133.4, 113.6, 101.1, 58.0, 54.0, 49.8, 39.4, 35.2, 34.5, 33.3, 30.4, 24.7, 22.8, 18.6, 17.3, 14.1.

**LRMS (ESI)** *m/z* calcd for $[C_{29}H_{36}N_8O_2 + H]^+$ = 529.3, found 529.4.

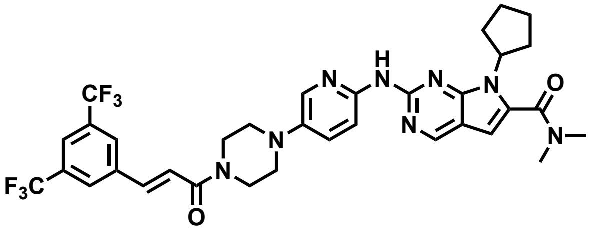

**KN1002**

**General Procedure A** was followed with *trans*-3,5-bis(trifluoromethyl)cinnamic acid (14.6 mg, 0.06 mmol), HATU (43.8 mg, 0.12 mmol), DIPEA (0.04 mL, 0.23 mmol), and ribociclib (25.0 mg, 0.06 mmol). The crude residue was purified by silica gel chromatography (0-10% MeOH in DCM) to afford 14.5 mg (40%) of the title compound as a yellow film.

**$^1$H NMR** (500 MHz, $CDCl_3$) δ 8.65 (s, 1H), 8.28 (s, 1H), 7.97 (d, *J* = 3.0 Hz, 2H), 7.87 (s, 2H), 7.79 (s, 1H), 7.68 (d, *J* = 15.4 Hz, 1H), 7.30 (dd, *J* = 9.2, 2.9 Hz, 1H), 6.99 (d, *J* = 15.4 Hz, 1H), 6.38 (s, 1H), 4.73 (t, *J* = 8.9 Hz, 1H), 3.88 (s, 2H), 3.82 (d, *J* = 13.7 Hz, 2H), 3.13 (d, *J* = 8.6 Hz, 4H), 3.09 (s, 6H), 2.50 (dq, *J* = 15.6, 7.3 Hz, 2H), 2.06 – 1.92 (m, 4H), 1.65 (dt, *J* = 12.6, 5.1 Hz, 2H).

**$^{13}$C NMR** (151 MHz, $CDCl_3$) δ 164.3, 164.0, 151.9, 142.0, 139.8, 137.3, 132.4 (q, $^{2}J_{CF}$ = 33.4 Hz), 127.4, 123.1 (q, $^{1}J_{CF}$ = 273.0 Hz), 122.9, 120.8, 112.9, 112.6, 101.0, 57.9, 50.3, 45.9, 42.3, 31.9, 30.2, 29.4, 24.7, 22.7, 14.1.

**LRMS (ESI)** *m/z* calcd for $[C_{34}H_{34}F_6N_8O_2 + H]^+$ = 701.3, found 701.4.

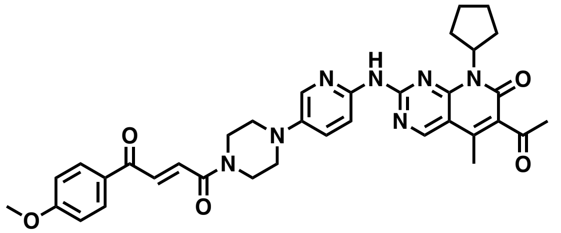

**EST1089**

**General Procedure A** was followed with 4-methoxycinnamic acid (13.8 mg, 0.06 mmol), HATU (42.5 mg, 0.11 mmol), DIPEA (0.04 mL, 0.23 mmol), and palbociclib (25.0 mg, 0.06 mmol). The crude residue was purified by silica gel chromatography (0-10% MeOH in DCM) to afford 26.1 mg (74%) of the title compound as a yellow film.

**$^{1}$H NMR** (500 MHz, $CDCl_3$) δ 8.86 (s, 1H), 8.22 (d, *J* = 9.0 Hz, 1H), 8.09 (d, *J* = 2.9 Hz, 1H), 8.08 – 8.02 (m, 2H), 8.00 (d, *J* = 14.9 Hz, 1H), 7.52 (d, *J* = 14.8 Hz, 1H), 7.35 (dd, *J* = 9.1, 3.0 Hz, 1H), 7.01 – 6.95 (m, 2H), 5.88 (p, *J* = 8.9 Hz, 1H), 3.94 (t, *J* = 5.2 Hz, 2H), 3.89 (s, 3H), 3.85 (t, *J* = 5.1 Hz, 2H), 3.22 (t, *J* = 5.2 Hz, 4H), 2.54 (s, 3H), 2.38 (s, 3H), 2.36 – 2.30 (m, 1H), 2.12 – 2.01 (m, 2H), 1.89 (dddd, *J* = 15.4, 13.0, 7.6, 4.7 Hz, 2H), 1.75 – 1.64 (m, 3H).

**$^{13}$C NMR** (151 MHz, $CDCl_3$) δ 202.6, 187.6, 164.3, 164.1, 161.4, 158.0, 157.2, 155.5, 145.8, 143.0, 141.7, 137.3, 134.9, 131.3, 131.1, 130.9, 129.9, 126.9, 114.2, 113.6, 107.9, 55.6, 54.1, 53.4, 50.4, 49.8, 45.8, 42.1, 31.5, 28.1, 25.8, 22.7, 14.0.

**HRMS (ESI)** *m/z* calcd for $[C_{35}H_{37}N_7O_5 + H]^+$ = 636.2929, found 636.2935.

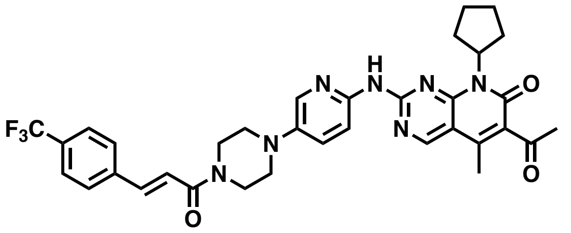

**EST1090**

**General Procedure A** was followed with *trans*-4-(trifluoromethyl)cinnamic acid (14.5 mg, 0.07 mmol), HATU (42.5 mg, 0.11 mmol), DIPEA (0.04 mL, 0.23 mmol), and palbociclib (0.25, 0.06 mmol). The crude residue was purified by silica gel chromatography (0-100% EtOAc in Hexanes) to afford 22.8 mg (63%) of the title compound as a yellow film.

**$^{1}$H NMR** (500 MHz, $CDCl_3$) δ 8.87 (s, 1H), 8.74 (s, 1H), 8.22 (d, *J* = 9.1 Hz, 1H), 8.09 (d, *J* = 2.9 Hz, 1H), 7.72 (d, *J* = 15.4 Hz, 1H), 7.64 (s, 4H), 7.37 (dd, *J* = 9.1, 3.0 Hz, 1H), 7.00 (d, *J* = 15.5 Hz, 1H), 5.88 (p, *J* = 8.9 Hz, 1H), 4.01 – 3.81 (m, 4H), 3.23 (t, *J* = 5.1 Hz, 4H), 2.54 (s, 3H), 2.38 (s, 3H), 2.37 – 2.31 (m, 2H), 2.07 (tdd, *J* = 11.9, 9.9, 5.2 Hz, 2H), 1.92 – 1.84 (m, 2H), 1.69 (tdd, *J* = 10.9, 6.6, 4.0 Hz, 2H).

**$^{13}$C NMR** (126 MHz, $CDCl_3$) δ 202.6, 164.9, 161.4, 158.0, 157.2, 155.6, 145.8, 143.1, 141.8, 141.6, 138.5 (d, $^{5}J_{CF}$ = 1.4 Hz) , 137.1, 131.4 (q, $^{2}J_{CF}$ = 32.7 Hz), 130.9, 128.0, 126.9, 125.8 (q, $^{3}J_{CF}$ = 3.8 Hz), 123.9 (d, $^{1}J_{CF}$ = 272.1 Hz), 119.2, 113.7, 107.9, 54.1, 50.3, 49.8, 45.7, 42.1, 31.5, 28.1, 25.8, 14.0.

**HRMS (ESI)** *m/z* calcd for $[C_{34}H_{34}F_3N_7O_3 + H]^+$ = 646.2748, found 646.2741.

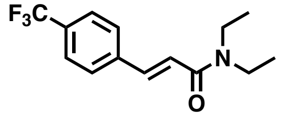


**KN1026**

**General Procedure A** was followed with *trans*-4-(trifluoromethyl)cinnamic acid (100.0 mg, 0.06 mmol), HATU (43.8 mg, 0.12 mmol), DIPEA (0.04 mL, 0.23 mmol), and *N,N*-diethylamine (0.05, 0.48 mmol). The crude residue was purified by silica gel chromatography (0-60% EtOAC in Hexanes) to afford 66.0 mg (50%) of the title compound as a yellow film.

**$^{1}$H NMR** (500 MHz, $CDCl_3$) δ 7.63 (d, *J* = 15.4 Hz, 1H), 7.54 (s, 4H), 6.83 (d, *J* = 15.4 Hz, 1H), 3.42 (dq, *J* = 9.8, 7.1 Hz, 4H), 1.19 (t, *J* = 7.2 Hz, 3H), 1.12 (t, *J* = 7.1 Hz, 3H).

**$^{13}$C NMR** (126 MHz, $CDCl_3$) δ 165.1, 140.5, 138.9 (d, $^{5}J_{CF}$ = 1.4 Hz), 131.0 (q, $^{2}J_{CF}$ = 32.5 Hz), 127.9, 125.7 (q, $^{3}J_{CF}$ = 3.8 Hz), 123.9 (q, $^{1}J_{CF}$ = 272.1 Hz), 120.3, 42.4, 41.2, 15.1, 13.1.

**LRMS (ESI)** *m/z* calcd for $[C_{14}H_{16}F_3NO + H]^+$ = 272.1, found 272.2.

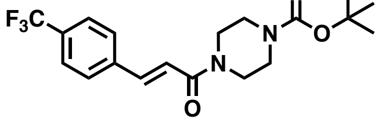


**EST1096**

**General Procedure A** was followed with *trans*-4-(trifluoromethyl)cinnamic acid (100.0 mg, 0.46 mmol), HATU (43.8 mg, 0.12 mmol), DIPEA (0.04 mL, 0.23 mmol), and 1-boc-piperazine (94.7 mg, 0.51 mmol). The crude residue was purified by silica gel chromatography (0-70% EtOAc in Hexanes) to afford 72.5 mg (41%) of the title compound as a white powder.

**$^1$H NMR** (500 MHz, $CDCl_3$) δ 7.62 (d, *J* = 15.4 Hz, 1H), 7.55 (s, 4H), 6.87 (d, *J* = 15.4 Hz, 1H), 3.60 (s, 4H), 3.42 (dd, *J* = 6.6, 3.9 Hz, 4H), 1.41 (s, 9H).

**$^{13}$C NMR** (151 MHz, $CDCl_3$) δ 165.0, 154.5, 141.4, 138.5, 131.3 (q, $^2J_{CF}$ = 32.6 Hz), 127.9, 125.8 (q, $^3J_{CF}$ = 3.7 Hz), 123.9 (q, $^1J_{CF}$ = 272.3 Hz), 119.3, 80.4, 45.7, 42.0, 28.4.

**LRMS (ESI)** *m/z* calcd for $[C_{19}H_{23}F_3N_2O_3 + Na]^+$ = 407.2, found 407.2.

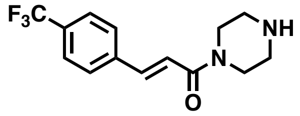


**EST1102**

**General Procedure E** was followed with **EST1096** (47.3 mg, 0.12 mmol), and TFA (0.3 mL, 3.94 mmol) for 2 hours. The crude residue was purified by silica gel chromatography (0-17% MeOH in DCM) using a Biotage® Sfär KP-Amino D cartridge to afford 14.5 mg (41%) of the title compound as a white film.

**$^1$H NMR** (500 MHz, $CDCl_3$) δ 7.64 (d, *J* = 15.5 Hz, 1H), 7.59 (s, 4H), 6.93 (d, *J* = 15.4 Hz, 1H), 3.72 – 3.66 (m, 2H), 3.64 – 3.58 (m, 2H), 2.89 (t, *J* = 5.0, 5.0 Hz, 4H), 1.80 (s, 1H).

**$^{13}$C NMR** (151 MHz, $CDCl_3$) δ 164.9, 140.8, 138.7, 131.1 (q, $^2J_{CF}$ = 32.6 Hz), 127.8, 125.7 (q, $^3J_{CF}$ = 3.8 Hz), 123.9 (q, $^1J_{CF}$ = 272.0 Hz), 119.8, 47.2, 46.6, 45.9, 43.4.

**LRMS (ESI)** *m/z* calcd for $[C_{14}H_{15}F_3N_2O + H]+$ = 285.1, found 285.2.

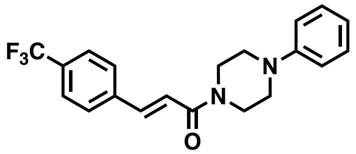


**KN1023**

**General Procedure A** was followed with *trans*-4-(trifluoromethyl)cinnamic acid (200.0 mg, 0.93 mmol), HATU (703.6 mg, 1.85 mmol), DIPEA (0.64 mL, 3.7 mmol), and 1-phenylpiperazine (150.1 mg, 0.93 mmol). The crude residue was purified by silica gel chromatography (15-55% EtOAc in Hexanes) to afford 125.0 mg (37%) of the title compound as a yellow powder.

**$^1$H NMR** (500 MHz, $CDCl_3$) δ 7.62 (d, *J* = 15.4 Hz, 1H), 7.53 (s, 4H), 7.23 – 7.16 (m, 2H), 6.91 (d, *J* = 15.4 Hz, 1H), 6.83 (dd, *J* = 16.5, 7.9 Hz, 3H), 3.80 (br s, 2H), 3.72 (br s, 2H), 3.13 (m, 4H).

**$^{13}$C NMR** (126 MHz, $CDCl_3$) δ 164.9, 150.9, 141.3, 138.7, 131.2 (q, $^{2}J_{CF}$ = 32.5 Hz), 129.3, 128.0, 125.8 (q, $^{3}J_{CF}$ = 3.8 Hz), 124.0 (q, $^{1}J_{CF}$ = 272.2 Hz), 120.6, 119.6, 116.7, 49.9, 49.4, 45.9, 42.2.

**LRMS (ESI)** *m/z* calcd for $[C_{20}H_{19}F_3N_2O + H]^+$ = 361.1, found 361.2.

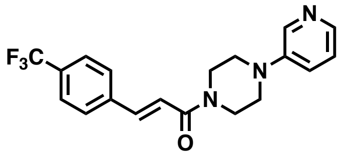

**KN1017**

**General Procedure A** was followed with *trans*-4-(trifluoromethyl)cinnamic acid (200.0 mg, 0.93 mmol), HATU (703.6 mg, 1.85 mmol), DIPEA (0.64 mL, 3.7 mmol), and 1-(3-pyridinyl)piperazine (151.1 mg, 0.93 mmol). The crude residue was purified by silica gel chromatography (0-10% MeOH in DCM) to afford 53.1 mg (16%) of the title compound as a yellow powder.

**$^{1}$H NMR** (500 MHz, $CDCl_3$) δ 8.34 – 8.29 (m, 1H), 8.14 (dd, *J* = 3.9, 2.1 Hz, 1H), 7.70 (d, *J* = 15.4 Hz, 1H), 7.61 (s, 4H), 7.23 – 7.14 (m, 2H), 6.98 (d, *J* = 15.5 Hz, 1H), 3.90 (br s, 2H), 3.83 (br s, 2H), 3.25 (m, 4H).

**$^{13}$C NMR** (126 MHz, $CDCl_3$) δ 164.9, 146.5, 141.7, 141.5, 139.1, 138.5, 131.3 (q, $^{2}J_{CF}$ = 32.6 Hz), 128.0, 125.8 (q, $^{3}J_{CF}$ = 3.8 Hz), 123.9 (q, $^{1}J_{CF}$ = 272.2 Hz), 123.6, 123.1, 119.2, 49.2, 48.7, 45.6, 41.9.

**LRMS (ESI)** *m/z* calcd for $[C_{19}H_{18}F_3N_3O + H]^+$ = 362.1, found 362.1.

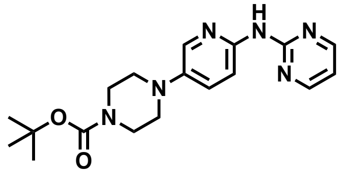

**JP-2-195**

A combination of 2-chloropyrimidine (100.0 mg, 0.95 mmol), tert-butyl 4-(6-aminopyridin-3-yl)piperazine-1-carboxylate (265.2 mg, 0.95 mmol), palladium (II) acetate (21.4 mg, 0.10 mmol), caesium carbonate (465.6 mg, 1.4 mmol), and Xantphos (82.7 mg, 0.14 mmol) was suspended in dioxane (10 mL) and heated to 120 °C overnight. The volatiles were removed *in vacuo*, and the crude residue was purified by silica gel chromatography (0-5% MeOH in DCM) to afford 282.1 mg (91%) of the title compound as a beige solid.

**$^{1}$H NMR** (700 MHz, $CDCl_3$) δ 8.50 (d, *J* = 4.8 Hz, 2H), 8.30 (d, *J* = 9.0 Hz, 1H), 8.19 (s, 1H), 8.07 (d, *J* = 2.9 Hz, 1H), 7.35 (dd, *J* = 9.1, 3.0 Hz, 1H), 6.78 (t, *J* = 4.8 Hz, 1H), 3.63 (t, *J* = 5.1 Hz, 4H), 3.11 (t, *J* = 5.2 Hz, 4H), 1.51 (s, 9H).

**$^{13}$C NMR** (151 MHz, $CDCl_3$) δ 159.3, 158.0, 154.7, 146.6, 143.0, 137.4, 127.2, 113.0, 112.9, 80.0, 50.1, 28.4.

**LRMS (ESI)** *m/z* calcd for $[C_{18}H_{24}N_6O_2 + H]^+$ = 375.2, found 375.3.

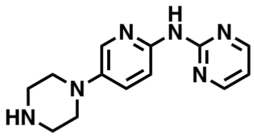

**JP-2-198**

**General Procedure E** was followed with **JP-2-195** (152.4 mg, 0.43 mmol), and TFA (1.1 mL, 13.7 mmol) for 2 hours. The crude material was used without further purification.

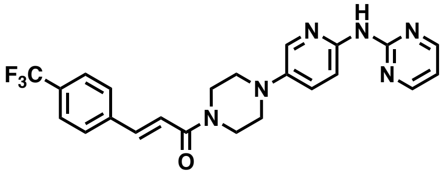

**JP-2-200**

**General Procedure A** was followed with *trans*-4-(trifluoromethyl)cinnamic acid (43.0 mg, 0.20 mmol), HATU (82.6 mg, 0.22 mmol), DIPEA (0.09 mL, 0.54 mmol), and **JP-2-198** (46.4 mg, 0.18 mmol). The crude residue was purified by silica gel chromatography (50-100% EtOAc in Hexanes followed by 0-5% MeOH in DCM) to afford 13.3 mg (16%, 2 steps) of the title compound as a yellow residue.

**$^1$H NMR** (700 MHz, $CDCl_3$) δ 8.48 (dd, *J* = 4.8, 1.6 Hz, 2H), 8.30 (d, *J* = 8.9 Hz, 1H), 8.14 (s, 1H), 8.05 (t, *J* = 2.2 Hz, 1H), 7.72 (dd, *J* = 15.5, 1.7 Hz, 1H), 7.64 (d, *J* = 1.7 Hz, 3H), 7.35 (dt, *J* = 9.1, 2.3 Hz, 1H), 7.00 (dd, *J* = 15.5, 1.6 Hz, 1H), 6.77 (td, *J* = 4.8, 1.7 Hz, 1H), 3.94 (br s, 2H), 3.85 (br s, 2H), 3.19 (m, 3H).

**$^{13}$C NMR** (126 MHz, $CDCl_3$) δ 164.9, 159.2, 158.0, 147.0, 142.5, 141.5, 138.6 (d, $^5J_{CF}$ = 1.3 Hz), 137.5, 131.3 (q, $^2J_{CF}$ = 32.8), 127.9, 127.4, 125.8 (q, $^3J_{CF}$ = 3.8 Hz), 123.9 (q, $^1J_{CF}$ = 272.2 Hz), 119.3, 113.1, 113.0, 50.7, 50.1, 45.8, 42.1, 29.7, 28.4.

**LRMS (ESI)** *m/z* calcd for $[C_{23}H_{21}F_3N_6O + H]^+$ = 455.2, found 455.2.

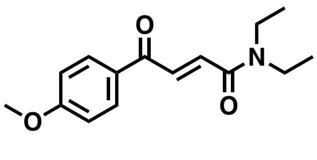

**KN1025**

**General Procedure A** was followed with *trans*-3-(4-methoxybenzoyl)acrylic acid (100.1 mg, 0.47 mmol), HATU (175.9 mg, 0.48 mmol), DIPEA (0.08 mL, 0.46 mmol), and *N,N*-diethylamine (25.0 mg, 0.06 mmol). The crude residue was purified by silica gel chromatography (0-55% EtOAc in Hexanes) to afford 16.3 mg (13%) of the title compound as a yellow film.

**$^1$H NMR** (700 MHz, $CDCl_3$) δ 8.06 – 8.04 (m, 2H), 7.99 (d, $J$ = 14.8 Hz, 1H), 7.43 (d, $J$ = 14.8 Hz, 1H), 6.99 – 6.95 (m, 2H), 3.89 (s, 3H), 3.52 – 3.45 (m, 4H), 1.24 (t, $J$ = 7.2 Hz, 3H), 1.19 (t, $J$ = 7.1 Hz, 3H).

**$^{13}$C NMR** (126 MHz, $CDCl_3$) δ 187.9, 164.5, 164.1, 134.0, 132.2, 131.3, 131.0, 130.1, 114.1, 55.6, 42.5, 41.2, 15.1, 13.0.

**LRMS (ESI)** *m/z* calcd for $[C_{15}H_{19}NO_3 + H]^+$ = 262.1, found 262.3.

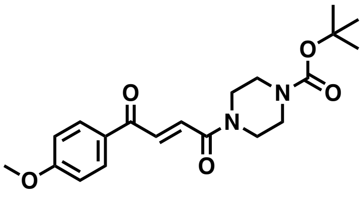

**JP-2-190**

**General Procedure A** was followed with *trans*-3-(4-methoxybenzoyl)acrylic acid (977 mg, 7.3 mmol), HATU (3.1 g, 8.1 mmol), DIPEA (3.8 mL, 21.8 mmol), and 1-boc-piperazine (1.5 g, 8.0 mmol). The crude residue was purified by silica gel chromatography (0-50% EtOAC in Hexanes) to afford 1.59 g (58%) of the title compound as a yellow solid.

**$^1$H NMR** (500 MHz, $CDCl_3$) δ 8.04 (d, $J$ = 8.6 Hz, 2H), 7.96 (dd, $J$ = 15.4, 3.9 Hz, 1H), 7.46 (d, $J$ = 14.9 Hz, 1H), 6.98 (d, $J$ = 8.7 Hz, 2H), 3.89 (s, 3H), 3.72 (t, $J$ = 5.4, 5.4 Hz, 2H), 3.62 (t, $J$ = 5.2, 5.2 Hz, 2H), 3.49 (t, $J$ = 5.2, 5.2 Hz, 4H), 1.48 (s, 9H).

**$^{13}$C NMR** (126 MHz, $CDCl_3$) δ 187.6, 164.2, 154.5, 134.8, 131.3, 131.2, 131.1, 129.9, 114.1, 114.0, 80.5, 55.6, 45.9, 42.1, 28.4.

**LRMS (ESI)** *m/z* calcd for $[C_{20}H_{26}N_2O_5 + Na]^+$ = 397.2, found 397.2.

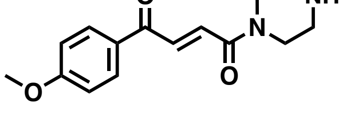

**JP-2-196**

**General Procedure E** was followed with **JP-2-190** (78.6 mg, 0.21 mmol), and TFA (0.51 mL, 6.7 mmol) for 1.5 hours. The reaction mixture was concentrated *in vacuo*, redissolved in DCM, and stirred with a saturated solution of $NaHCO_3$ (2 mL) for 30 min. The organic layer was separated, washed with brine, dried over

$Na_2SO_4$, and concentrated *in vacuo*. The crude residue was purified by silica gel chromatography (0-12% MeOH in DCM) to afford 20.8 mg (81%) of the title compound as a white solid.

**$^1$H NMR** (700 MHz, $CDCl_3$) δ 7.99 (dd, *J* = 8.9, 1.9 Hz, 2H), 7.93 (dd, *J* = 14.9, 1.8 Hz, 1H), 7.37 (dd, *J* = 14.9, 1.8 Hz, 1H), 6.94 (dd, *J* = 9.0, 2.0 Hz, 2H), 3.97 (t, *J* = 5.4 Hz, 2H), 3.92 (t, *J* = 5.4 Hz, 2H), 3.85 (d, *J* = 1.9 Hz, 3H), 3.22 (t, *J* = 5.2 Hz, 4H), 2.73 (s, 1H).

**$^{13}$C NMR** (151 MHz, $CDCl_3$) δ 187.5, 164.5, 136.0, 131.5, 130.0, 129.7, 114.3, 55.7, 43.4, 43.1, 42.8, 38.9.

**LRMS (ESI)** *m/z* calcd for $[C_{15}H_{18}N_2O_3 + H]^+$ = 275.1, found 275.1.

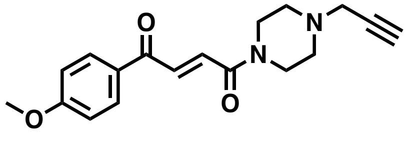

**JP-2-253**

A combination of **JP-2-196** (25.9 mg, 0.094 mmol) and potassium carbonate (15.7 mg, 0.11 mmol) was suspended in DMF (4 mL) and stirred at ambient temperature for 5 minutes. Propargyl bromide (0.01 mL, 0.13 mmol) was added, and the reaction mixture was heated to 90 °C overnight. The reaction mixture was concentrated *in vacuo*, and the crude residue was purified by silica gel chromatography (0-10% MeOH in DCM) to afford 11.7 mg (40%) of the title compound as a yellow oil.

**$^1$H NMR** (700 MHz, $CDCl_3$) δ 8.05 – 8.02 (m, 2H), 7.95 (d, *J* = 14.9 Hz, 1H), 7.47 (d, *J* = 14.9 Hz, 1H), 6.99 – 6.95 (m, 2H), 3.89 (s, 3H), 3.79 (t, *J* = 5.1 Hz, 2H), 3.68 (t, *J* = 5.1 Hz, 2H), 3.35 (d, *J* = 2.4 Hz, 2H), 2.60 (q, *J* = 4.6 Hz, 4H), 2.28 (d, *J* = 2.4 Hz, 1H).

**$^{13}$C NMR** (151 MHz, $CDCl_3$) δ 187.7, 164.2, 164.0, 134.4, 131.5, 131.3, 130.0, 114.1, 78.0, 73.8, 55.6, 51.9, 51.3, 46.8, 45.9, 42.1.

**LRMS (ESI)** *m/z* calcd for $[C_{18}H_{20}N_2O_3 + H]^+$ = 213.1, found 213.2.

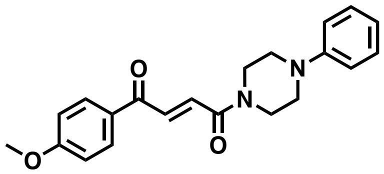

**KN1021**

**General Procedure A** was followed with *trans*-3-(4-methoxybenzoyl)acrylic acid (201.2 mg, 0.07 mmol), HATU (737.6 mg, 1.94 mmol), DIPEA (0.68 mL, 3.88 mmol), and 1-phenylpiperazine (25.0 mg, 0.06 mmol). The crude residue was purified by silica gel chromatography (15-55% EtOAc in Hexanes) to afford 255.6 mg (75%) of the title compound as an orange film.

**$^1$H NMR** (500 MHz, $CDCl_3$) δ 8.05 (d, *J* = 8.6 Hz, 2H), 7.99 (d, *J* = 14.9 Hz, 1H), 7.52 (d, *J* = 14.9 Hz, 1H), 7.30 (t, *J* = 7.7 Hz, 2H), 7.01 – 6.89 (m, 5H), 3.92 (m, 2H), 3.90 (s, 3H), 3.81 (t, *J* = 5.1 Hz, 2H), 3.23 (m, 4H).

**$^{13}$C NMR** (126 MHz, $CDCl_3$) δ 187.7, 164.2, 164.1, 150.8, 134.6, 131.4, 131.3, 130.0, 129.3, 120.8, 116.8, 114.1, 55.6, 50.0, 49.4, 46.0, 42.2.

**LRMS (ESI)** *m/z* calcd for $[C_{21}H_{22}N_2O_3 + H]^+$ = 351.2, found 351.1.

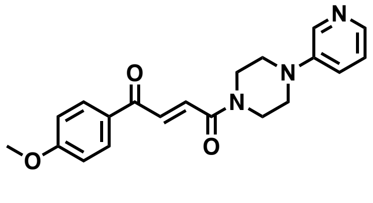


**KN1018**

**General Procedure A** was followed with *trans*-3-(4-methoxybenzoyl)acrylic acid (200.2 mg, 0.97 mmol), HATU (740.2 mg, 1.96 mmol), DIPEA (0.68 mL, 3.90 mmol), and 1-(3-pyridinyl)piperazine (158.3 mg, 0.97 mmol). The crude residue was purified by silica gel chromatography (0-8% MeOH in DCM) to afford 40.4 mg (12%) of the title compound as a yellow film.

**$^1$H NMR** (500 MHz, $CDCl_3$) δ 8.33 – 8.27 (m, 1H), 8.14 (dd, *J* = 3.8, 2.2 Hz, 1H), 8.06 – 8.00 (m, 2H), 7.97 (d, *J* = 14.9 Hz, 1H), 7.49 (d, *J* = 14.8 Hz, 1H), 7.22 – 7.14 (m, 2H), 6.99 – 6.92 (m, 2H), 3.90 (m, 2H), 3.86 (s, 3H), 3.80 (t, *J* = 5.2 Hz, 2H), 3.27 – 3.21 (m, 4H).

**$^{13}$C NMR** (126 MHz, $CDCl_3$) δ 187.6, 164.2, 164.1, 146.5, 141.7, 139.2, 134.8, 131.3, 131.1, 129.9, 123.6, 123.1, 114.1, 55.6, 49.2, 48.7, 45.7, 42.0.

**LRMS (ESI)** *m/z* calcd for $[C_{20}H_{21}N_3O_3 + H]^+$ = 352.2, found 352.1.

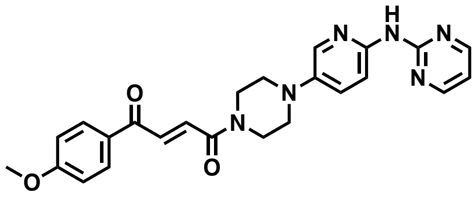


**JP-2-199**

**General Procedure A** was followed with *trans*-3-(4-methoxybenzoyl)acrylic acid (39.2 mg, 0.19 mmol), HATU (78.9 mg, 0.21 mmol), DIPEA (0.09 mL, 0.52 mmol), and **JP-2-198** (44.3 mg, 0.17 mmol). The crude residue was purified by silica gel chromatography (50-100% EtOAc in Hexanes followed by 0-5% MeOH in DCM) to afford 37.9 mg (49%) of the title compound as a yellow residue.

**$^1$H NMR** (700 MHz, $CDCl_3$) δ 8.77 (s, 1H), 8.50 (d, *J* = 4.8 Hz, 2H), 8.31 (d, *J* = 9.1 Hz, 1H), 8.10 (d, *J* = 2.9 Hz, 1H), 8.05 (d, *J* = 8.4 Hz, 2H), 7.99 (d, *J* = 14.8 Hz, 1H), 7.52 (d, *J* = 14.6 Hz, 1H), 7.34 (dd, *J* = 9.0, 2.9 Hz, 1H), 6.98 (d, *J* = 8.4 Hz, 2H), 6.76 (t, *J* = 4.9 Hz, 1H), 3.93 (t, *J* = 5.4 Hz, 2H), 3.89 (s, 3H), 3.83 (t, *J* = 5.1 Hz, 2H), 3.18 (q, *J* = 4.5 Hz, 4H).

**$^{13}$C NMR** (126 MHz, $CDCl_3$) δ 187.6, 164.2, 164.1, 159.3, 158.0, 147.2, 142.3, 137.5, 134.7, 131.3, 131.3, 130.0, 127.5, 114.1, 113.2, 112.9, 55.6, 50.7, 50.1, 45.9, 42.2, 29.7, 28.4.

**LRMS (ESI)** *m/z* calcd for $[C_{24}H_{24}N_6O_3 + H]^+$ = 445.2, found 445.2.

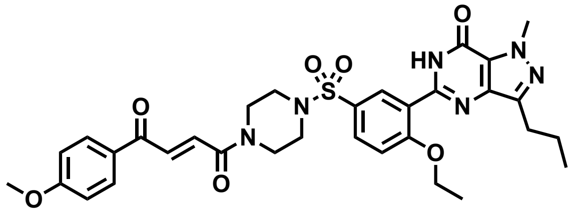

**JP-2-201**

The secondary amine **JP-2-196** (23.2 mg, 0.08 mmol) was dissolved in DCM (1 mL). DIPEA (0.1 mL, 0.5 mmol) was added, and the reaction mixture was stirred at ambient temperature for 5 minutes. A solution of 5-(5-Chlorosulfonyl-2-ethoxyphenyl)-1-methyl-3-propyl-1,6-dihydro-7H-pyrazolo[4,3-d]pyrimidin-7-one (34.7 mg, 0.08 mmol) in DCM (0.5 mL) was added dropwise to the reaction mixture and allowed to stir at ambient temperature for 2 hours. The volatiles were removed *in vacuo*, and the crude residue was purified by silica gel chromatography (25-100% EtOAc in Hexanes) to afford 49.5 mg (90%) of the title compound as a yellow oil.

**$^1$H NMR** (500 MHz, $CDCl_3$) δ 10.73 (s, 1H), 8.73 (d, *J* = 2.4 Hz, 1H), 7.92 – 7.89 (m, 2H), 7.82 (d, *J* = 14.9 Hz, 1H), 7.75 (dd, *J* = 8.7, 2.4 Hz, 1H), 7.27 (d, *J* = 14.8 Hz, 1H), 7.09 (d, *J* = 8.8 Hz, 1H), 6.89 – 6.84 (m, 2H), 4.30 (q, *J* = 7.0 Hz, 2H), 4.19 (s, 3H), 3.80 (s, 3H), 3.79 – 3.76 (m, 2H), 3.68 (t, *J* = 5.0 Hz, 2H), 3.05 (q, *J* = 5.4 Hz, 4H), 2.86 (t, *J* = 7.6 Hz, 2H), 1.79 (h, *J* = 7.4 Hz, 2H), 1.57 (t, *J* = 7.0 Hz, 3H), 0.96 (t, *J* = 7.4 Hz, 3H).

**$^{13}$C NMR** (126 MHz, $CDCl_3$) δ 187.3, 164.3, 164.0, 159.6, 153.6, 147.0, 146.2, 138.3, 135.2, 131.5, 131.3, 131.1, 130.5, 129.7, 128.5, 124.5, 121.4, 114.1, 113.3, 77.3, 66.2, 55.6, 46.4, 45.8, 45.4, 41.6, 38.2, 27.7, 22.3, 14.5, 14.1.

**HRMS (ESI)** *m/z* calcd for $[C_{32}H_{36}N_6O_7S + H]^+$ = 649.2366, found 649.2450.

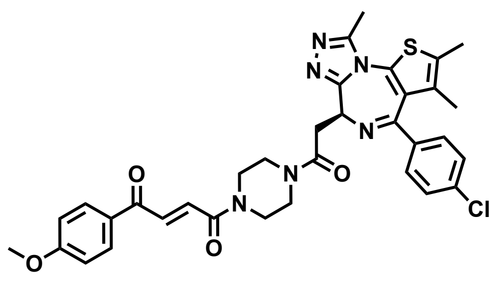

**JP-2-197**

**General Procedure A** was followed with (6S)-4-(4-chlorophenyl)-2,3,9-trimethyl-6H-thieno[3,2-f][1,2,4]triazolo[4,3-a][1,4]diazepine-6-acetic acid (JQ1-Acid) (92.5 mg, 0.23 mmol), HATU (93.0 mg, 0.24 mmol), DIPEA (0.1 mL, 0.6 mmol), and **JP-2-196** (55.0 mg, 0.2 mmol). The crude residue was purified by silica gel chromatography (0-5% MeOH in DCM) to afford 108.6 mg (83%, 2 steps) of the title compound as a yellow powder.

**$^{1}$H NMR** (700 MHz, $CDCl_3$) δ 8.08 (d, *J* = 8.6 Hz, 2H), 8.02 (dd, *J* = 14.9, 6.5 Hz, 1H), 7.51 (dd, *J* = 14.9, 6.7 Hz, 1H), 7.42 (d, *J* = 7.8 Hz, 2H), 7.39 – 7.34 (m, 2H), 7.01 (d, *J* = 8.5 Hz, 2H), 4.83 (td, *J* = 6.9, 2.9 Hz, 1H), 4.08 – 3.95 (m, 2H), 3.93 (s, 3H), 3.91 – 3.70 (m, 5H), 3.71 – 3.49 (m, 3H), 2.69 (s, 3H), 2.43 (s, 3H), 1.70 (s, 3H).

**$^{13}$C NMR** (151 MHz, $CDCl_3$) δ 187.8, 187.7, 169.3, 169.2, 164.5, 164.2, 164.2, 164.1, 155.6, 150.0, 136.7, 136.5, 134.9, 131.9, 131.3, 131.0, 131.0, 131.0, 130.9, 130.5, 129.7, 129.7, 128.6, 114.1, 114.0, 55.5, 54.2, 54.1, 45.8, 45.8, 45.6, 45.2, 42.1, 42.0, 41.8, 41.4, 35.1, 29.6, 14.2, 13.0, 11.6.

**HRMS (ESI)** *m/z* calcd for $[C_{34}H_{33}ClN_6O_4S + H]^+$ = 657.1973, found 657.2054.

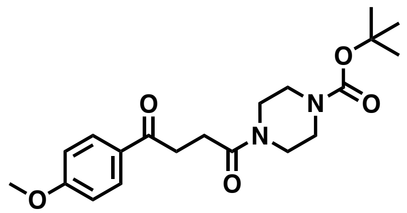

**JP-2-229**

**General Procedure A** was followed with 3-(4-methoxybenzoyl)propionic acid (100.7 mg, 0.48 mmol), HATU (206.8 mg, 0.54 mmol), DIPEA (0.25 mL, 1.45 mmol), and 1-boc-piperazine (104.7 mg, 0.56 mmol). The crude residue was purified by silica gel chromatography (0-100% EtOAc in Hexanes) to afford 132.8 mg (73%) of the title compound as a white powder.

**$^1$H NMR** (700 MHz, $CDCl_3$) δ 7.93 (d, *J* = 8.6 Hz, 2H), 6.87 (d, *J* = 8.5 Hz, 2H), 3.80 (s, 3H), 3.58 – 3.52 (m, 2H), 3.52 – 3.47 (m, 2H), 3.44 (d, *J* = 7.5 Hz, 2H), 3.35 (t, *J* = 5.5 Hz, 2H), 3.26 (t, *J* = 6.5 Hz, 2H), 2.71 (t, *J* = 6.6 Hz, 2H), 1.43 (s, 9H).

**$^{13}$C NMR** (151 MHz, $CDCl_3$) δ 197.5, 170.6, 163.5, 154.5, 130.3, 129.8, 113.7, 80.2, 55.4, 45.2, 41.6, 33.1, 29.7, 28.4, 27.1.

**LRMS (ESI)** *m/z* calcd for $[C_{20}H_{28}N_2O_5 + H]^+$ = 377.5, found 377.2.

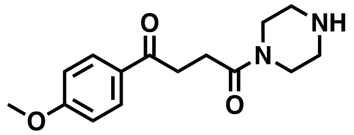


**JP-2-231**

**General Procedure E** was followed with **JP-2-229** (23.6 mg, 0.06 mmol), and TFA (0.13 mL, 1.7 mmol) for 2.5 hours. The crude material was used without further purification.

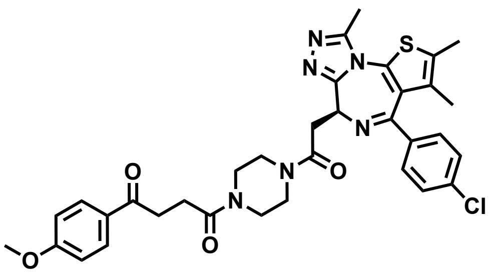


**JP-2-232**

**General Procedure A** was followed with (6S)-4-(4-chlorophenyl)-2,3,9-trimethyl-6H-thieno[3,2-f][1,2,4]triazolo[4,3-a][1,4]diazepine-6-acetic acid (JQ1-Acid) (26.1 mg, 0.07 mmol), HATU (27.3 mg, 0.07 mmol), DIPEA (0.4 mL, 0.19 mmol), and **JP-2-231** (17.3 mg, 0.06 mmol). The crude residue was purified by silica gel chromatography (0-7% MeOH in DCM) to afford 38.8 mg (94%, 2 steps) of the title compound as a yellow-white foam.

**$^1$H NMR** (700 MHz, $CDCl_3$) δ 7.99 (d, *J* = 8.8 Hz, 2H), 7.39 (d, *J* = 8.2 Hz, 2H), 7.32 (dd, *J* = 8.6, 4.2 Hz, 2H), 6.95 – 6.91 (m, 2H), 4.79 (q, *J* = 6.7 Hz, 1H), 4.01 – 3.94 (m, 1H), 3.86 (s, 3H), 3.85 – 3.80 (m, 2H), 3.79 – 3.69 (m, 4H), 3.69 – 3.63 (m, 1H), 3.60 – 3.55 (m, 1H), 3.54 – 3.48 (m, 1H), 3.42 – 3.35 (m, 1H), 3.32 – 3.26 (m, 1H), 2.84 (tt, *J* = 16.0, 6.5 Hz, 1H), 2.78 – 2.73 (m, 1H), 2.65 (d, *J* = 2.8 Hz, 3H), 2.39 (s, 3H), 1.67 (s, 3H).

**$^{13}$C NMR** (151 MHz, $CDCl_3$) δ 197.5, 170.7, 169.4, 169.2, 163.9, 163.8, 163.6, 155.8, 149.9, 149.9, 136.8, 136.7, 132.2, 130.9, 130.9, 130.7, 130.7, 130.5, 130.4, 129.9, 129.8, 128.7, 113.7, 55.5, 54.6, 54.4, 45.9, 45.6, 45.5, 45.2, 41.8, 41.7, 41.6, 35.4, 35.3, 33.2, 33.2, 31.9, 29.7, 27.2, 27.1, 22.7, 14.4, 14.1, 13.1, 11.8.

**HRMS (ESI)** *m/z* calcd for $[C_{34}H_{35}ClN_6O_4S + H]^+$ = 659.2129, found 659.2207.

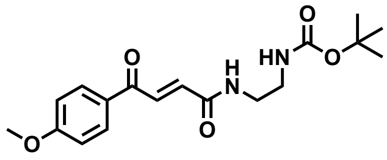


**JP-2-216**

**General Procedure A** was followed with *trans*-3-(4-methoxybenzoyl)acrylic acid (54.2 mg, 0.26 mmol), HATU (124.1 mg, 0.33 mmol), DIPEA (0.13 mL, 0.79 mmol), and *N*-boc-ethylenediamine (0.1 mL, 0.62 mmol). The crude residue was purified by silica gel chromatography (0-5% MeOH in DCM followed by 0-100% EtOAc in Hexanes) to afford 38.0 mg (42%) of the title compound as a brown residue, and 101.8 mg of a double addition byproduct. LRMS (ESI) of byproduct *m/z* calcd for $[C_{25}H_{40}N_4O_7 + H]^+$ = 509.3, found 509.3.

**$^{1}$H NMR** (700 MHz, $CDCl_3$) δ 8.03 (d, *J* = 8.4 Hz, 2H), 7.94 (d, *J* = 14.9 Hz, 1H), 6.98 – 6.93 (m, 3H), 4.99 (t, *J* = 6.2 Hz, 1H), 3.89 (s, 3H), 3.51 (q, *J* = 5.6 Hz, 2H), 3.36 (q, *J* = 5.9 Hz, 2H), 2.80 (s, 1H), 1.43 (s, 9H).

**$^{13}$C NMR** (151 MHz, $CDCl_3$) δ 187.9, 164.9, 164.2, 134.5, 133.2, 131.3, 130.0, 114.1, 80.0, 55.6, 41.5, 40.1, 38.6, 37.1, 32.8, 31.9, 30.0, 29.7, 29.7, 29.4, 28.4, 27.1, 22.7, 19.7, 14.1.

**LRMS (ESI)** *m/z* calcd for $[C_{18}H_{24}N_2O_5 + Na]^+$ = 371.2, found 371.1.

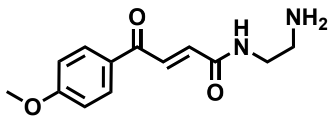


**JP-2-218**

**General Procedure E** was followed with **JP-2-216** (19.0 mg, 0.05 mmol), and TFA (0.13 mL, 1.8 mmol) for 40 minutes. The crude material was used without further purification.

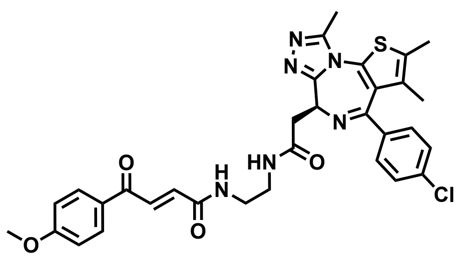

**JP-2-219**

**General Procedure A** was followed with (6S)-4-(4-chlorophenyl)-2,3,9-trimethyl-6H-thieno[3,2-f][1,2,4]triazolo[4,3-a][1,4]diazepine-6-acetic acid (JQ1-Acid) (24.6 mg, 0.06 mmol), HATU (27.0 mg, 0.07 mmol), DIPEA (0.04 mL, 0.23 mmol), and **JP-2-218** (13.5 mg, 0.05 mmol). The crude residue was purified by silica gel chromatography (0-6% MeOH in DCM) to afford 17.4 mg (51%, 2 steps) of the title compound as a yellow foam.

**$^1$H NMR** (600 MHz, $CDCl_3$) δ 8.02 – 7.98 (m, 2H), 7.95 (t, *J* = 5.6 Hz, 1H), 7.88 (s, 1H), 7.82 (q, *J* = 5.4 Hz, 1H), 7.39 – 7.36 (m, 2H), 7.29 (d, *J* = 7.4 Hz, 2H), 6.96 – 6.91 (m, 2H), 6.89 (d, *J* = 15.0 Hz, 1H), 4.71 (dd, *J* = 7.9, 6.2 Hz, 1H), 3.87 (s, 3H), 3.55 (qdt, *J* = 10.9, 6.8, 3.5 Hz, 5H), 3.46 (dd, *J* = 14.7, 6.2 Hz, 1H), 2.73 (s, 3H), 2.39 (s, 3H), 1.66 (s, 3H).

**$^{13}$C NMR** (151 MHz, $CDCl_3$) δ 188.0, 171.3, 164.8, 164.3, 164.0, 156.0, 150.4, 136.9, 136.4, 135.1, 132.6, 132.0, 131.2, 131.2, 131.0, 130.6, 130.2, 129.9, 128.7, 114.0, 55.5, 54.2, 40.3, 39.0, 38.9, 29.7, 14.4, 13.1, 11.8.

**HRMS (ESI)** *m/z* calcd for $[C_{32}H_{31}ClN_6O_4S + H]^+$ = 631.1816, found 631.1899.

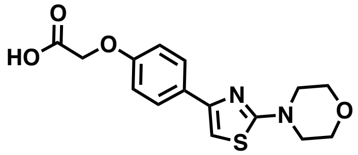

**JP-2-215**

A solution of *tert*-butyl 2-(4-(2-morpholinothiazol-4-yl)phenoxy)acetate (40.0 mg, 0.11 mmol) in 4 M hydrochloric acid in dioxane (0.7 mL) was stirred at ambient temperature overnight. The reaction mixture was diluted with toluene and the volatiles were removed *in vacuo* to yield a yellow-white solid which was used without further purification.

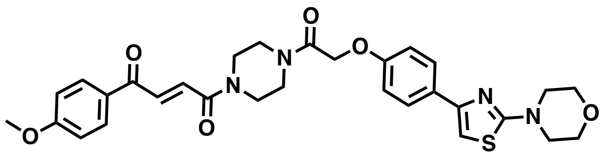

**JP-2-217**

**General Procedure A** was followed with 2-(4-(2-morpholinothiazol-4-yl)phenoxy)acetic acid (35.00 mg, 0.11 mmol), HATU (49.9 mg, 0.13 mmol), DIPEA (0.06 mL, 0.33 mmol), and **JP-2-196** (36.9 mg, 0.13 mmol). The crude residue was purified by silica gel chromatography (0-5% MeOH in DCM) to afford 34.6 mg (55%, 2 steps) of the title compound as a beige powder.

**$^1$H NMR** (700 MHz, $CDCl_3$) δ 8.03 (d, *J* = 8.6 Hz, 2H), 7.96 (dd, *J* = 14.9, 4.6 Hz, 1H), 7.79 – 7.76 (m, 2H), 7.43 (d, *J* = 14.7 Hz, 1H), 6.99 – 6.94 (m, 4H), 6.68 (s, 1H), 4.76 (d, *J* = 3.3 Hz, 2H), 3.89 (s, 3H), 3.85 – 3.82 (m, 4H), 3.73 (q, *J* = 6.3 Hz, 1H), 3.68 (p, *J* = 5.5 Hz, 5H), 3.64 – 3.59 (m, 2H), 3.54 – 3.50 (m, 4H).

**$^{13}$C NMR** (151 MHz, $CDCl_3$) δ 187.5, 171.2, 166.9, 164.4, 164.3, 157.1, 151.2, 135.1, 131.3, 130.8, 129.9, 129.3, 127.6, 114.5, 114.1, 100.5, 68.2, 68.0, 66.2, 55.6, 48.6, 46.1, 45.7, 45.2, 42.4, 42.0.

**HRMS (ESI)** *m/z* calcd for $[C_{30}H_{32}N_4O_6S + H]^+$ = 577.2043, found 577.2122.

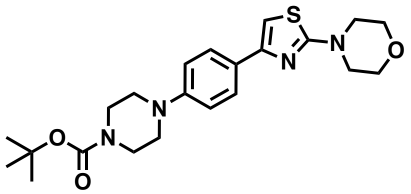

**JP-2-221**

A mixture of 4-(4-bromo-2-thiazolyl)morpholine (120.0 mg, 0.51 mmol), 4-(4-*tert*-butoxycarbonylpiperazinyl)phenylboronic acid pinacol ester (210.6, 0.54 mmol), caesium carbonate (485.9 mg, 1.5 mmol), and tetrakis(triphenylphosphine)palladium(0) (66.4 mg, 0.05 mmol) was suspended in DMF (12 mL) and stirred at 100 °C overnight. The reaction was quenched with 5% LiCl $_{(aq)}$ (60 mL) and extracted 3 times with DCM. The organic extracts were washed again with 5% LiCl $_{(aq)}$ and the organic extract was dried over $Na_2SO_4$, vacuum filtered, and the volatiles were removed *in vacuo*. The crude residue was purified by silica gel chromatography (0-50% EtOAc in Hexanes) to afford 28.8 mg (32%, BRSM) of the title compound as a white solid.

**$^1$H NMR** (700 MHz, $CDCl_3$) δ 7.75 – 7.72 (m, 2H), 6.93 – 6.90 (m, 2H), 6.65 (s, 1H), 3.85 – 3.81 (m, 4H), 3.58 (t, *J* = 5.2 Hz, 4H), 3.52 (dd, *J* = 5.9, 3.9 Hz, 4H), 3.16 (t, *J* = 5.1 Hz, 4H), 1.49 (s, 9H).

**$^{13}$C NMR** (126 MHz, $CDCl_3$) δ 171.1, 151.8, 150.7, 127.3, 127.2, 127.0, 116.8, 116.3, 99.7, 79.9, 66.3, 49.2, 48.6, 28.5.

**LRMS (ESI)** *m/z* calcd for $[C_{22}H_{30}N_4O_3S + H]^+$ = 431.6, found 431.2.

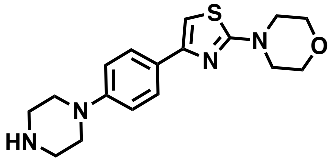

**JP-2-223**

**General Procedure E** was followed with **JP-2-221** (33.6 mg, 0.07 mmol), and TFA (0.18 mL, 2.4 mmol) for 30 minutes. The crude material was used without further purification.

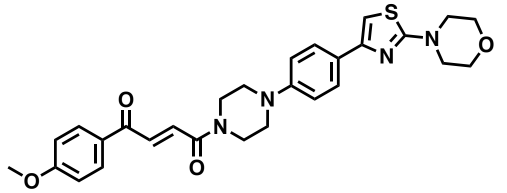

**JP-2-224**

**General Procedure A** was followed with *trans*-3-(4-methoxybenzoyl)acrylic acid (19.3 mg, 0.09 mmol), HATU (43.8 mg, 0.10 mmol), DIPEA (0.04 mL, 0.23 mmol), and **JP-2-223** (23.0 mg, 0.07 mmol). The crude residue was purified by silica gel chromatography (0-100% EtOAc in Hexanes) to afford 18.6 mg (51%, 2 steps) of the title compound as a light orange foam.

**$^{1}$H NMR** (700 MHz, $CDCl_3$) δ 8.07 – 8.03 (m, 2H), 7.99 (d, *J* = 14.8 Hz, 1H), 7.77 – 7.73 (m, 2H), 7.54 – 7.50 (m, 1H), 7.00 – 6.96 (m, 2H), 6.95 – 6.91 (m, 2H), 6.66 (s, 1H), 3.91 (t, *J* = 5.2 Hz, 2H), 3.89 (s, 3H), 3.83 (t, *J* = 4.9 Hz, 4H), 3.81 (d, *J* = 5.1 Hz, 2H), 3.52 (t, *J* = 4.9 Hz, 4H), 3.26 (q, *J* = 4.9 Hz, 4H).

**$^{13}$C NMR** (126 MHz, $CDCl_3$) δ 187.7, 171.2, 164.2, 164.1, 151.6, 150.1, 134.6, 131.3, 130.0, 127.7, 127.1, 116.4, 114.1, 99.9, 66.3, 55.6, 49.7, 49.1, 48.6, 45.9, 42.1.

**HRMS (ESI)** *m/z* calcd for $[C_{28}H_{30}N_4O_4S + H]^+$ = 519.1988, found 519.2067.

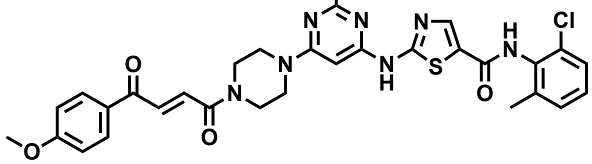

**JP-2-227**

**General Procedure A** was followed with *trans*-3-(4-methoxybenzoyl)acrylic acid (7.0 mg, 0.03 mmol), HATU (12.8 mg, 0.03 mmol), DIPEA (0.02 mL, 0.13 mmol), and *N*-deshydroxyethyl dasatinib (10.0 mg, 0.02 mmol). The crude residue was purified by silica gel chromatography (0-6% MeOH in DCM) to afford 12.8 mg (90%) of the title compound as a yellow solid.

**$^1$H NMR** (700 MHz, $CDCl_3$) δ 8.00 – 7.95 (m, 3H), 7.92 (d, *J* = 14.9 Hz, 1H), 7.42 (d, *J* = 14.9 Hz, 1H), 7.24 (d, *J* = 7.7 Hz, 1H), 7.15 – 7.07 (m, 2H), 6.95 – 6.91 (m, 2H), 5.87 (s, 1H), 3.83 (s, 3H), 3.77 (t, *J* = 5.3 Hz, 2H), 3.70 (p, *J* = 5.2 Hz, 4H), 3.62 (d, *J* = 5.5 Hz, 2H), 2.45 (s, 3H), 2.26 (s, 3H).

**$^{13}$C NMR** (126 MHz, $CDCl_3$) δ 188.0, 166.5, 164.6, 164.4, 162.8, 156.9, 140.4, 138.6, 135.0, 132.4, 131.4, 131.0, 129.7, 129.1, 128.1, 127.1, 125.6, 114.2, 83.1, 55.5, 45.5, 43.9, 43.6, 41.8, 29.6, 25.4, 18.7.

**HRMS (ESI)** *m/z* calcd for $[C_{31}H_{30}ClN_7O_4S + H]^+$ = 632.1769, found 632.1848.

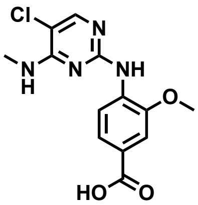

**JP-2-241**

A mixture of 2,5-dichloro-N-methyl pyrimidin-4-amine (106.2 mg, 0.60 mmol) and 4-amino-3-methoxybenzoic acid (111.8 mg, 0.67 mmol) was dissolved in a 1:1 mixture of Dioxane:$H_2O$ (4 mL). A solution of 4 M HCl in dioxane (0.15 mL) was added and the reaction mixture was stirred at 100 °C overnight. A white precipitate formed upon cooling which was filtered and washed with $H_2O$ to afford 78.7 mg (43%) of the title compound as a white powder which product was used without further purification.

**$^1$H NMR** (500 MHz, DMSO) δ 9.17 (s, 1H), 8.55 (d, *J* = 5.8 Hz, 1H), 8.29 (d, *J* = 8.5 Hz, 1H), 8.18 (s, 1H), 7.63 (d, *J* = 8.5 Hz, 1H), 7.55 (s, 1H), 3.93 (s, 3H), 2.98 (d, *J* = 4.6 Hz, 3H).

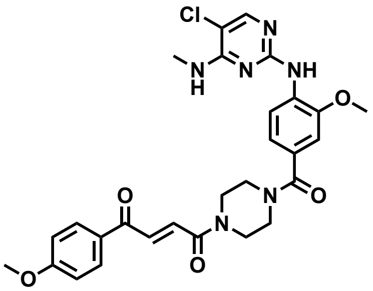

**JP-2-244**

**General Procedure A** was followed with **JP-2-241** (21.3 mg, 0.07 mmol), HATU (34.2 mg, 0.07 mmol), DIPEA (0.05 mL, 0.3 mmol), and **JP-2-196** (20.8 mg, 0.07 mmol). The crude residue was purified by silica gel chromatography (0-8% MeOH in DCM) to afford 23.3 mg (60%, 2 steps) of the title compound as a yellow film.

**$^1$H NMR** (700 MHz, $CDCl_3$) δ 8.58 (d, *J* = 8.2 Hz, 1H), 8.06 – 8.02 (m, 2H), 7.99 (d, *J* = 14.9 Hz, 1H), 7.93 (s, 1H), 7.67 (s, 1H), 7.47 (d, *J* = 14.9 Hz, 1H), 7.04 (d, *J* = 1.8 Hz, 1H), 7.02 (dd, *J* = 8.3, 1.8 Hz, 1H), 7.00 – 6.96 (m, 2H), 5.34 (q, *J* = 4.9 Hz, 1H), 3.93 (s, 3H), 3.89 (s, 3H), 3.81 – 3.59 (m, 8H), 3.11 (d, *J* = 4.9 Hz, 3H).

**$^{13}$C NMR** (151 MHz, $CDCl_3$) δ 187.5, 170.9, 164.3, 164.3, 158.6, 157.7, 152.6, 147.5, 135.0, 131.9, 131.3, 130.9, 129.9, 126.7, 120.3, 116.7, 114.2, 114.1, 109.6, 105.7, 55.9, 55.6, 46.0, 42.4, 28.1.

**HRMS (ESI)** *m/z* calcd for $[C_{28}H_{29}ClN_6O_5 + H]^+$ = 565.1888, found 565.1968.

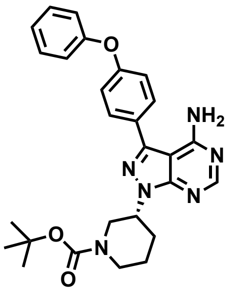

**JP-2-242**

A mixture of (4-phenoxyphenyl)boronic acid (60.6 mg, 0.28 mmol), *tert*-butyl (*R*)-3-(4-amino-3-iodo-1*H*-pyrazolo[3,4-*d*]pyrimidin-1-yl)piperidine-1-carboxylate (117.1 mg, 0.26 mmol), tetrakis(triphenylphosphine)palladium (28.4 mg, 0.03 mmol), and potassium carbonate (105.0 mg, 0.79 mmol) was suspended in a 1:1 mixture of Dioxane:$H_2O$ (4 mL) and the reaction mixture was stirred at 90 °C for 5 hours. The solvent was removed *in vacuo*, and the crude residue was purified by silica gel chromatography (0-70% EtOAc in Hexanes) to afford 127.1 mg (99%) of the title compound as a yellow foam.

**$^1$H NMR** (500 MHz, $CDCl_3$) δ 8.37 (s, 1H), 7.65 (dd, *J* = 8.1, 6.1 Hz, 2H), 7.41 – 7.36 (m, 2H), 7.15 (dd, *J* = 8.4, 6.4 Hz, 3H), 7.08 (d, *J* = 7.9 Hz, 2H), 5.64 (s, 2H), 4.84 (tt, *J* = 10.9, 4.4 Hz, 1H), 4.47 – 4.03 (m, 2H), 2.85 (td, *J* = 13.0, 3.0 Hz, 1H), 2.32 – 2.14 (m, 2H), 1.88 (d, *J* = 7.9 Hz, 1H), 1.75 – 1.63 (m, 1H), 1.44 (s, 9H).

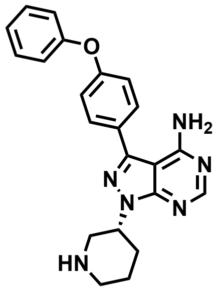

**JP-2-246**

**General Procedure E** was followed with **JP-2-242** (37.2 mg, 0.08 mmol), and TFA (0.19 mL, 2.5 mmol) for 30 minutes. The crude material was used without further purification.

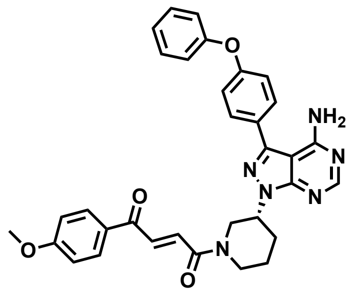

**JP-2-247**

**General Procedure A** was followed with *trans*-3-(4-methoxybenzoyl)acrylic acid (24.5 mg, 0.09 mmol), HATU (40.7 mg, 0.11 mmol), DIPEA (0.05 mL, 0.29 mmol), and **JP-2-246** (27.8 mg, 0.07 mmol). The crude residue was purified by silica gel chromatography (0-5% MeOH in DCM) and then by reverse phase silica gel chromatography (5-95% MeCN in $H_2O$) to afford 16.9 mg (41%, 2 steps) of the title compound as a white solid.
**$^1$H NMR** (700 MHz, $CDCl_3$) δ 8.38 – 8.31 (m, 1H), 8.07 – 8.03 (m, 1H), 8.00 – 7.97 (m, 1H), 7.97 – 7.85 (m, 1H), 7.67 – 7.62 (m, 2H), 7.55 – 7.41 (m, 1H), 7.40 – 7.36 (m, 2H), 7.17 (t, *J* = 7.3 Hz, 1H), 7.16 – 7.13 (m, 2H), 7.08 (d, *J* = 8.0 Hz, 2H), 7.00 – 6.96 (m, 1H), 6.96 – 6.92 (m, 1H), 5.64 (s, 2H), 4.96 – 4.87 (m, 2H), 4.55 – 4.49 (m, 1H), 4.22 (dd, *J* = 13.6, 4.1 Hz, 1H), 4.11 (d, *J* = 13.8 Hz, 1H), 3.92 (dd, *J* = 13.4, 9.9 Hz, 1H), 3.88 (d, *J* = 13.3 Hz, 3H), 3.48 (dd, *J* = 12.6, 10.6 Hz, 1H), 3.28 (td, *J* = 13.2, 2.9 Hz, 1H), 3.08 (ddd, *J* = 13.7, 11.1, 3.2 Hz, 1H), 2.45 – 2.33 (m, 1H), 2.28 (dd, *J* = 13.7, 4.1 Hz, 1H), 2.05 (ddt, *J* = 12.1, 7.5, 3.7 Hz, 1H), 1.81 – 1.71 (m, 1H).

**$^{13}$C NMR** (151 MHz, $CDCl_3$) δ 187.7, 187.6, 164.7, 164.4, 164.2, 164.1, 158.6, 158.6, 157.8, 156.4, 156.3, 155.8, 155.7, 154.3, 154.3, 144.1, 144.0, 134.4, 134.1, 132.0, 131.9, 131.3, 131.3, 130.0, 127.8, 127.6, 124.1, 119.6, 119.1, 114.1, 114.0, 98.6, 98.6, 55.6, 55.6, 53.4, 52.4, 50.1, 46.3, 46.2, 42.5, 30.3, 30.0, 25.3, 23.7.

**HRMS (ESI)** *m/z* calcd for $[C_{33}H_{30}N_6O_4 + H]^+$ = 575.2329, found 575.2406.

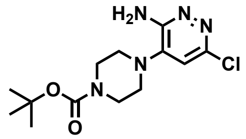


**JP-2-238**

A mixture of 3-amino-4-bromo-6-chloropyridazine (108.5 mg, 0.52 mmol) and 1-boc-piperazine (461.9 mg, 2.5 mmol) was dissolved in a THF (1 mL) and the reaction mixture was stirred at 80 °C for overnight. The solvent was removed *in vacuo*, and the crude residue was purified by silica gel chromatography (0-100% EtOAc in Hexanes) to afford 147.4 mg (90%) of the title compound as a yellow foam.

**$^{1}$H NMR** (500 MHz, $CDCl_3$) δ 6.73 (s, 1H), 5.12 (s, 2H), 3.60 – 3.55 (m, 4H), 3.00 (t, *J* = 5.0 Hz, 4H), 1.48 (s, 9H).

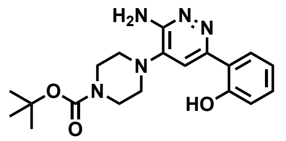


**JP-2-243**

A mixture of **JP-2-238** (122.3 mg, 0.39 mmol), 2-hydroxyphenyl boronic acid (138.4 mg, 1.00 mmol), [1,1′-Bis(diphenylphosphino)ferrocene]dichloropalladium(II) (34.5 mg, 0.05 mmol), and potassium carbonate (186.1 mg, 1.35 mmol) was suspended in a 1:1 mixture of MeCN:$H_2O$ (2.5 mL) in a sealed tube, and the reaction mixture was stirred at 120 °C for 45 minutes. The solvent was removed *in vacuo*, and the crude residue was purified by silica gel chromatography (0-100% EtOAc in Hexanes) to afford 21.2 mg (15%) of the title compound as a white-yellow powder.

**$^{1}$H NMR** (500 MHz, $CDCl_3$) δ 7.57 (dd, *J* = 8.0, 1.6 Hz, 1H), 7.29 (ddd, *J* = 8.5, 7.2, 1.6 Hz, 1H), 7.05 (dd, *J* = 8.2, 1.2 Hz, 1H), 6.91 (td, *J* = 7.6, 1.3 Hz, 1H), 4.88 (s, 2H), 3.65 (t, *J* = 5.0 Hz, 4H), 3.10 (t, *J* = 5.0 Hz, 4H), 1.50 (s, 9H).

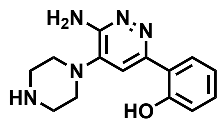

**JP-2-248**

**General Procedure E** was followed with **JP-2-243** (11.2 mg, 0.03 mmol), and TFA (0.09 mL, 0.96 mmol) for 1 hour. The solvent was removed *in vacuo*, and the crude residue was purified by silica gel chromatography (0-20% MeOH in DCM with 0.1% $Et_3N$) to afford 6.2 mg (76%) of the title compound as a yellow oil.

**$^1$H NMR** (600 MHz, DMSO) δ 7.92 (dd, *J* = 8.3, 1.6 Hz, 1H), 7.48 (s, 1H), 7.24 (ddd, *J* = 8.4, 7.3, 1.6 Hz, 1H), 6.89 (dtd, *J* = 8.2, 3.6, 1.2 Hz, 2H), 6.23 (d, *J* = 6.0 Hz, 1H), 4.01 (s, 1H), 3.04 (t, *J* = 5.0 Hz, 4H), 2.98 – 2.91 (m, 4H).

**LRMS (ESI)** *m/z* calcd for $[C_{14}H_{17}N_5O + H]^+$ = 272.3, found 272.1.

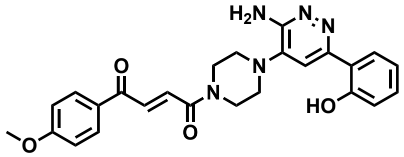

**JP-2-249**

**General Procedure A** was followed with *trans*-3-(4-methoxybenzoyl)acrylic acid (12.5 mg, 0.06 mmol), HATU (27.7 mg, 0.07 mmol), DIPEA (0.05 mL, 0.29 mmol), and **JP-2-248** (16.4 mg, 0.06 mmol). The crude residue was purified by silica gel chromatography (0-10% MeOH in DCM) to afford 25.5 mg (92%, 2 steps) of the title compound as a white-yellow powder.

**$^1$H NMR** (700 MHz, $CDCl_3$) δ 8.02 (d, *J* = 8.5 Hz, 2H), 7.97 (d, *J* = 14.9 Hz, 1H), 7.54 (d, *J* = 8.0 Hz, 1H), 7.48 (d, *J* = 14.9 Hz, 1H), 7.30 (s, 1H), 7.26 (t, *J* = 1.7 Hz, 1H), 7.00 (d, *J* = 8.1 Hz, 1H), 6.96 (d, *J* = 8.6 Hz, 2H), 6.89 (t, *J* = 7.5 Hz, 1H), 3.92 (t, *J* = 5.0 Hz, 2H), 3.87 (s, 3H), 3.86 – 3.83 (m, 2H), 3.16 (t, *J* = 5.2 Hz, 4H), 2.47 (br s, 2H).

**$^{13}$C NMR** (151 MHz, $CDCl_3$) δ 187.8, 164.6, 164.5, 158.7, 155.0, 154.1, 140.6, 135.3, 131.5, 131.1, 130.9, 129.8, 125.3, 119.1, 118.4, 117.5, 114.3, 111.6, 55.7, 49.6, 49.1, 49.1, 45.9, 42.1.

**HRMS (ESI)** *m/z* calcd for $[C_{25}H_{25}N_5O_4 + H]^+$ = 460.1979, found 460.1983.

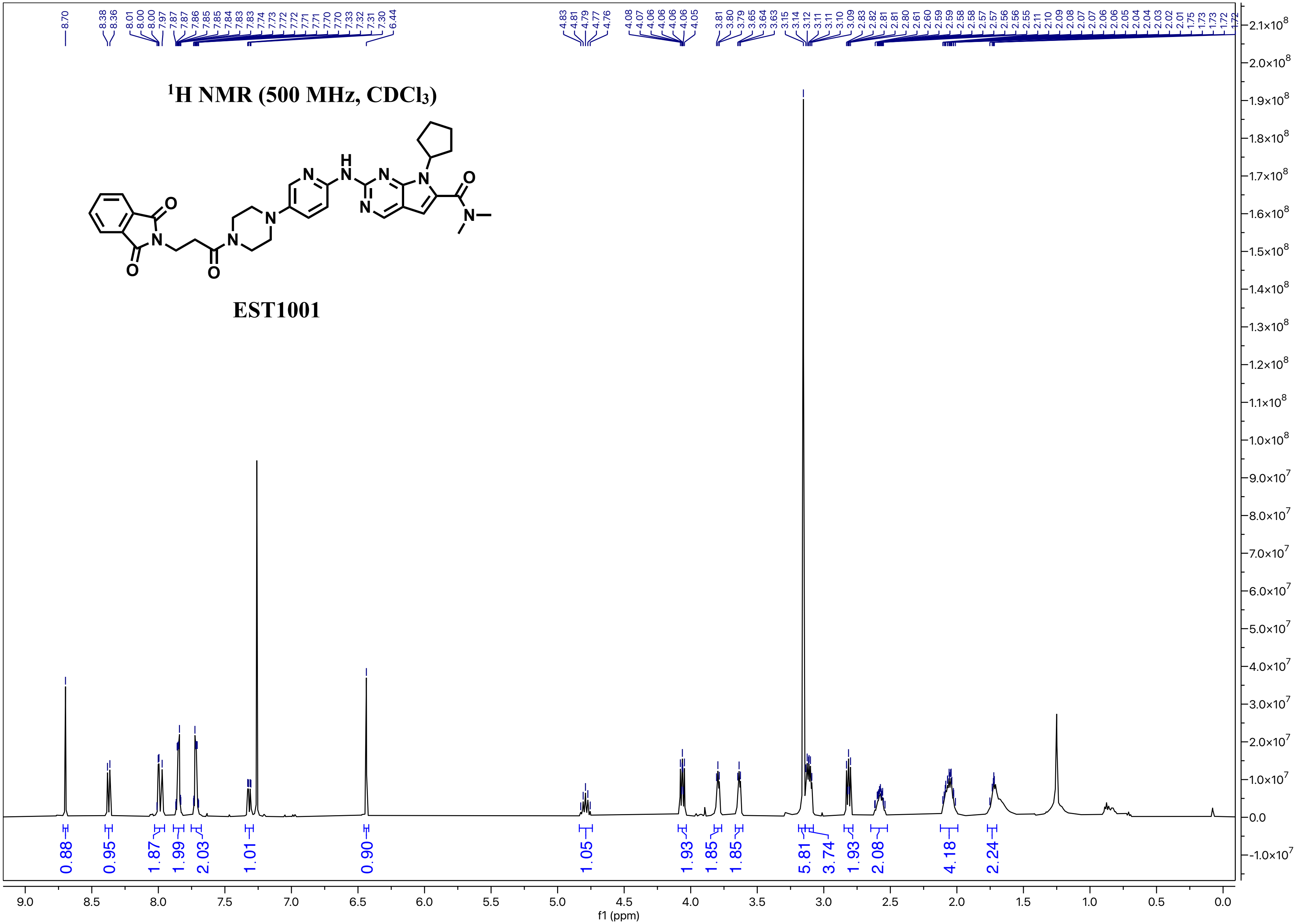

$^1$H NMR (500 MHz, CDCl$_3$)
EST1001
8.70
8.38
8.36
8.01
8.00
8.00
7.97
7.87
7.87
7.86
7.85
7.85
7.84
7.83
7.83
7.74
7.73
7.72
7.72
7.71
7.71
7.70
7.70
7.33
7.32
7.31
7.30
6.44
4.83
4.81
4.79
4.77
4.76
4.08
4.07
4.06
4.06
4.06
4.06
4.05
3.81
3.80
3.79
3.65
3.64
3.63
3.15
3.14
3.12
3.11
3.11
3.10
3.09
2.83
2.82
2.81
2.81
2.80
2.61
2.60
2.59
2.59
2.58
2.58
2.57
2.57
2.56
2.56
2.55
2.11
2.10
2.09
2.08
2.07
2.07
2.06
2.06
2.05
2.04
2.04
2.03
2.02
2.01
1.75
1.73
1.73
1.72
1.72
0.88
0.95
1.87
1.99
2.03
1.01
0.90
1.05
1.93
1.85
1.85
5.81
3.74
1.93
2.08
4.18
2.24
f1 (ppm)

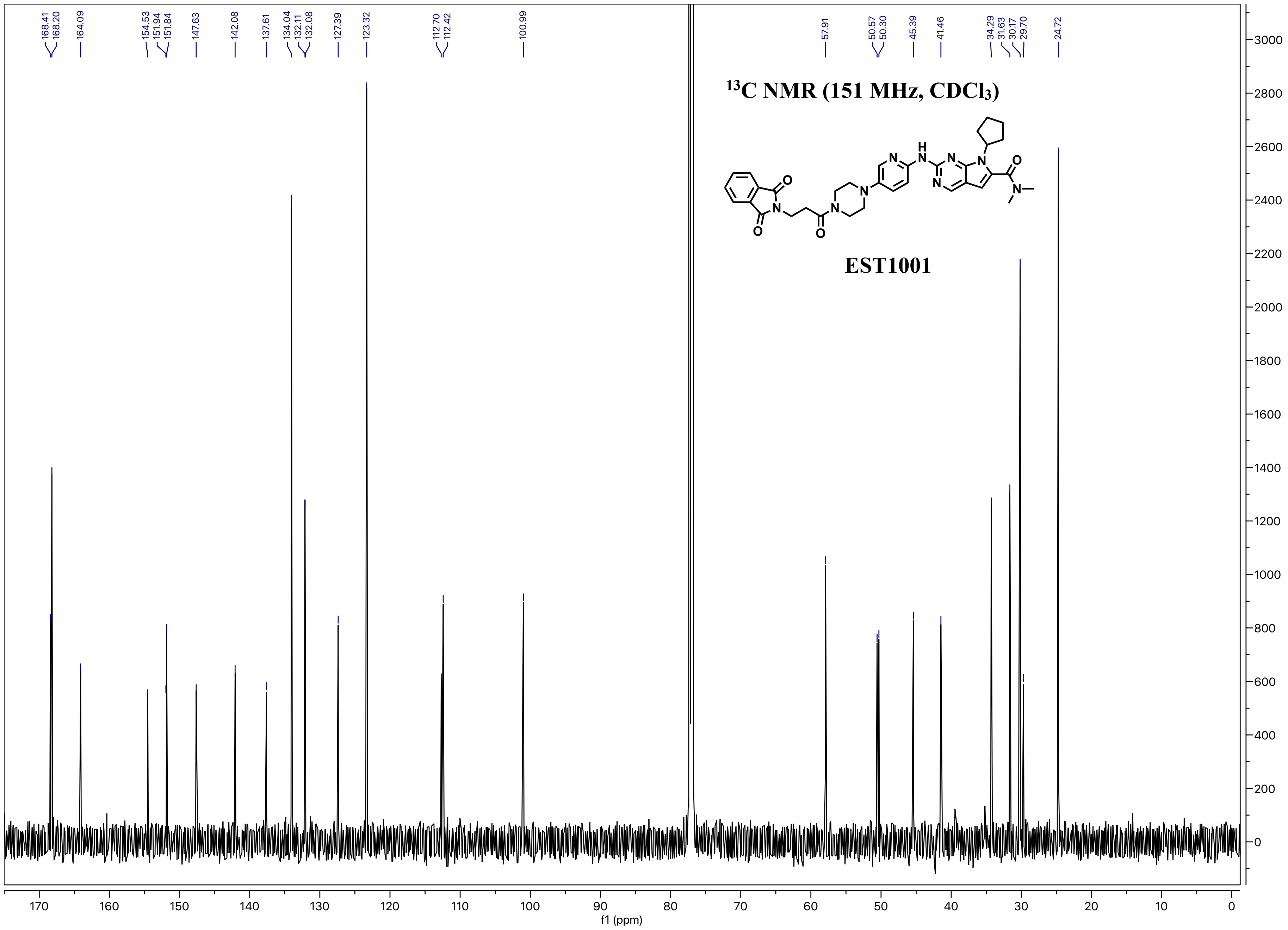

168.41
168.20
164.09
154.53
151.94
151.84
147.63
142.08
137.61
134.04
132.11
132.08
127.39
123.32
112.70
112.42
100.99
57.91
50.57
50.30
45.39
41.46
34.29
31.63
30.17
29.70
24.72
13C NMR (151 MHz, CDCl3)
EST1001
f1 (ppm)

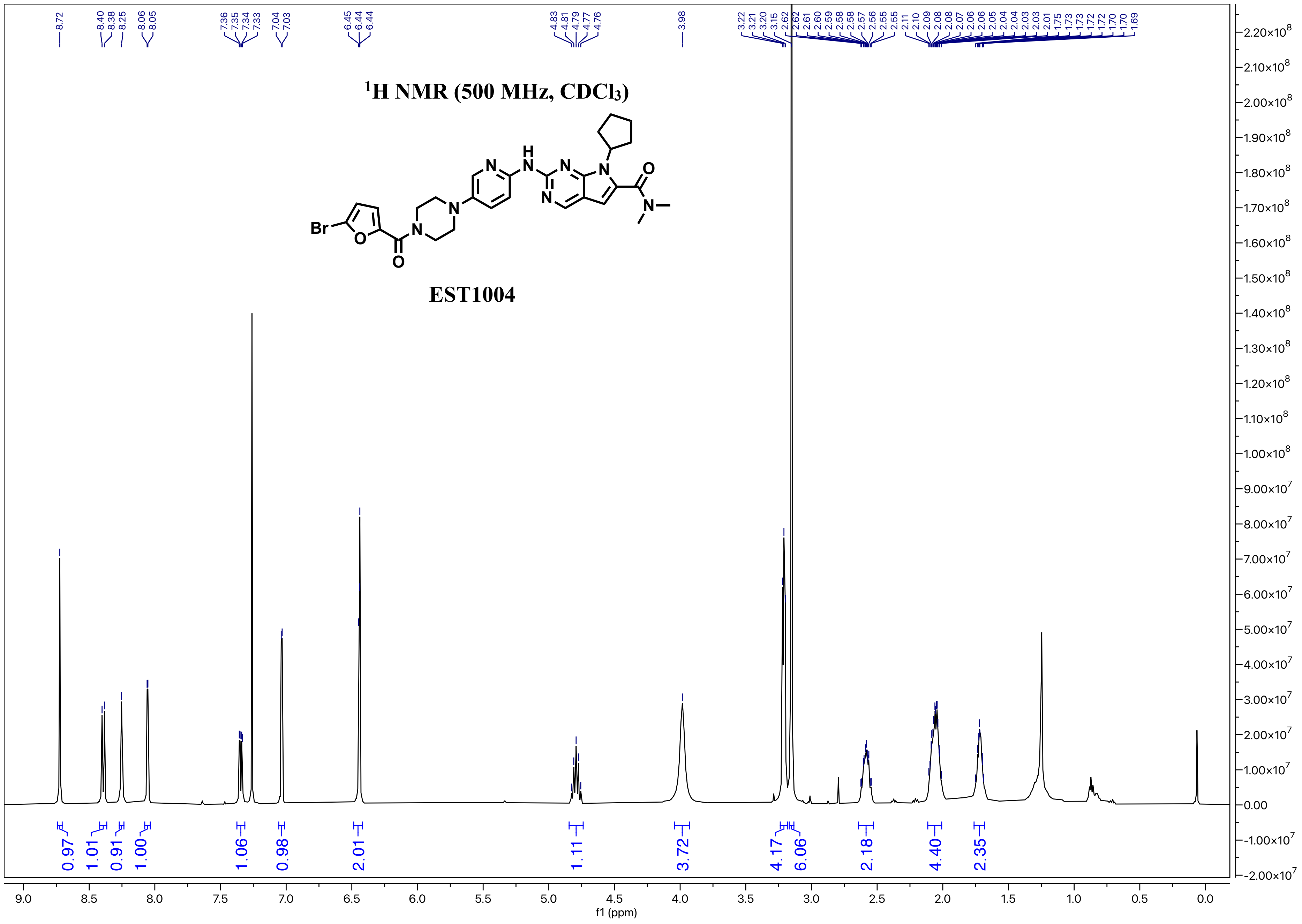
¹H NMR (500 MHz, CDCl3)
EST1004
Br
O
N
H
f1 (ppm)

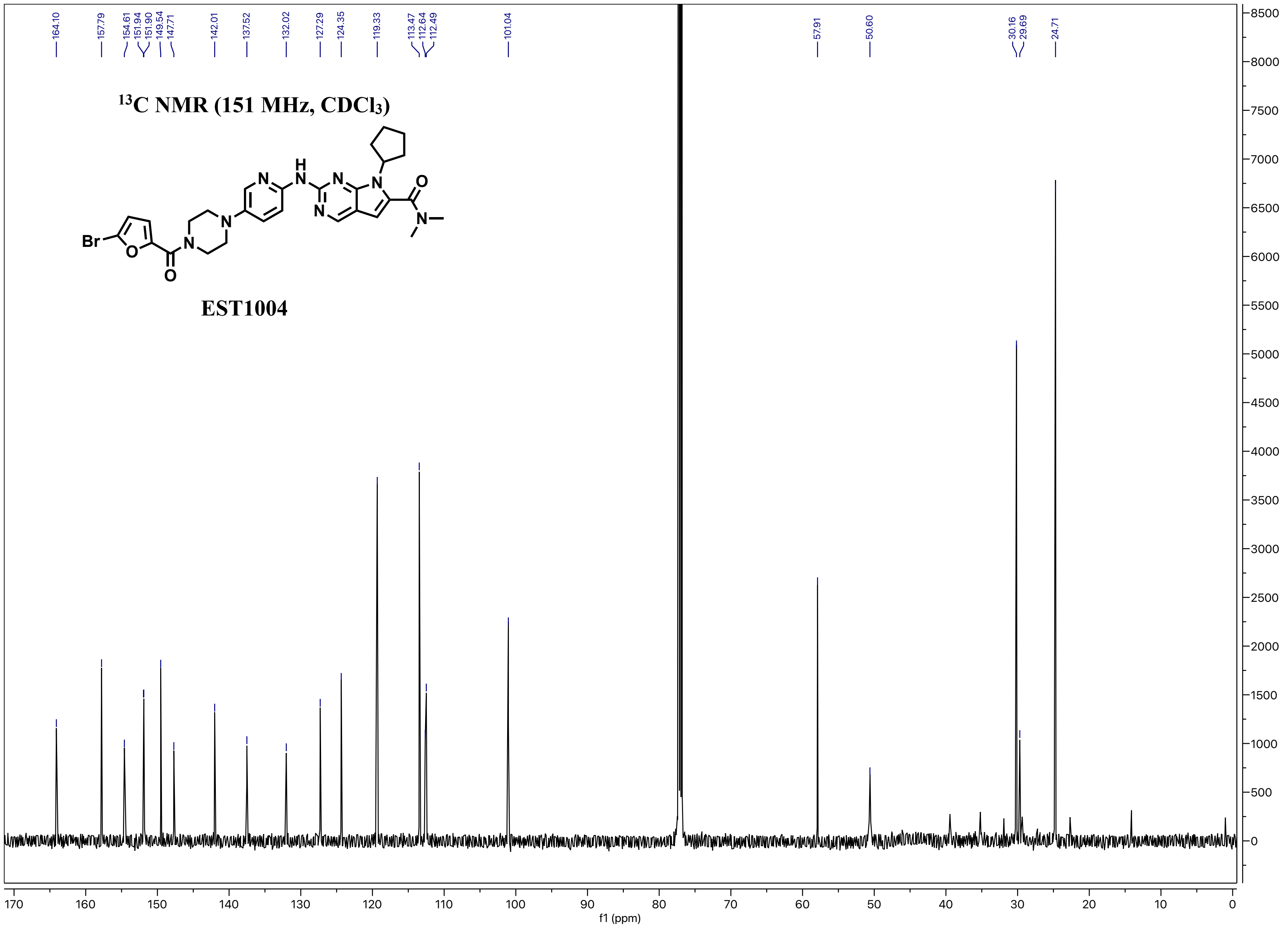

164.10
157.79
154.61
151.94
151.90
149.54
147.71
142.01
137.52
132.02
127.29
124.35
119.33
113.47
112.64
112.49
101.04
57.91
50.60
30.16
29.69
24.71
13C NMR (151 MHz, CDCl3)
EST1004
f1 (ppm)

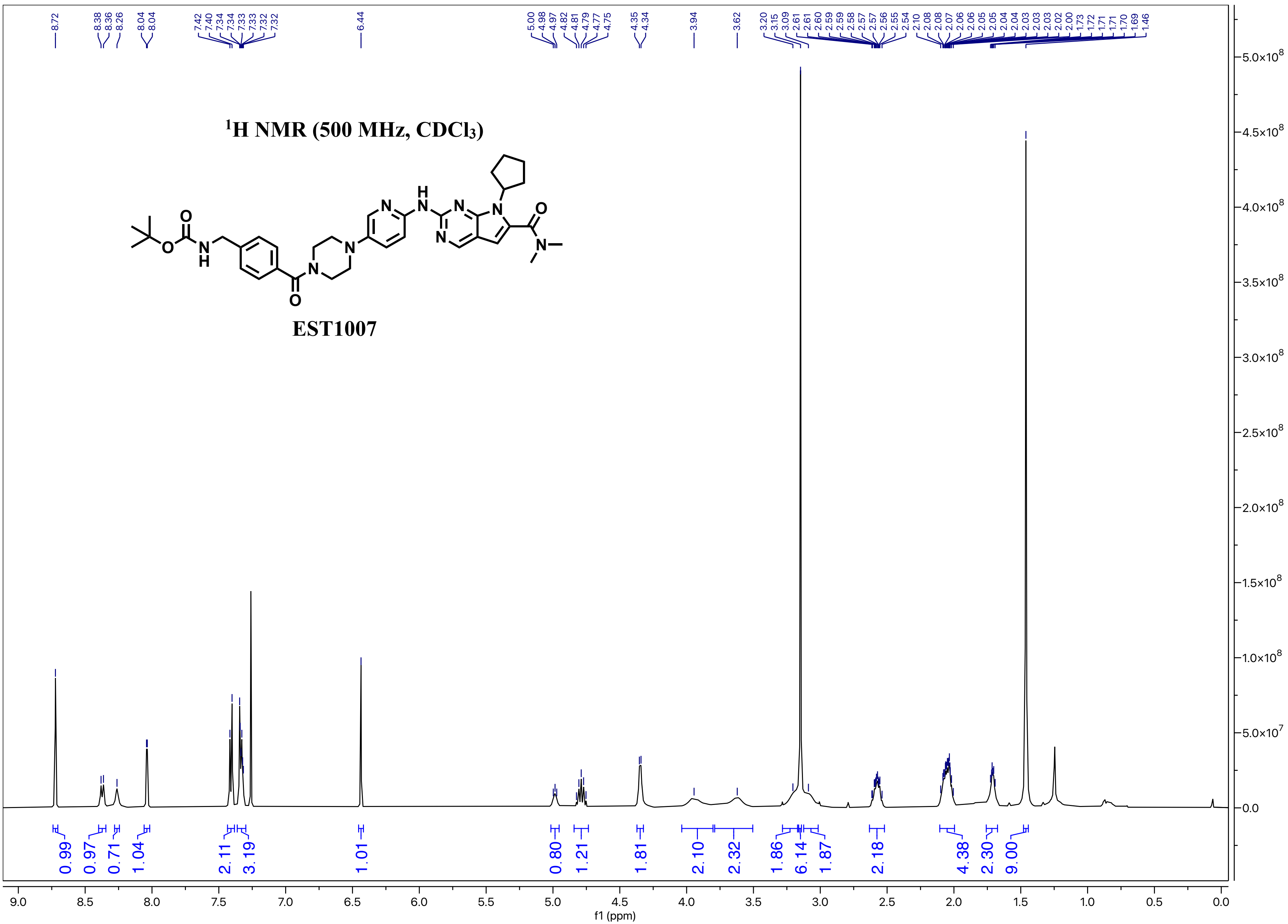

$^1$H NMR (500 MHz, CDCl$_3$)
EST1007
f1 (ppm)

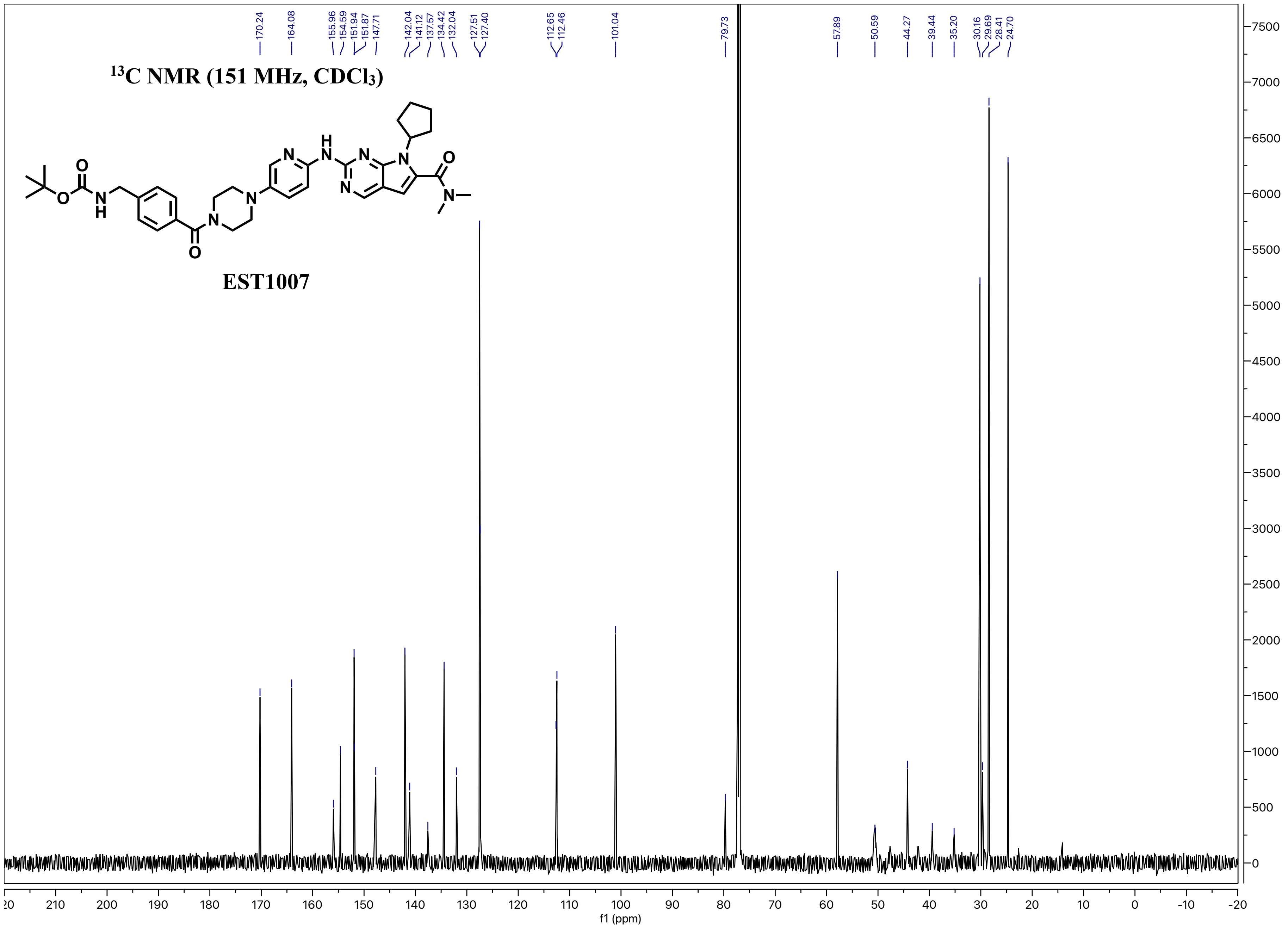

$^{13}C$ NMR (151 MHz, $CDCl_3$)
EST1007
170.24
164.08
155.96
154.59
151.94
151.87
147.71
142.04
141.12
137.57
134.42
132.04
127.51
127.40
112.65
112.46
101.04
79.73
57.89
50.59
44.27
39.44
35.20
30.16
29.69
28.41
24.70
f1 (ppm)

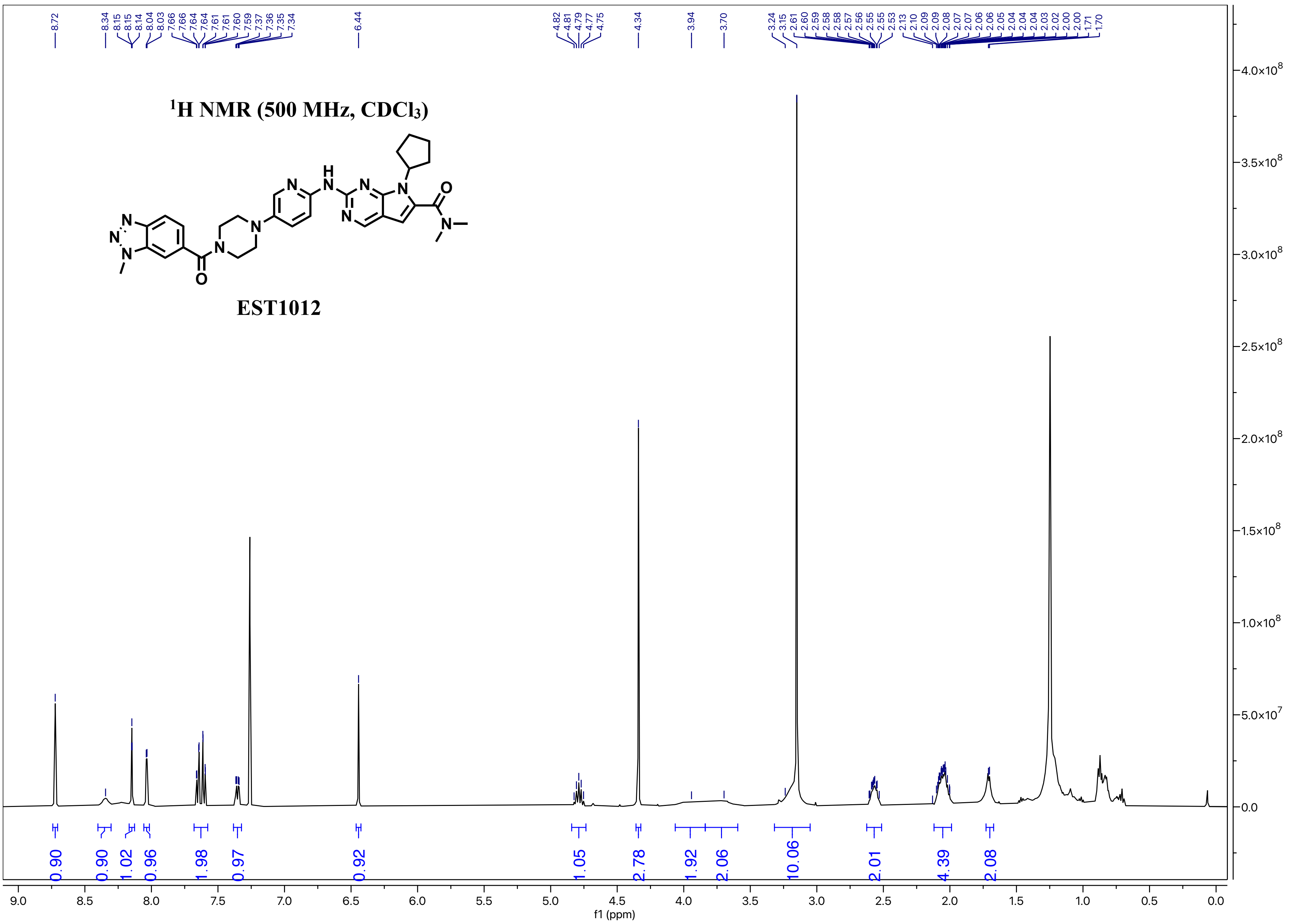

$^1$H NMR (500 MHz, CDCl$_3$)
EST1012
8.72
8.34
8.15
8.15
8.14
8.04
8.03
7.66
7.66
7.64
7.64
7.61
7.61
7.60
7.59
7.37
7.36
7.35
7.34
6.44
4.82
4.81
4.79
4.77
4.75
4.34
3.94
3.70
3.24
3.15
2.61
2.60
2.59
2.58
2.58
2.57
2.56
2.55
2.55
2.53
2.13
2.10
2.09
2.09
2.08
2.07
2.07
2.06
2.06
2.05
2.04
2.04
2.04
2.03
2.02
2.00
2.00
1.71
1.70
0.90
0.90
1.02
0.96
1.98
0.97
0.92
1.05
2.78
1.92
2.06
10.06
2.01
4.39
2.08
f1 (ppm)

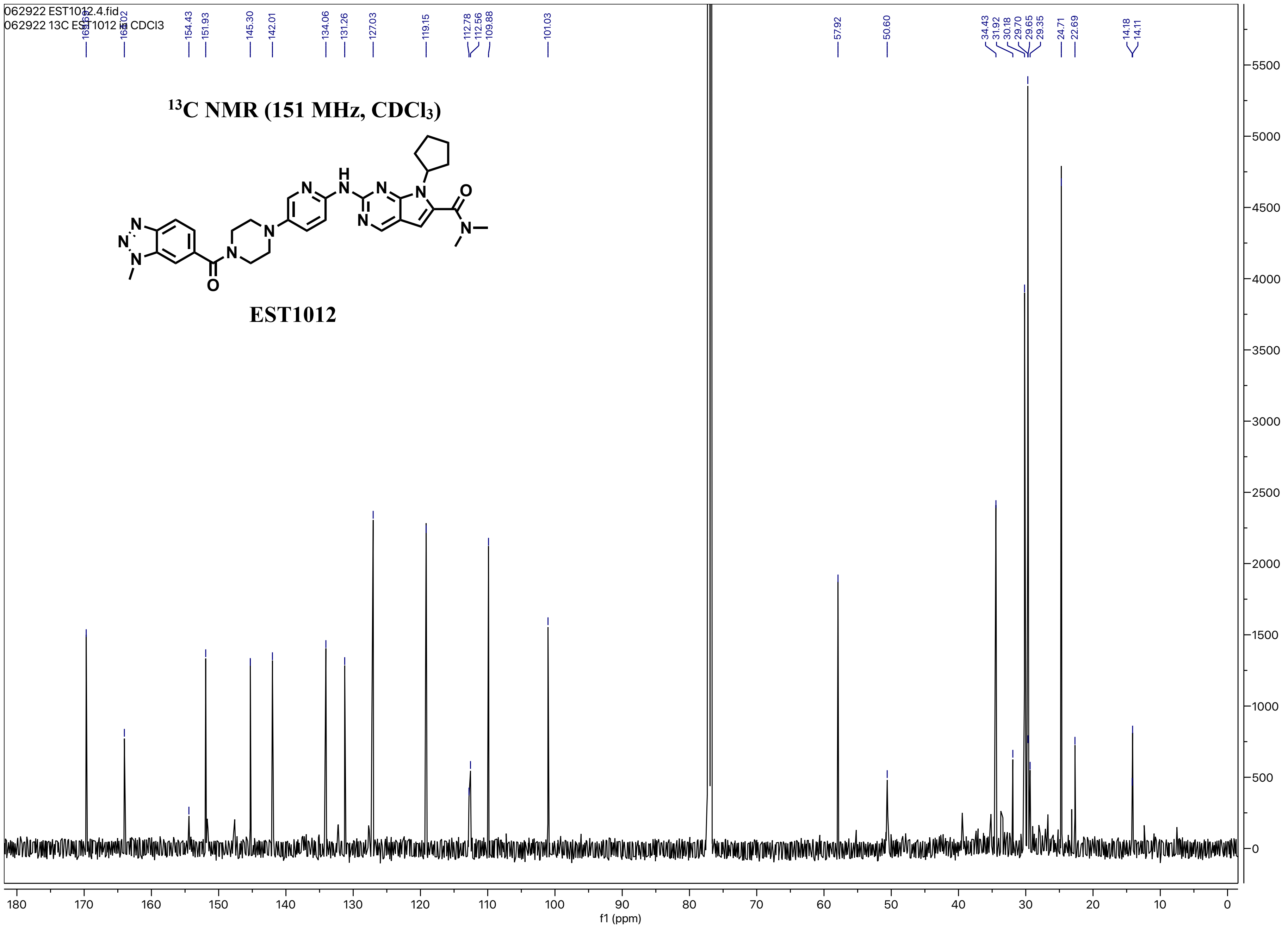
062922 EST1012.4.fid
062922 13C EST1012 in CDCl3
169.69
164.02
154.43
151.93
145.30
142.01
134.06
131.26
127.03
119.15
112.78
112.56
109.88
101.03
57.92
50.60
34.43
31.92
30.18
29.70
29.65
29.35
24.71
22.69
14.18
14.11
13C NMR (151 MHz, CDCl3)
EST1012
f1 (ppm)

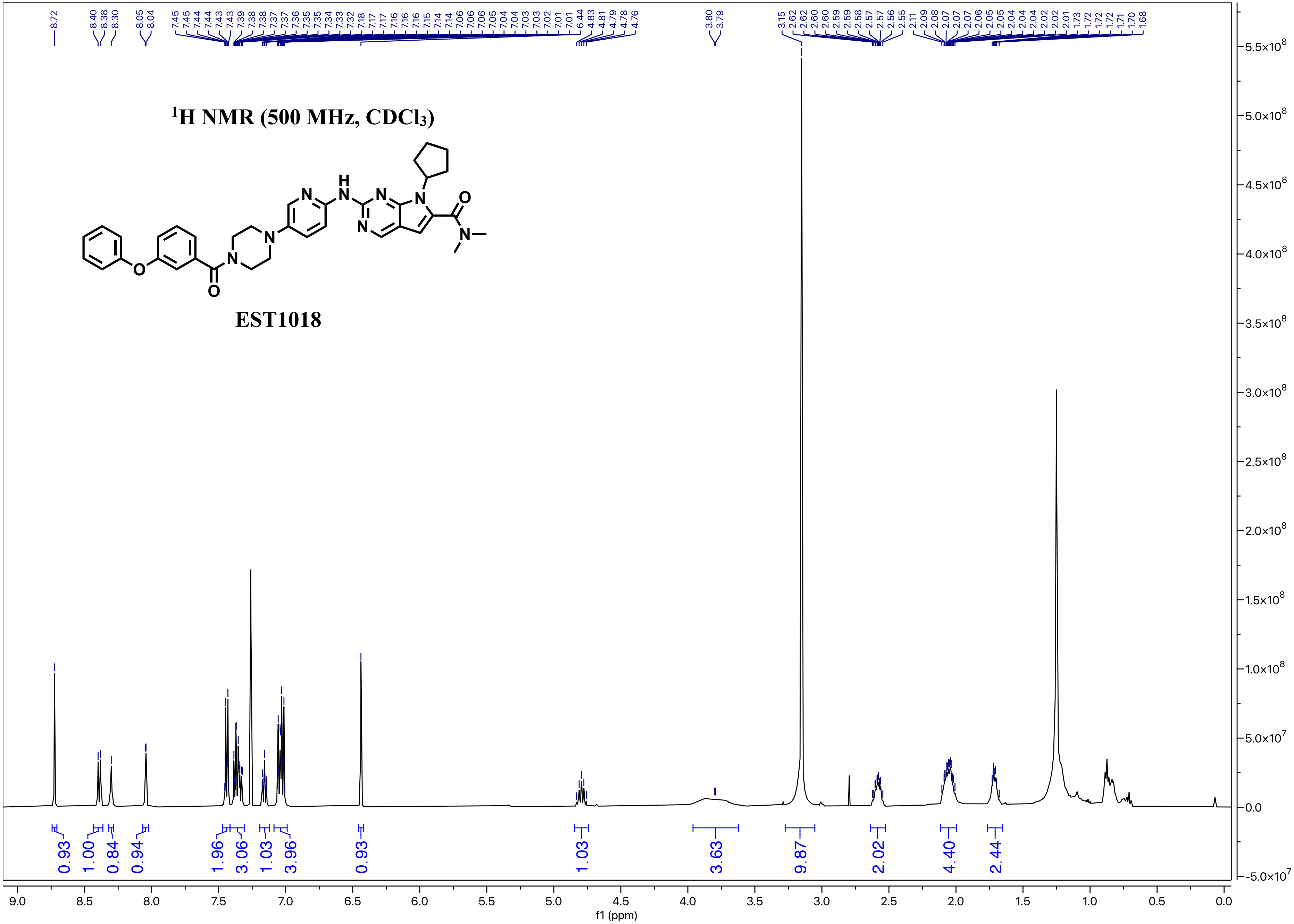

1H NMR (500 MHz, CDCl3)
EST1018
f1 (ppm)

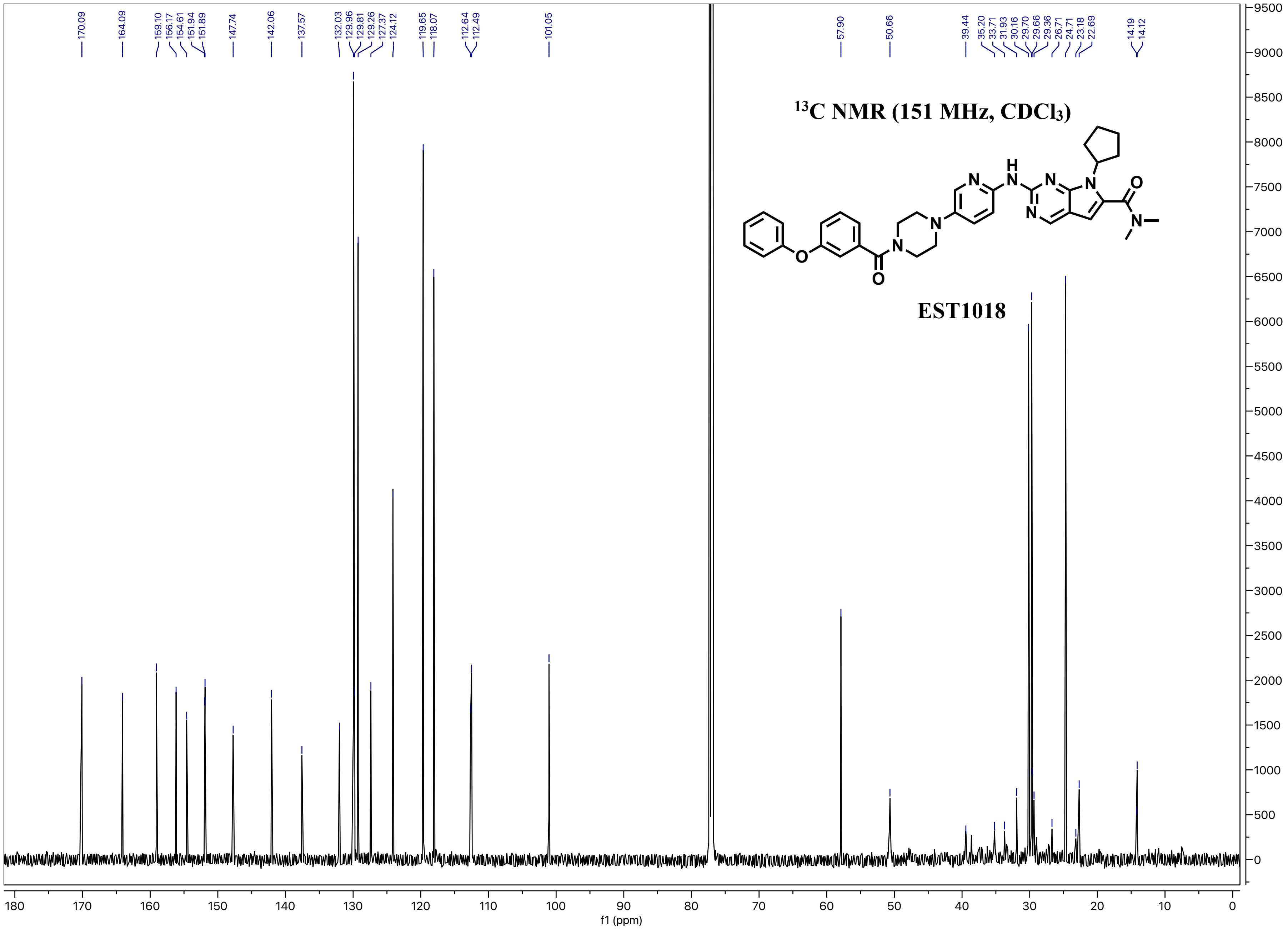

13C NMR (151 MHz, CDCl3)
EST1018
170.09
164.09
159.10
156.17
154.61
151.94
151.89
147.74
142.06
137.57
132.03
129.96
129.81
129.26
127.37
124.12
119.65
118.07
112.64
112.49
101.05
57.90
50.66
39.44
35.20
33.71
31.93
30.16
29.70
29.66
29.36
26.71
24.71
23.18
22.69
14.19
14.12
f1 (ppm)

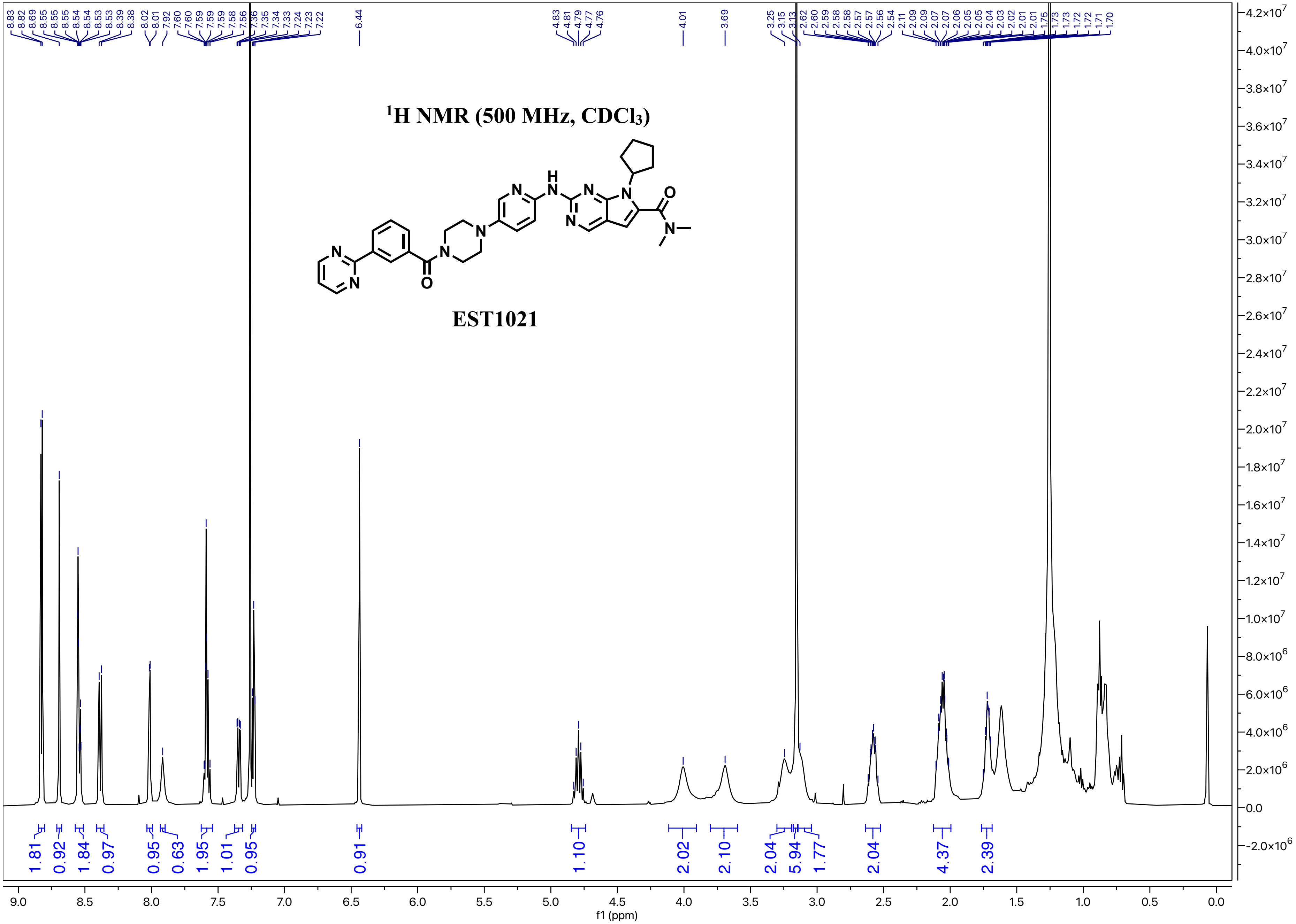

1H NMR (500 MHz, CDCl3)
EST1021
f1 (ppm)

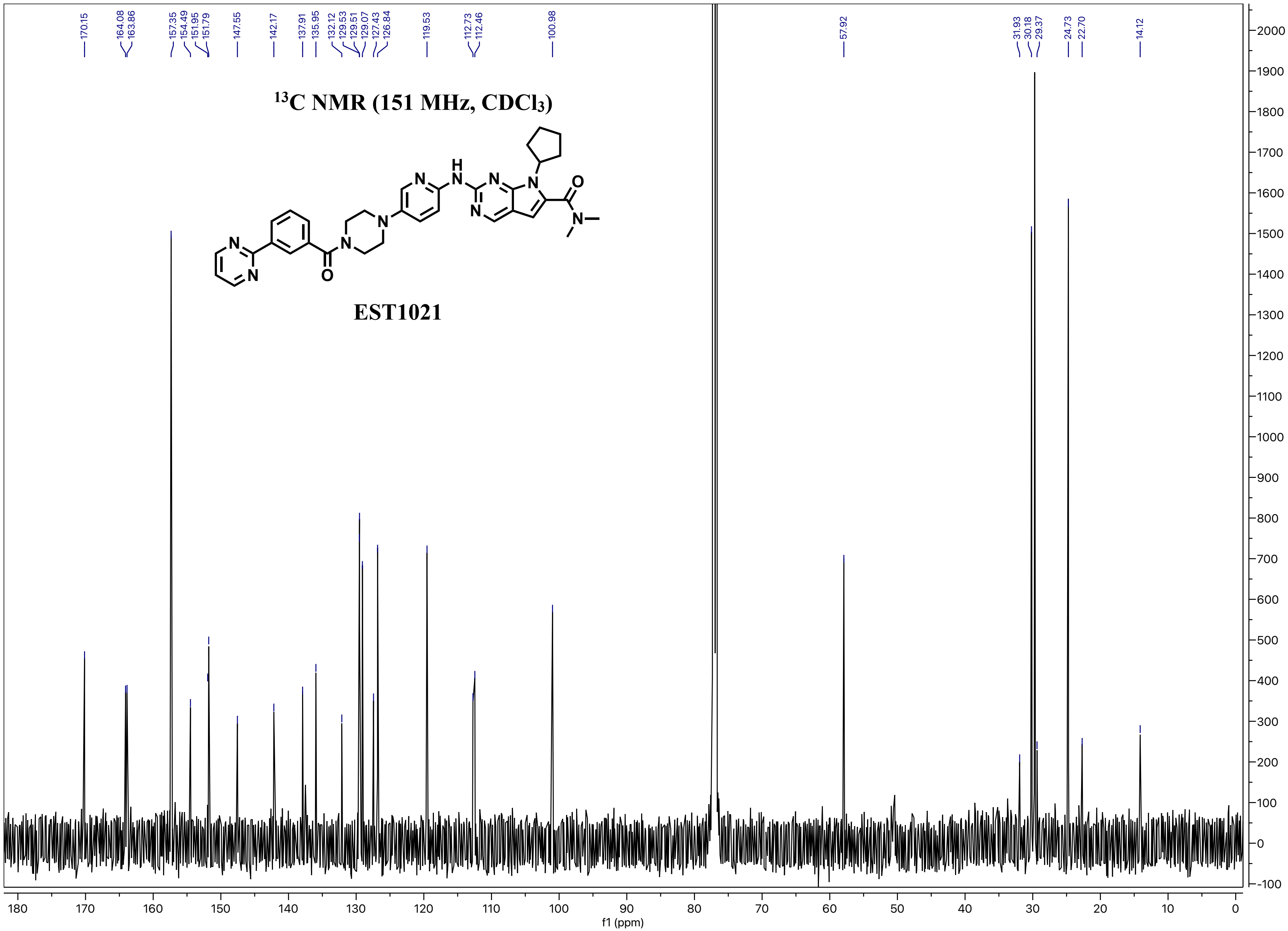

13C NMR (151 MHz, CDCl3)
EST1021
170.15
164.08
163.86
157.35
154.49
151.95
151.79
147.55
142.17
137.91
135.95
132.12
129.53
129.51
129.07
127.43
126.84
119.53
112.73
112.46
100.98
57.92
31.93
30.18
29.37
24.73
22.70
14.12
f1 (ppm)

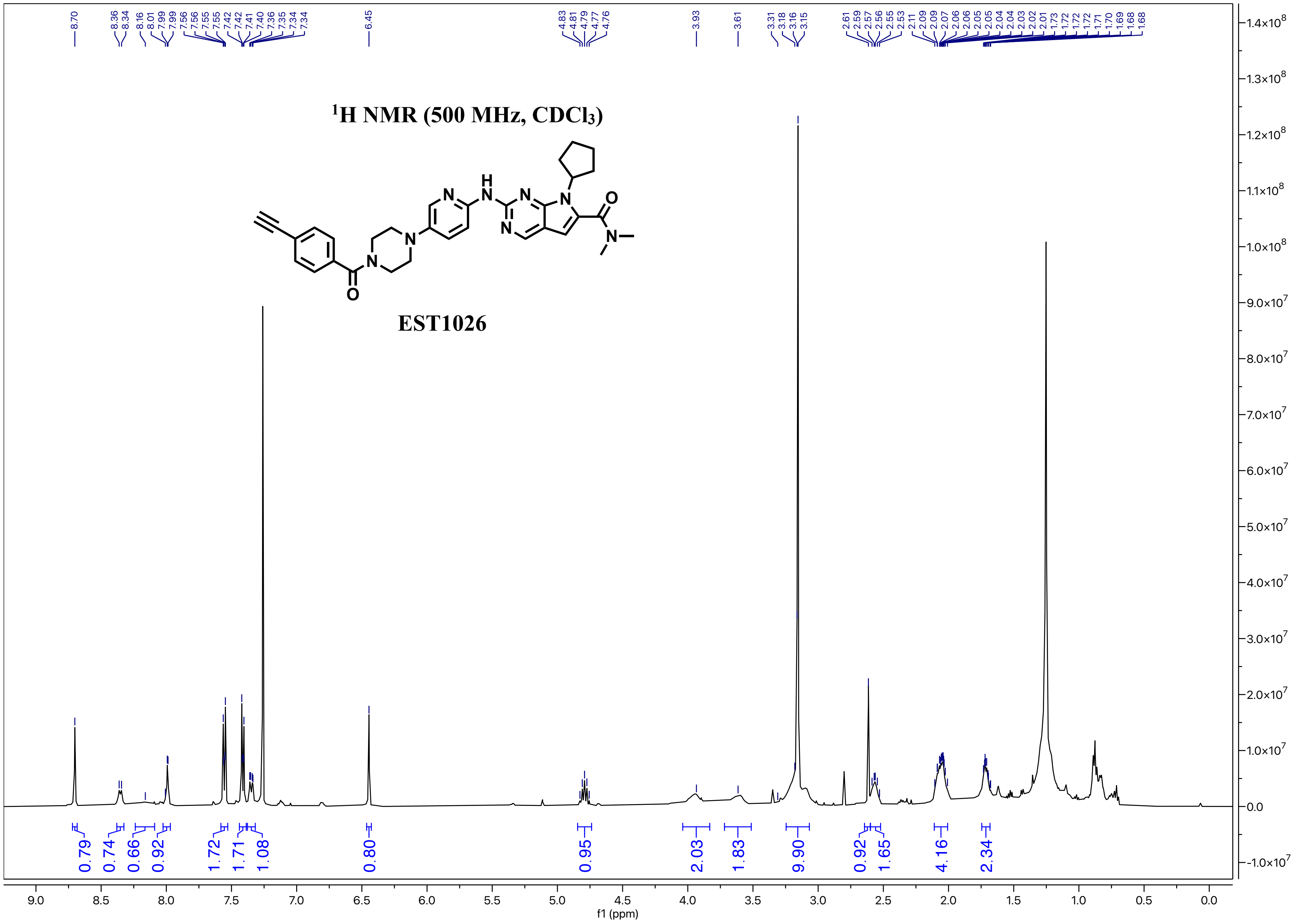

1H NMR (500 MHz, CDCl3)
EST1026
8.70
8.36
8.34
8.16
8.01
7.99
7.99
7.56
7.56
7.55
7.55
7.42
7.42
7.41
7.40
7.36
7.35
7.34
7.34
6.45
4.83
4.81
4.79
4.77
4.76
3.93
3.61
3.31
3.18
3.16
3.15
2.61
2.59
2.57
2.56
2.55
2.53
2.11
2.09
2.07
2.06
2.06
2.05
2.05
2.04
2.04
2.03
2.02
2.01
1.73
1.72
1.72
1.72
1.71
1.70
1.69
1.68
1.68
0.79
0.74
0.66
0.92
1.72
1.71
1.08
0.80
0.95
2.03
1.83
9.90
0.92
1.65
4.16
2.34
f1 (ppm)

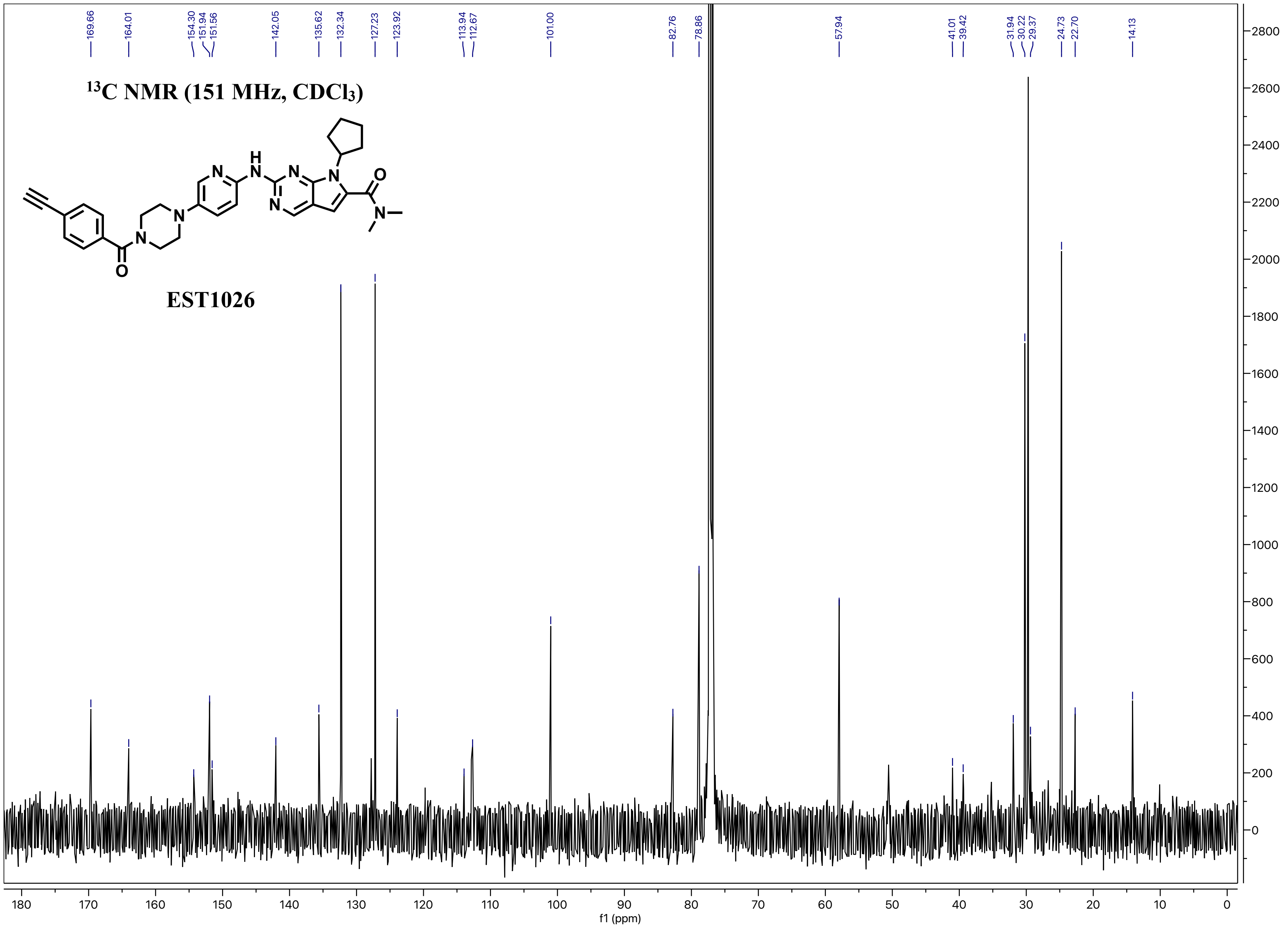

$^{13}$C NMR (151 MHz, CDCl$_3$)
EST1026
169.66
164.01
154.30
151.94
151.56
142.05
135.62
132.34
127.23
123.92
113.94
112.67
101.00
82.76
78.86
57.94
41.01
39.42
31.94
30.22
29.37
24.73
22.70
14.13
f1 (ppm)

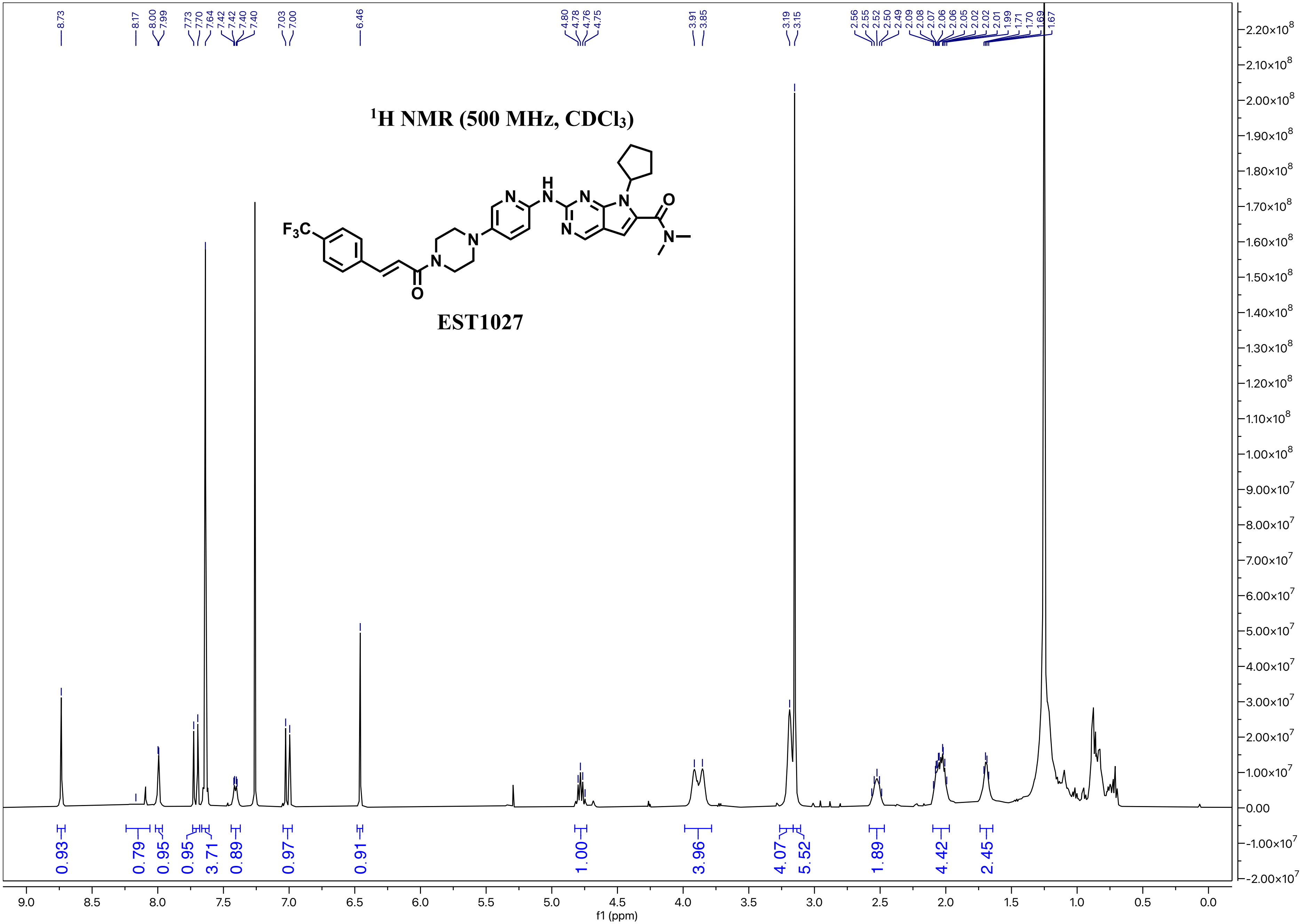

$^1$H NMR (500 MHz, $CDCl_3$)
EST1027
8.73
8.17
8.00
7.99
7.73
7.70
7.64
7.42
7.42
7.40
7.40
7.03
7.00
6.46
4.80
4.78
4.76
4.75
3.91
3.85
3.19
3.15
2.56
2.55
2.52
2.50
2.49
2.09
2.08
2.07
2.06
2.06
2.05
2.02
2.02
2.01
1.99
1.71
1.70
1.69
1.67
0.93
0.79
0.95
0.95
3.71
0.89
0.97
0.91
1.00
3.96
4.07
5.52
1.89
4.42
2.45
f1 (ppm)

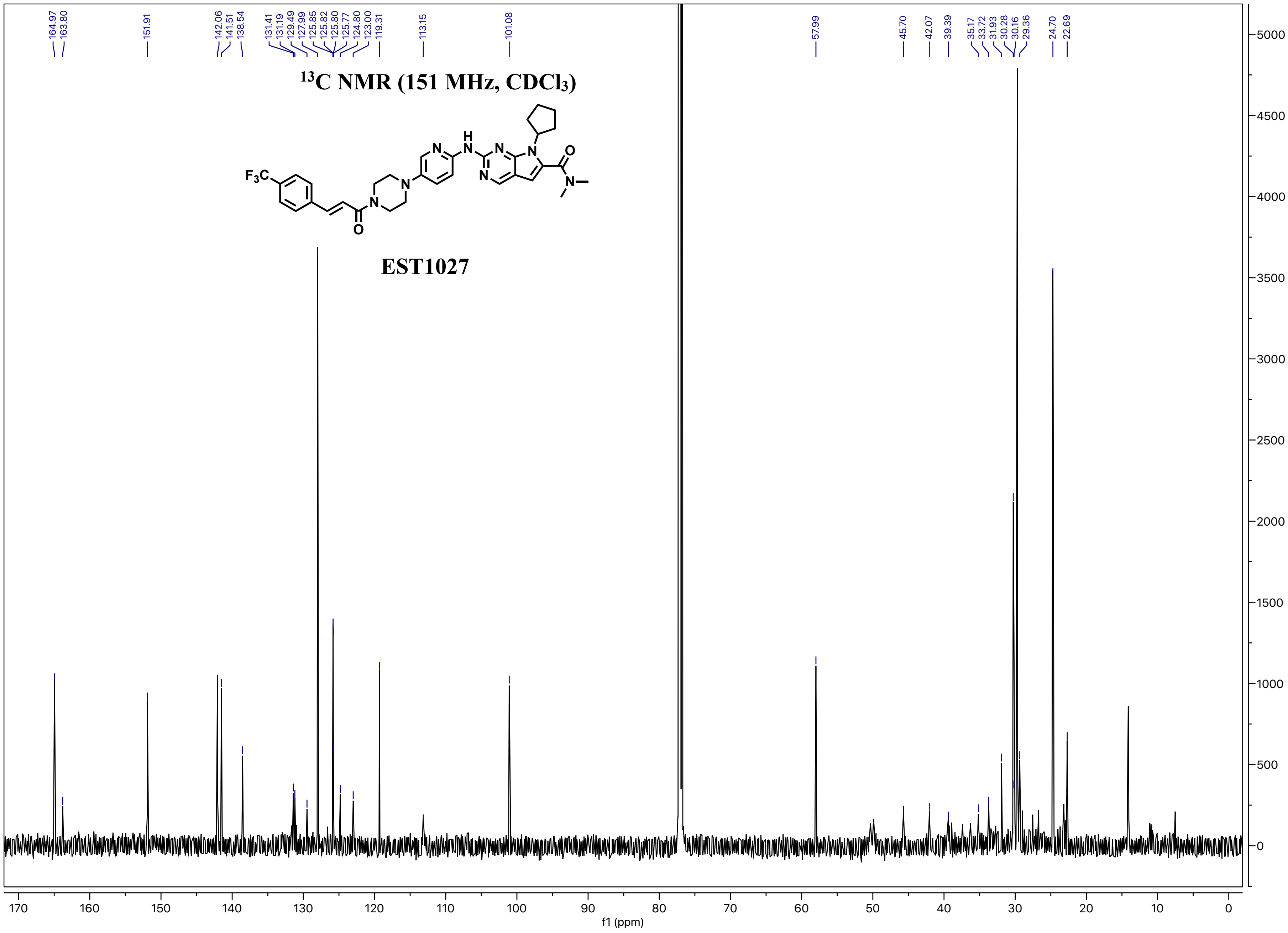
13C NMR (151 MHz, CDCl3)
EST1027
F3C
164.97
163.80
151.91
142.06
141.51
138.54
131.41
131.19
129.49
127.99
125.85
125.82
125.80
125.77
124.80
123.00
119.31
113.15
101.08
57.99
45.70
42.07
39.39
35.17
33.72
31.93
30.28
30.16
29.36
24.70
22.69
f1 (ppm)

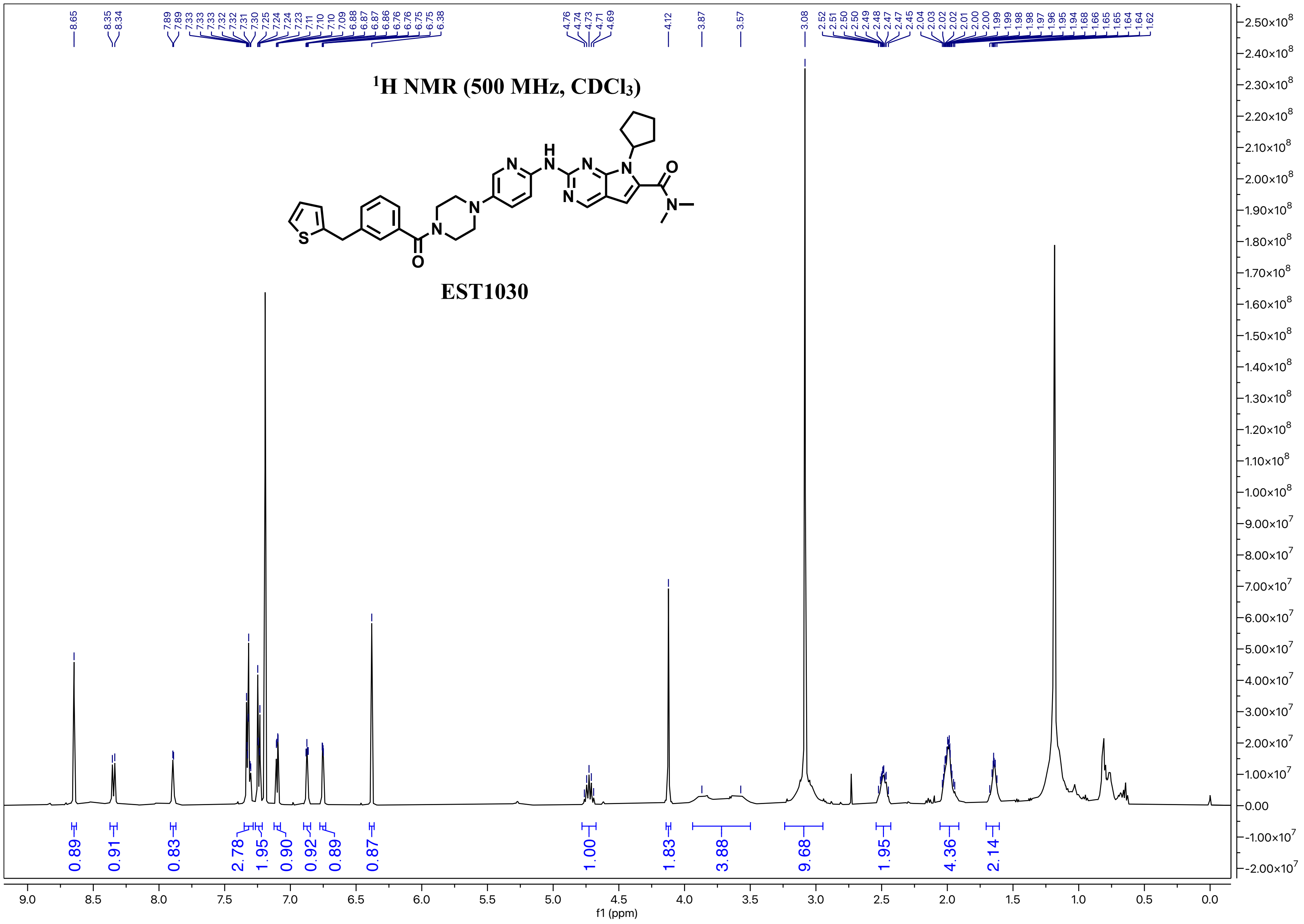

$^1$H NMR (500 MHz, CDCl$_3$)
EST1030
f1 (ppm)

#### $^{13}C$ NMR (151 MHz, $CDCl_3$)

**EST1030**

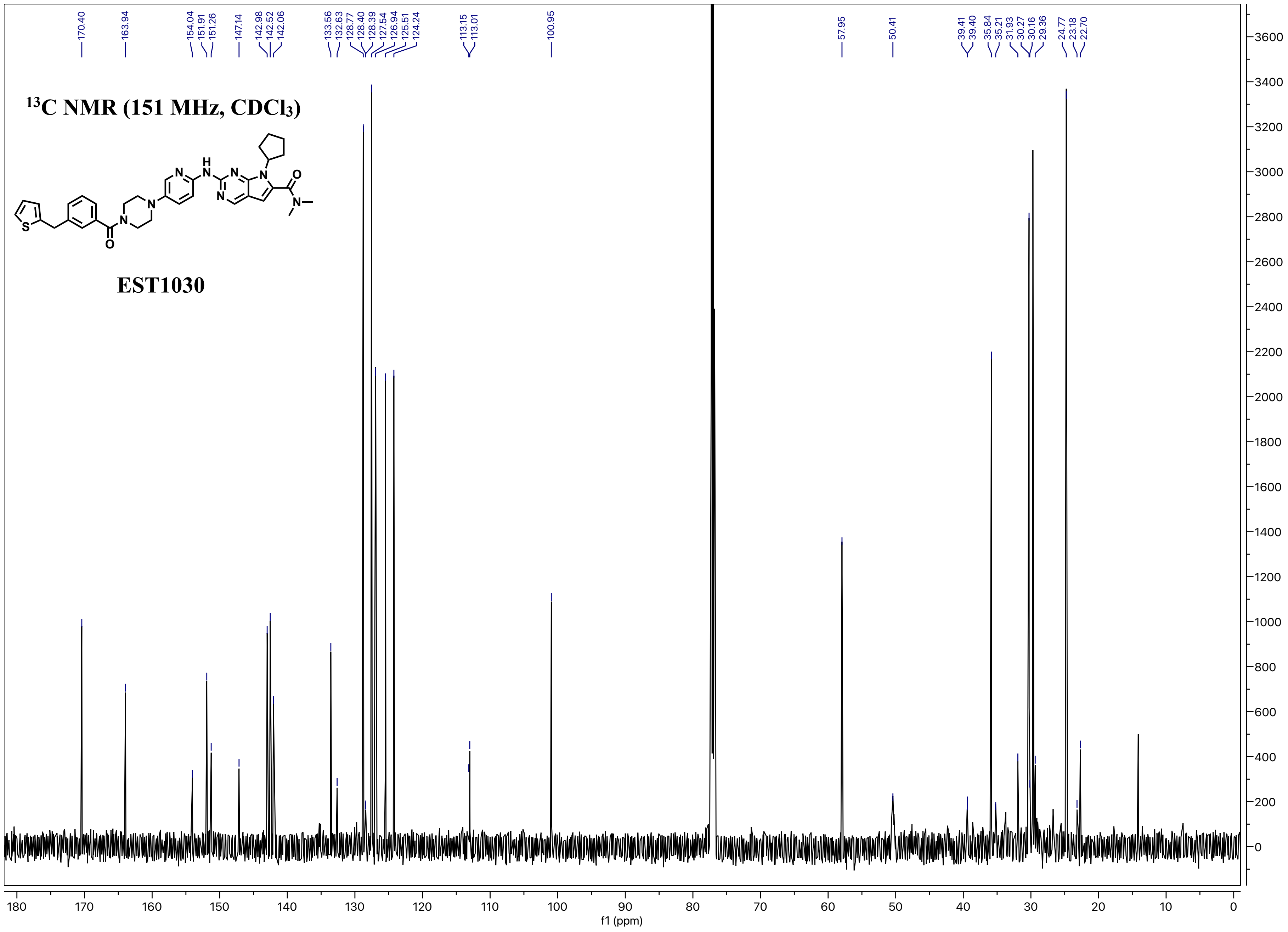

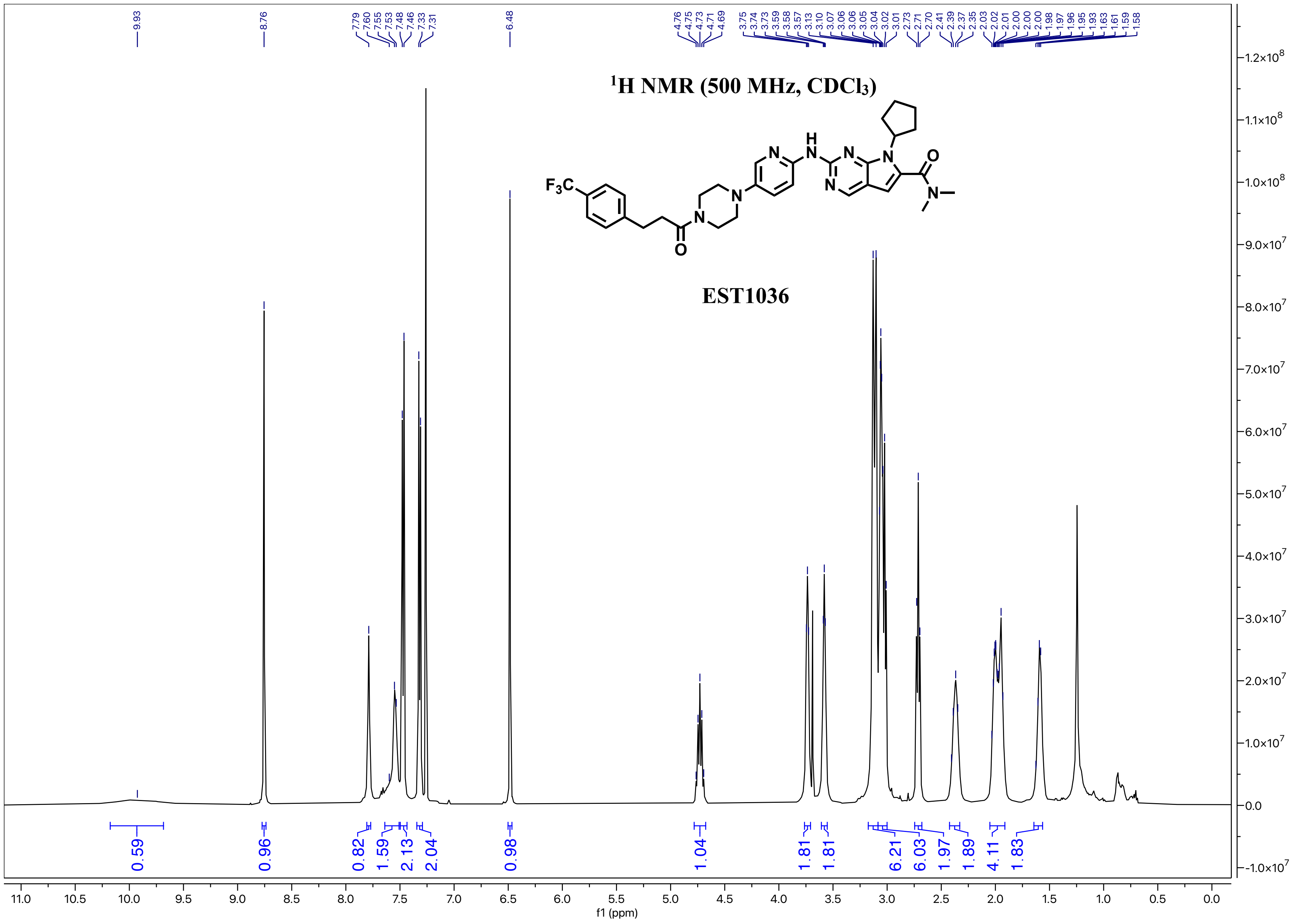

$^{1}$H NMR (500 MHz, CDCl$_3$)
EST1036
f1 (ppm)

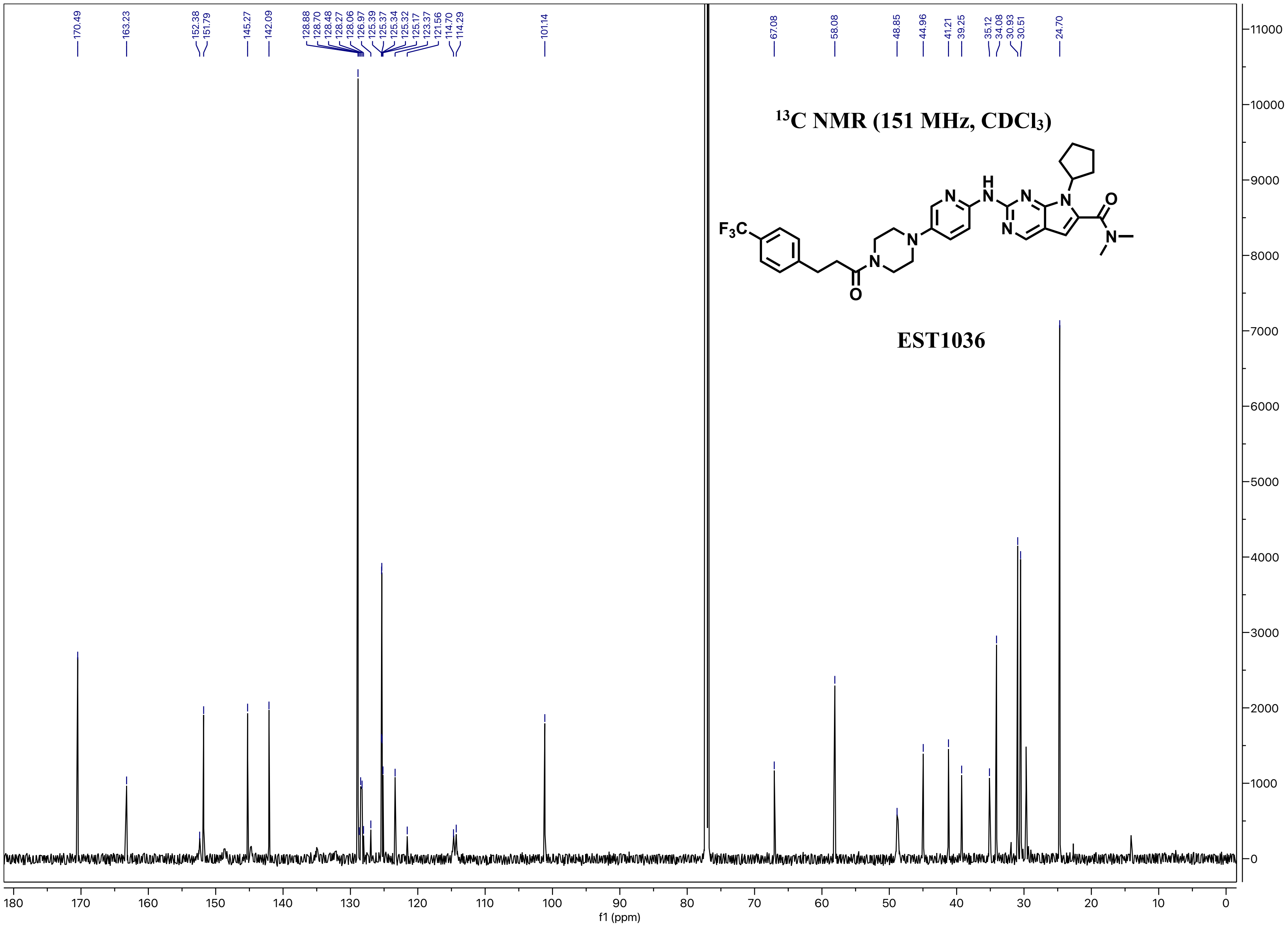

170.49
163.23
152.38
151.79
145.27
142.09
128.88
128.70
128.48
128.27
128.06
126.97
125.39
125.37
125.34
125.32
125.17
123.37
121.56
114.70
114.29
101.14
67.08
58.08
48.85
44.96
41.21
39.25
35.12
34.08
30.93
30.51
24.70
13C NMR (151 MHz, CDCl3)
F3C
EST1036
f1 (ppm)

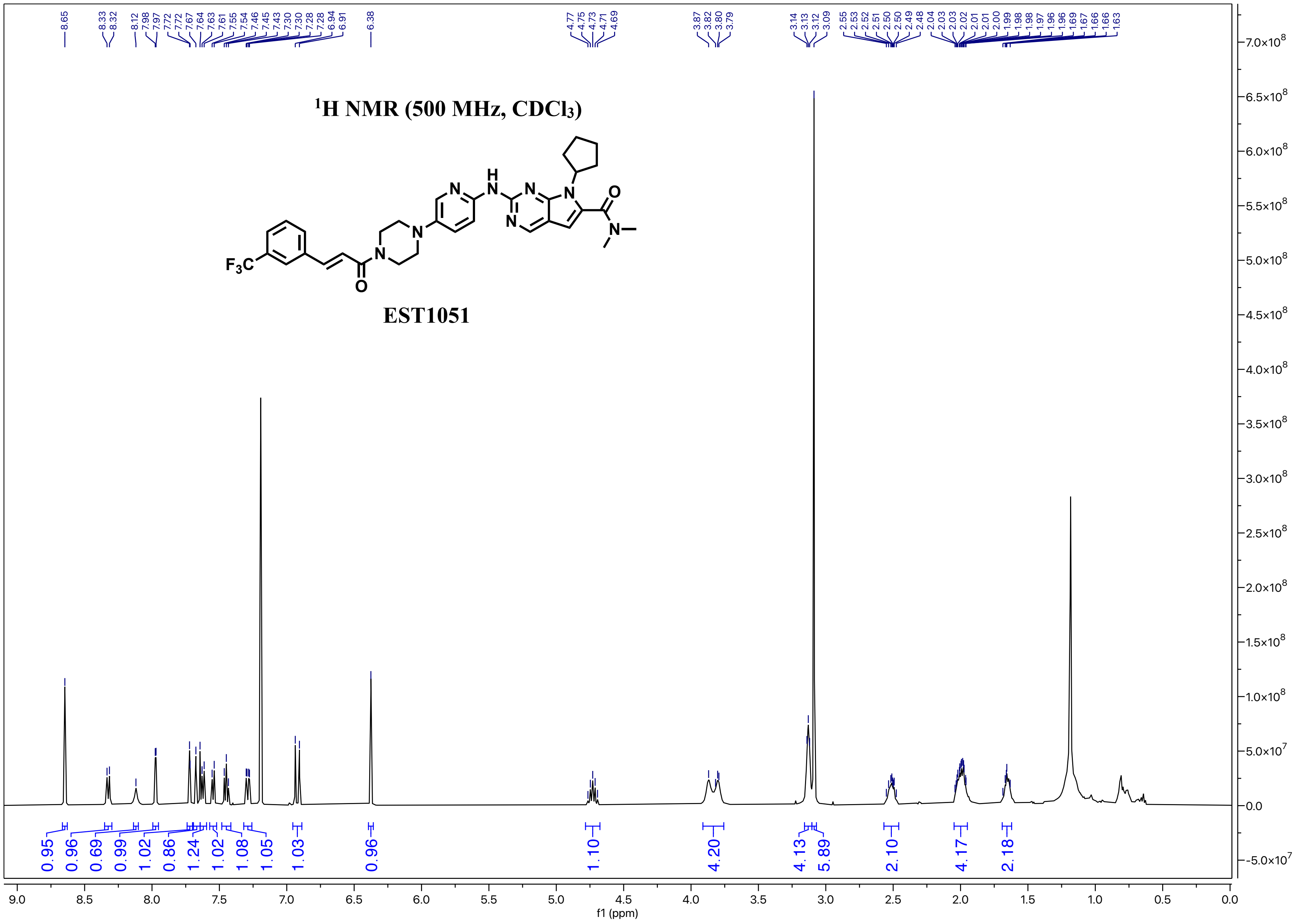

1H NMR (500 MHz, CDCl3)
EST1051
f1 (ppm)

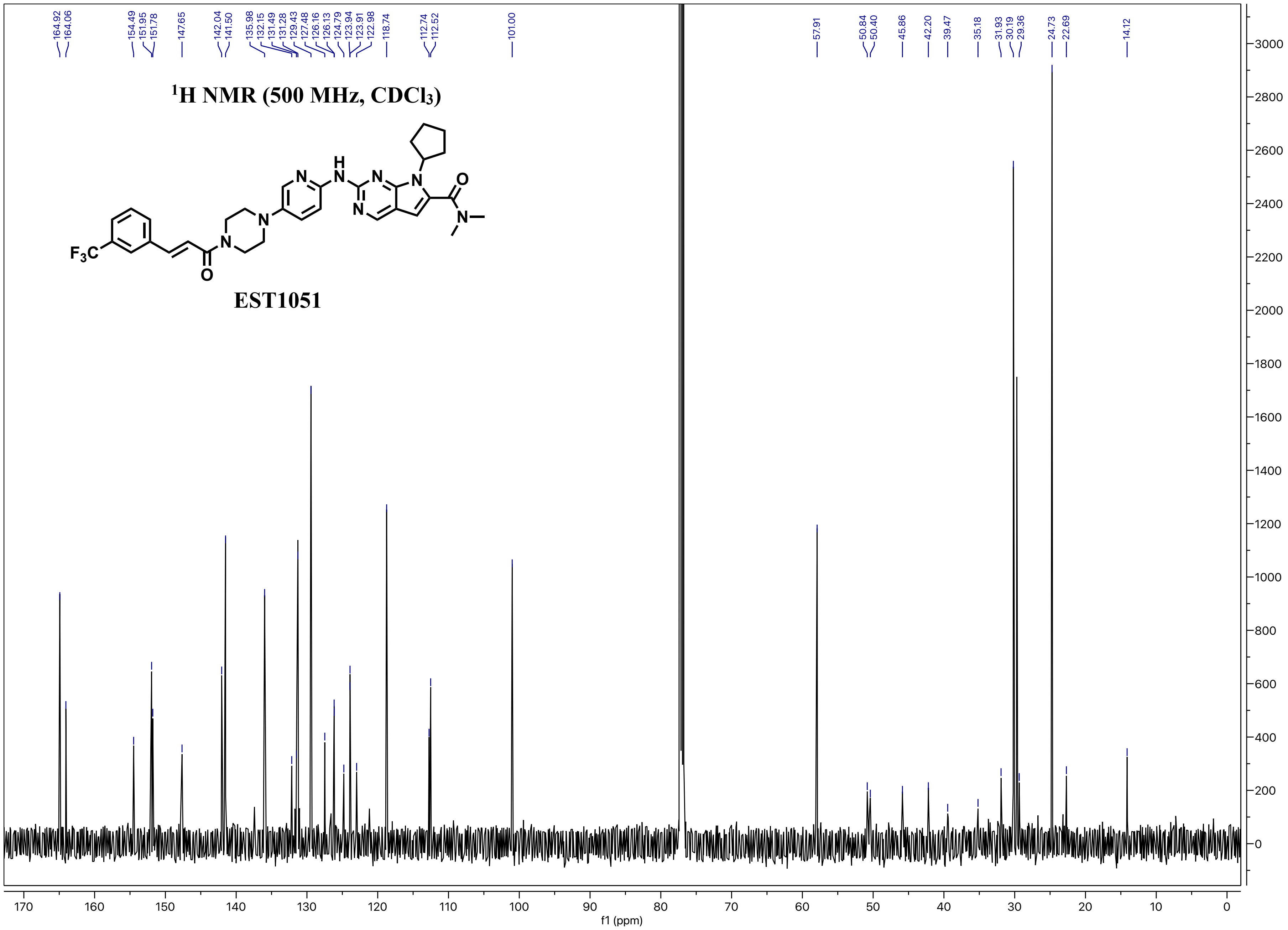
164.92
164.06
154.49
151.95
151.78
147.65
142.04
141.50
135.98
132.15
131.49
131.28
129.43
127.48
126.16
126.13
124.79
123.94
123.91
122.98
118.74
112.74
112.52
101.00
57.91
50.84
50.40
45.86
42.20
39.47
35.18
31.93
30.19
29.36
24.73
22.69
14.12
$^{1}H$ NMR (500 MHz, $CDCl_3$)
F3C
N
N
H
N
N
N
O
N
EST1051
f1 (ppm)

### [1]H NMR (500 MHz, CDCl3)

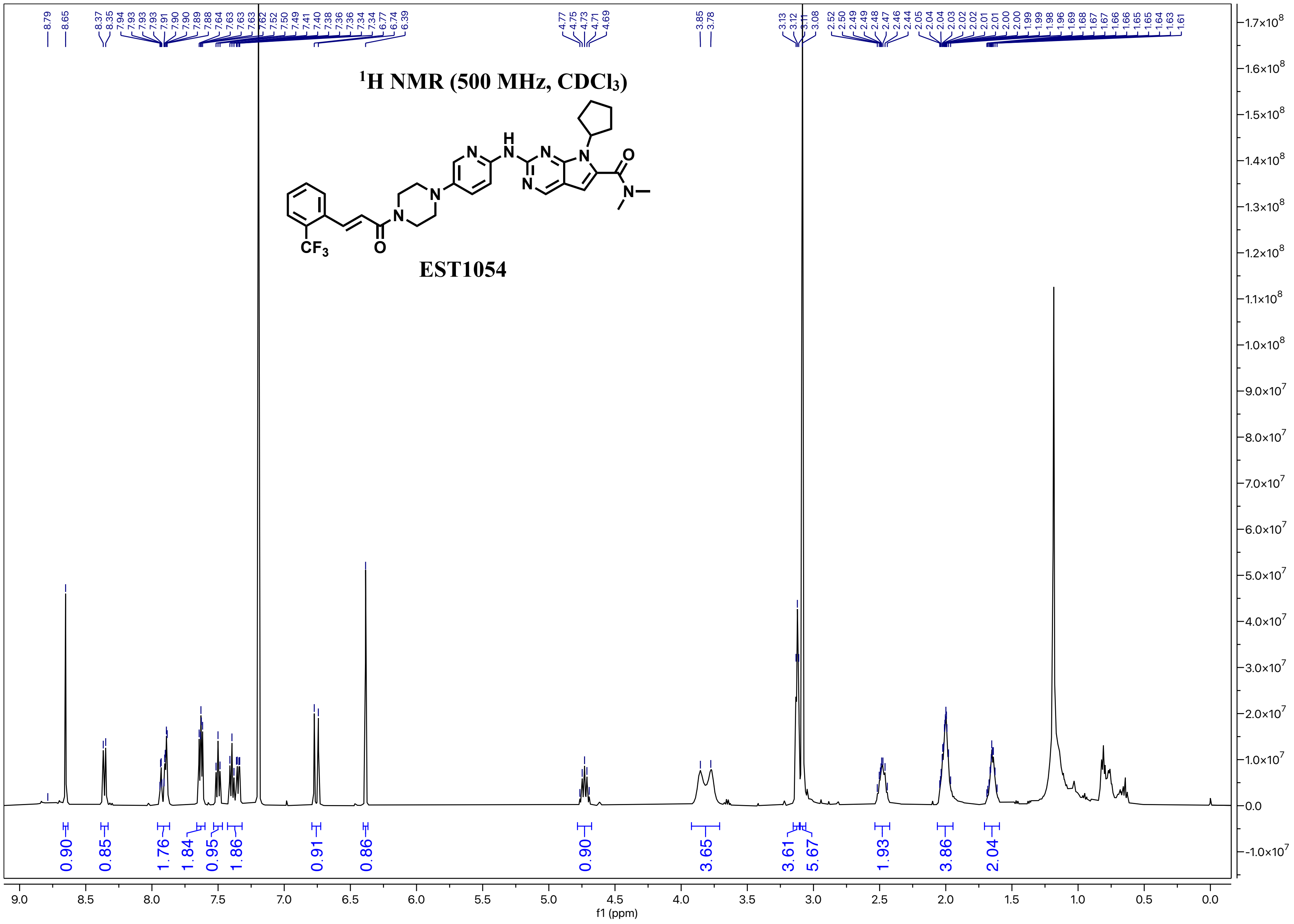


EST1054

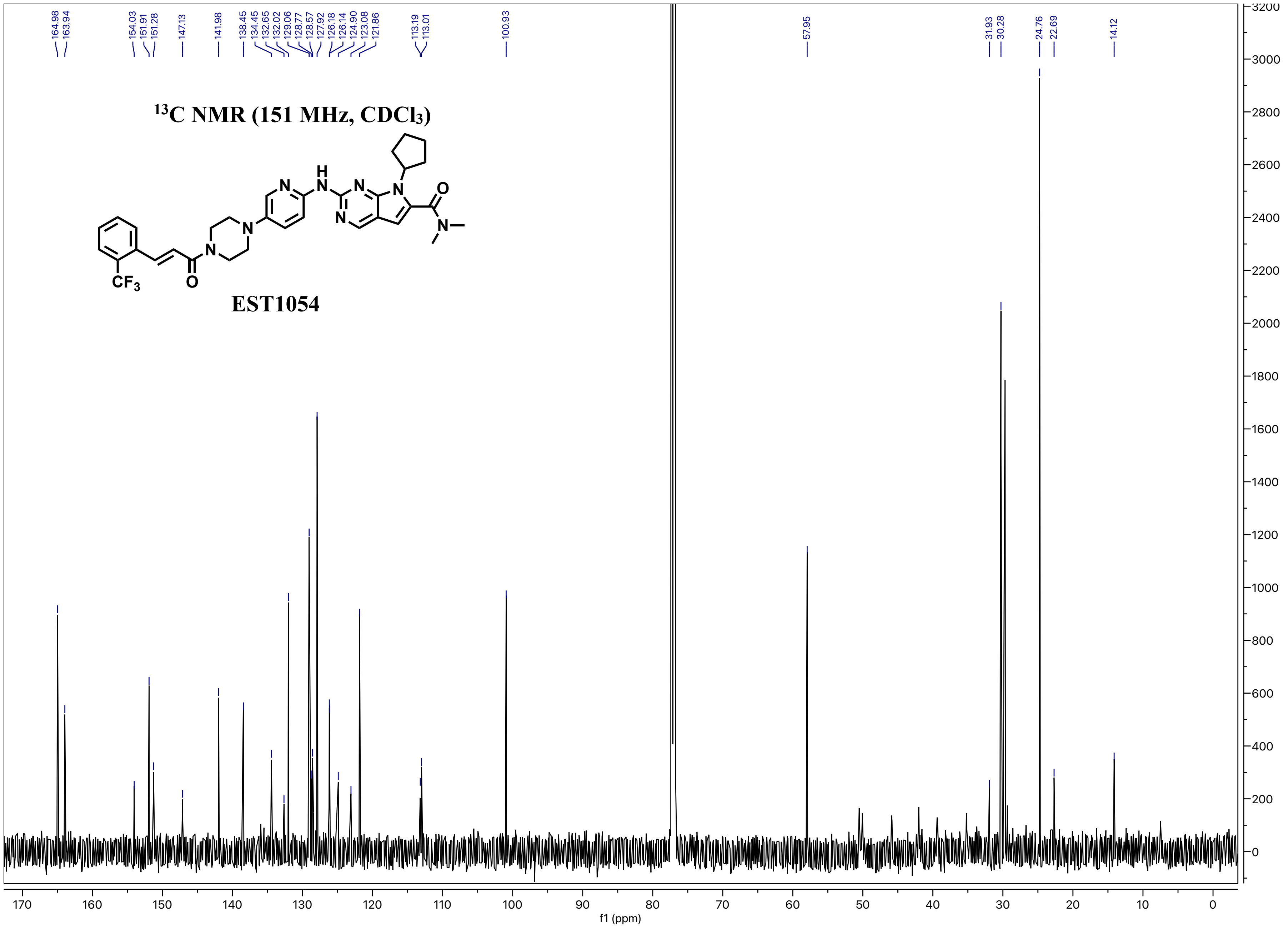

164.98
163.94
154.03
151.91
151.28
147.13
141.98
138.45
134.45
132.65
132.02
129.06
128.77
128.57
127.92
126.18
126.14
124.90
123.08
121.86
113.19
113.01
100.93
57.95
31.93
30.28
24.76
22.69
14.12
13C NMR (151 MHz, CDCl3)
EST1054
f1 (ppm)

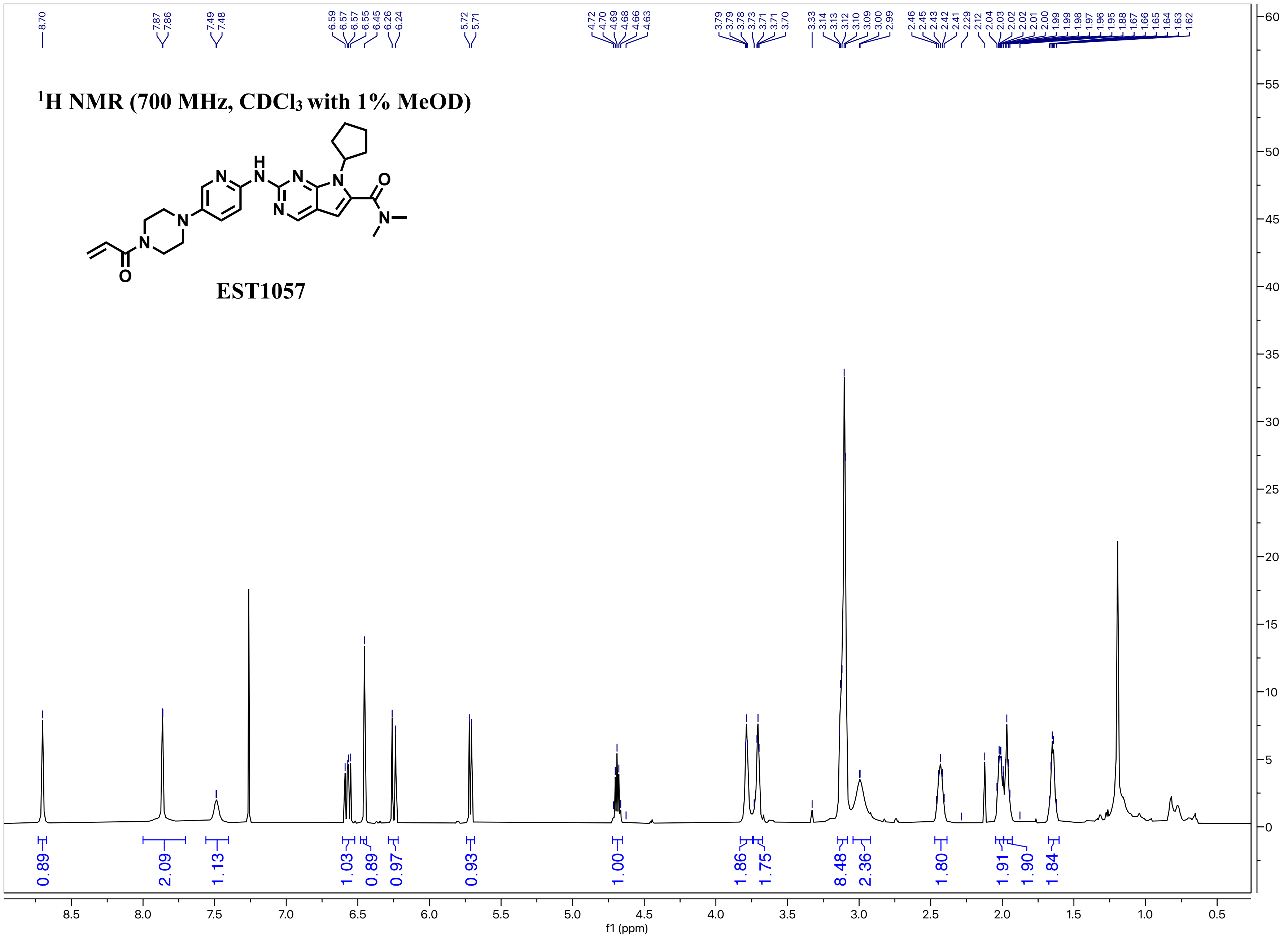

$^{1}$H NMR (700 MHz, $CDCl_3$ with 1% MeOD)
EST1057
8.70
7.87 7.86
7.49 7.48
6.59 6.57 6.57 6.55 6.45 6.26 6.24
5.72 5.71
4.72 4.70 4.69 4.68 4.66 4.63
3.79 3.79 3.78 3.73 3.71 3.71 3.70
3.33 3.14 3.13 3.12 3.10 3.09 3.00 2.99
2.46 2.45 2.43 2.42 2.41 2.29 2.12 2.04 2.03 2.02 2.02 2.01 2.00 1.99 1.99 1.98 1.97 1.96 1.95 1.88 1.67 1.66 1.65 1.64 1.63 1.62
0.89 2.09 1.13 1.03 0.89 0.97 0.93 1.00 1.86 1.75 8.48 2.36 1.80 1.91 1.90 1.84
f1 (ppm)

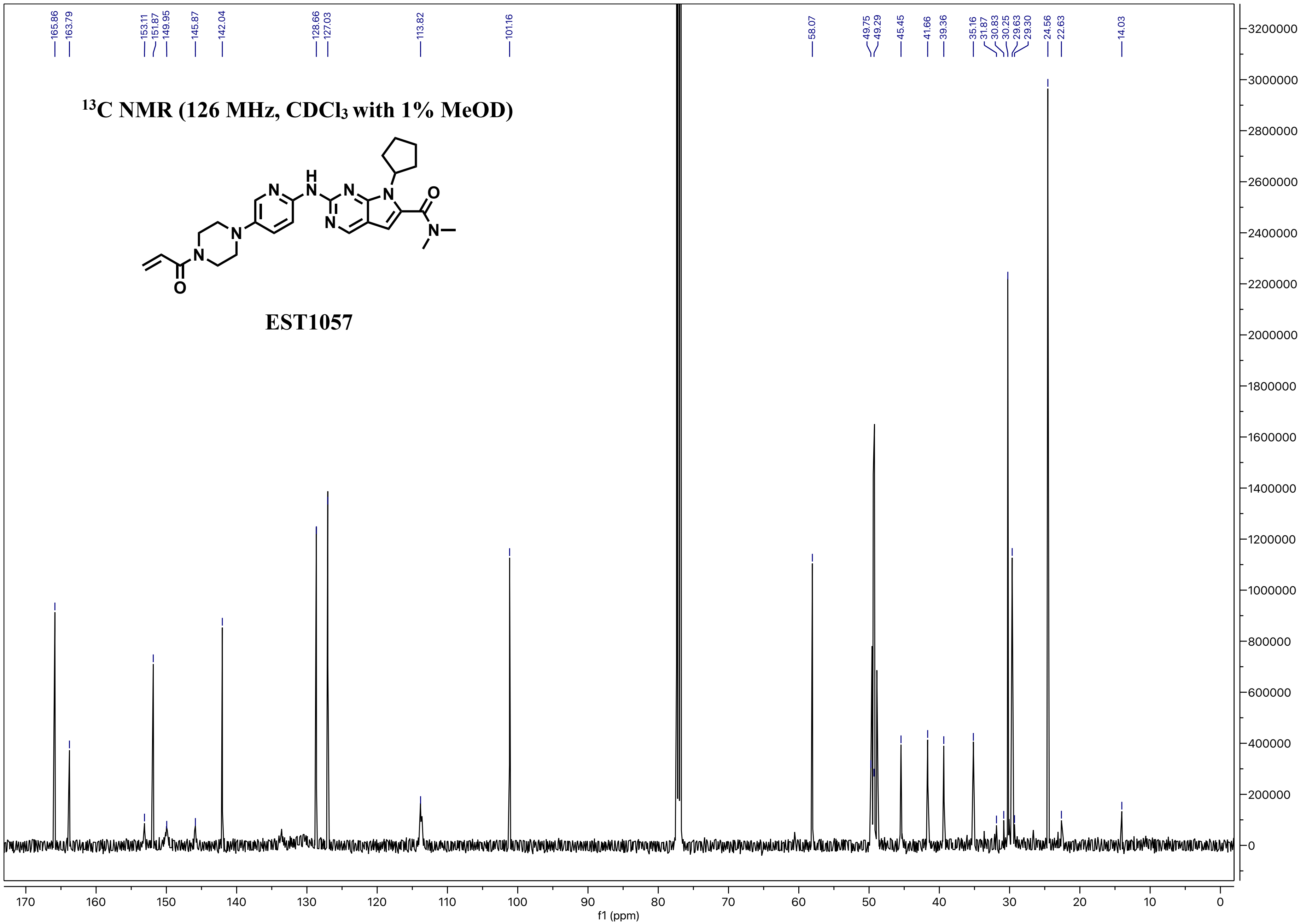

$^{13}C$ NMR (126 MHz, $CDCl_3$ with 1% MeOD)
EST1057
165.86
163.79
153.11
151.87
149.95
145.87
142.04
128.66
127.03
113.82
101.16
58.07
49.75
49.29
45.45
41.66
39.36
35.16
31.87
30.83
30.25
29.63
29.30
24.56
22.63
14.03
f1 (ppm)

**$^1$H NMR (700 MHz, DMSO-d$_6$)**

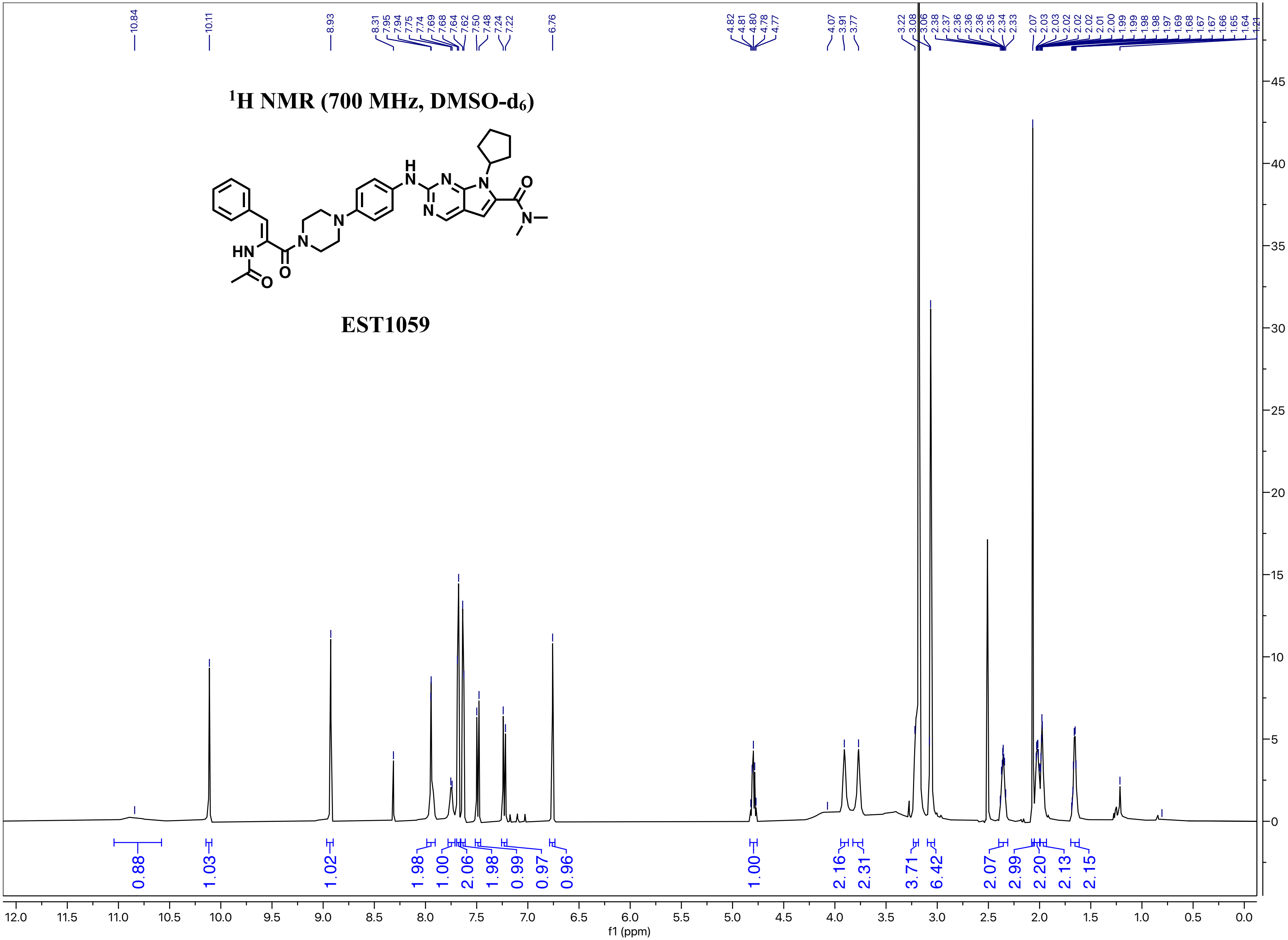


**EST1059**

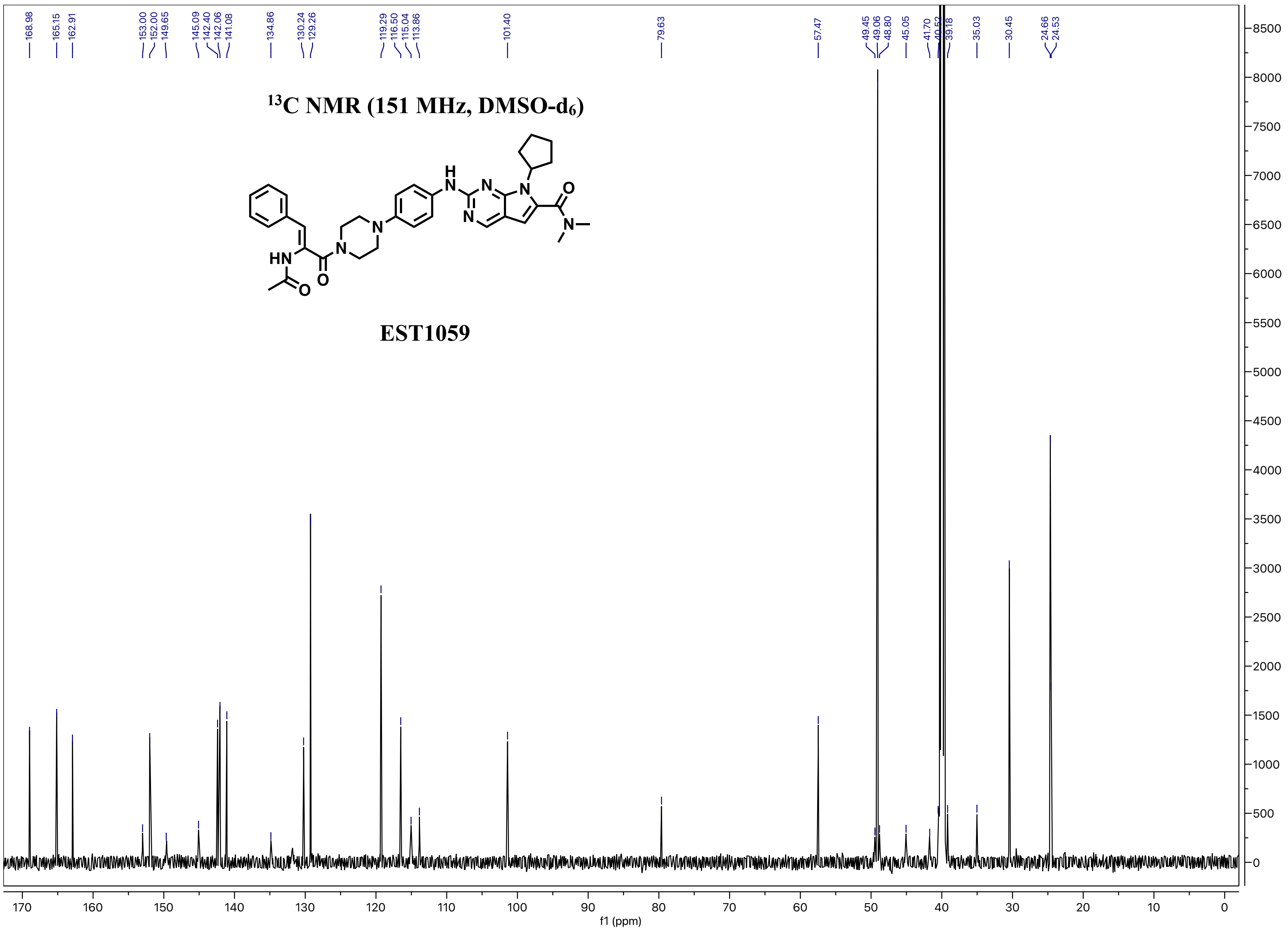

168.98
165.15
162.91
153.00
152.00
149.65
145.09
142.40
142.06
141.08
134.86
130.24
129.26
119.29
116.50
115.04
113.86
101.40
79.63
57.47
49.45
49.06
48.80
45.05
41.70
40.52
39.18
35.03
30.45
24.66
24.53
13C NMR (151 MHz, DMSO-d6)
EST1059
f1 (ppm)
170 160 150 140 130 120 110 100 90 80 70 60 50 40 30 20 10 0
8500 8000 7500 7000 6500 6000 5500 5000 4500 4000 3500 3000 2500 2000 1500 1000 500 0

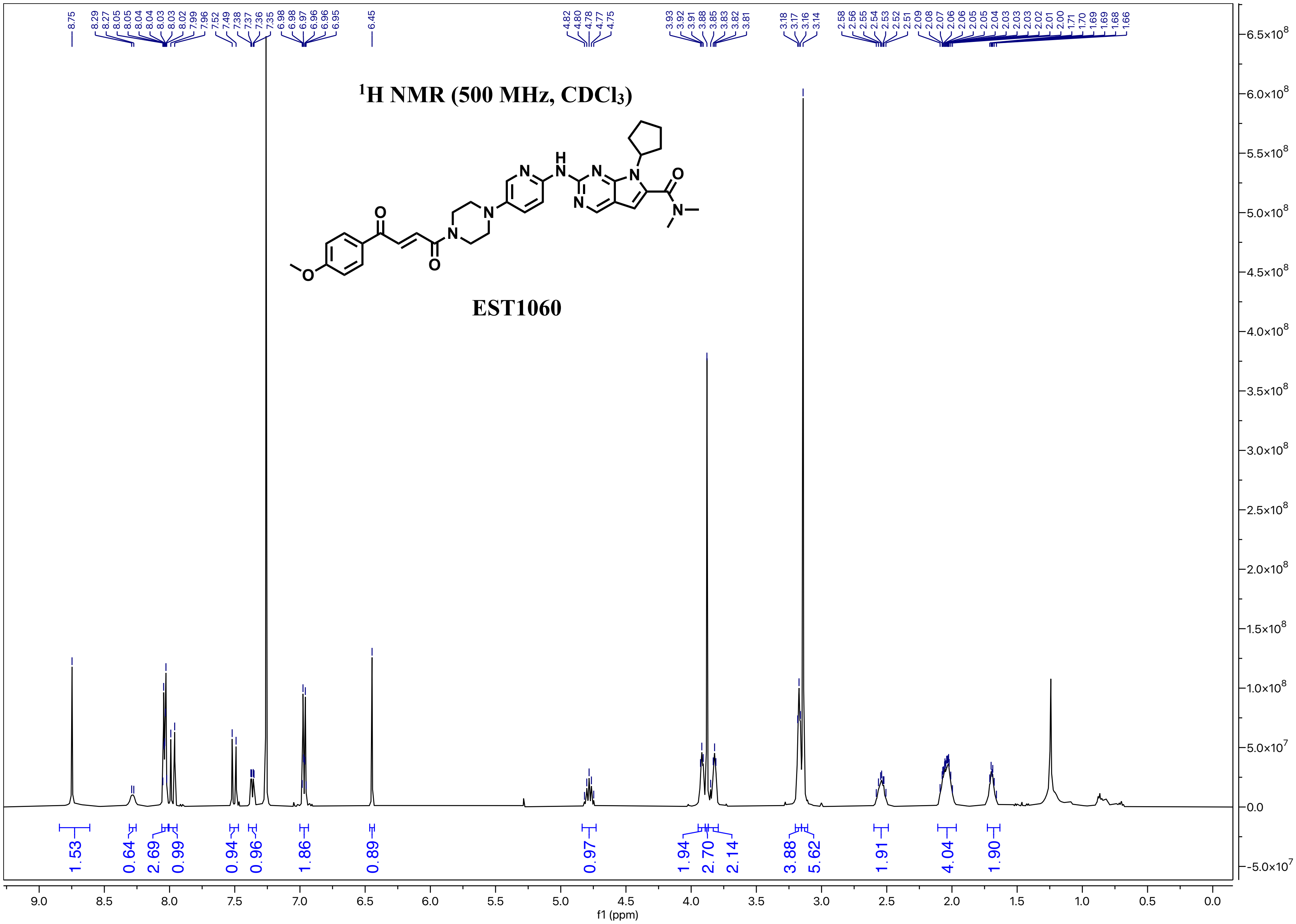

1H NMR (500 MHz, CDCl3)
EST1060
f1 (ppm)

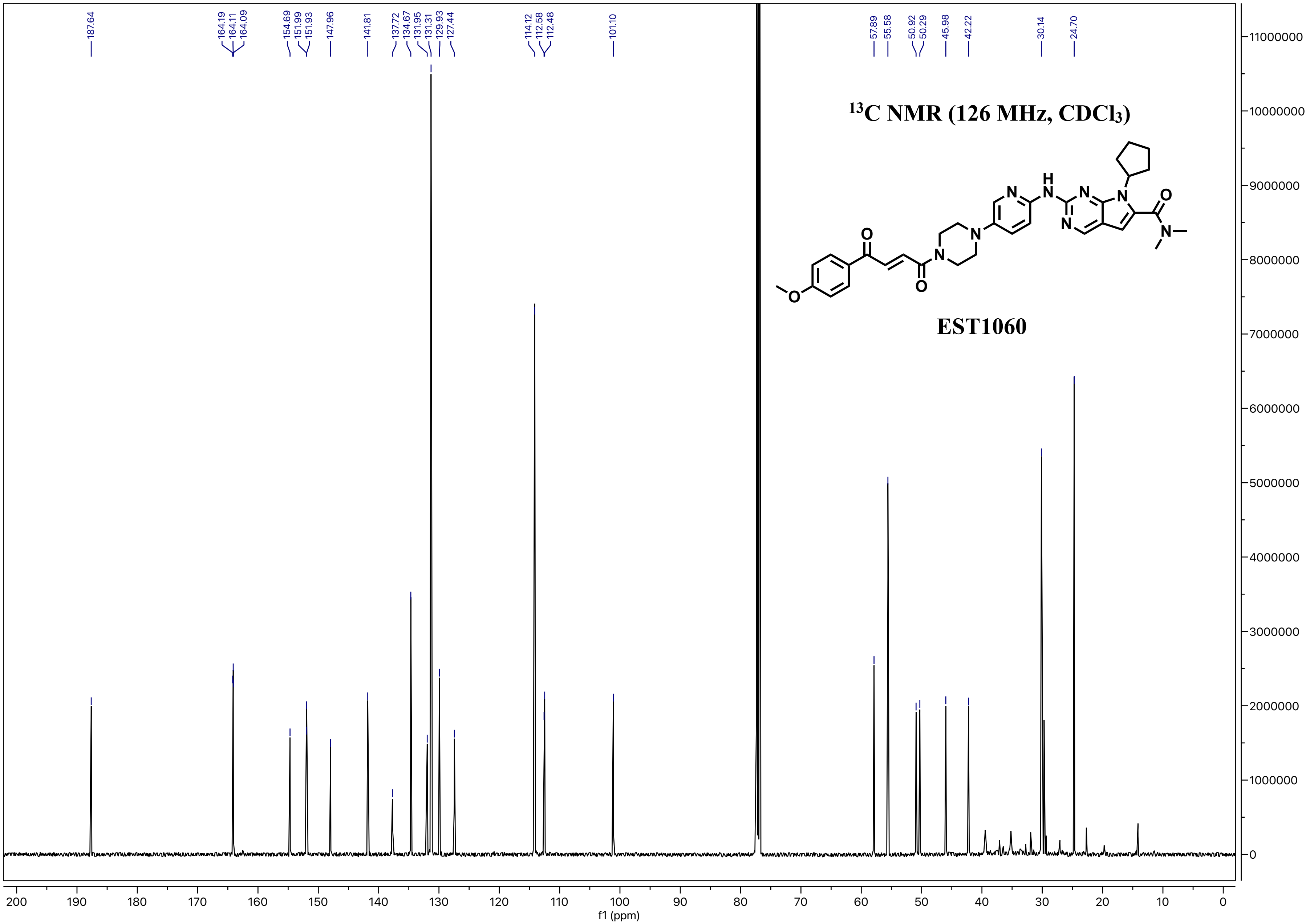

187.64
164.19
164.11
164.09
154.69
151.99
151.93
147.96
141.81
137.72
134.67
131.95
131.31
129.93
127.44
114.12
112.58
112.48
101.10
57.89
55.58
50.92
50.29
45.98
42.22
30.14
24.70
$^{13}$C NMR (126 MHz, CDCl$_3$)
EST1060
f1 (ppm)

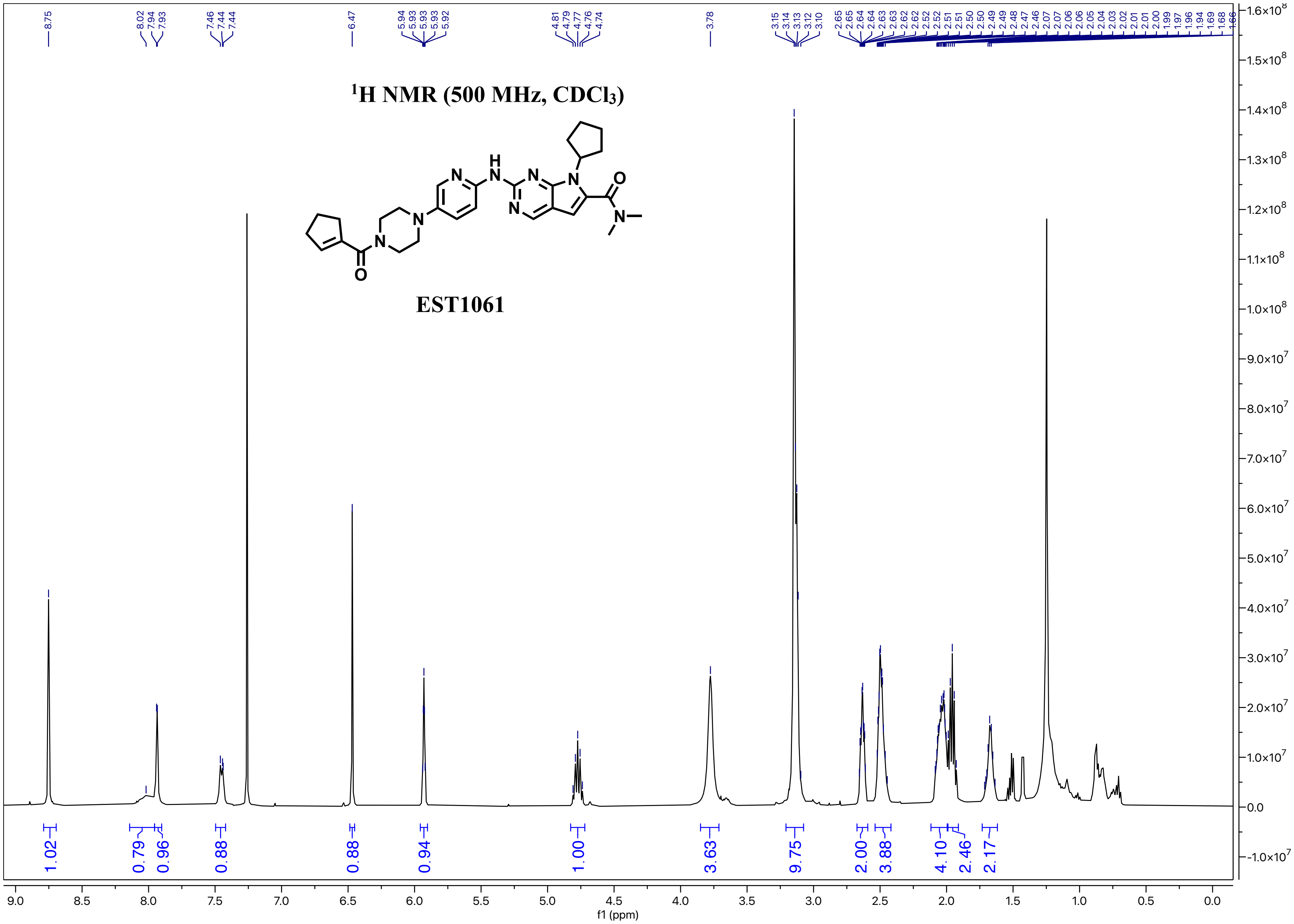

1H NMR (500 MHz, CDCl3)
EST1061
f1 (ppm)

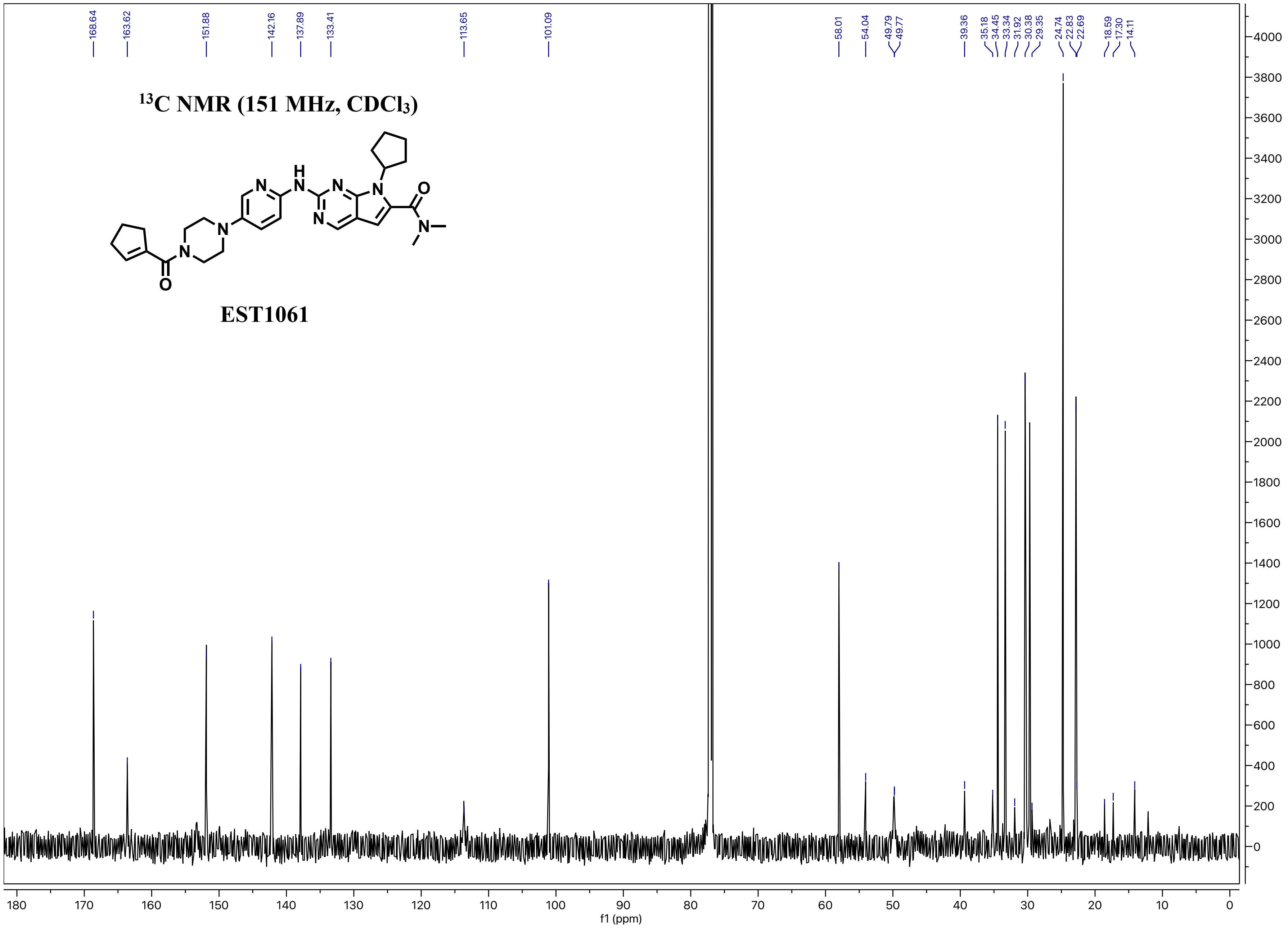

13C NMR (151 MHz, CDCl3)
EST1061
168.64
163.62
151.88
142.16
137.89
133.41
113.65
101.09
58.01
54.04
49.79
49.77
39.36
35.18
34.45
33.34
31.92
30.38
29.35
24.74
22.83
22.69
18.59
17.30
14.11
f1 (ppm)

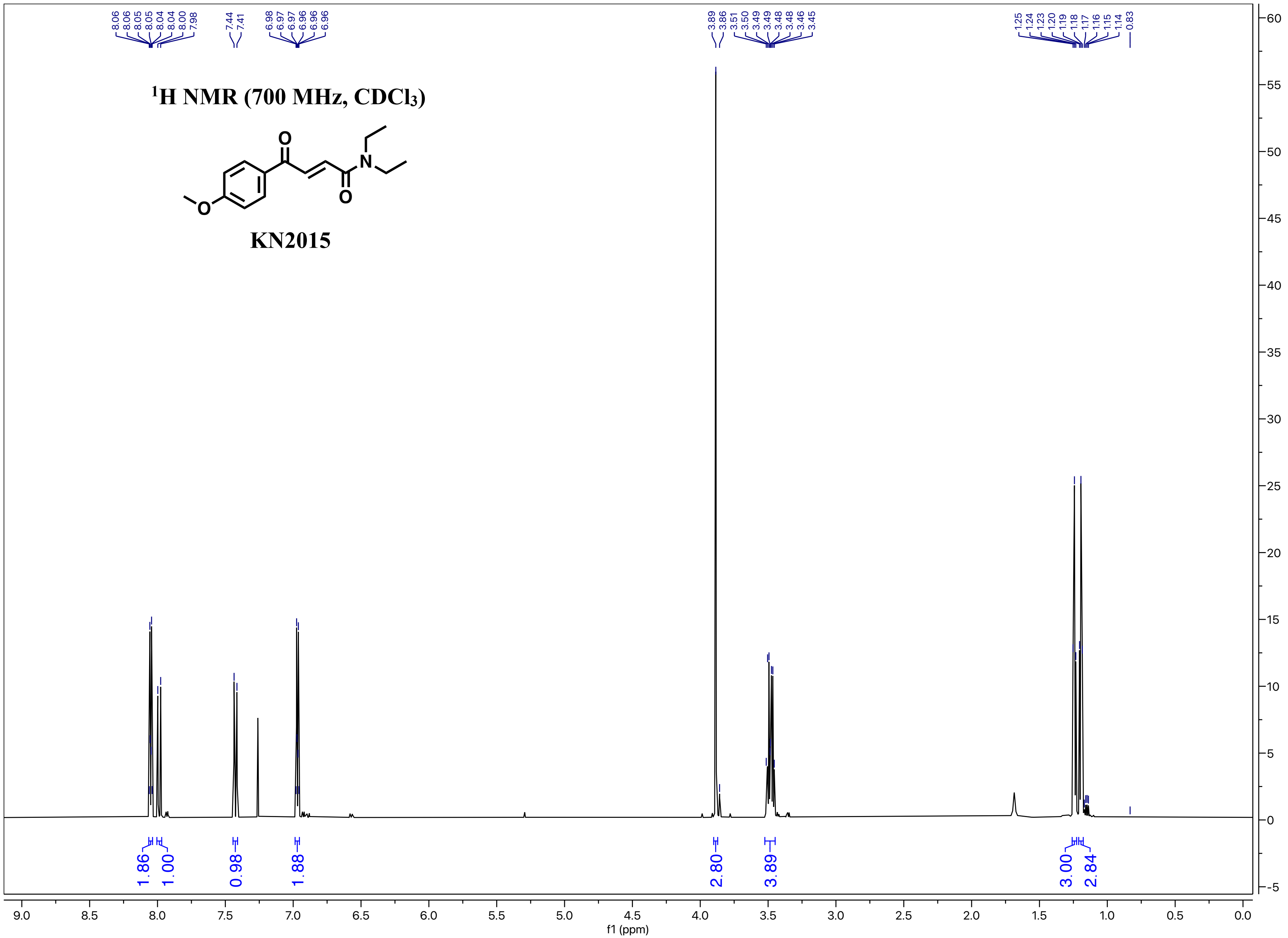

¹H NMR (700 MHz, CDCl₃)
KN2015
8.06 8.06 8.05 8.04 8.04 8.00 7.98
7.44 7.41
6.98 6.97 6.97 6.96 6.96 6.96
3.89 3.86 3.51 3.50 3.49 3.49 3.48 3.46 3.45
1.25 1.24 1.23 1.20 1.19 1.18 1.17 1.16 1.15 1.14 0.83
1.86 1.00 0.98 1.88 2.80 3.89 3.00 2.84
f1 (ppm)

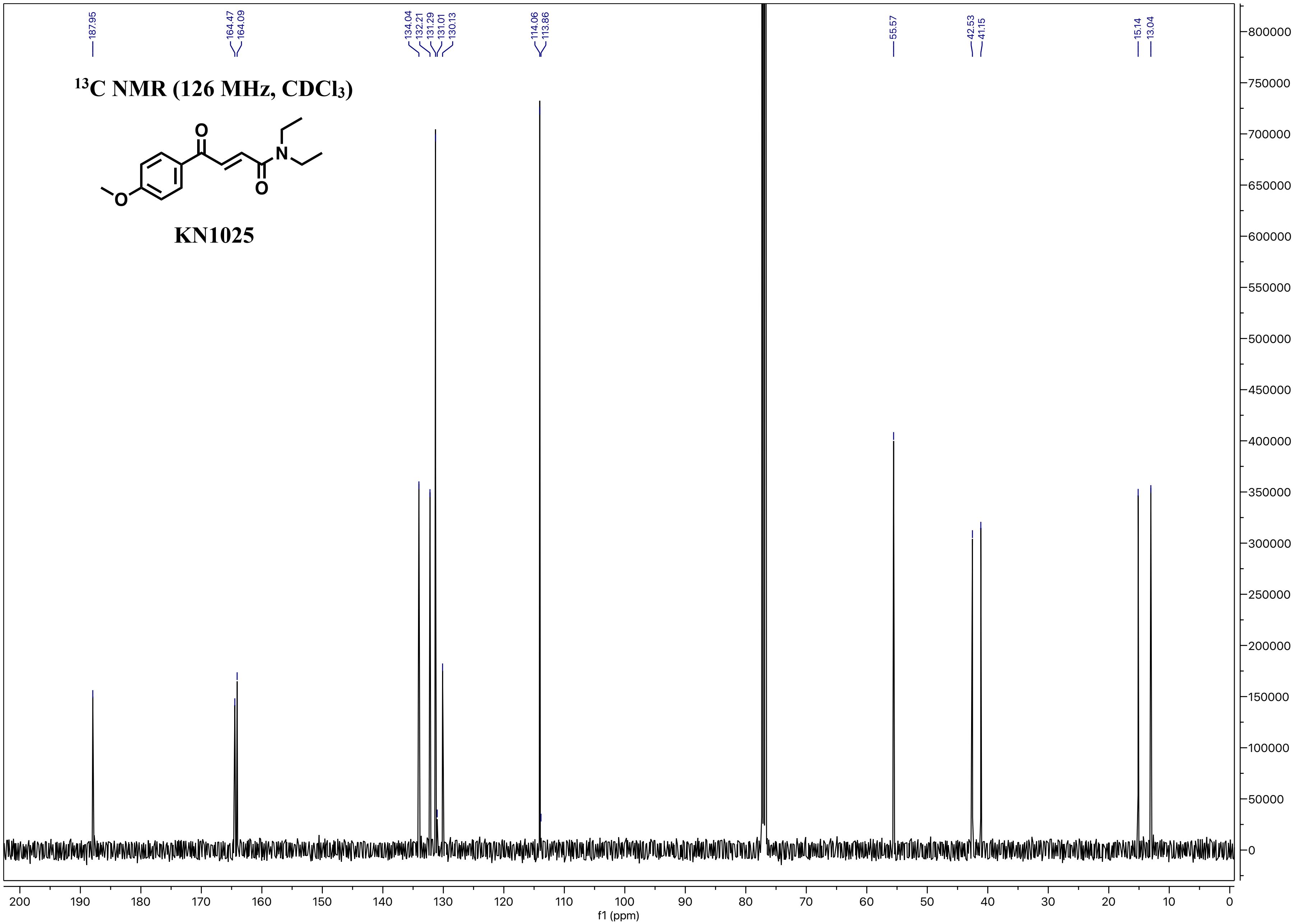

187.95
164.47
164.09
134.04
132.21
131.29
131.01
130.13
114.06
113.86
55.57
42.53
41.15
15.14
13.04
13C NMR (126 MHz, CDCl3)
KN1025
f1 (ppm)

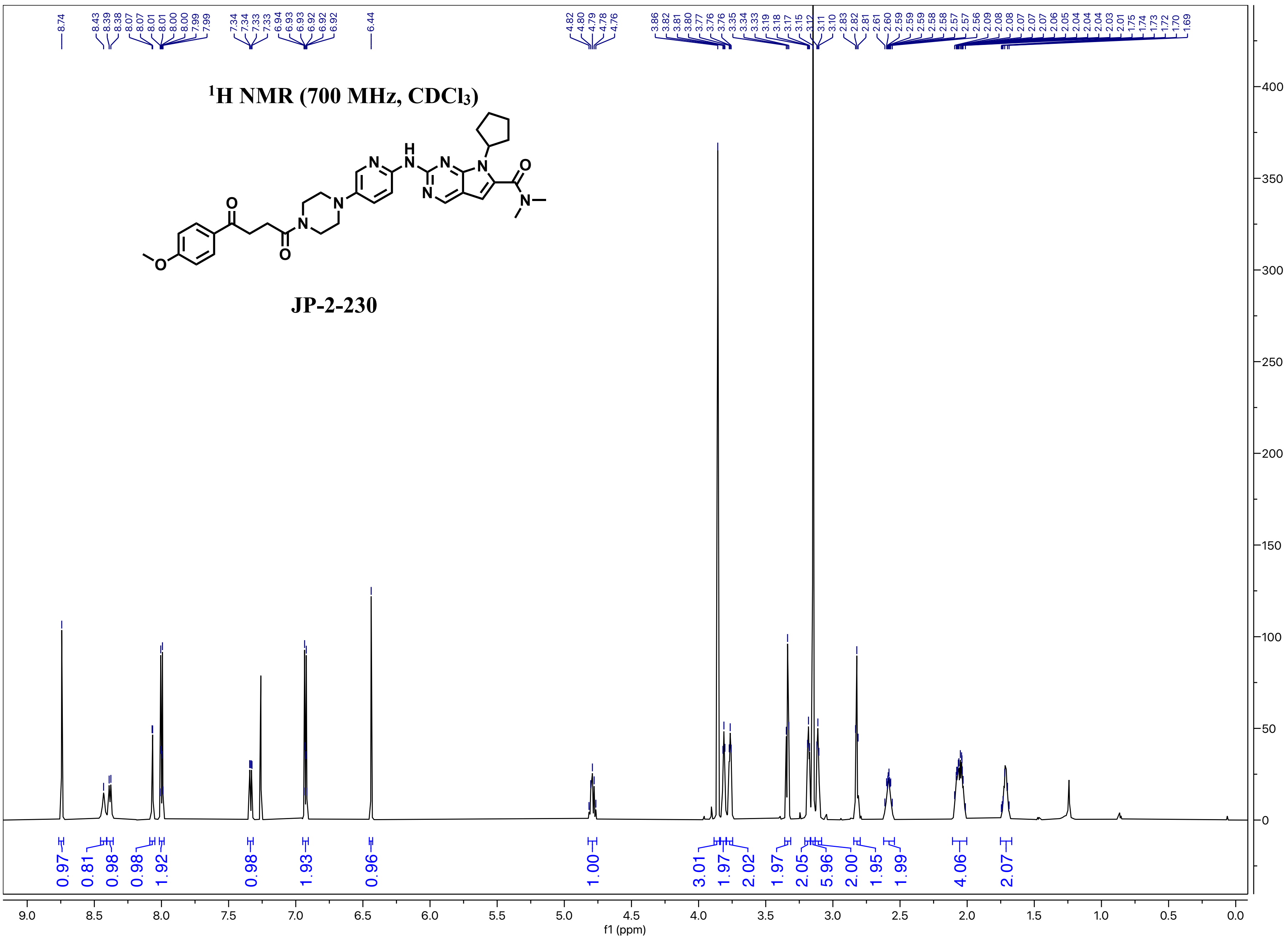

1H NMR (700 MHz, CDCl3)
JP-2-230
0.97
0.81
0.98
0.98
1.92
0.98
1.93
0.96
1.00
3.01
1.97
2.02
1.97
2.05
5.96
2.00
1.95
1.99
4.06
2.07
f1 (ppm)

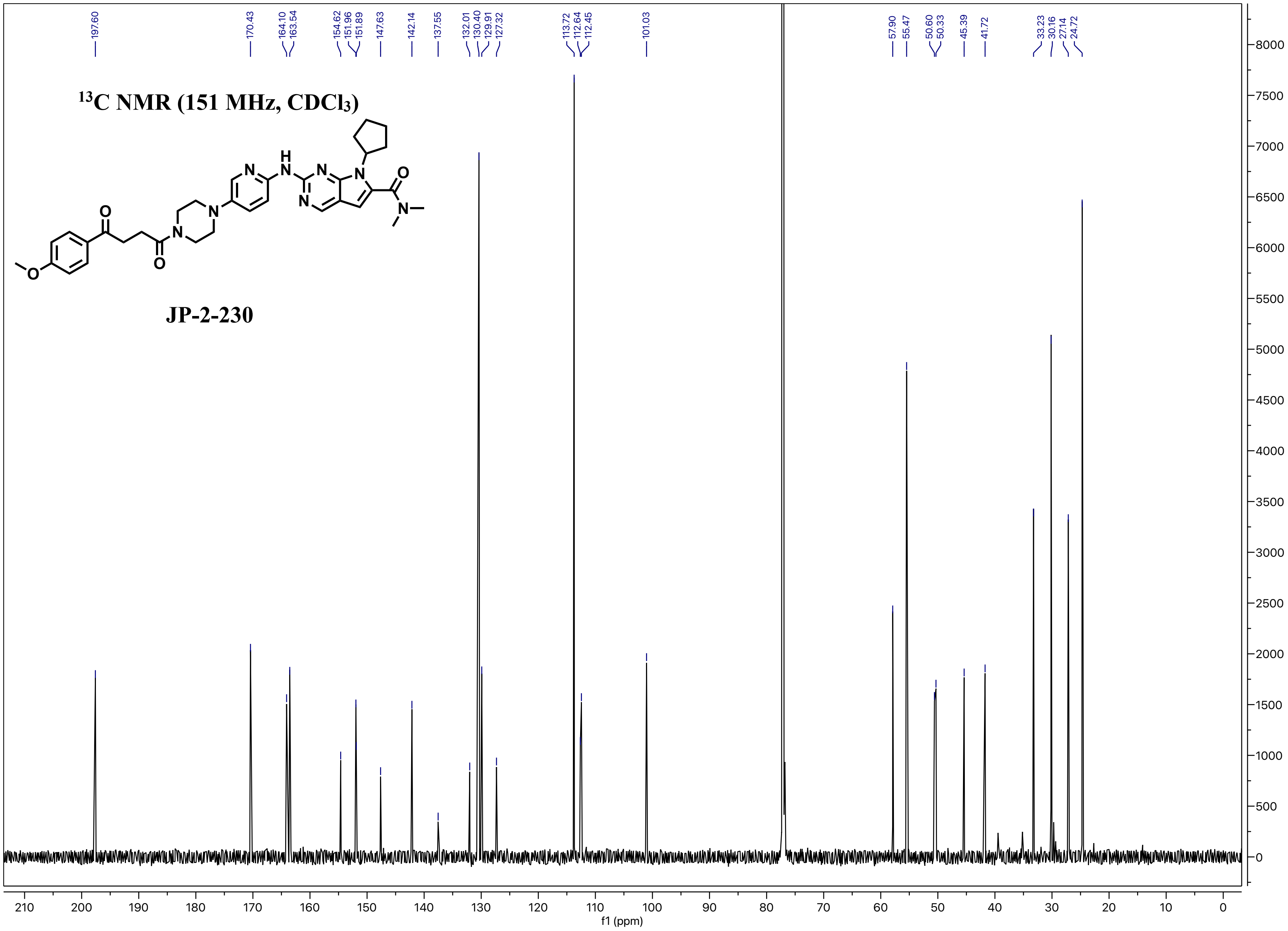
$^{13}C$ NMR (151 MHz, $CDCl_3$)
JP-2-230
197.60
170.43
164.10
163.54
154.62
151.96
151.89
147.63
142.14
137.55
132.01
130.40
129.91
127.32
113.72
112.64
112.45
101.03
57.90
55.47
50.60
50.33
45.39
41.72
33.23
30.16
27.14
24.72
f1 (ppm)

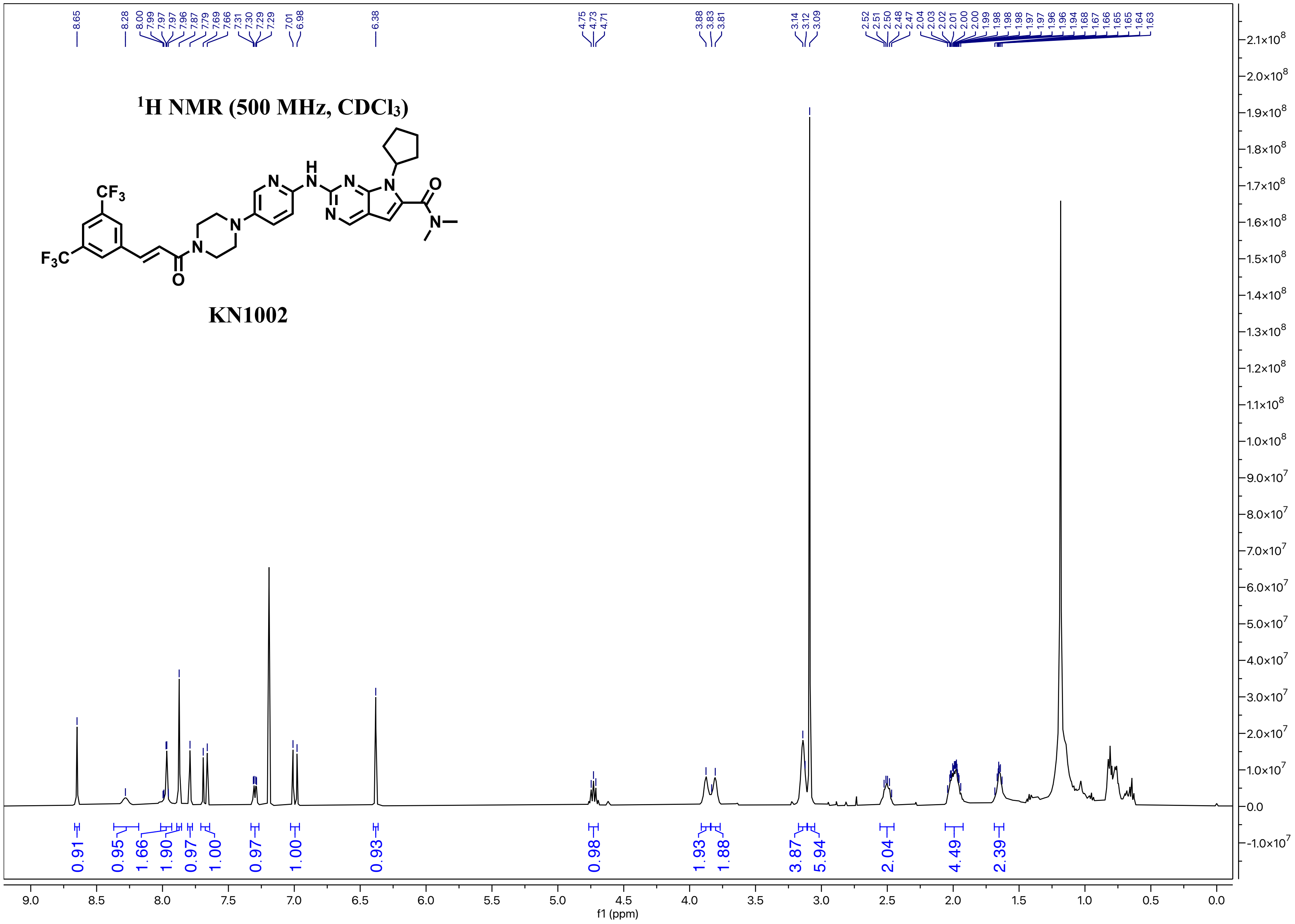

8.65 8.28 8.00 7.99 7.97 7.97 7.96 7.87 7.79 7.69 7.66 7.31 7.30 7.29 7.29 7.01 6.98 6.38 4.75 4.73 4.71 3.88 3.83 3.81 3.14 3.12 3.09 2.52 2.51 2.50 2.48 2.47 2.04 2.03 2.02 2.01 2.00 1.99 1.98 1.98 1.98 1.97 1.97 1.96 1.96 1.94 1.68 1.67 1.66 1.65 1.64 1.63
¹H NMR (500 MHz, CDCl₃)
CF₃
F₃C
O
N
N
H
N
N
N
N
O
N
KN1002
0.91 0.95 1.66 1.90 0.97 1.00 0.97 1.00 0.93 0.98 1.93 1.88 3.87 5.94 2.04 4.49 2.39
f1 (ppm)

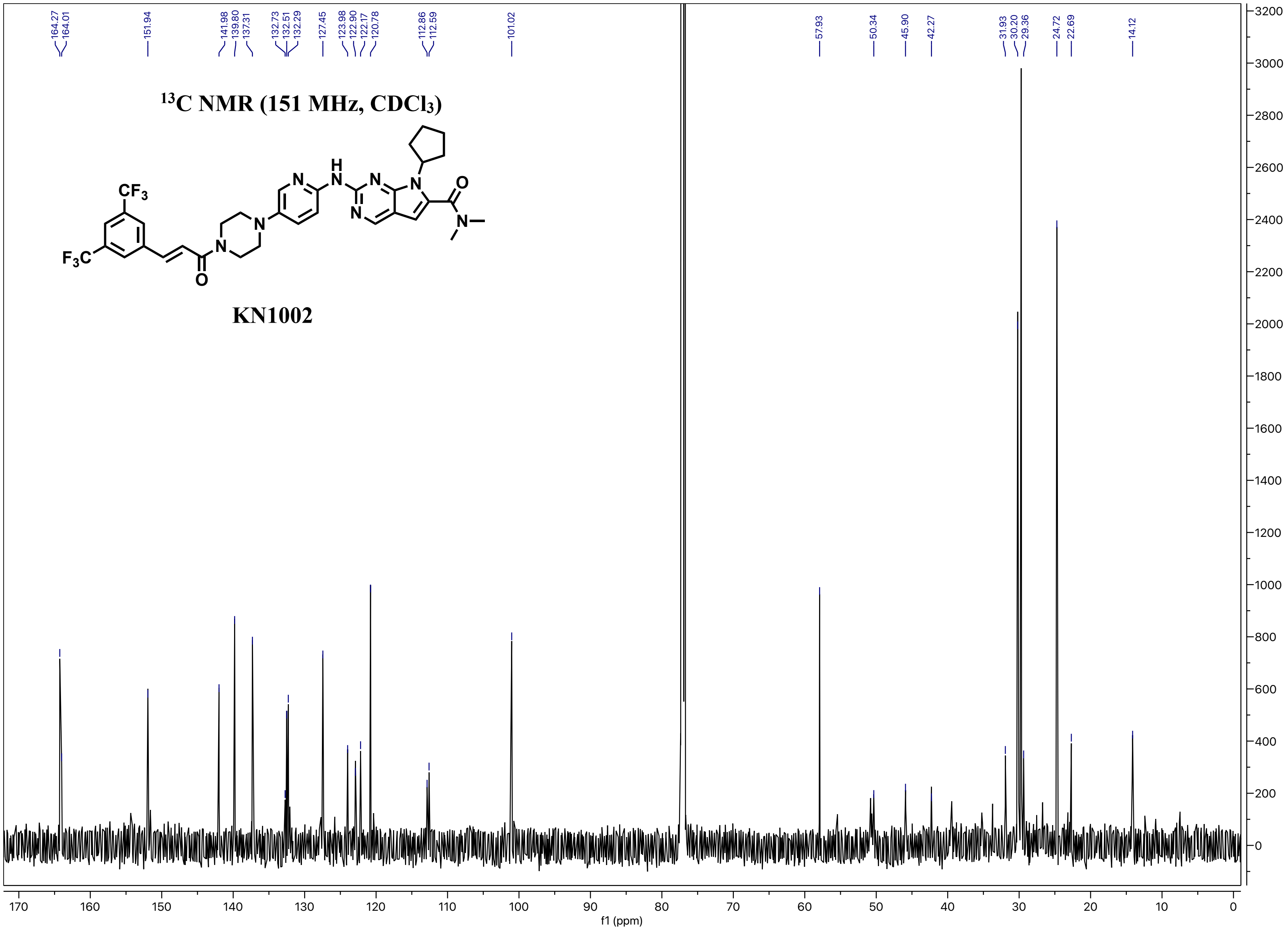

164.27
164.01
151.94
141.98
139.80
137.31
132.73
132.51
132.29
127.45
123.98
122.90
122.17
120.78
112.86
112.59
101.02
57.93
50.34
45.90
42.27
31.93
30.20
29.36
24.72
22.69
14.12
13C NMR (151 MHz, CDCl3)
KN1002
f1 (ppm)

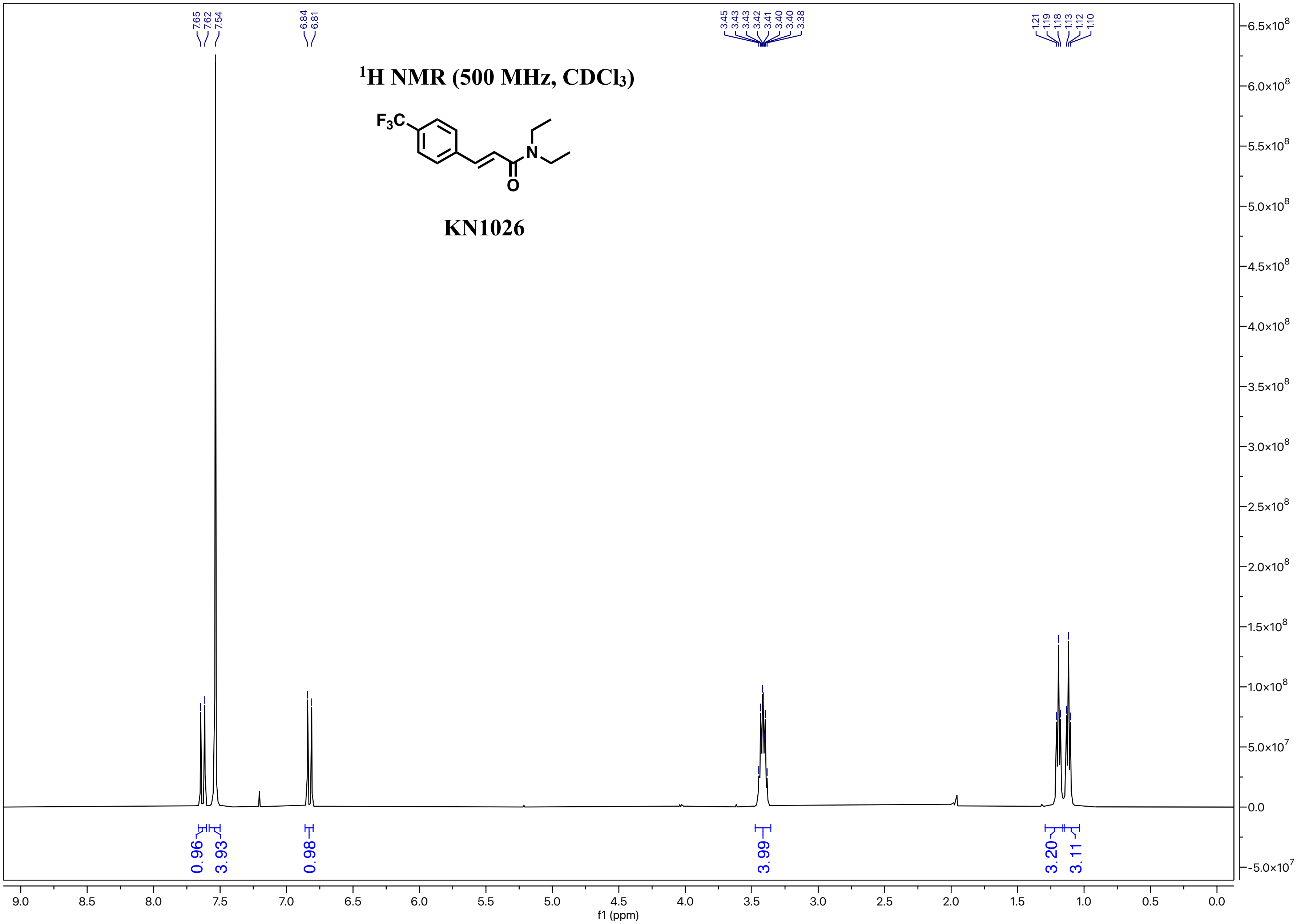

1H NMR (500 MHz, CDCl3)
F3C
N
O
KN1026
7.65
7.62
7.54
6.84
6.81
3.45
3.43
3.43
3.42
3.41
3.40
3.40
3.38
1.21
1.19
1.18
1.13
1.12
1.10
0.96
3.93
0.98
3.99
3.20
3.11
9.0
8.5
8.0
7.5
7.0
6.5
6.0
5.5
5.0
4.5
4.0
3.5
3.0
2.5
2.0
1.5
1.0
0.5
0.0
f1 (ppm)
6.5×10^8
6.0×10^8
5.5×10^8
5.0×10^8
4.5×10^8
4.0×10^8
3.5×10^8
3.0×10^8
2.5×10^8
2.0×10^8
1.5×10^8
1.0×10^8
5.0×10^7
0.0
-5.0×10^7

$^{13}$C NMR (126 MHz, $CDCl_3$) δ 165.1, 140.5, 138.9, 131.0 (q, $J$ = 32.5 Hz), 127.9, 125.7 (q, $J$ = 3.8 Hz), 123.9 (q, $J$ = 272.1 Hz), 120.3, 42.4, 41.2, 15.1, 13.1.

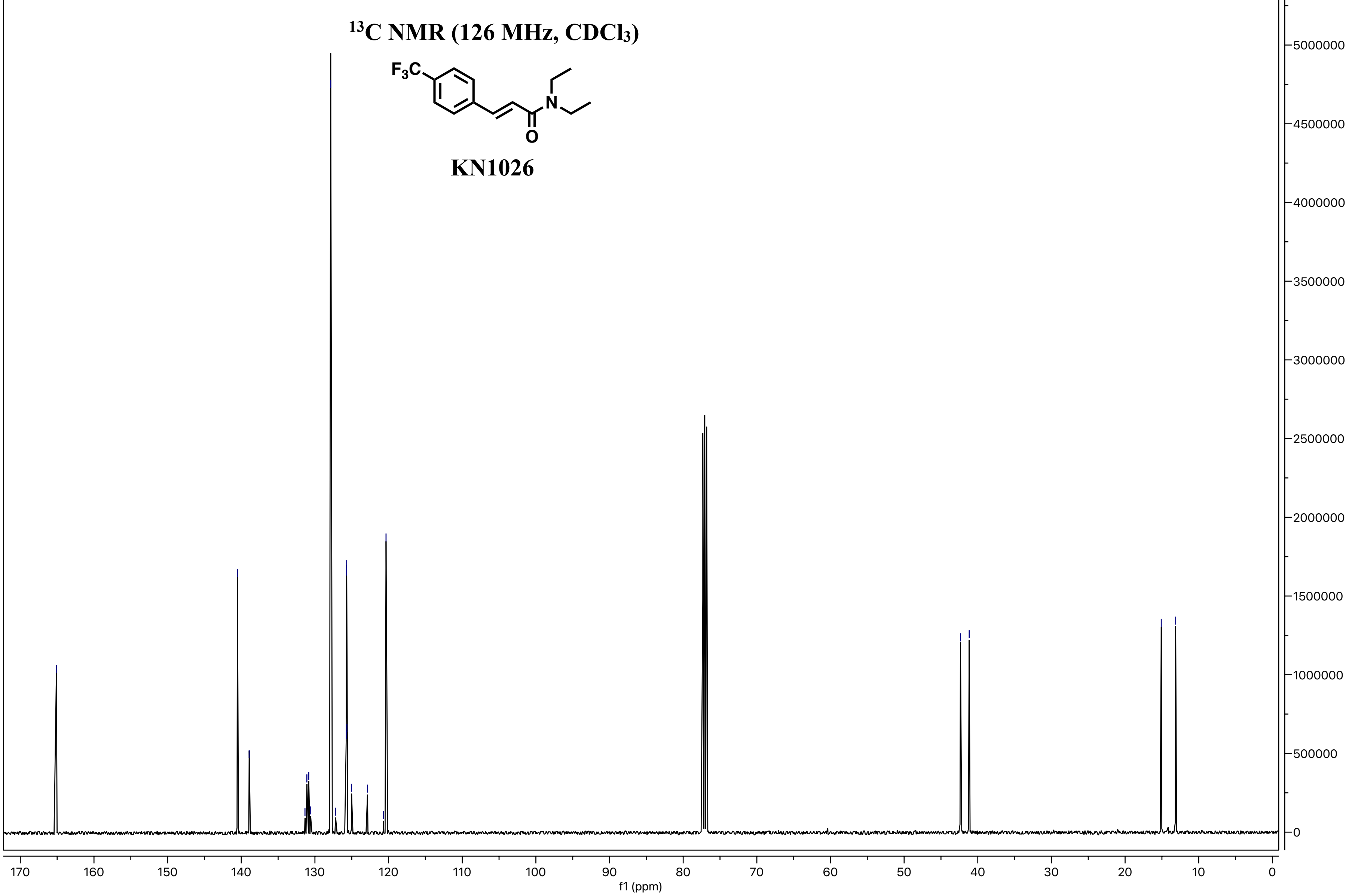

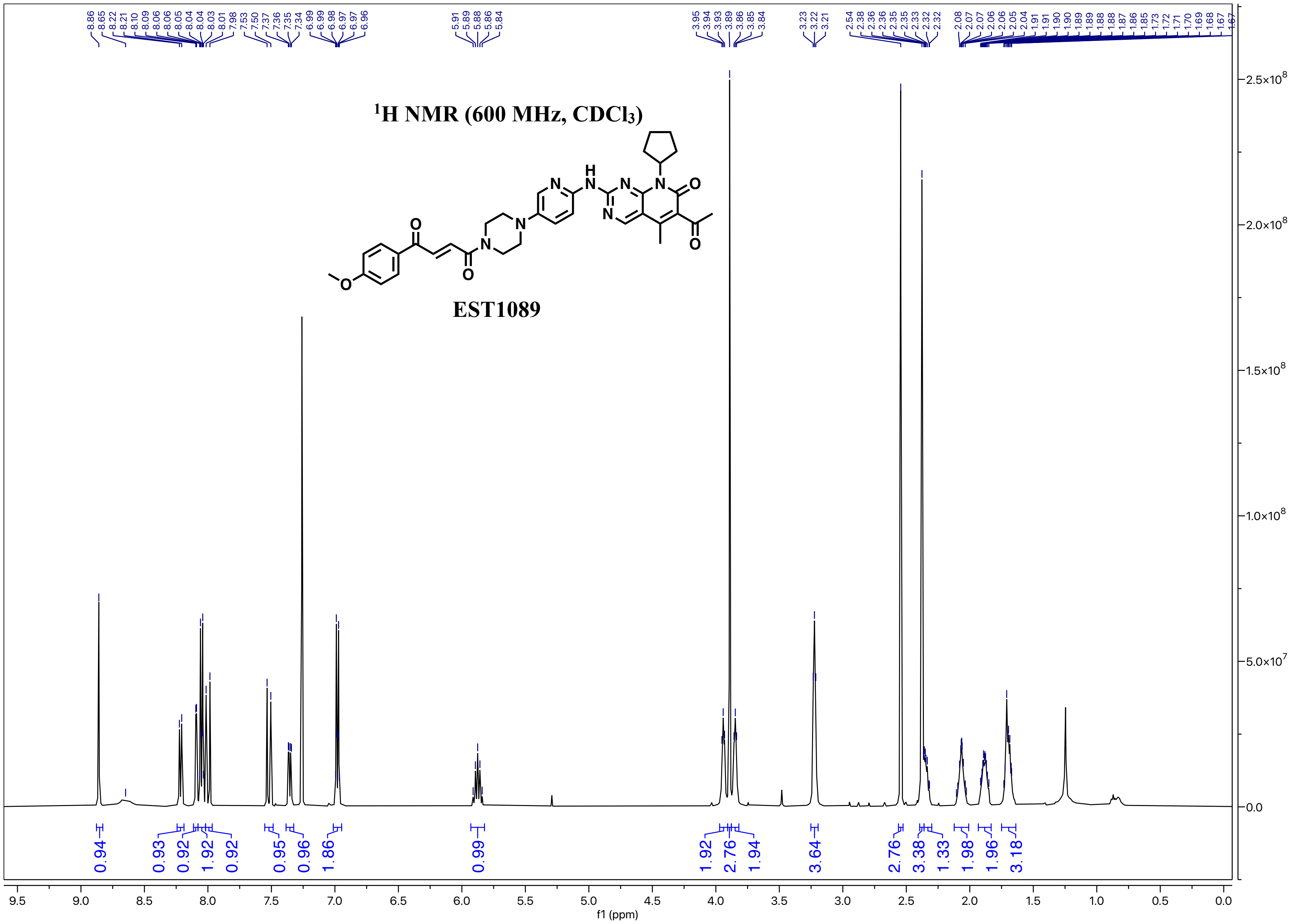

¹H NMR (600 MHz, CDCl₃)
EST1089
f1 (ppm)

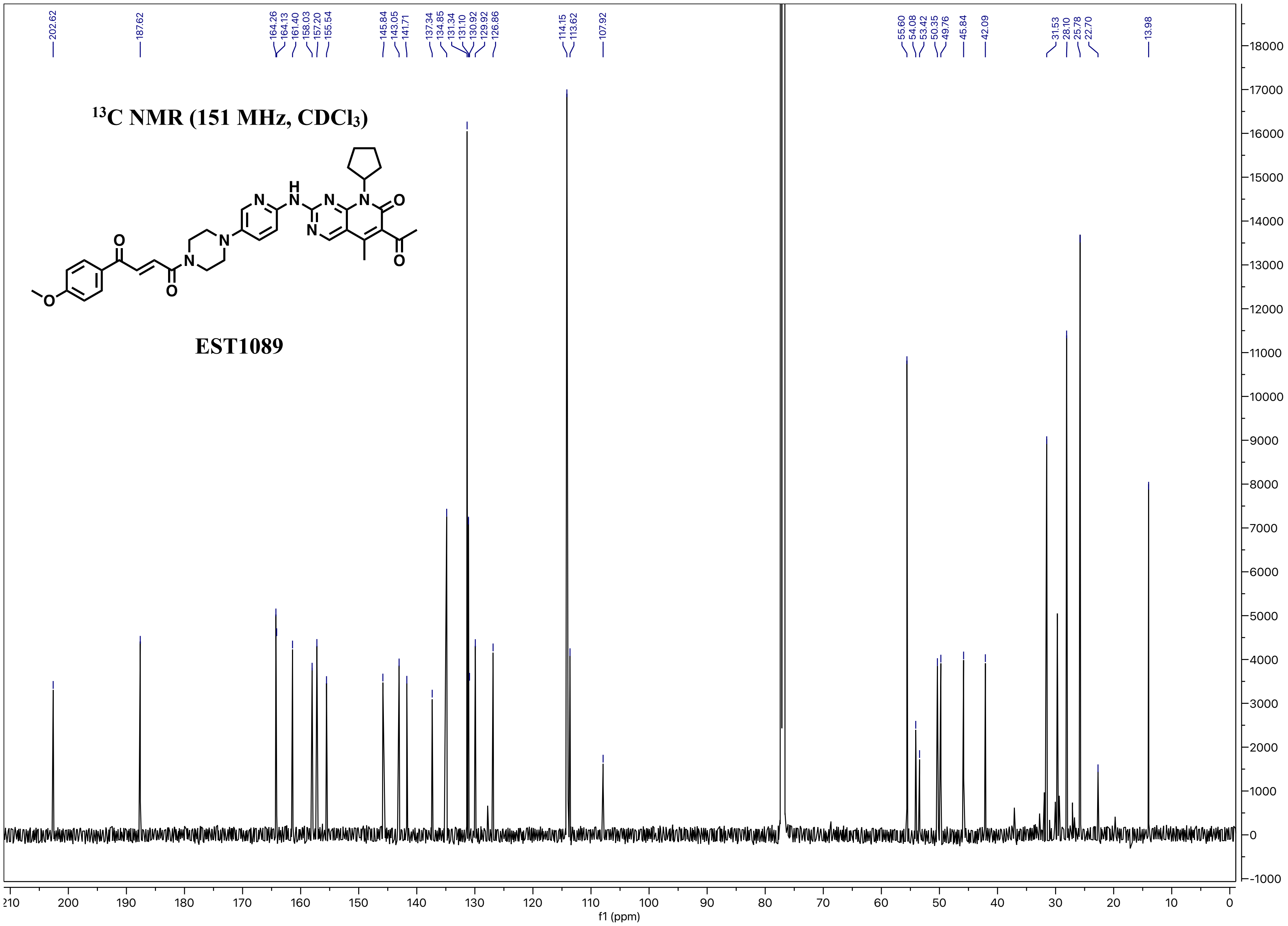

13C NMR (151 MHz, CDCl3)
EST1089
202.62
187.62
164.26
164.13
161.40
158.03
157.20
155.54
145.84
143.05
141.71
137.34
134.85
131.34
131.10
130.92
129.92
126.86
114.15
113.62
107.92
55.60
54.08
53.42
50.35
49.76
45.84
42.09
31.53
28.10
25.78
22.70
13.98
f1 (ppm)

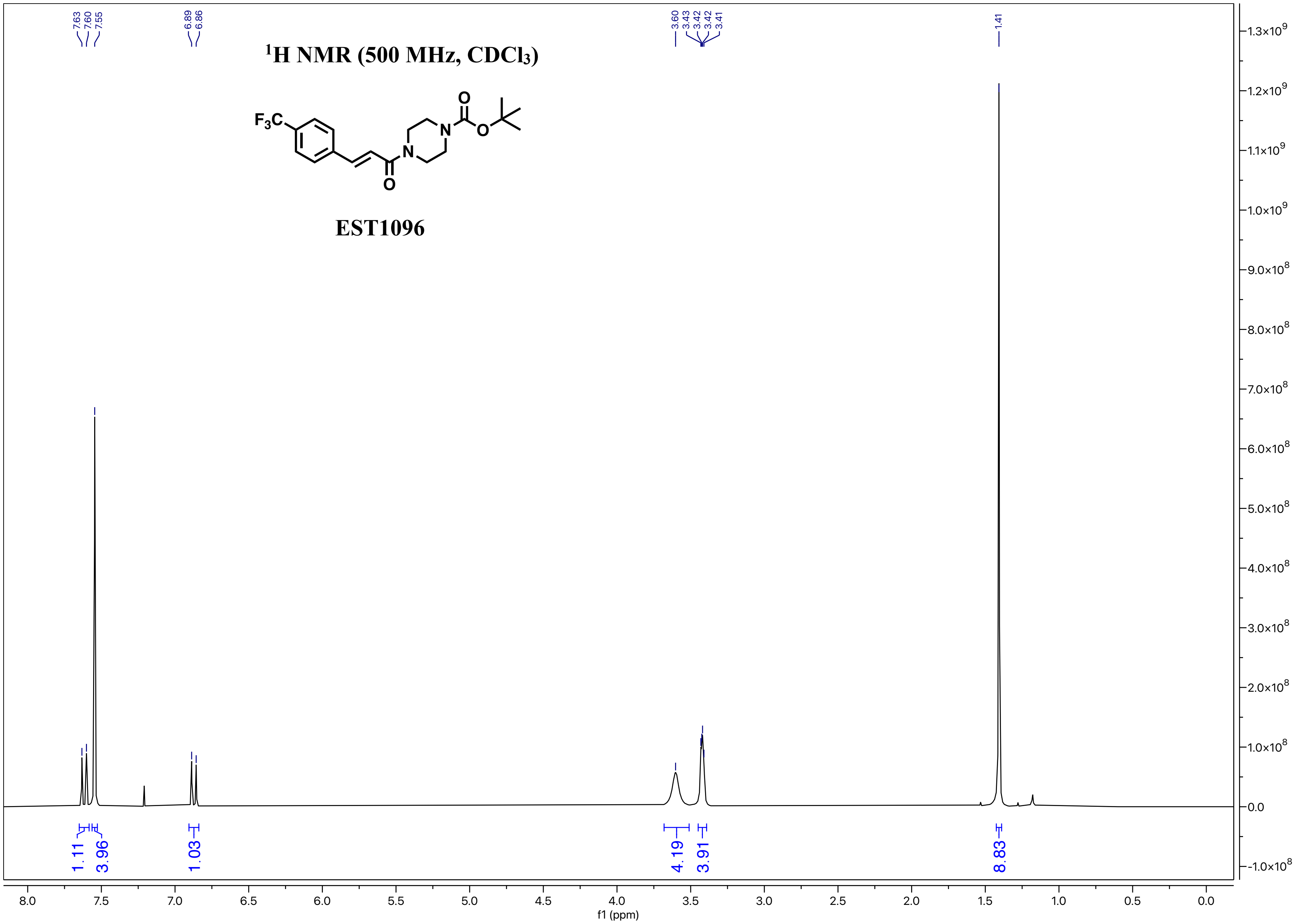

7.63
7.60
7.55
6.89
6.86
3.60
3.43
3.42
3.42
3.41
1.41
$^{1}$H NMR (500 MHz, CDCl$_3$)
F$_3$C
O
N
N
O
O
EST1096
1.11
3.96
1.03
4.19
3.91
8.83
1.3×10$^9$
1.2×10$^9$
1.1×10$^9$
1.0×10$^9$
9.0×10$^8$
8.0×10$^8$
7.0×10$^8$
6.0×10$^8$
5.0×10$^8$
4.0×10$^8$
3.0×10$^8$
2.0×10$^8$
1.0×10$^8$
0.0
−1.0×10$^8$
8.0
7.5
7.0
6.5
6.0
5.5
5.0
4.5
4.0
3.5
3.0
2.5
2.0
1.5
1.0
0.5
0.0
f1 (ppm)

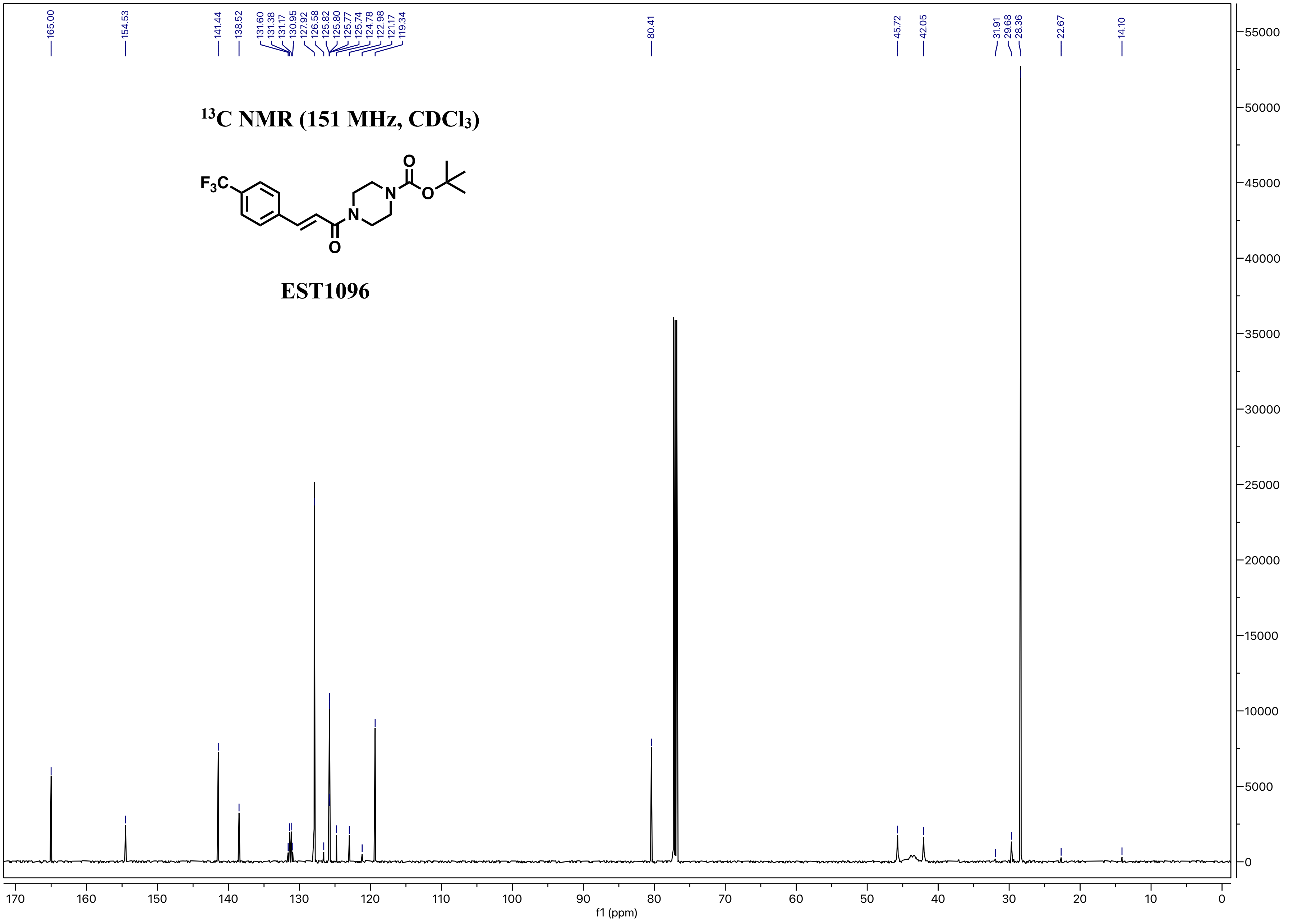

165.00
154.53
141.44
138.52
131.60
131.38
131.17
130.95
127.92
126.58
125.82
125.80
125.77
125.74
124.78
122.98
121.17
119.34
80.41
45.72
42.05
31.91
29.68
28.36
22.67
14.10
$^{13}$C NMR (151 MHz, $CDCl_3$)
$F_3C$
O
N
O
N
O
EST1096
f1 (ppm)

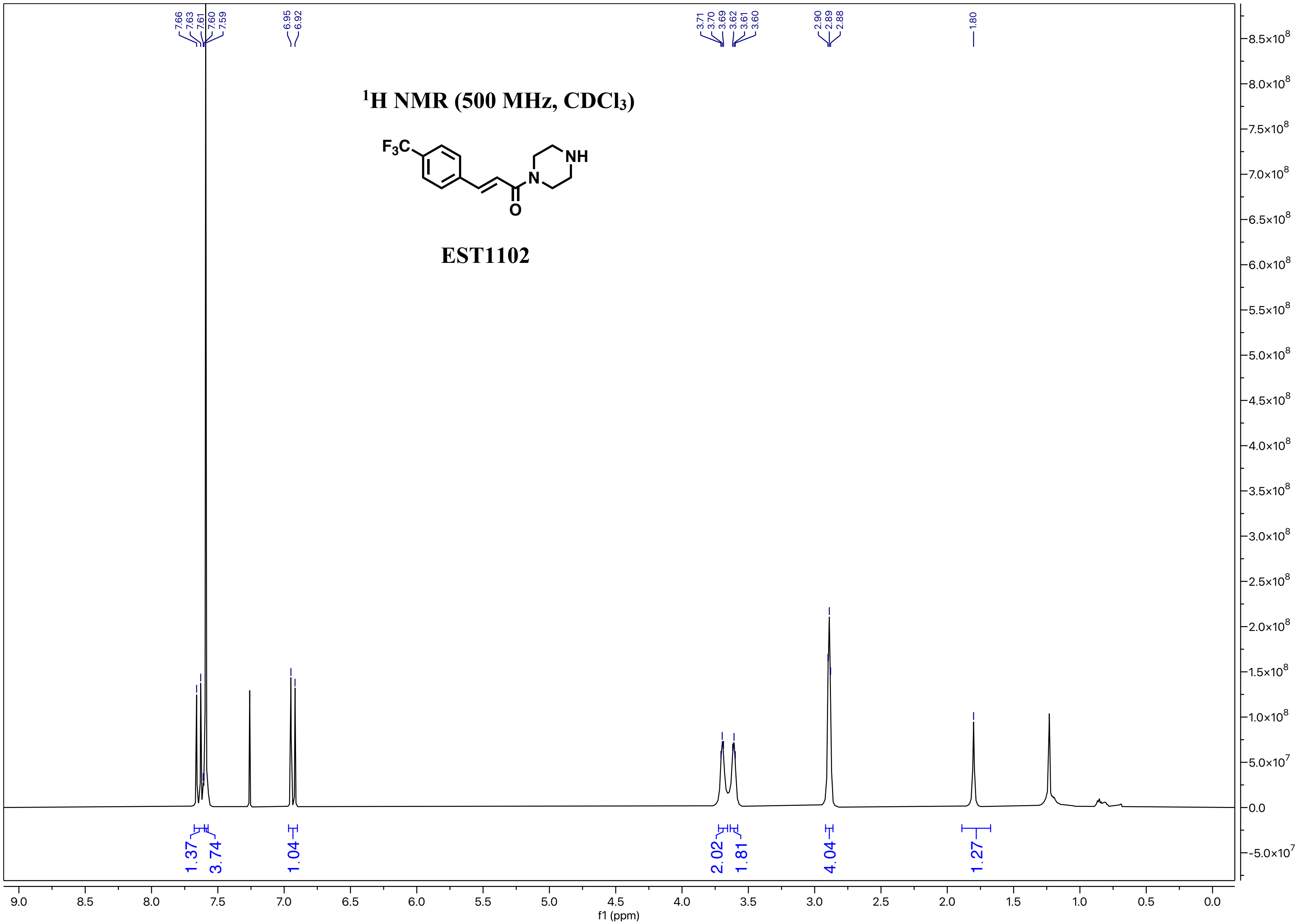

7.66
7.63
7.61
7.60
7.59
6.95
6.92
3.71
3.70
3.69
3.62
3.61
3.60
2.90
2.89
2.88
1.80
$^1$H NMR (500 MHz, CDCl$_3$)
F$_3$C
NH
N
O
EST1102
1.37
3.74
1.04
2.02
1.81
4.04
1.27
f1 (ppm)

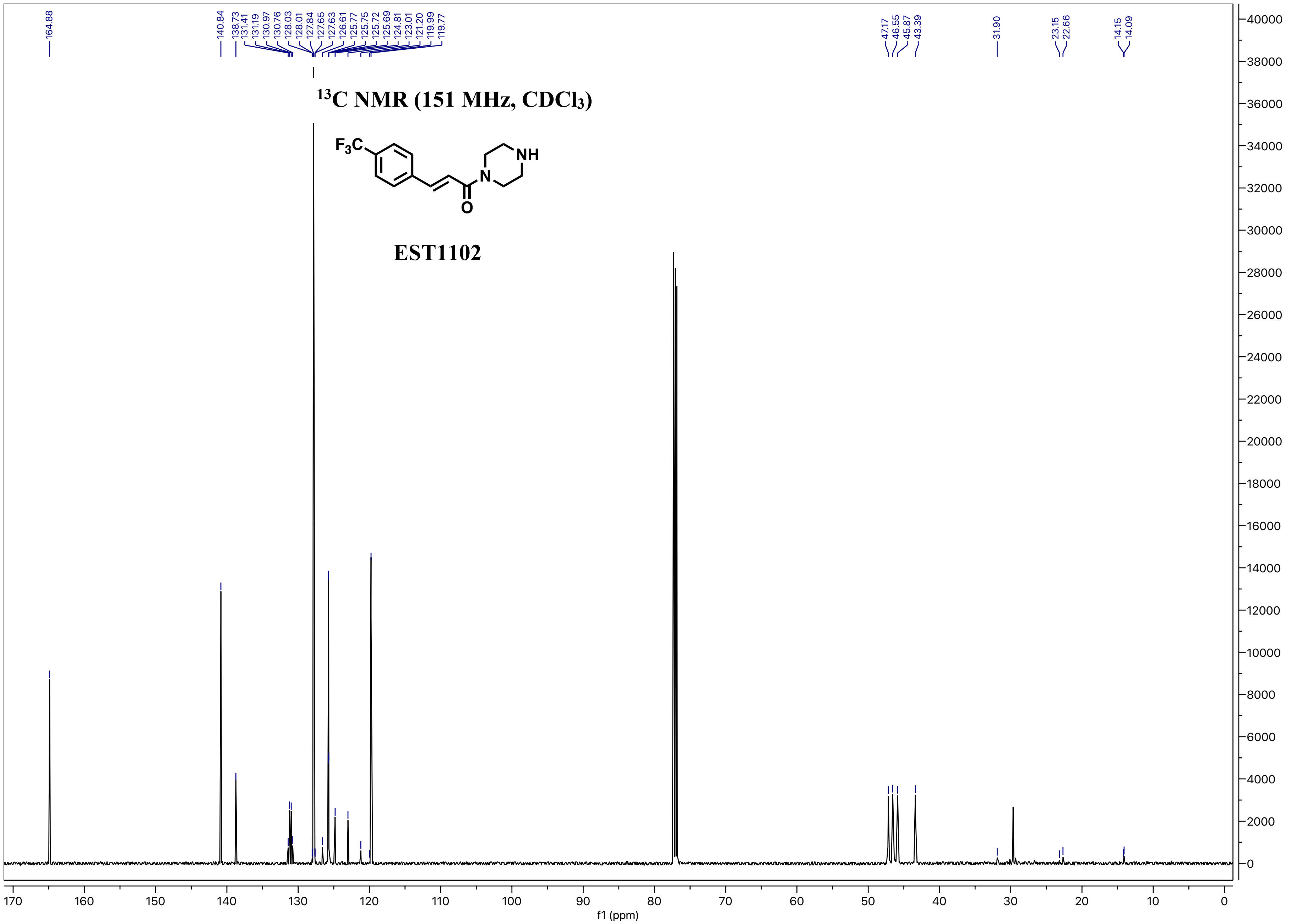

¹³C NMR (151 MHz, CDCl₃)
F₃C
NH
N
O
EST1102
164.88
140.84
138.73
131.41
131.19
130.97
130.76
128.03
128.01
127.84
127.65
127.63
126.61
125.77
125.75
125.72
125.69
124.81
123.01
121.20
119.99
119.77
47.17
46.55
45.87
43.39
31.90
23.15
22.66
14.15
14.09
f1 (ppm)

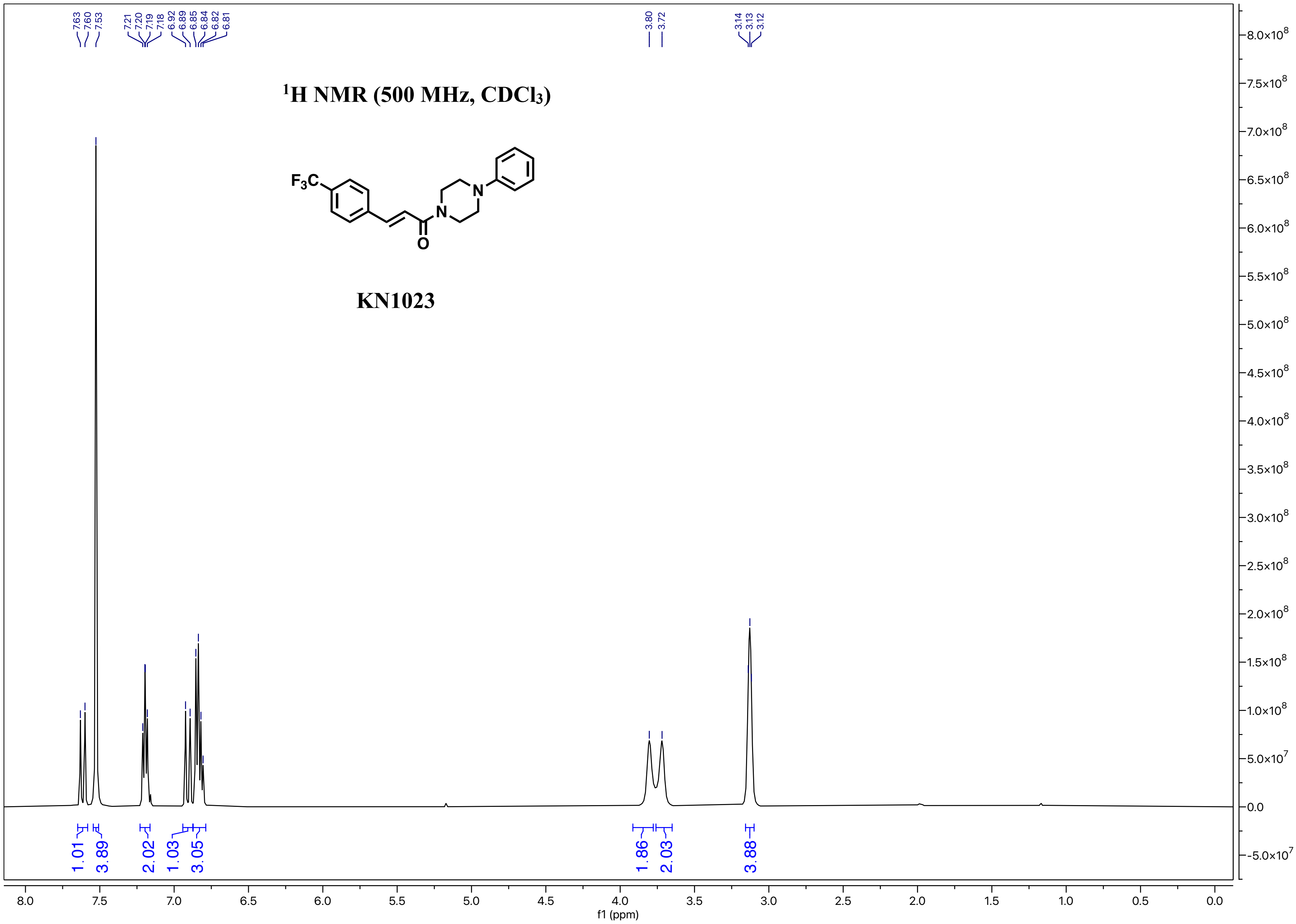

1H NMR (500 MHz, CDCl3)
F3C
N
N
O
KN1023
7.63
7.60
7.53
7.21
7.20
7.19
7.18
6.92
6.89
6.85
6.84
6.82
6.81
3.80
3.72
3.14
3.13
3.12
1.01
3.89
2.02
1.03
3.05
1.86
2.03
3.88
f1 (ppm)

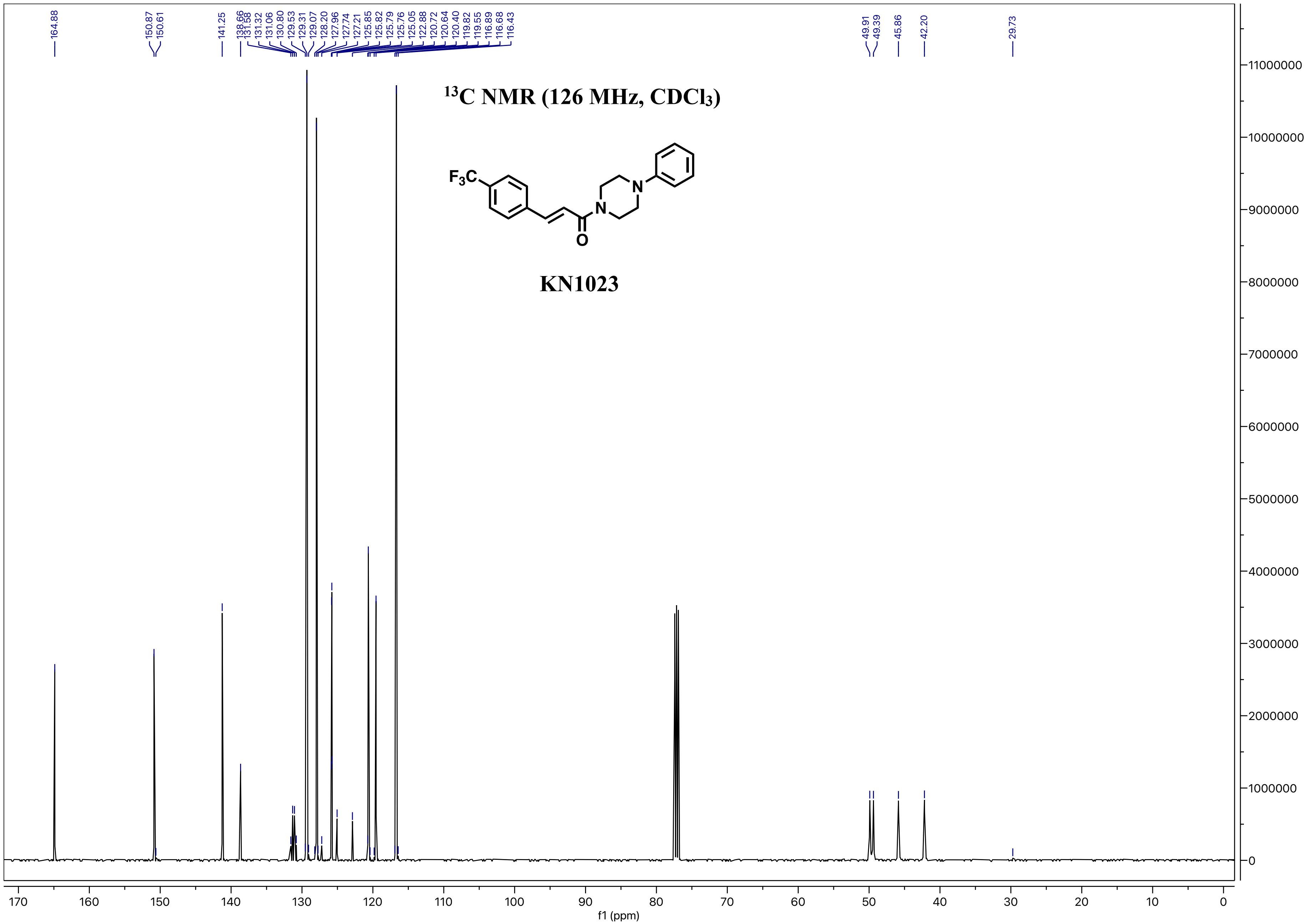

164.88
150.87
150.61
141.25
138.66
131.58
131.32
131.06
130.80
129.53
129.31
129.07
128.20
127.96
127.74
127.21
125.85
125.82
125.79
125.76
125.05
122.88
120.72
120.64
120.40
119.82
119.55
116.89
116.68
116.43
49.91
49.39
45.86
42.20
29.73
$^{13}$C NMR (126 MHz, CDCl$_3$)
$F_3C$
N
N
O
KN1023
f1 (ppm)

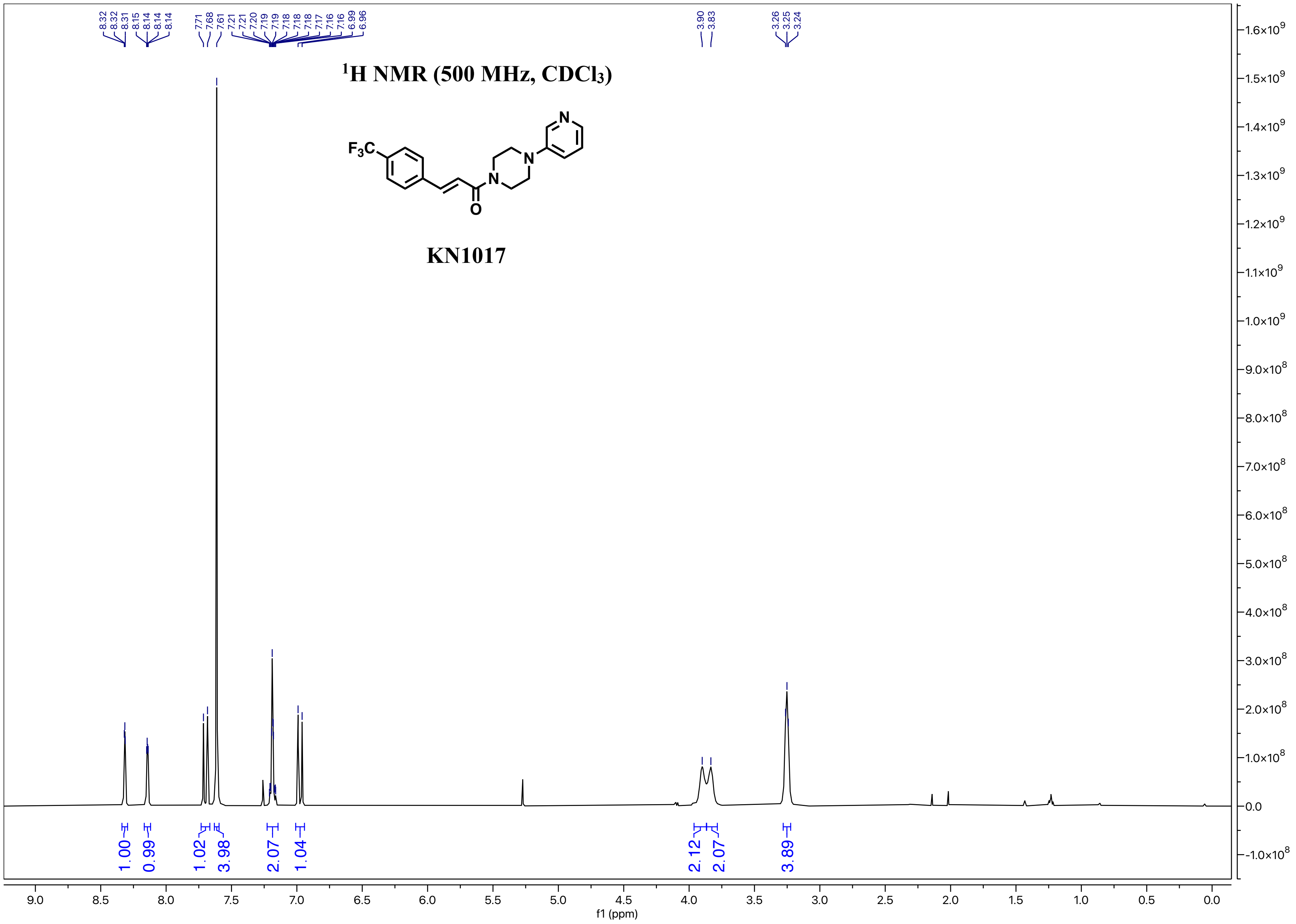

8.32
8.32
8.31
8.15
8.14
8.14
8.14
7.71
7.68
7.61
7.21
7.21
7.20
7.19
7.19
7.18
7.18
7.18
7.17
7.16
7.16
6.99
6.96
3.90
3.83
3.26
3.25
3.24
$^1$H NMR (500 MHz, $CDCl_3$)
$F_3C$
N
N
N
O
KN1017
1.00
0.99
1.02
3.98
2.07
1.04
2.12
2.07
3.89
f1 (ppm)

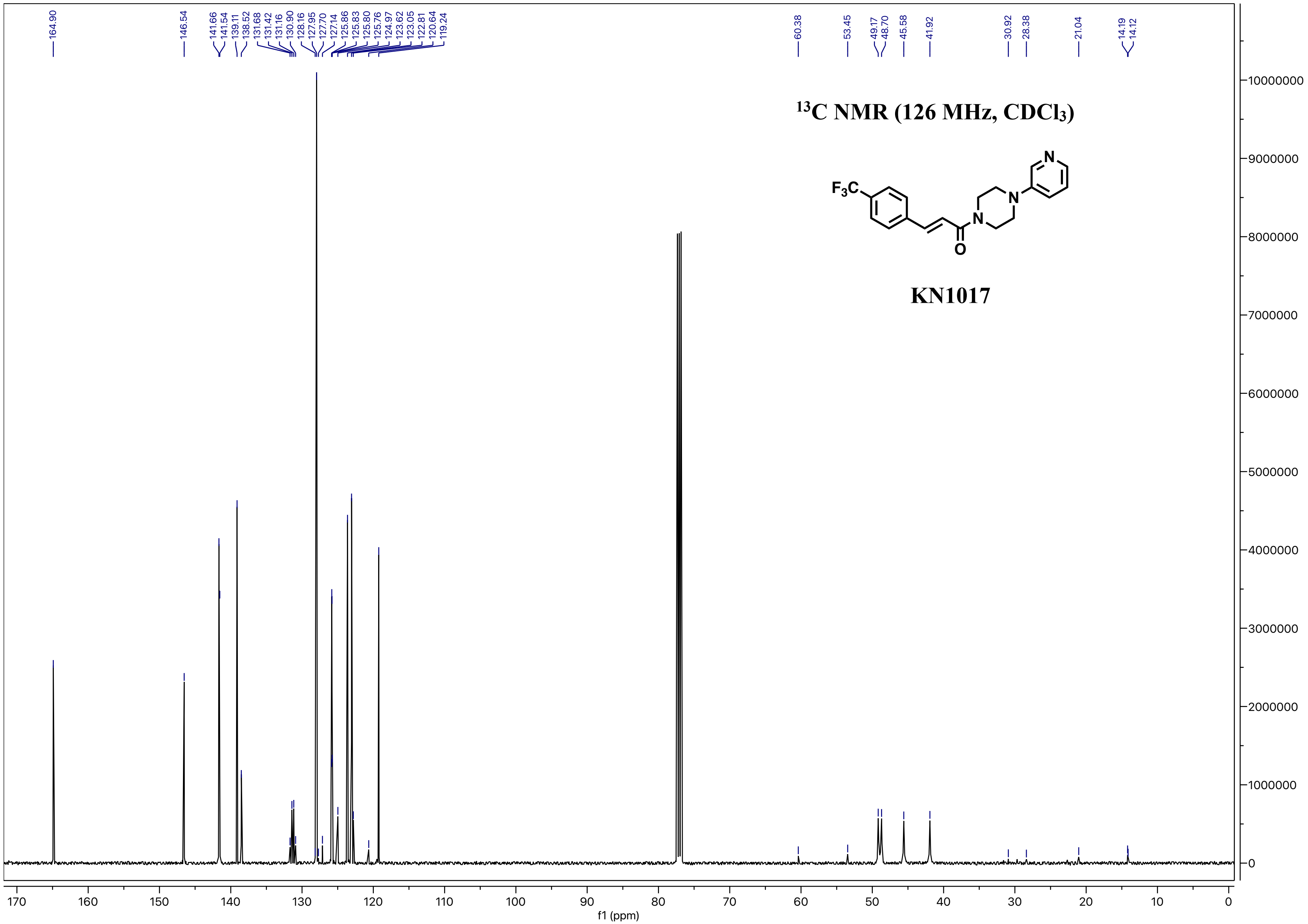

164.90
146.54
141.66
141.54
139.11
138.52
131.68
131.42
131.16
130.90
128.16
127.95
127.70
127.14
125.86
125.83
125.80
125.76
124.97
123.62
123.05
122.81
120.64
119.24
60.38
53.45
49.17
48.70
45.58
41.92
30.92
28.38
21.04
14.19
14.12
$^{13}$C NMR (126 MHz, CDCl$_3$)
KN1017
f1 (ppm)

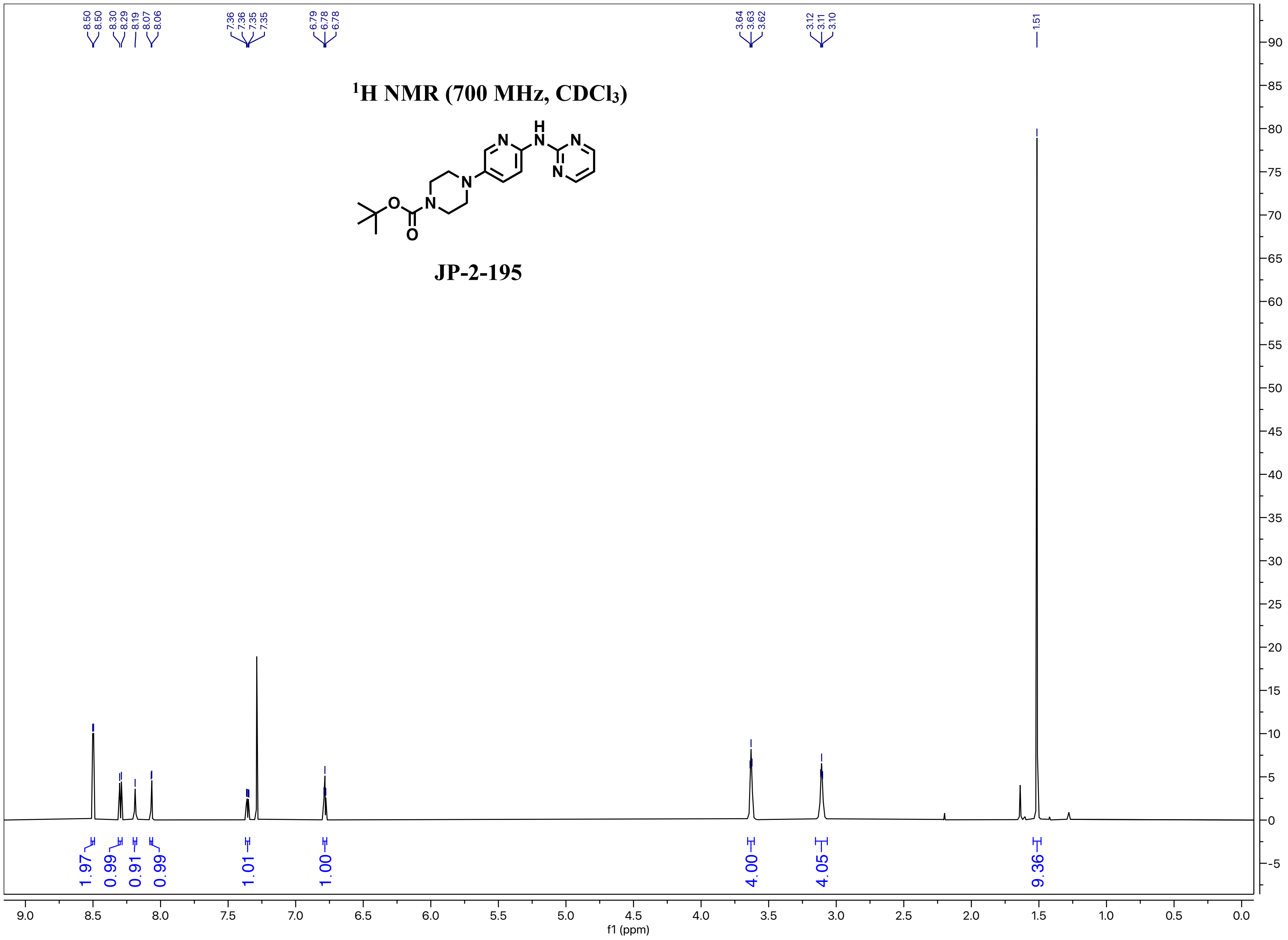

¹H NMR (700 MHz, CDCl₃)
JP-2-195
8.50 8.50 8.30 8.29 8.19 8.07 8.06
7.36 7.36 7.35 7.35
6.79 6.78 6.78
3.64 3.63 3.62
3.12 3.11 3.10
1.51
1.97 0.99 0.91 0.99 1.01 1.00 4.00 4.05 9.36
f1 (ppm)

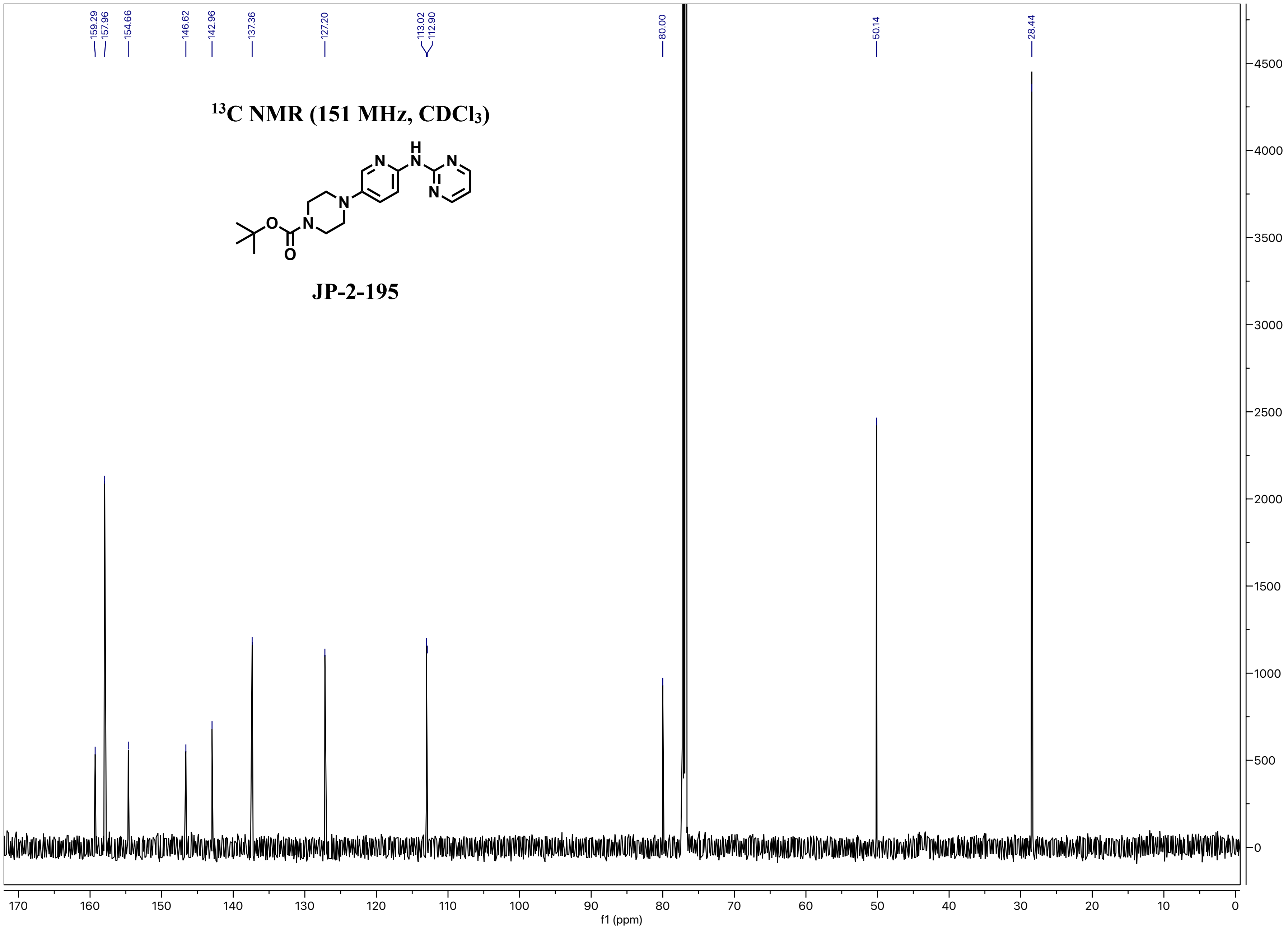
159.29
157.96
154.66
146.62
142.96
137.36
127.20
113.02
112.90
80.00
50.14
28.44
13C NMR (151 MHz, CDCl3)
JP-2-195
H
N
O
f1 (ppm)
170 160 150 140 130 120 110 100 90 80 70 60 50 40 30 20 10 0
4500 4000 3500 3000 2500 2000 1500 1000 500 0

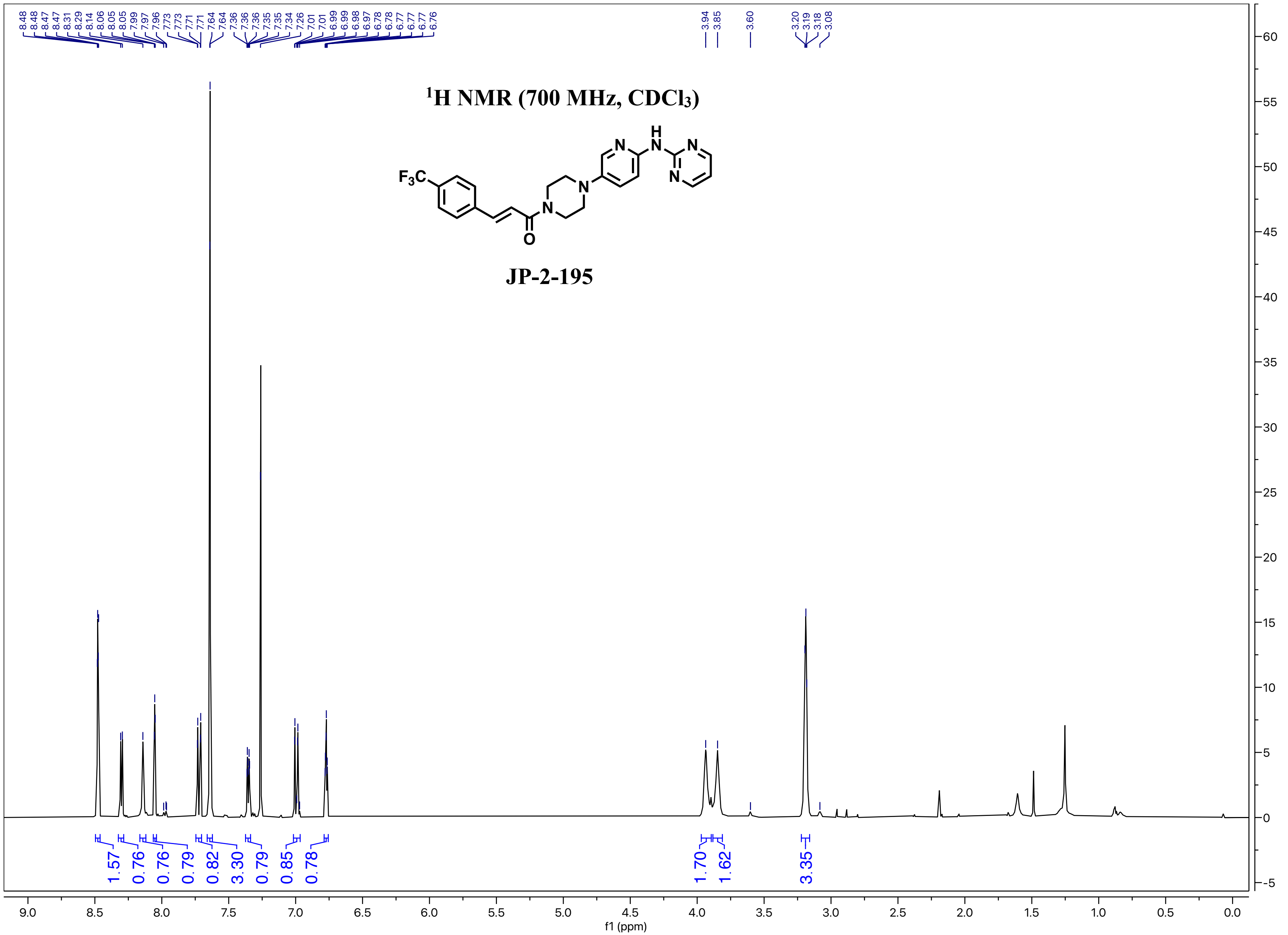

$^{1}$H NMR (700 MHz, CDCl$_3$)
JP-2-195
F3C
N
H
8.48 8.48 8.47 8.47 8.31 8.29 8.14 8.06 8.05 8.05 7.99 7.97 7.96 7.73 7.73 7.71 7.71 7.64 7.64 7.36 7.36 7.36 7.35 7.35 7.34 7.26 7.01 7.01 6.99 6.99 6.98 6.97 6.78 6.78 6.77 6.77 6.77 6.76
3.94 3.85 3.60 3.20 3.19 3.18 3.08
1.57 0.76 0.76 0.79 0.82 3.30 0.79 0.85 0.78
1.70 1.62 3.35
f1 (ppm)

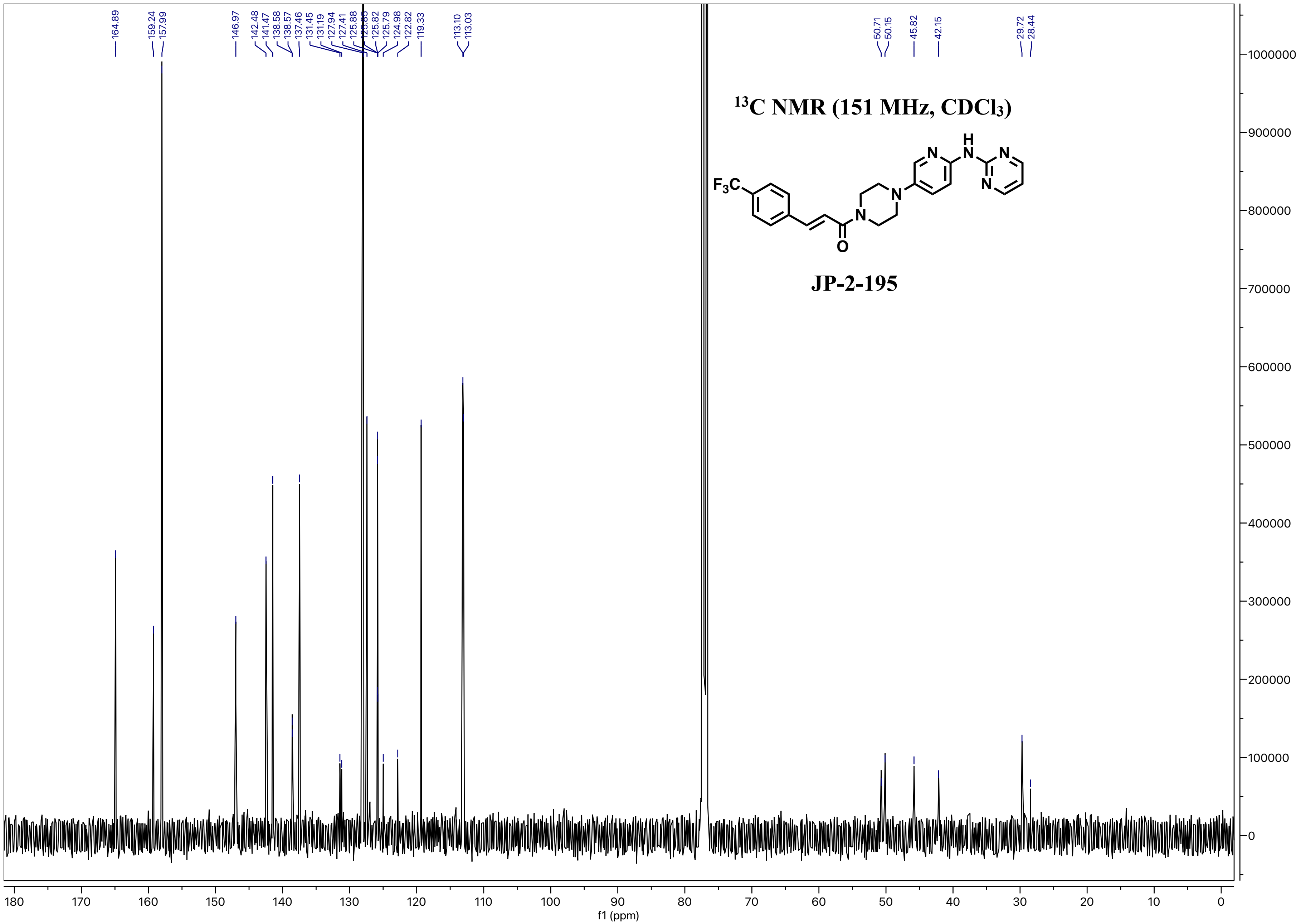

164.89
159.24
157.99
146.97
142.48
141.47
138.58
138.57
137.46
131.45
131.19
127.94
127.41
125.88
125.85
125.82
125.79
124.98
122.82
119.33
113.10
113.03
50.71
50.15
45.82
42.15
29.72
28.44
13C NMR (151 MHz, CDCl3)
F3C
N
H
O
JP-2-195
f1 (ppm)

$^1$H NMR (500 MHz, $CDCl_3$)

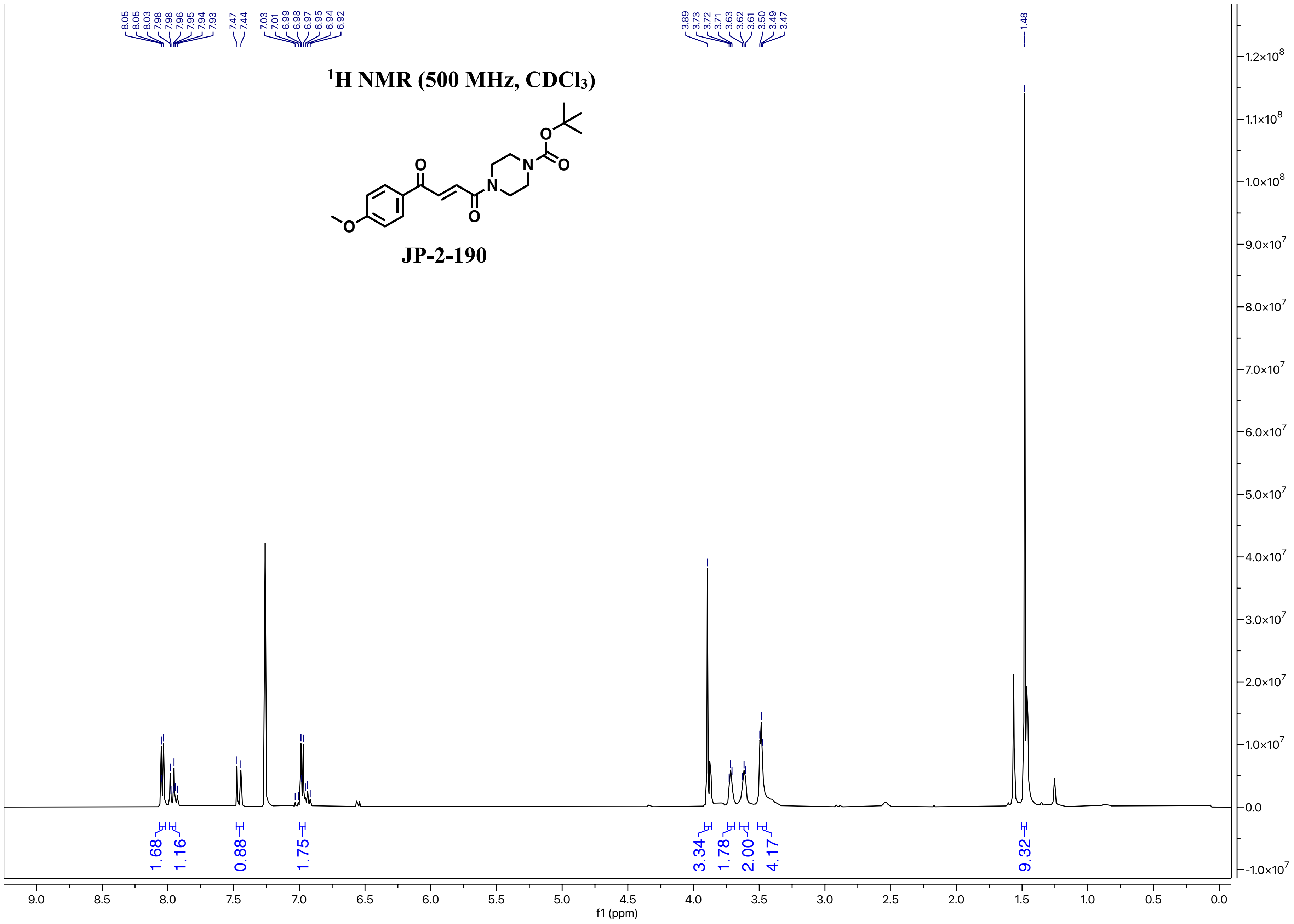


**JP-2-190**

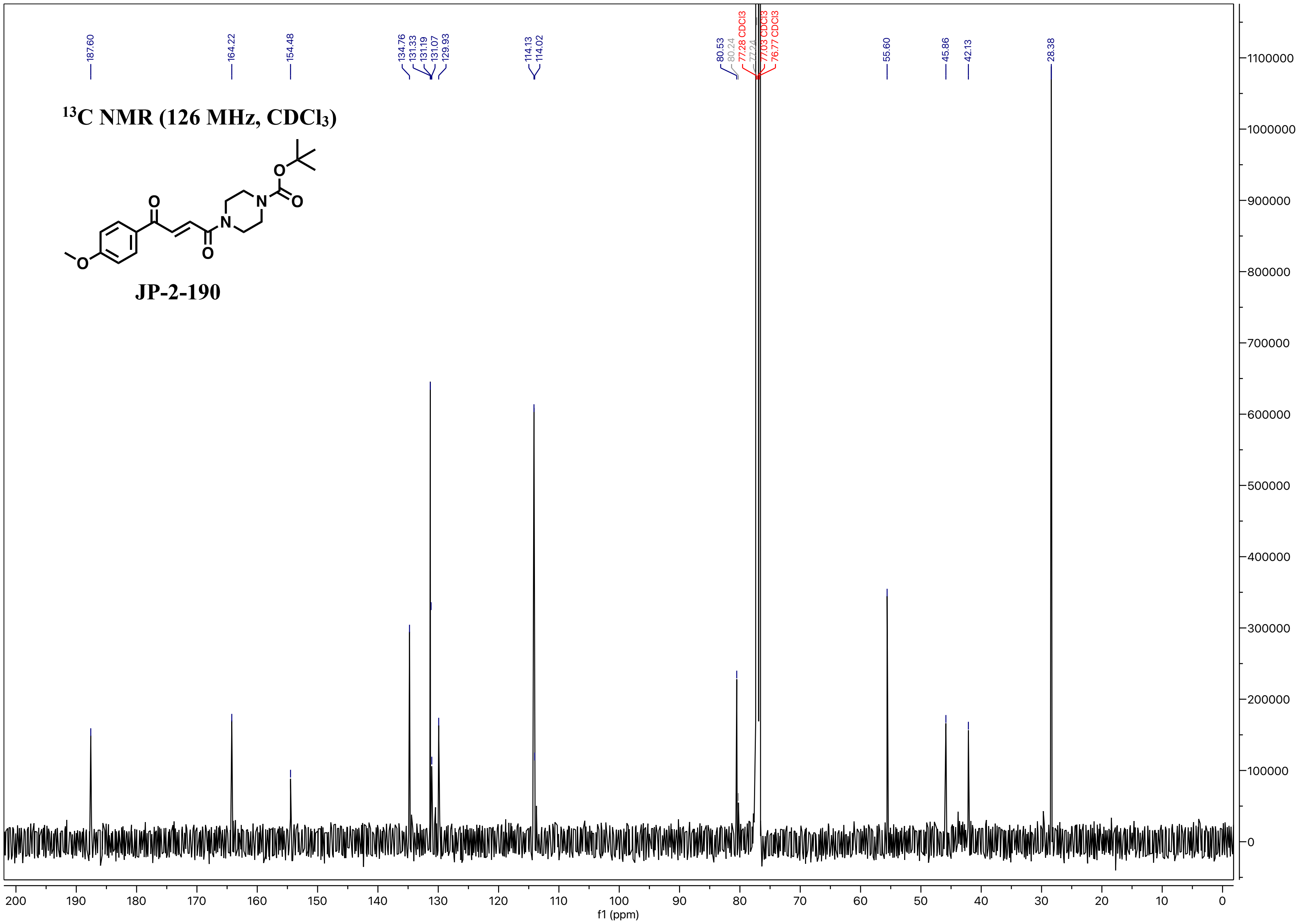

13C NMR (126 MHz, CDCl3)
JP-2-190
187.60
164.22
154.48
134.76
131.33
131.19
131.07
129.93
114.13
114.02
80.53
80.24
77.28 CDCl3
77.24
77.03 CDCl3
76.77 CDCl3
55.60
45.86
42.13
28.38
f1 (ppm)

$^{1}H$ NMR (700 MHz, $CDCl_3$)

JP-2-196

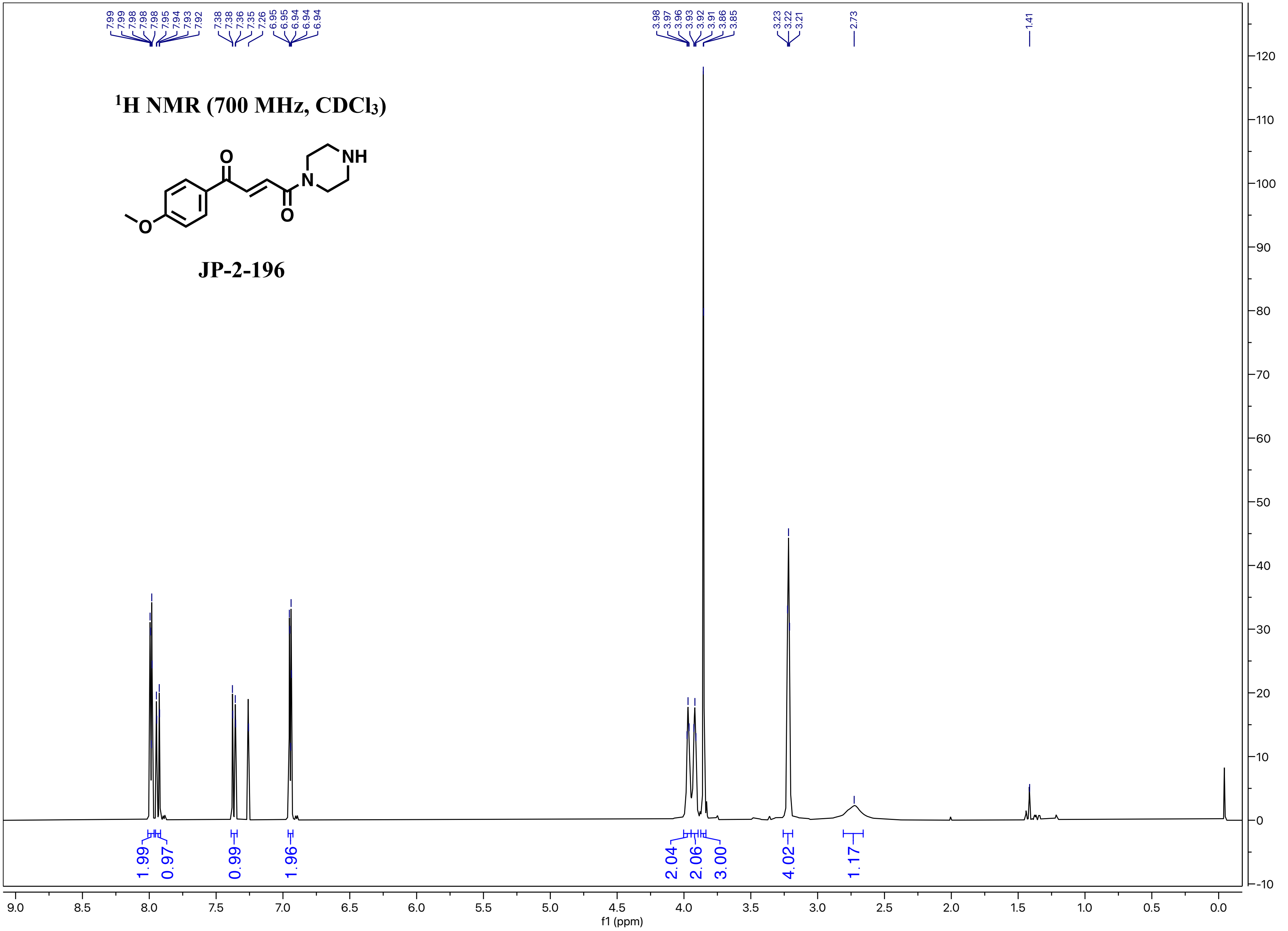

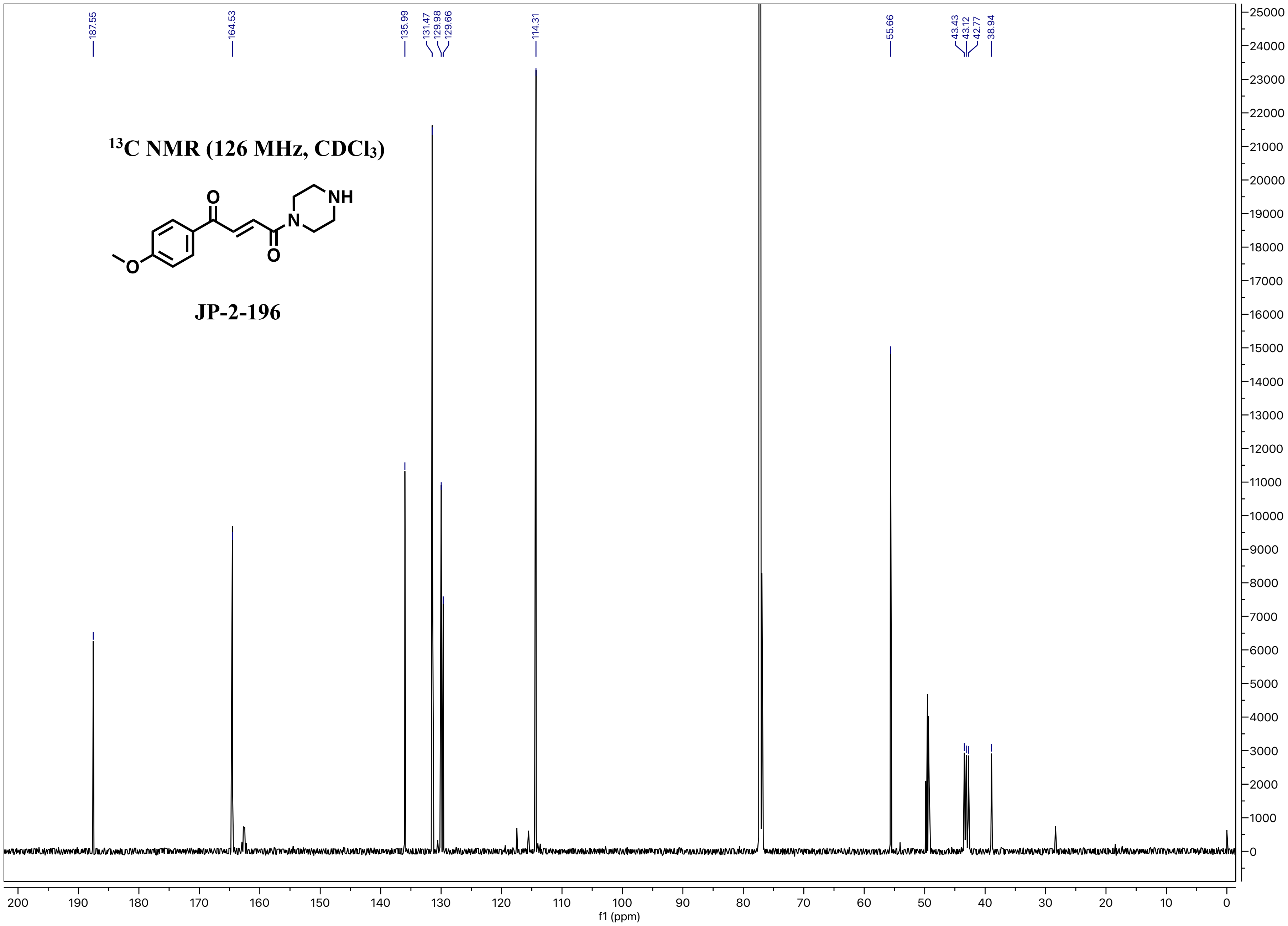

13C NMR (126 MHz, CDCl3)
JP-2-196
187.55
164.53
135.99
131.47
129.98
129.66
114.31
55.66
43.43
43.12
42.77
38.94
f1 (ppm)

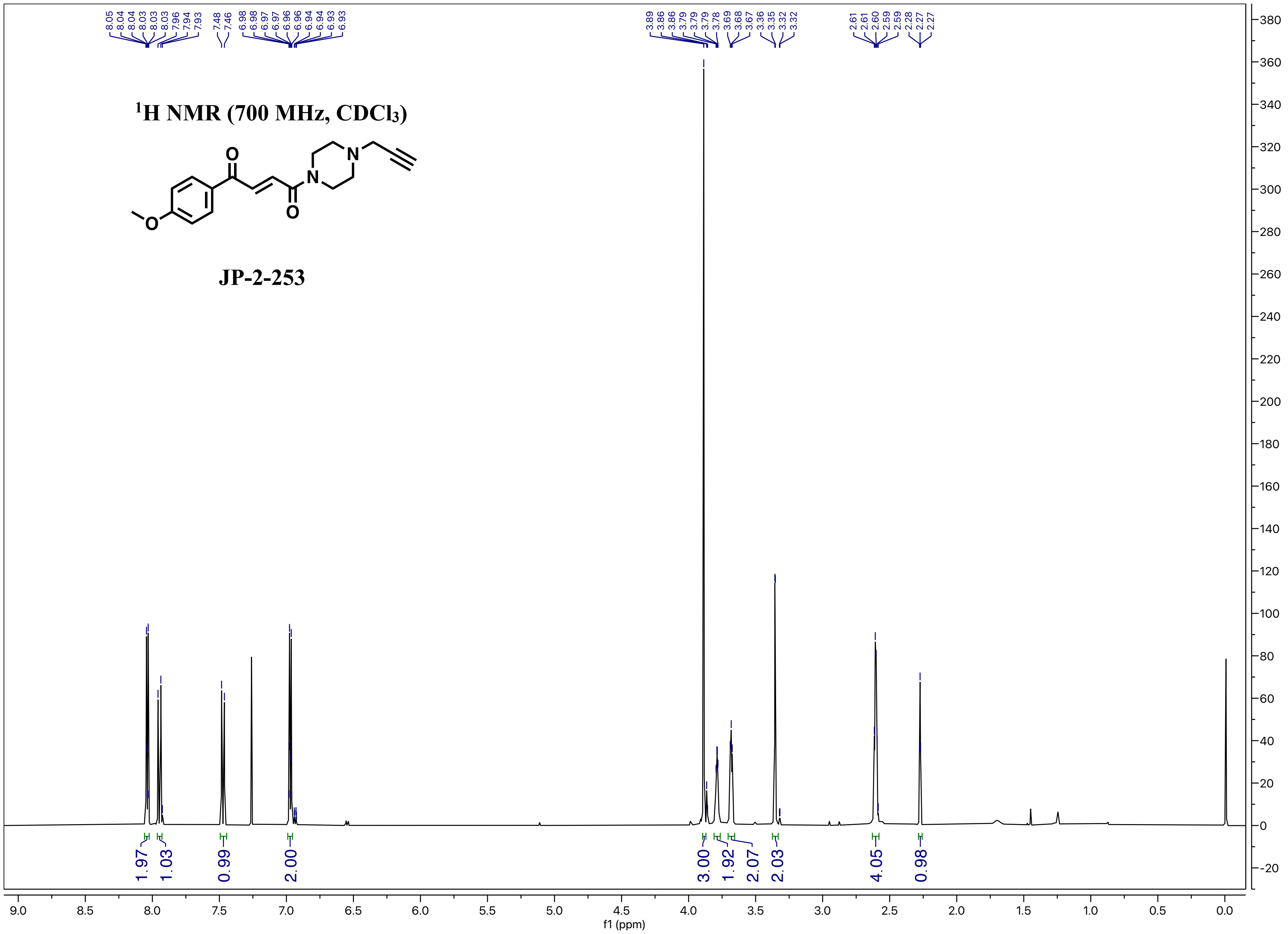

1H NMR (700 MHz, CDCl3)
JP-2-253
8.05 8.04 8.04 8.03 8.03 8.03 7.96 7.94 7.93 7.48 7.46 6.98 6.98 6.97 6.96 6.96 6.94 6.94 6.93 6.93
3.89 3.86 3.86 3.79 3.79 3.79 3.78 3.69 3.68 3.67 3.36 3.35 3.32 3.32
2.61 2.61 2.60 2.59 2.28 2.27 2.27
1.97 1.03 0.99 2.00 3.00 1.92 2.07 2.03 4.05 0.98
f1 (ppm)

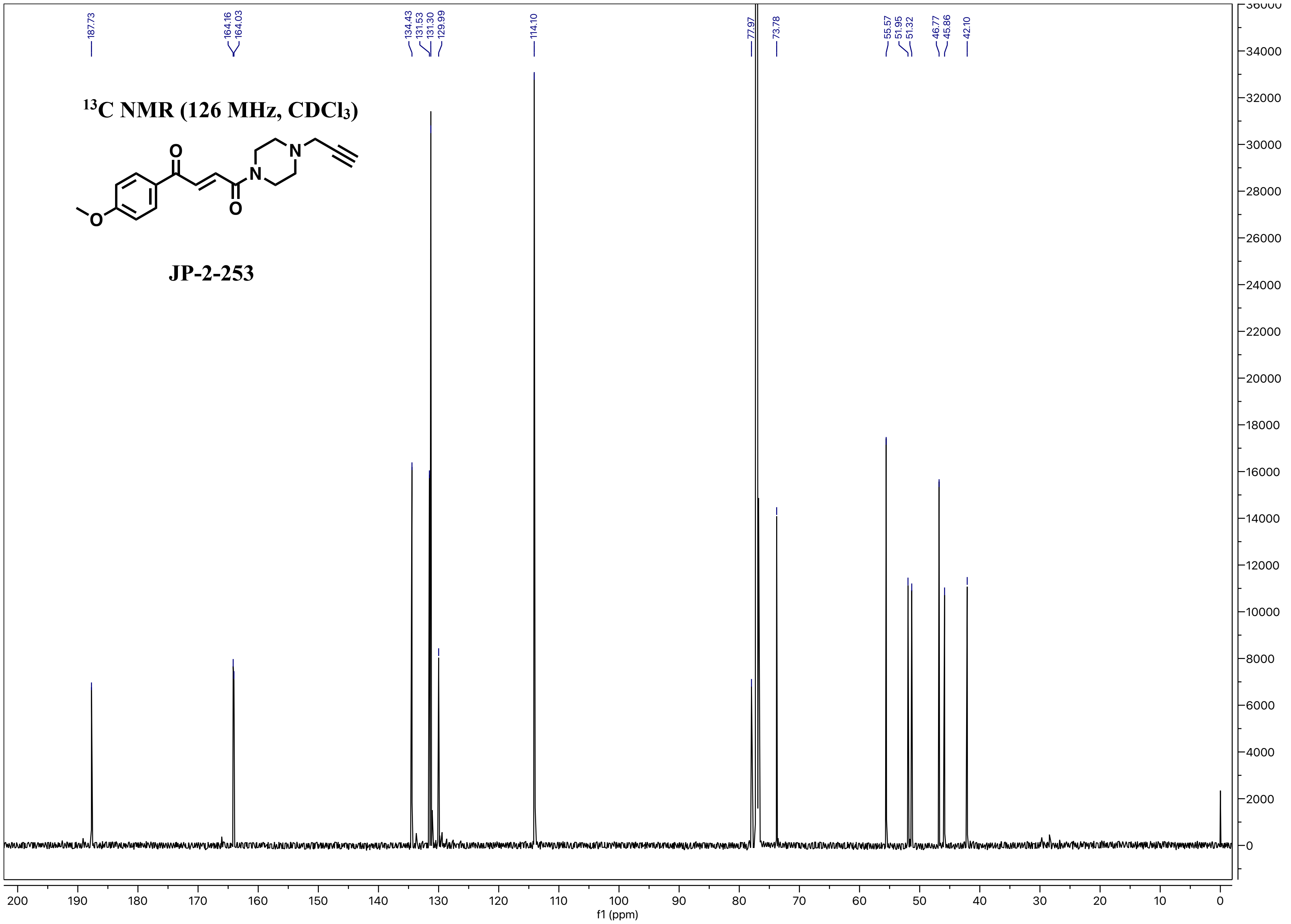

$^{13}$C NMR (126 MHz, CDCl$_3$)
JP-2-253
187.73
164.16
164.03
134.43
131.53
131.30
129.99
114.10
77.97
73.78
55.57
51.95
51.32
46.77
45.86
42.10
f1 (ppm)

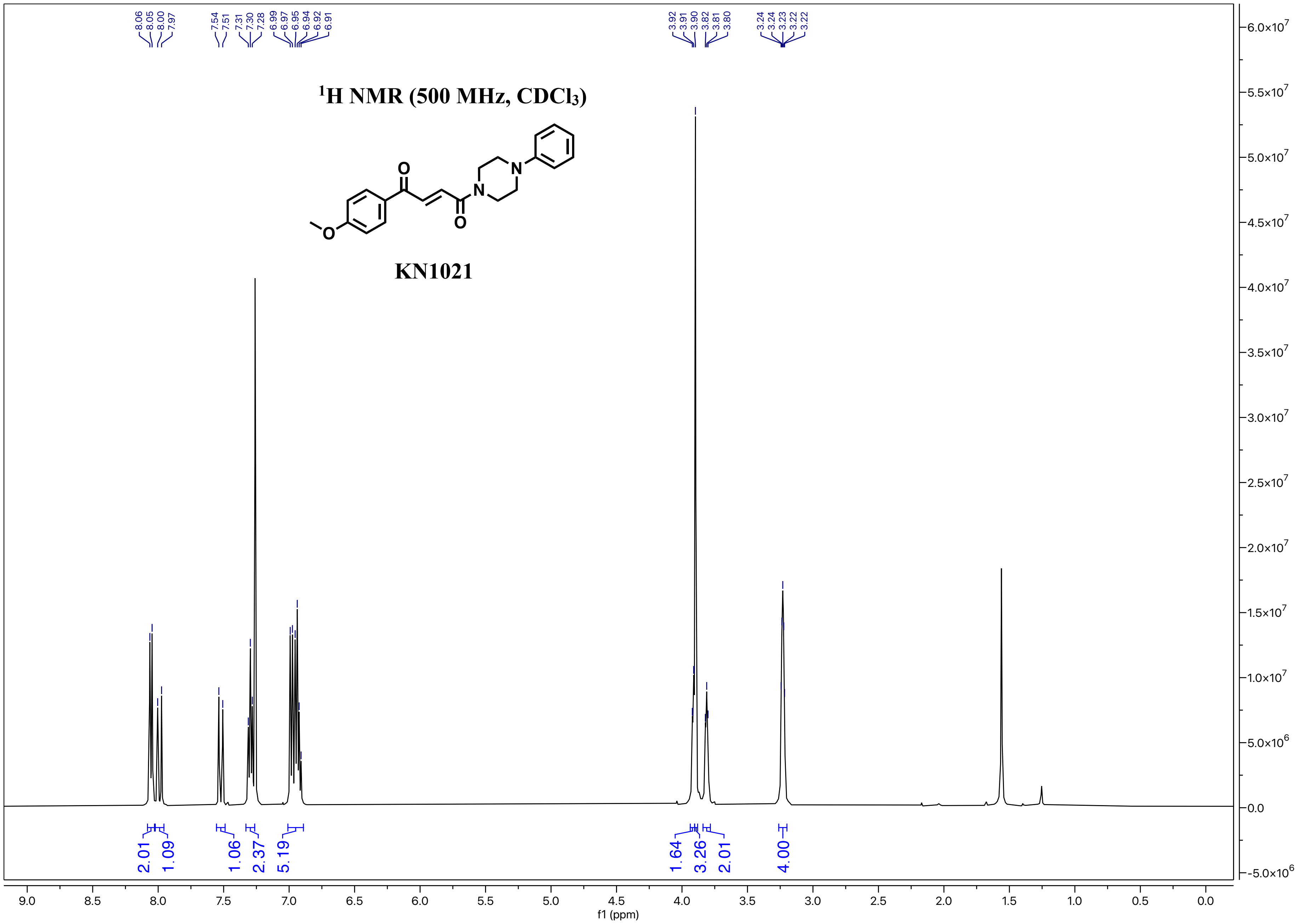

8.06 8.05 8.00 7.97 7.54 7.51 7.31 7.30 7.28 6.99 6.97 6.95 6.94 6.92 6.91
3.92 3.91 3.90 3.82 3.81 3.80 3.24 3.24 3.23 3.22 3.22
¹H NMR (500 MHz, CDCl₃)
O
N
N
O
O
KN1021
2.01 1.09 1.06 2.37 5.19 1.64 3.26 2.01 4.00
f1 (ppm)

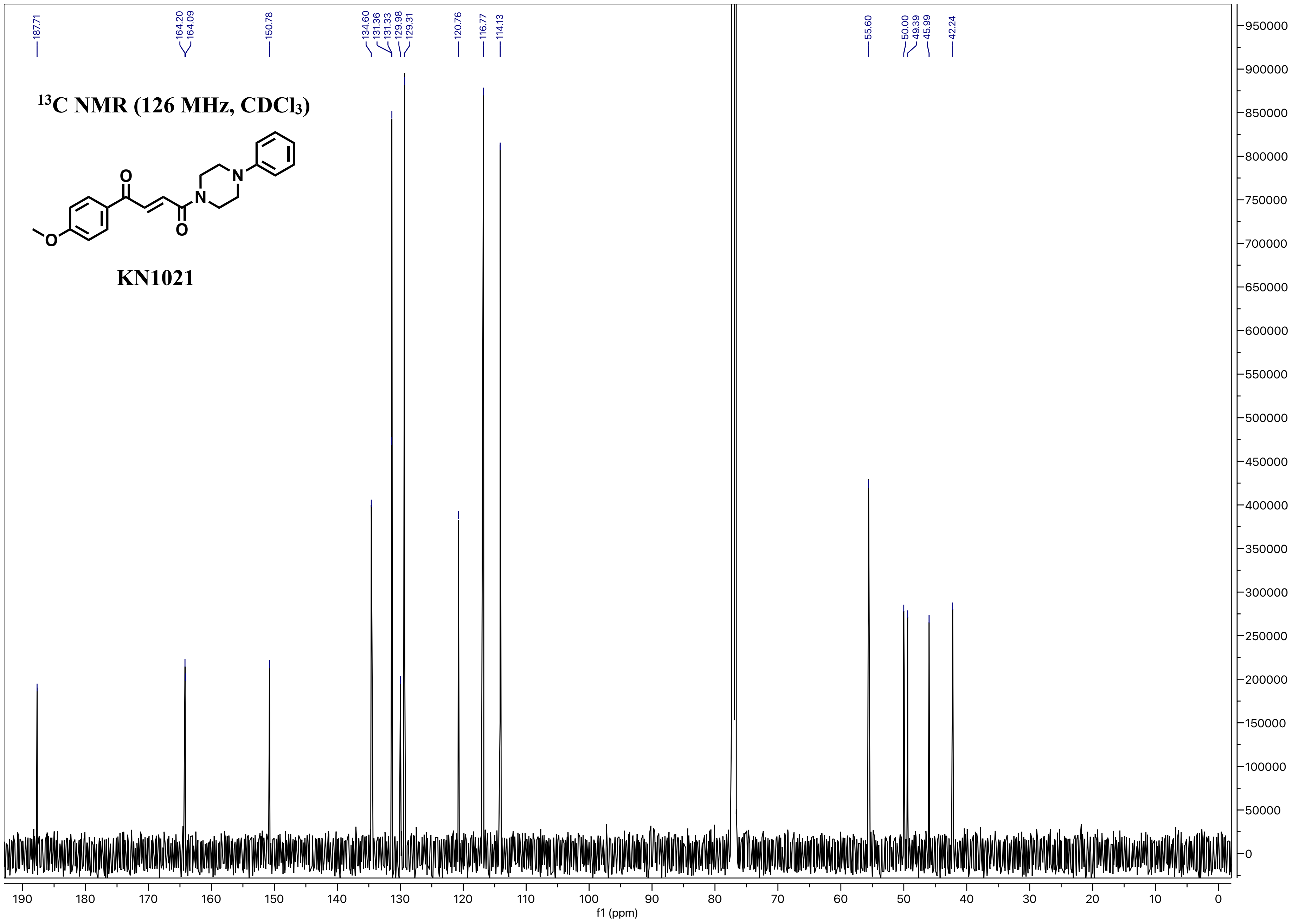

13C NMR (126 MHz, CDCl3)
KN1021
187.71
164.20
164.09
150.78
134.60
131.36
131.33
129.98
129.31
120.76
116.77
114.13
55.60
50.00
49.39
45.99
42.24
f1 (ppm)

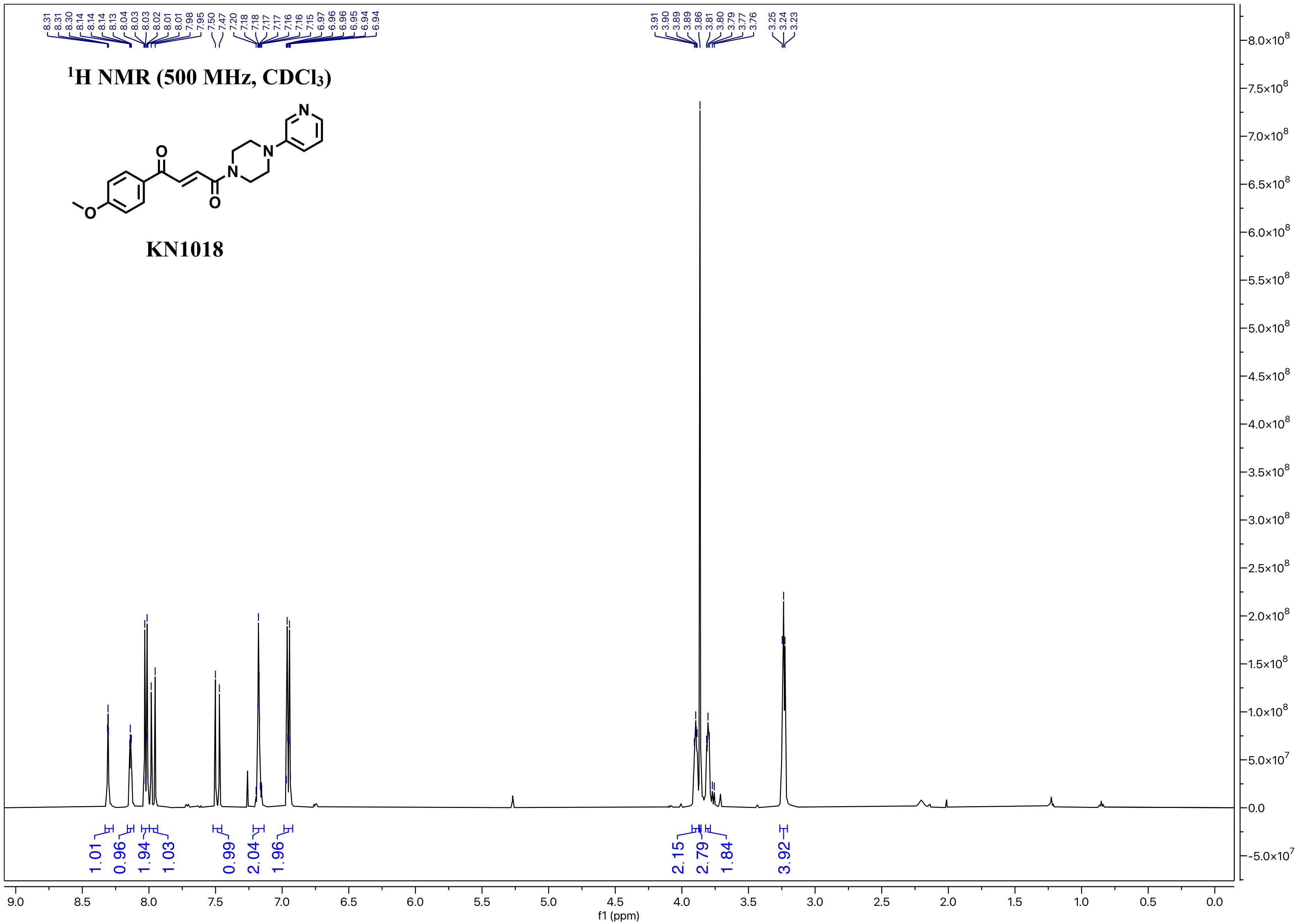

1H NMR (500 MHz, CDCl3)
KN1018
8.31 8.31 8.30 8.14 8.14 8.14 8.13 8.04 8.03 8.03 8.02 8.01 8.01 7.98 7.95 7.50 7.47 7.20 7.18 7.18 7.17 7.17 7.16 7.16 7.15 6.97 6.96 6.96 6.95 6.94 6.94
3.91 3.90 3.89 3.89 3.86 3.81 3.80 3.79 3.77 3.76 3.25 3.24 3.23
1.01 0.96 1.94 1.03 0.99 2.04 1.96 2.15 2.79 1.84 3.92
f1 (ppm)

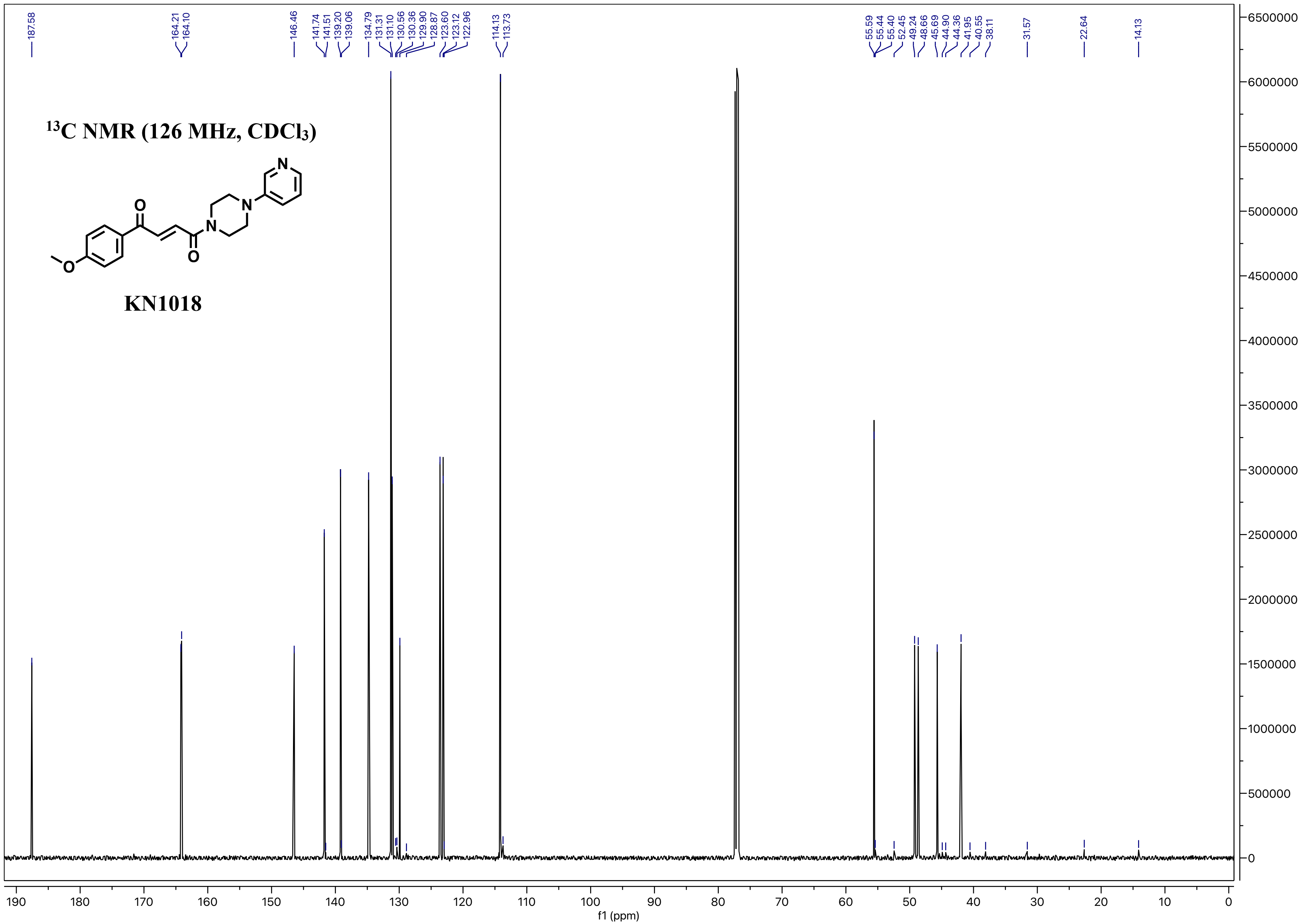

13C NMR (126 MHz, CDCl3)
187.58
164.21
164.10
146.46
141.74
141.51
139.20
139.06
134.79
131.31
131.10
130.56
130.36
129.90
128.87
123.60
123.12
122.96
114.13
113.73
55.59
55.44
55.40
52.45
49.24
48.66
45.69
44.90
44.36
41.95
40.55
38.11
31.57
22.64
14.13
f1 (ppm)


KN1018

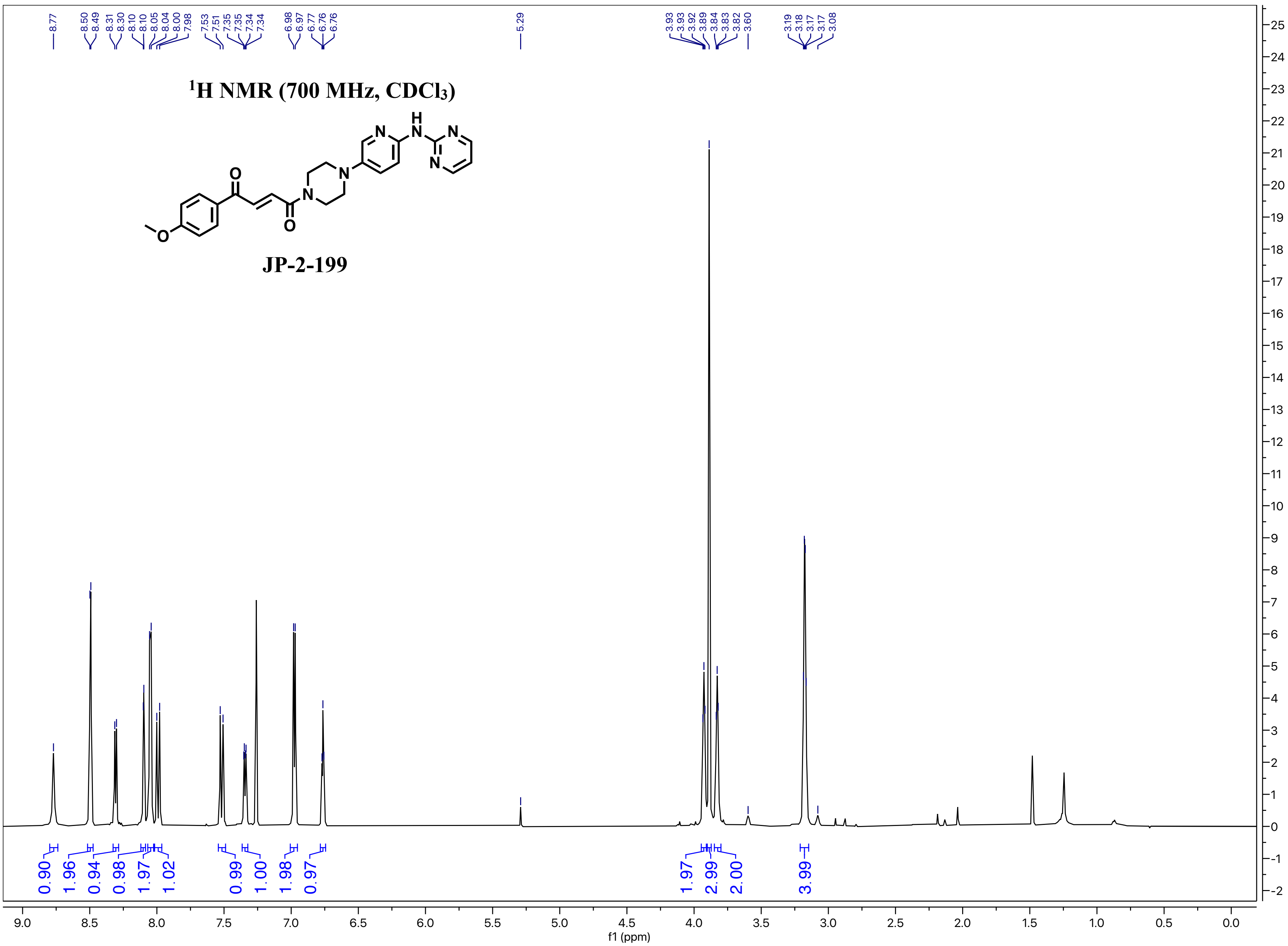

8.77
8.50
8.49
8.31
8.30
8.10
8.10
8.05
8.04
8.00
7.98
7.53
7.51
7.35
7.35
7.34
7.34
6.98
6.97
6.77
6.76
6.76
5.29
3.93
3.93
3.92
3.89
3.84
3.83
3.82
3.60
3.19
3.18
3.17
3.17
3.08
¹H NMR (700 MHz, CDCl₃)
JP-2-199
0.90
1.96
0.94
0.98
1.97
1.02
0.99
1.00
1.98
0.97
1.97
2.99
2.00
3.99
f1 (ppm)

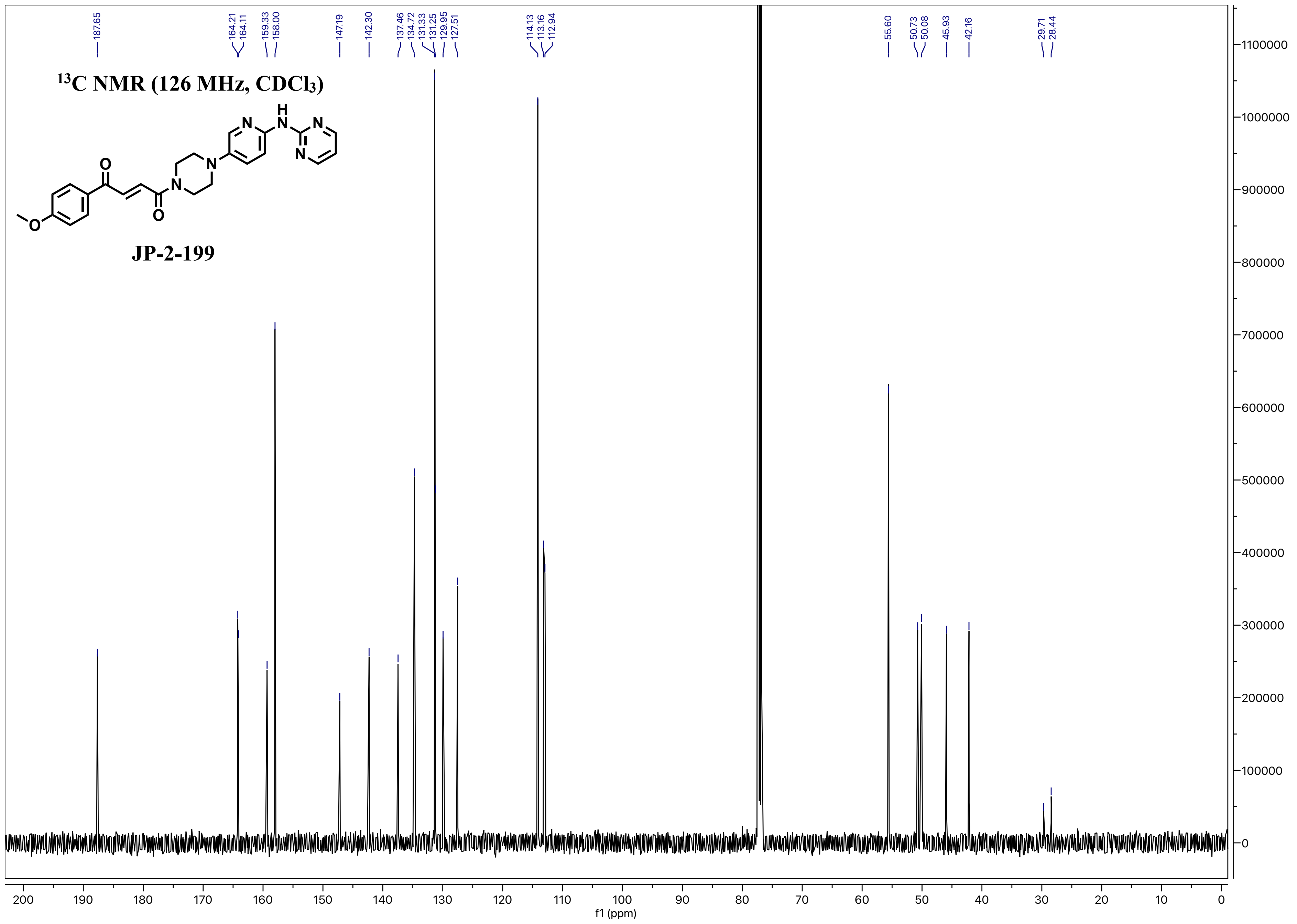
13C NMR (126 MHz, CDCl3)
JP-2-199
187.65
164.21
164.11
159.33
158.00
147.19
142.30
137.46
134.72
131.33
131.25
129.95
127.51
114.13
113.16
112.94
55.60
50.73
50.08
45.93
42.16
29.71
28.44
f1 (ppm)

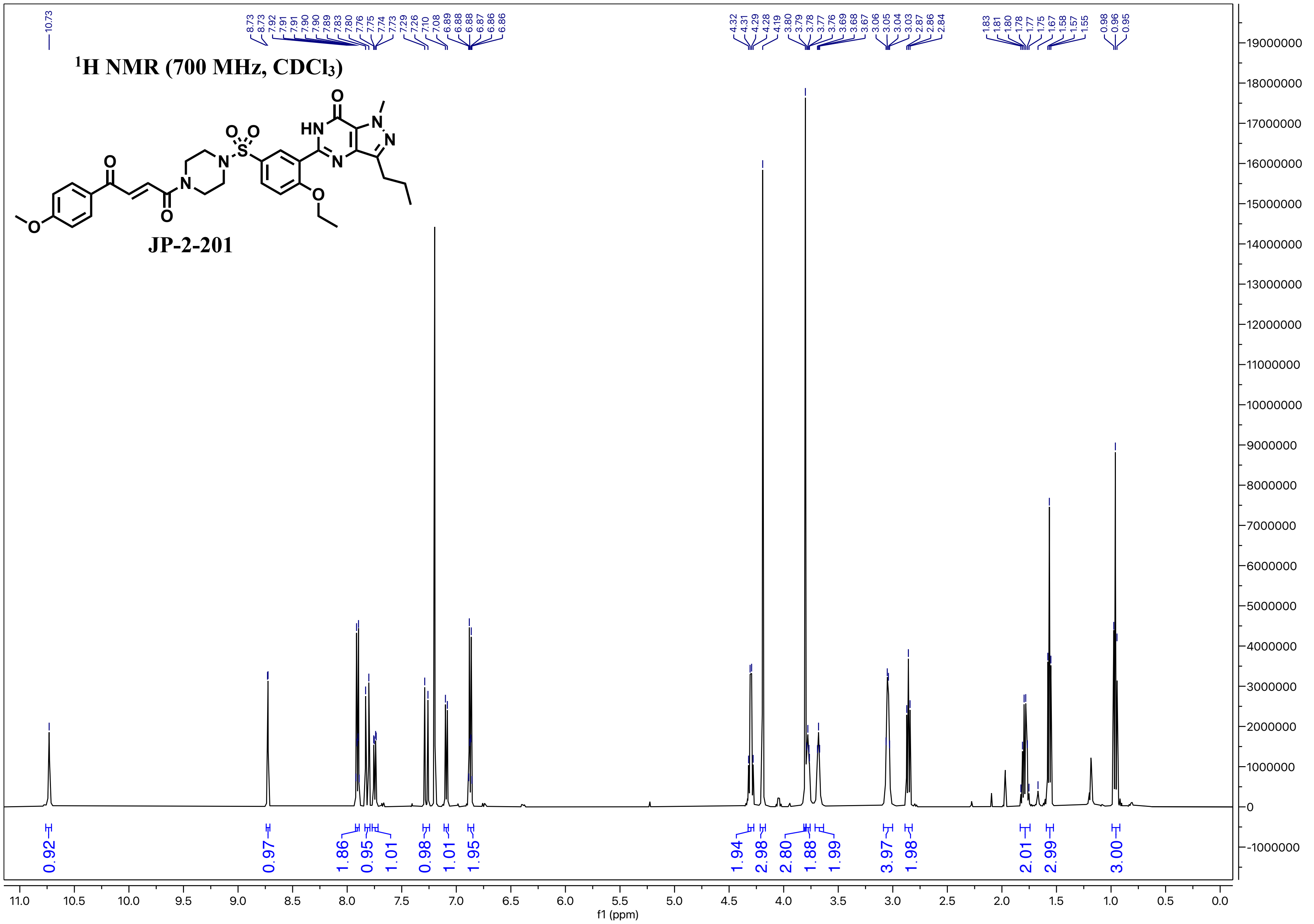

1H NMR (700 MHz, CDCl3)
JP-2-201
f1 (ppm)

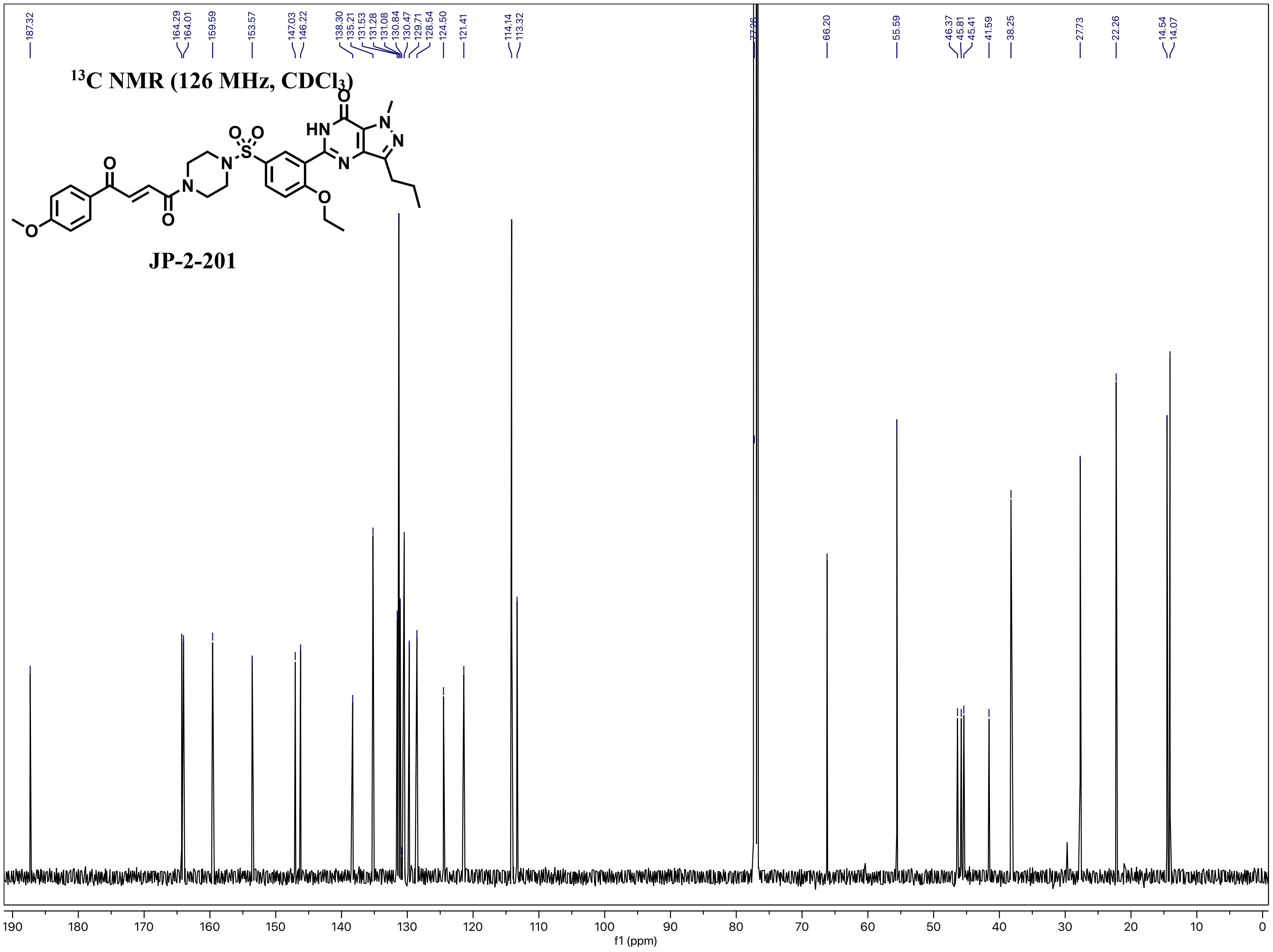

13C NMR (126 MHz, CDCl3)
JP-2-201
187.32
164.29
164.01
159.59
153.57
147.03
146.22
138.30
135.21
131.53
131.28
131.08
130.84
130.47
129.71
128.54
124.50
121.41
114.14
113.32
77.26
66.20
55.59
46.37
45.81
45.41
41.59
38.25
27.73
22.26
14.54
14.07
f1 (ppm)

$^1H$ NMR (700 MHz, $CDCl_3$)

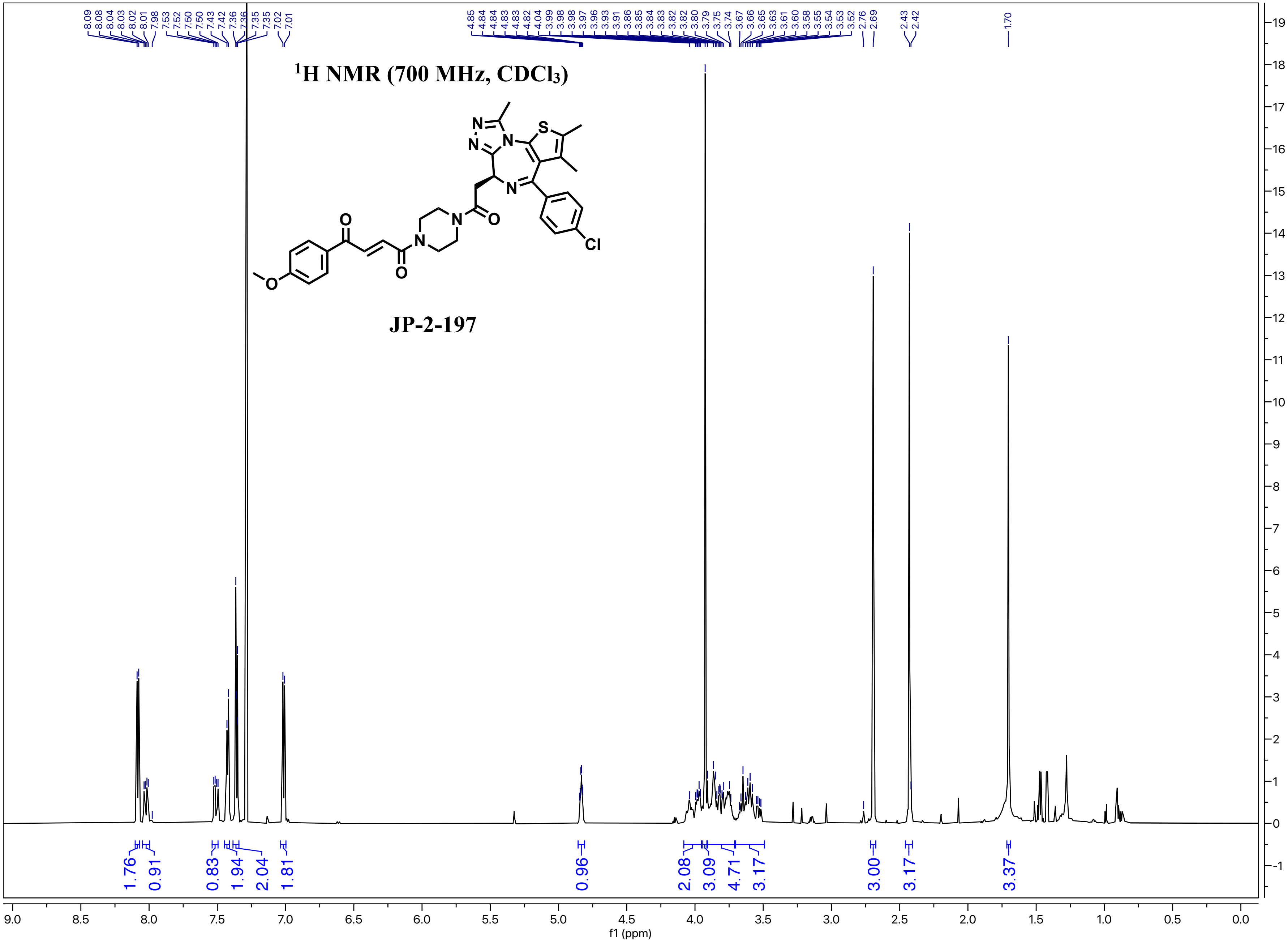


JP-2-197

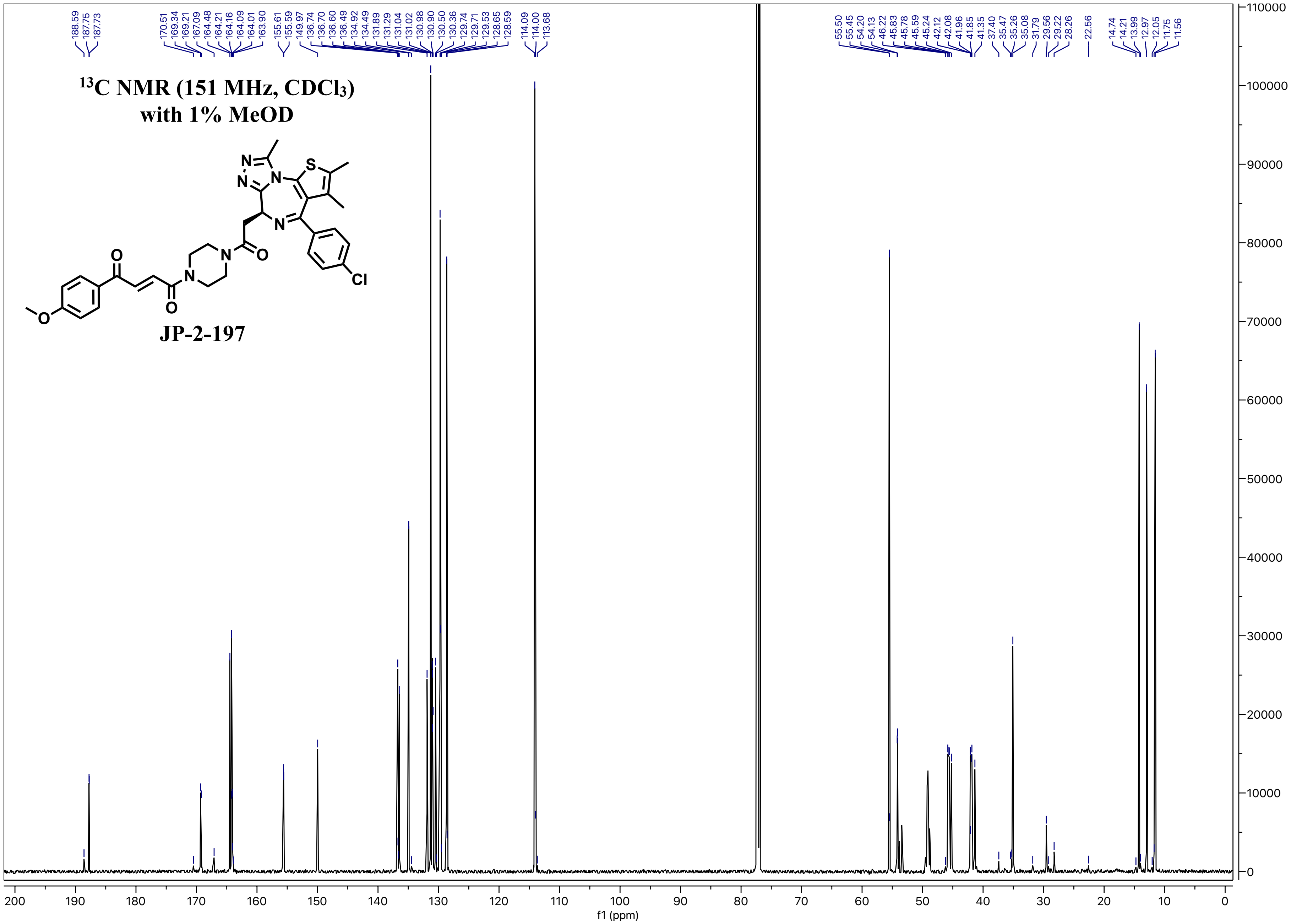

$^{13}$C NMR (151 MHz, CDCl$_3$)
with 1% MeOD
JP-2-197
188.59
187.75
187.73
170.51
169.34
169.21
167.09
164.48
164.21
164.16
164.09
164.01
163.90
155.61
155.59
149.97
136.74
136.70
136.60
136.49
134.92
134.49
131.89
131.29
131.04
131.02
130.98
130.90
130.50
130.36
129.74
129.71
129.53
128.65
128.59
114.09
114.00
113.68
55.50
55.45
54.20
54.13
46.22
45.83
45.78
45.59
45.24
42.12
42.08
41.96
41.85
41.35
37.40
35.47
35.26
35.08
31.79
29.56
29.22
28.26
22.56
14.74
14.21
13.99
12.97
12.05
11.75
11.56
f1 (ppm)

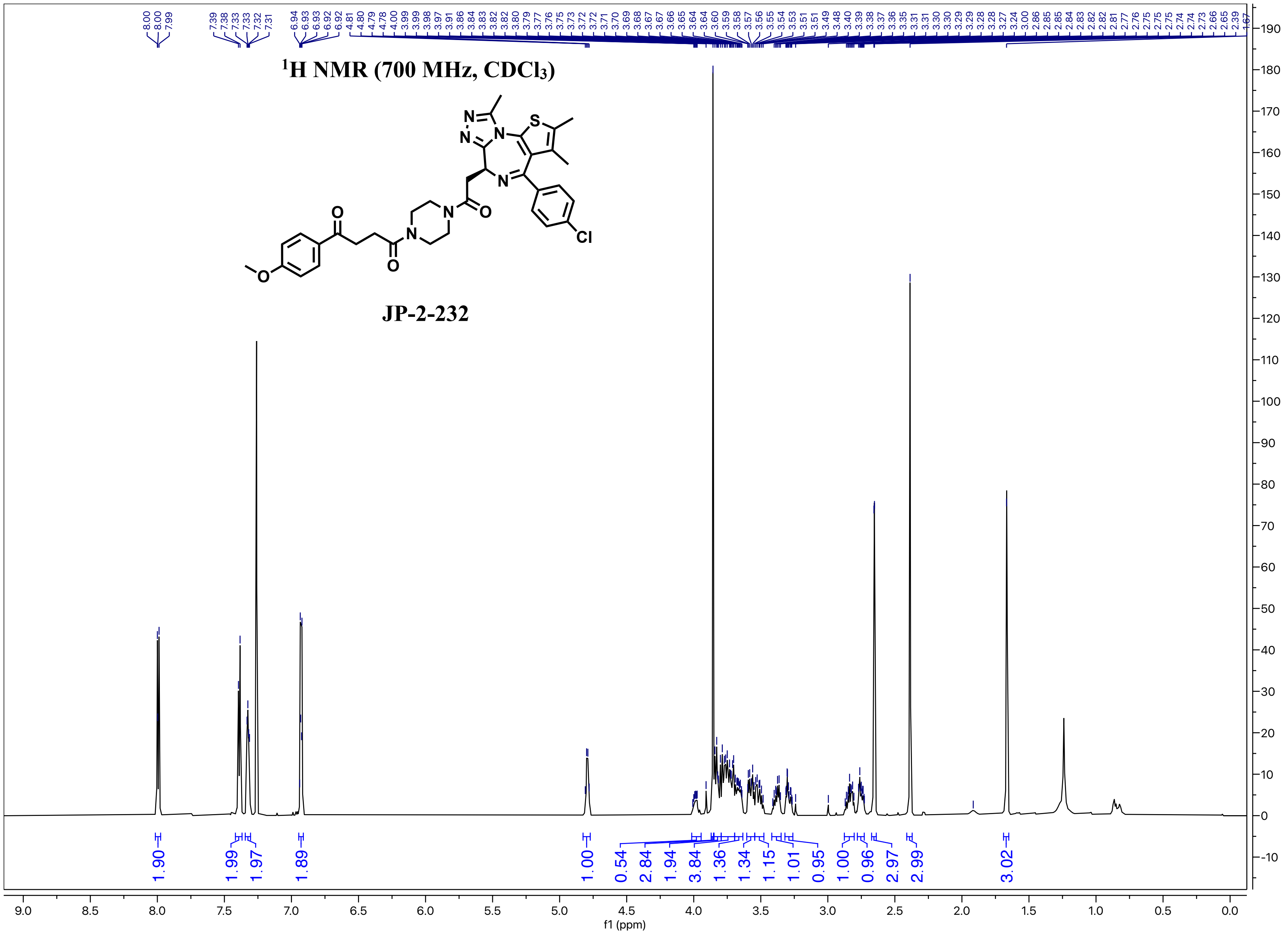

1H NMR (700 MHz, CDCl3)
JP-2-232
1.90
1.99
1.97
1.89
1.00
0.54
2.84
1.94
3.84
1.36
1.34
1.15
1.01
0.95
1.00
0.96
2.97
2.99
3.02
f1 (ppm)

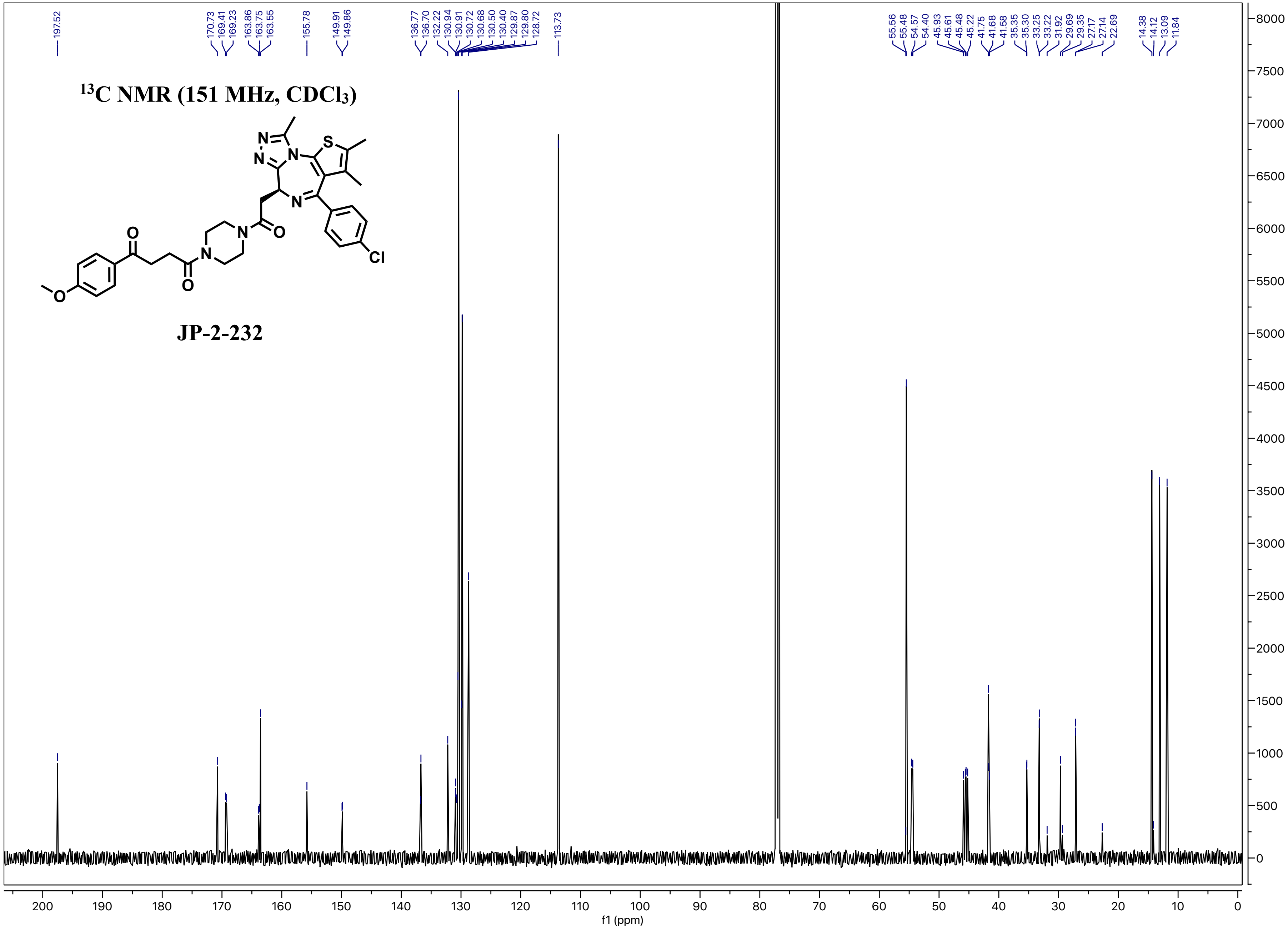

$^{13}C$ NMR (151 MHz, CDCl$_3$)
JP-2-232
197.52
170.73 169.41 169.23 163.86 163.75 163.55
155.78
149.91 149.86
136.77 136.70 132.22 130.94 130.91 130.72 130.68 130.50 130.40 129.87 129.80 128.72
113.73
55.56 55.48 54.57 54.40 45.93 45.61 45.48 45.22 41.75 41.68 41.58 35.35 35.30 33.25 33.22 31.92 29.69 29.35 27.17 27.14 22.69
14.38 14.12 13.09 11.84
f1 (ppm)

¹H NMR (700 MHz, CDCl₃)

**JP-2-216**

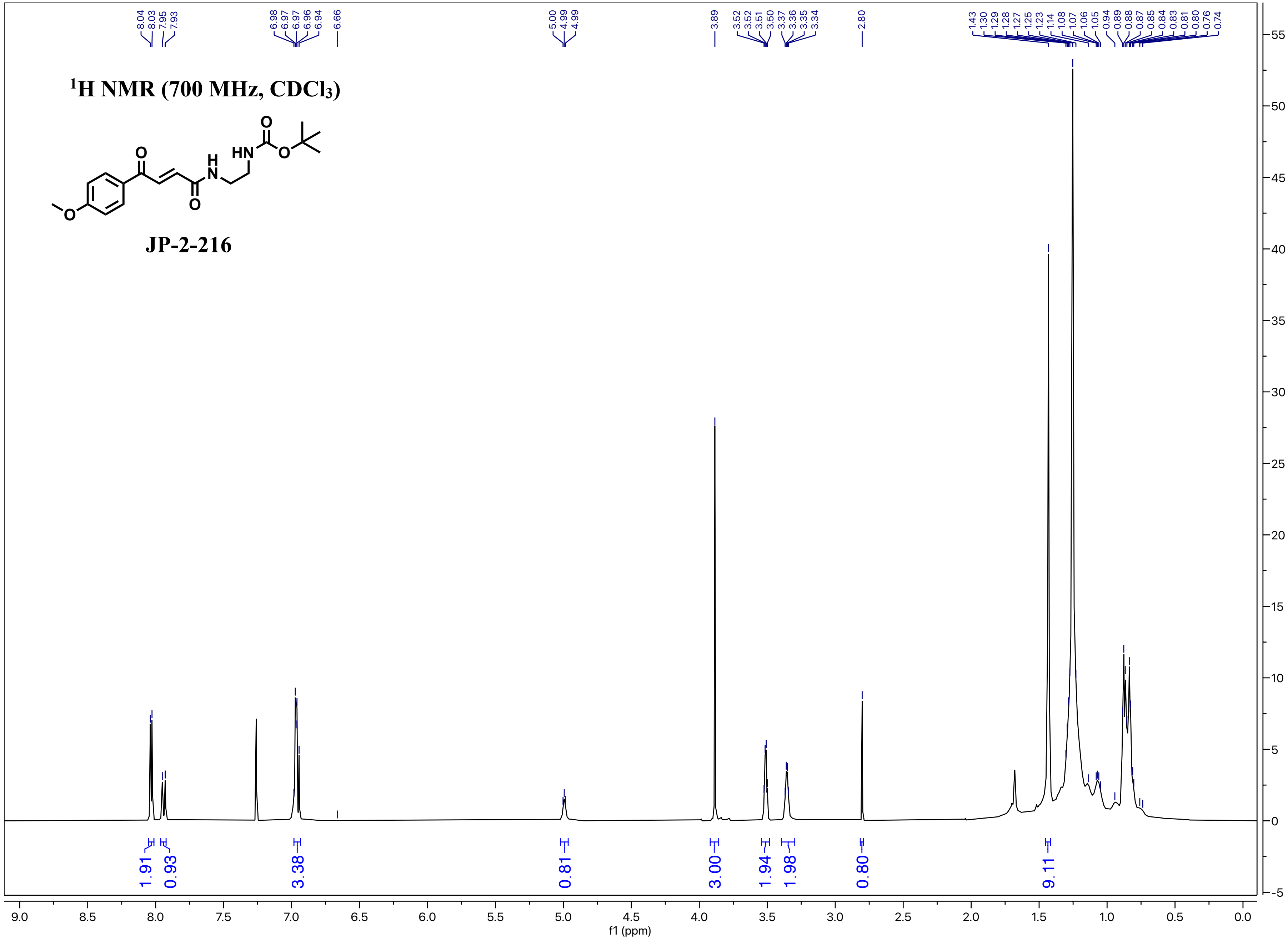

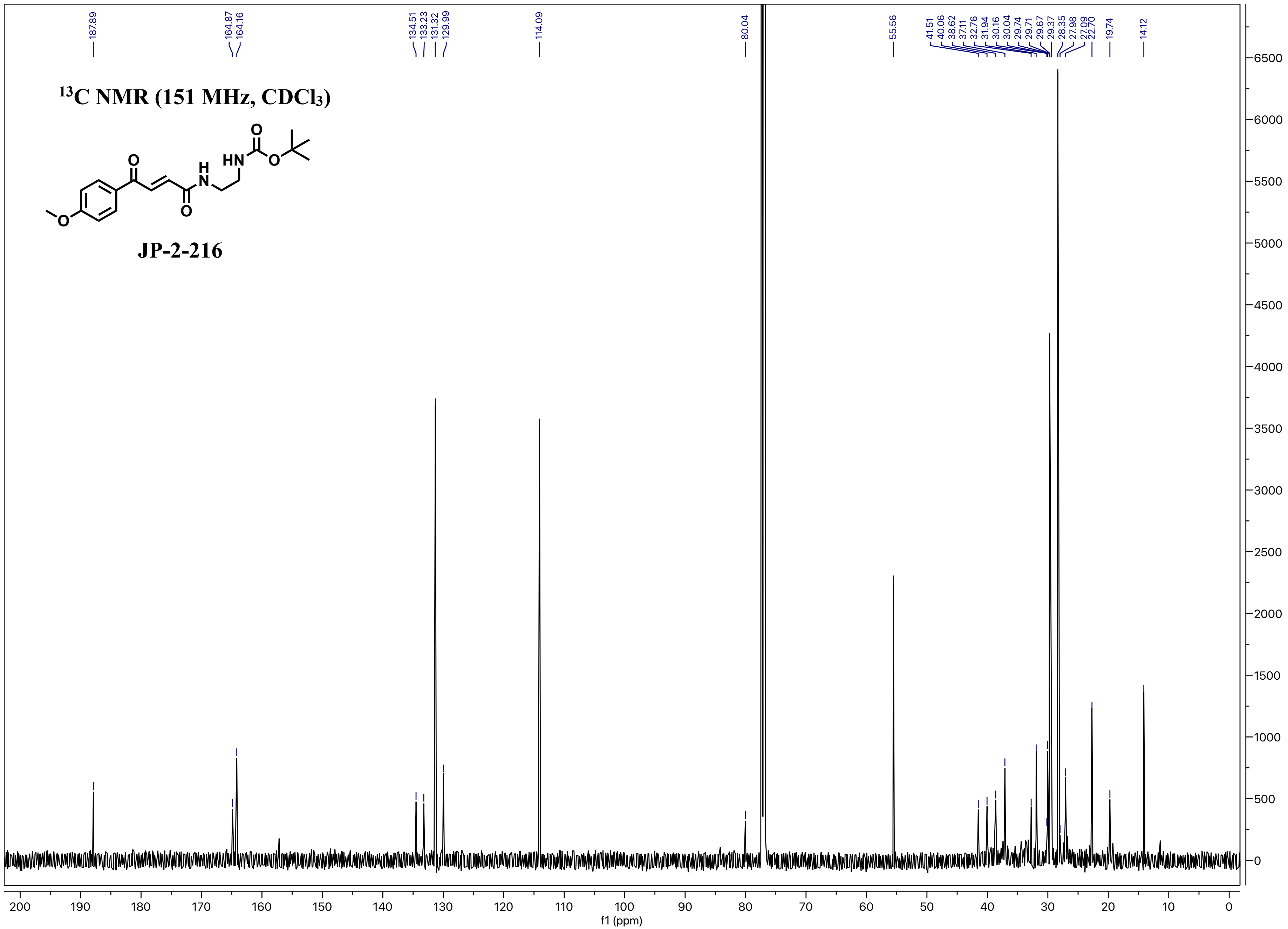

$^{13}$C NMR (151 MHz, CDCl$_3$)
JP-2-216
187.89
164.87
164.16
134.51
133.23
131.32
129.99
114.09
80.04
55.56
41.51
40.06
38.62
37.11
32.76
31.94
30.16
30.04
29.74
29.71
29.67
29.37
28.35
27.98
27.09
22.70
19.74
14.12
f1 (ppm)

#### [1]H NMR (700 MHz, $CDCl_3$)

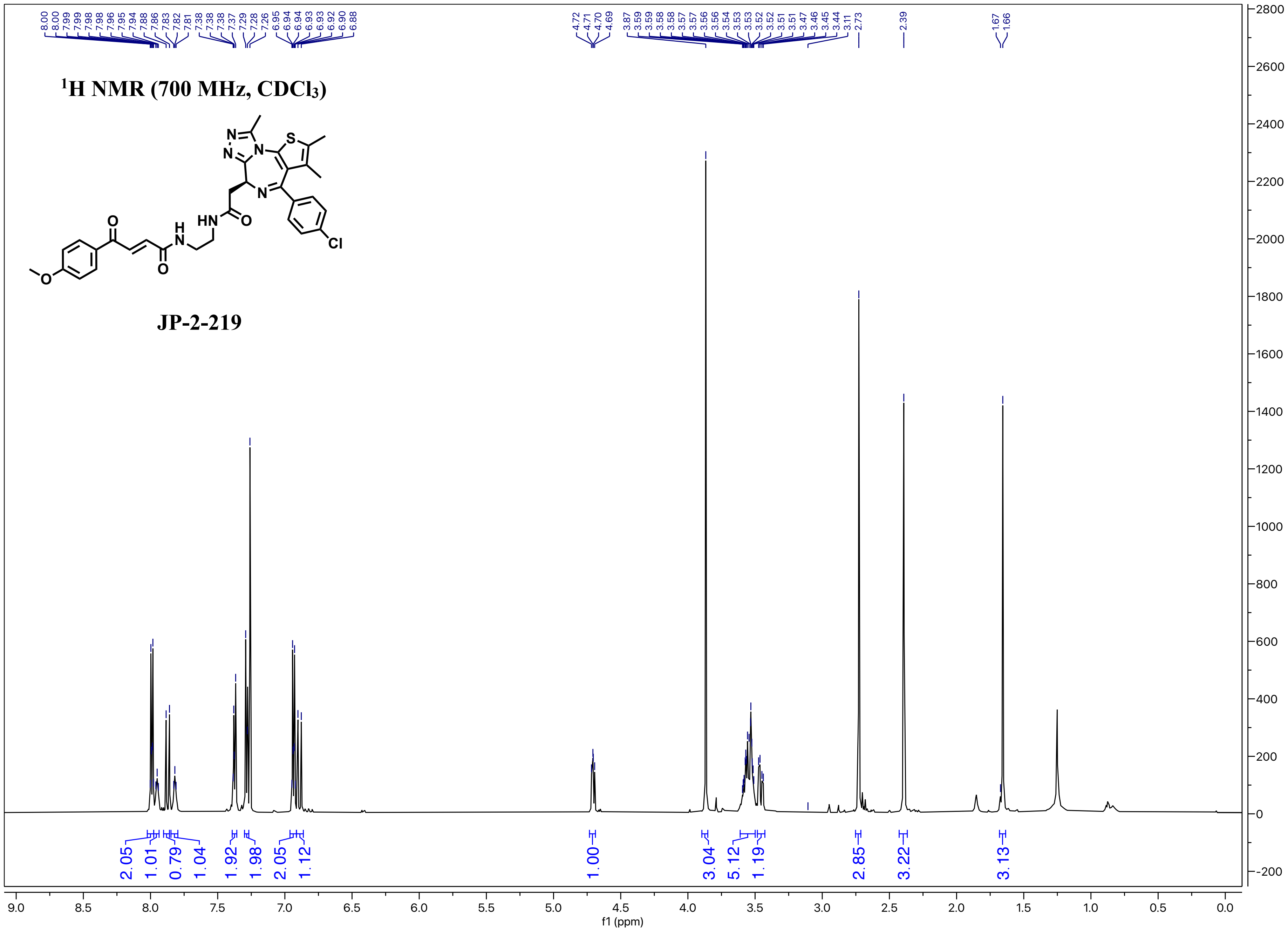

**JP-2-219**

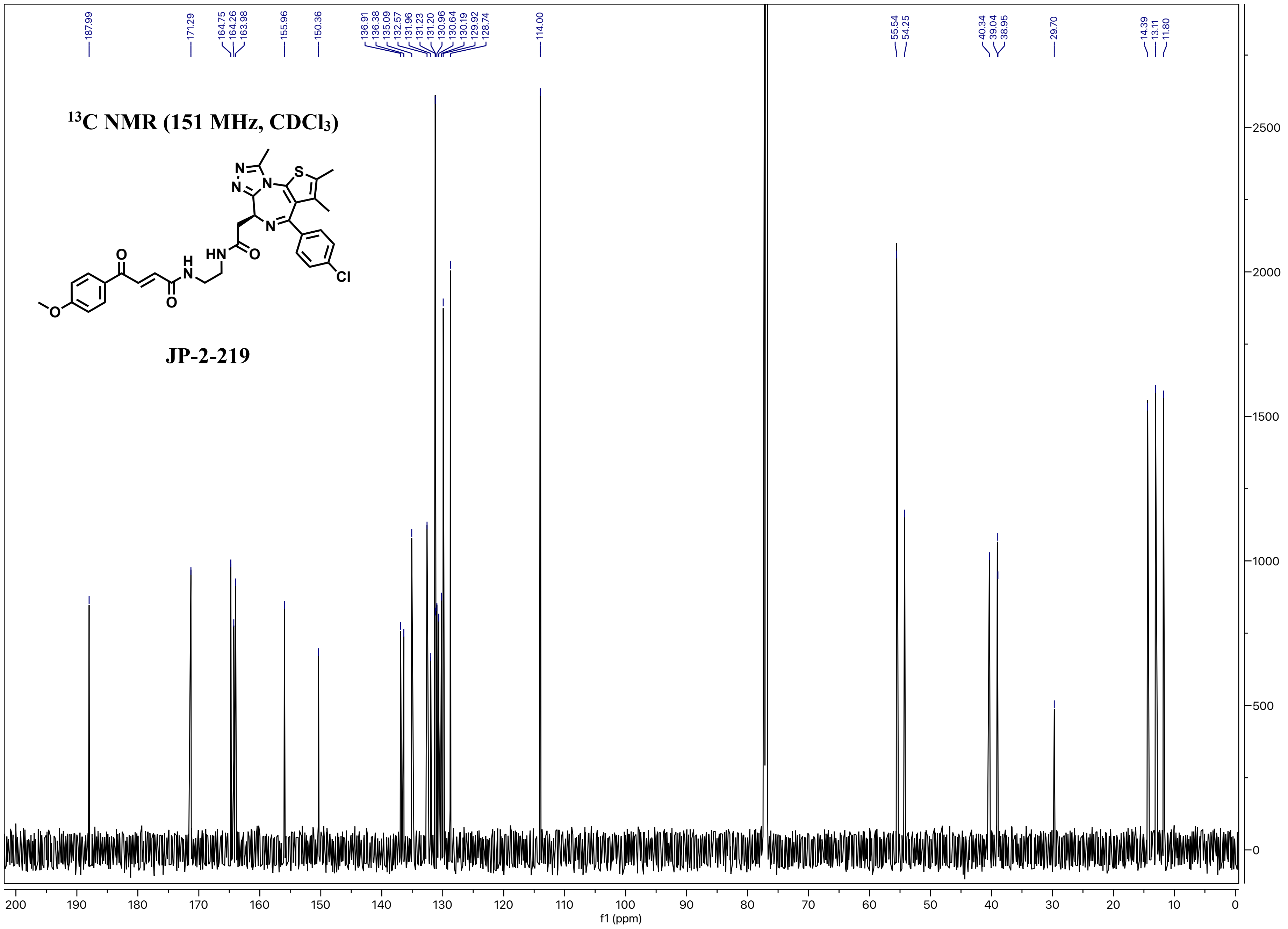

$^{13}C$ NMR (151 MHz, $CDCl_3$)
JP-2-219
187.99
171.29
164.75
164.26
163.98
155.96
150.36
136.91
136.38
135.09
132.57
131.96
131.23
131.20
130.96
130.64
130.19
129.92
128.74
114.00
55.54
54.25
40.34
39.04
38.95
29.70
14.39
13.11
11.80
f1 (ppm)

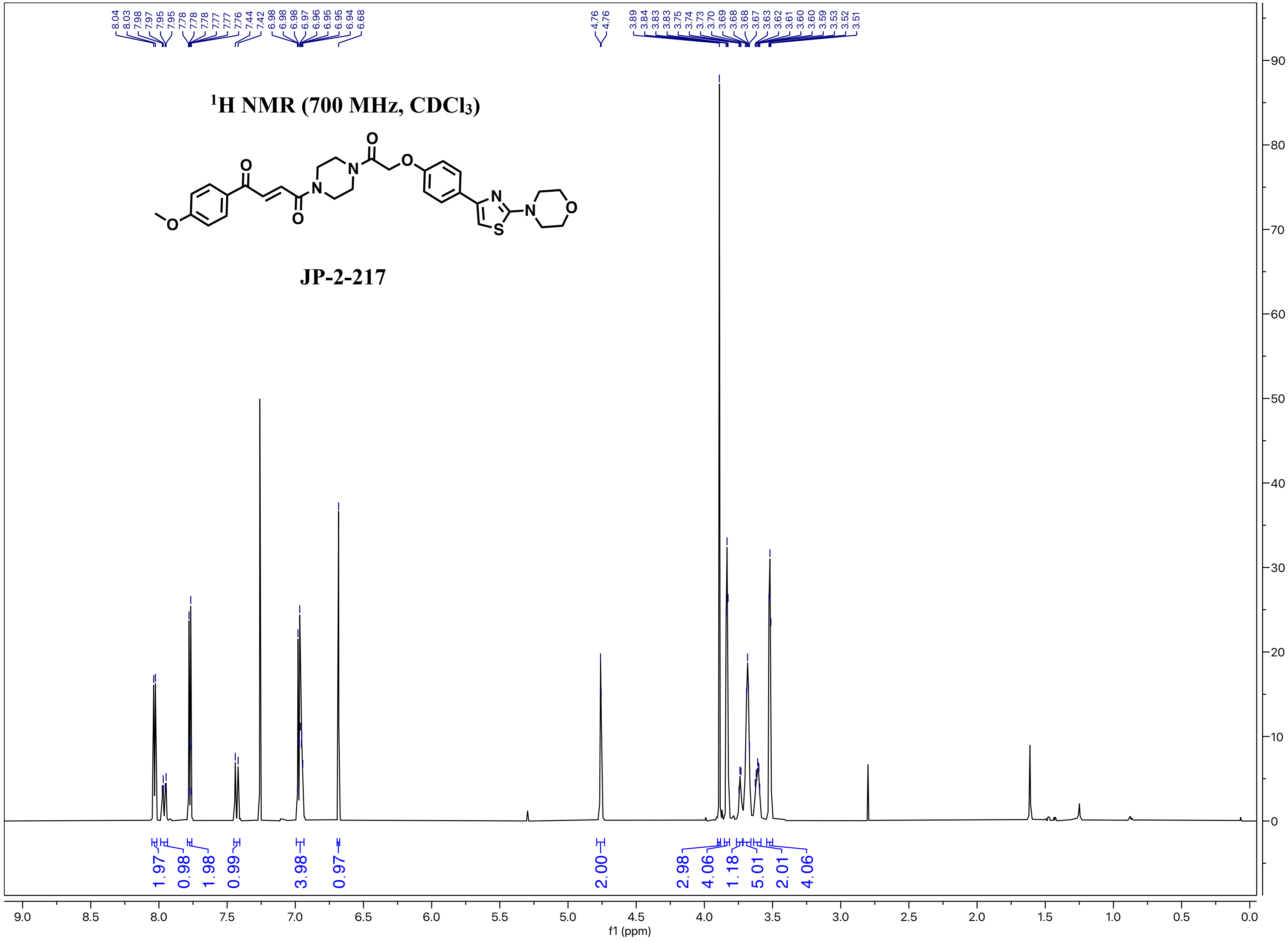

1H NMR (700 MHz, CDCl3)
JP-2-217
8.04 8.03 7.98 7.97 7.95 7.95 7.78 7.78 7.78 7.77 7.76 7.44 7.42 6.98 6.98 6.98 6.97 6.96 6.96 6.95 6.94 6.68
4.76 4.76
3.89 3.84 3.83 3.83 3.75 3.74 3.73 3.70 3.69 3.68 3.68 3.67 3.63 3.62 3.61 3.60 3.60 3.59 3.53 3.52 3.51
1.97 0.98 1.98 0.99 3.98 0.97 2.00 2.98 4.06 1.18 5.01 2.01 4.06
f1 (ppm)

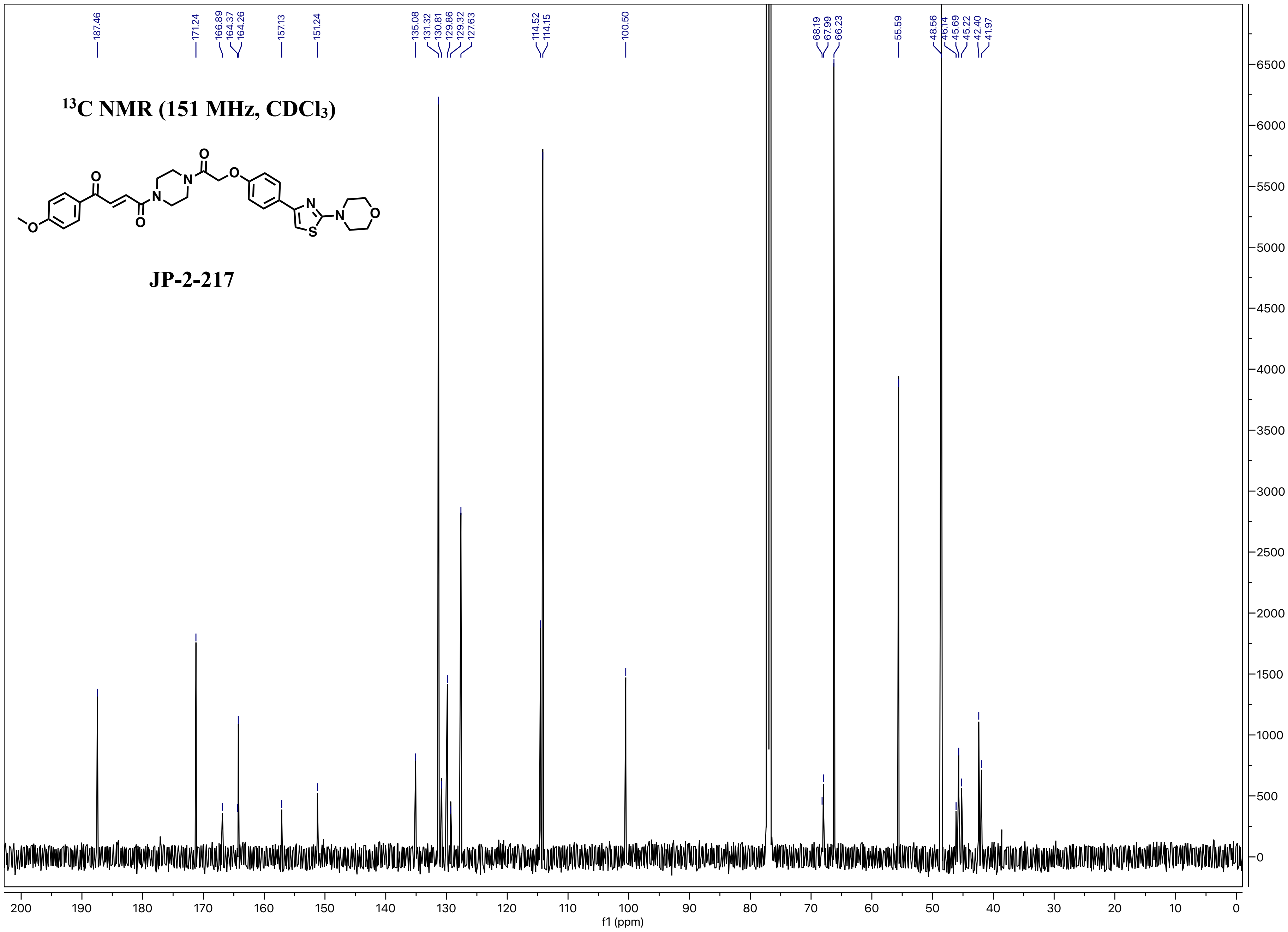
13C NMR (151 MHz, CDCl3)
JP-2-217
187.46
171.24
166.89
164.37
164.26
157.13
151.24
135.08
131.32
130.81
129.86
129.32
127.63
114.52
114.15
100.50
68.19
67.99
66.23
55.59
48.56
46.14
45.69
45.22
42.40
41.97
f1 (ppm)

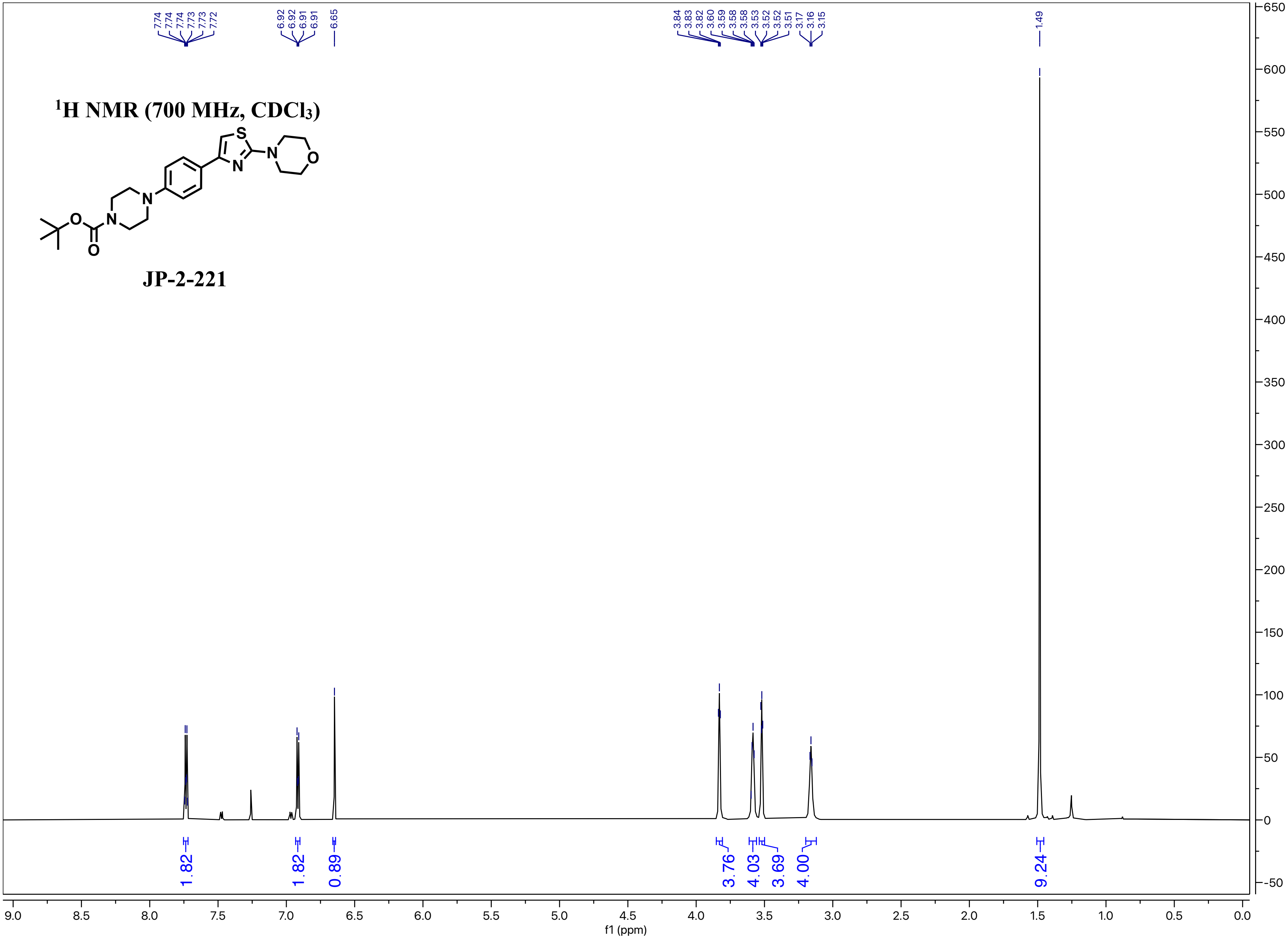

$^1$H NMR (700 MHz, CDCl$_3$)
JP-2-221
7.74 7.74 7.74 7.73 7.73 7.72
6.92 6.92 6.91 6.91
6.65
3.84 3.83 3.82 3.60 3.59 3.58 3.58 3.53 3.52 3.52 3.51 3.17 3.16 3.15
1.49
1.82
1.82
0.89
3.76
4.03
3.69
4.00
9.24
f1 (ppm)

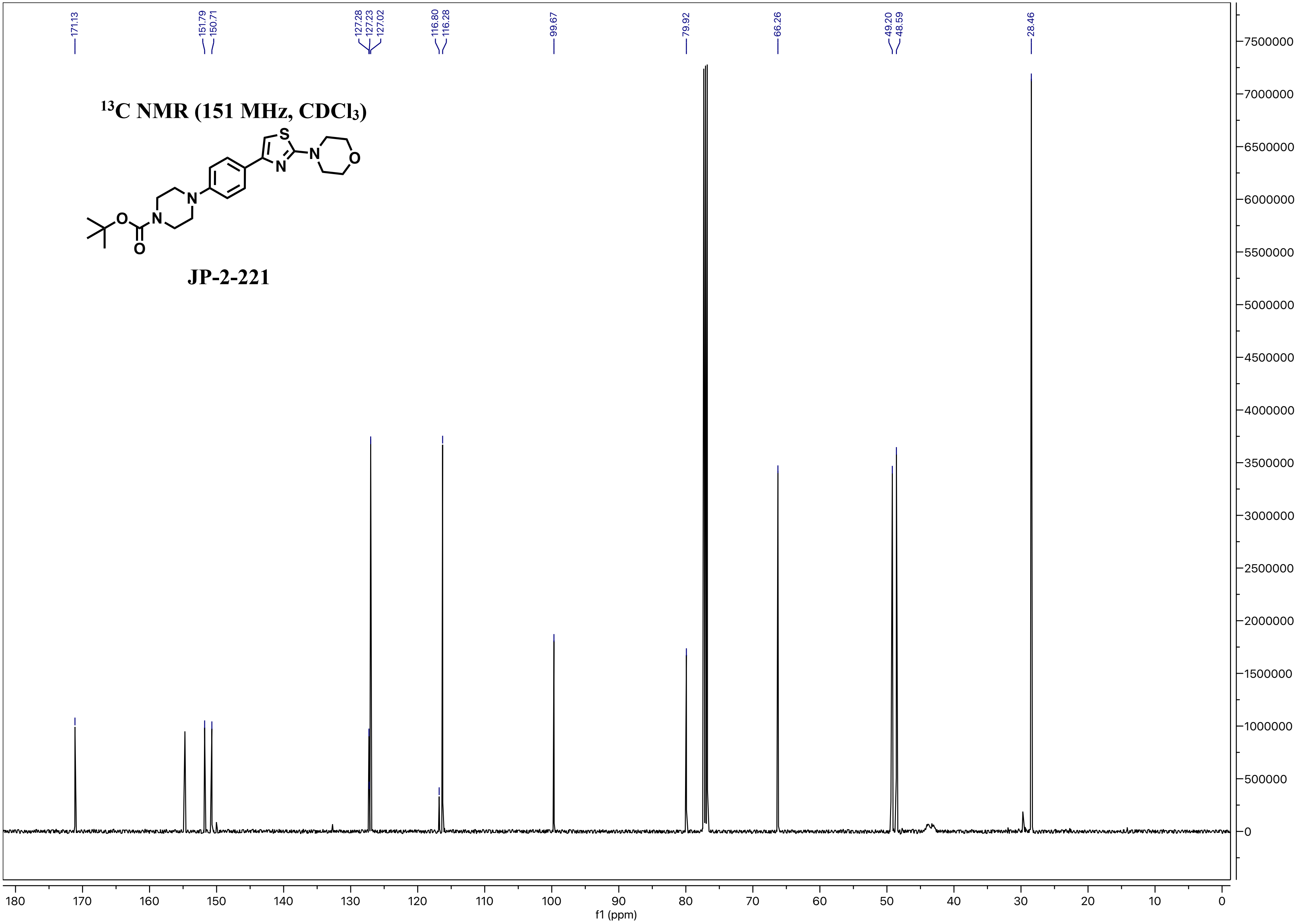

13C NMR (151 MHz, CDCl3)
JP-2-221
171.13
151.79
150.71
127.28
127.23
127.02
116.80
116.28
99.67
79.92
66.26
49.20
48.59
28.46
f1 (ppm)

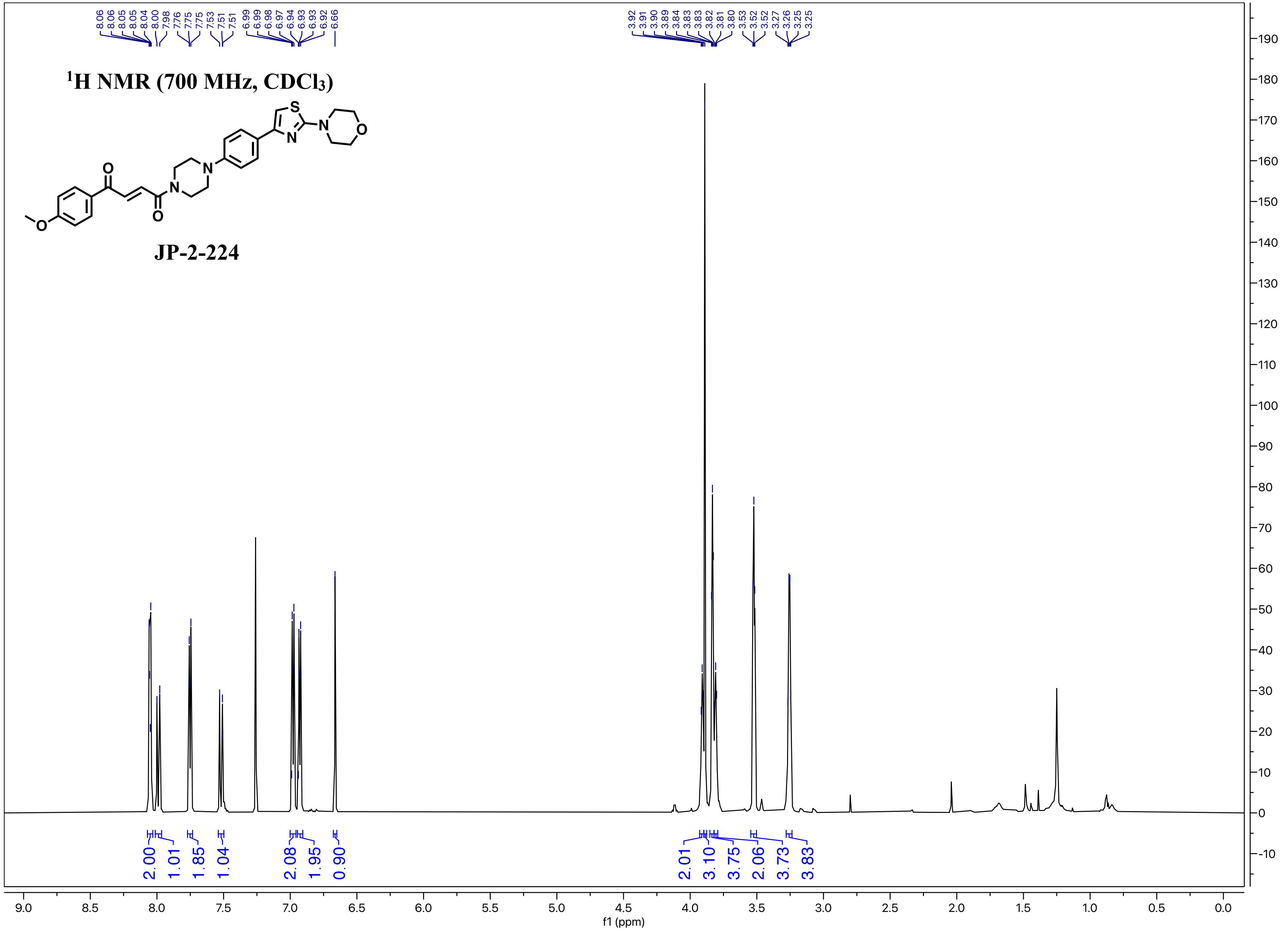

8.06 8.06 8.05 8.05 8.04 8.00 7.98 7.76 7.75 7.75 7.53 7.51 7.51 6.99 6.99 6.98 6.97 6.94 6.93 6.93 6.92 6.66
3.92 3.91 3.90 3.89 3.84 3.83 3.83 3.82 3.81 3.80 3.53 3.52 3.52 3.27 3.26 3.25 3.25
$^{1}$H NMR (700 MHz, $CDCl_3$)
JP-2-224
2.00 1.01 1.85 1.04 2.08 1.95 0.90
2.01 3.10 3.75 2.06 3.73 3.83
f1 (ppm)

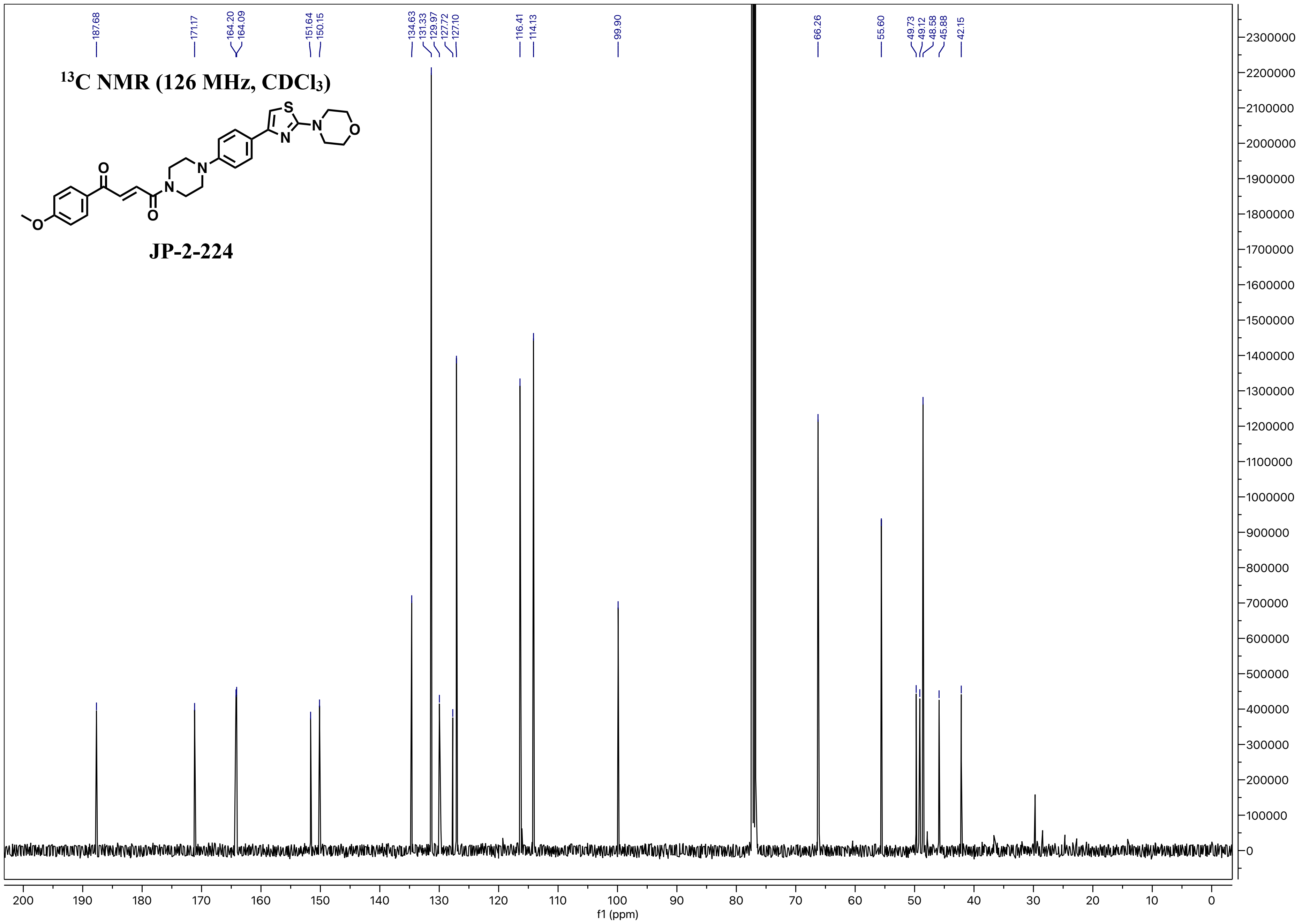

13C NMR (126 MHz, CDCl3)
JP-2-224
S
N
N
O
O
N
N
O
O
187.68
171.17
164.20
164.09
151.64
150.15
134.63
131.33
129.97
127.72
127.10
116.41
114.13
99.90
66.26
55.60
49.73
49.12
48.58
45.88
42.15
f1 (ppm)
200 190 180 170 160 150 140 130 120 110 100 90 80 70 60 50 40 30 20 10 0
2300000 2200000 2100000 2000000 1900000 1800000 1700000 1600000 1500000 1400000 1300000 1200000 1100000 1000000 900000 800000 700000 600000 500000 400000 300000 200000 100000 0

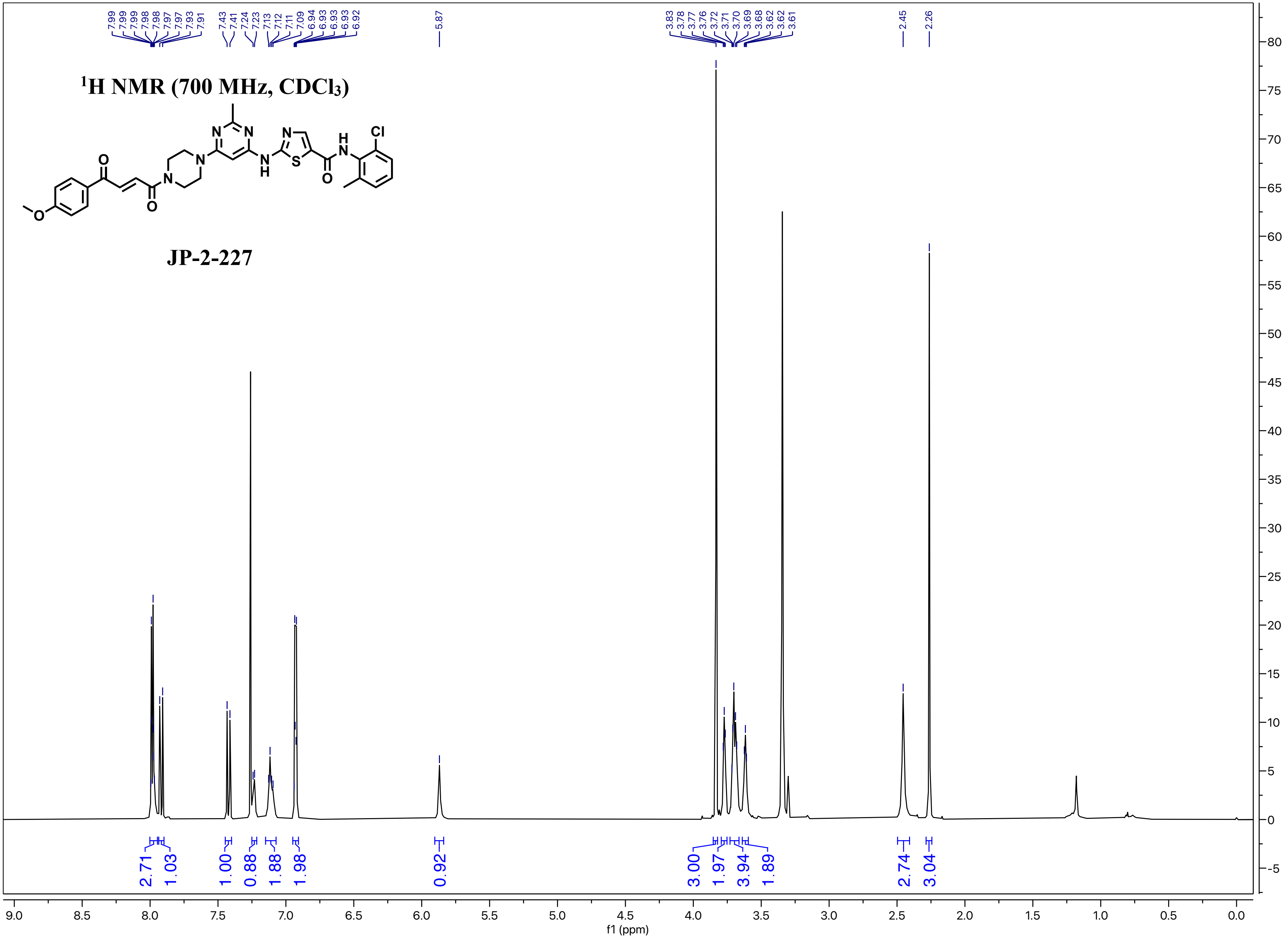

[1]H NMR (700 MHz, CDCl3)
JP-2-227
7.99 7.99 7.99 7.98 7.98 7.97 7.97 7.93 7.91
7.43 7.41 7.24 7.23 7.13 7.12 7.11 7.09 6.94 6.93 6.93 6.93 6.92
5.87
3.83 3.78 3.77 3.76 3.72 3.71 3.70 3.69 3.68 3.62 3.62 3.61
2.45
2.26
2.71 1.03 1.00 0.88 1.88 1.98 0.92 3.00 1.97 3.94 1.89 2.74 3.04
f1 (ppm)

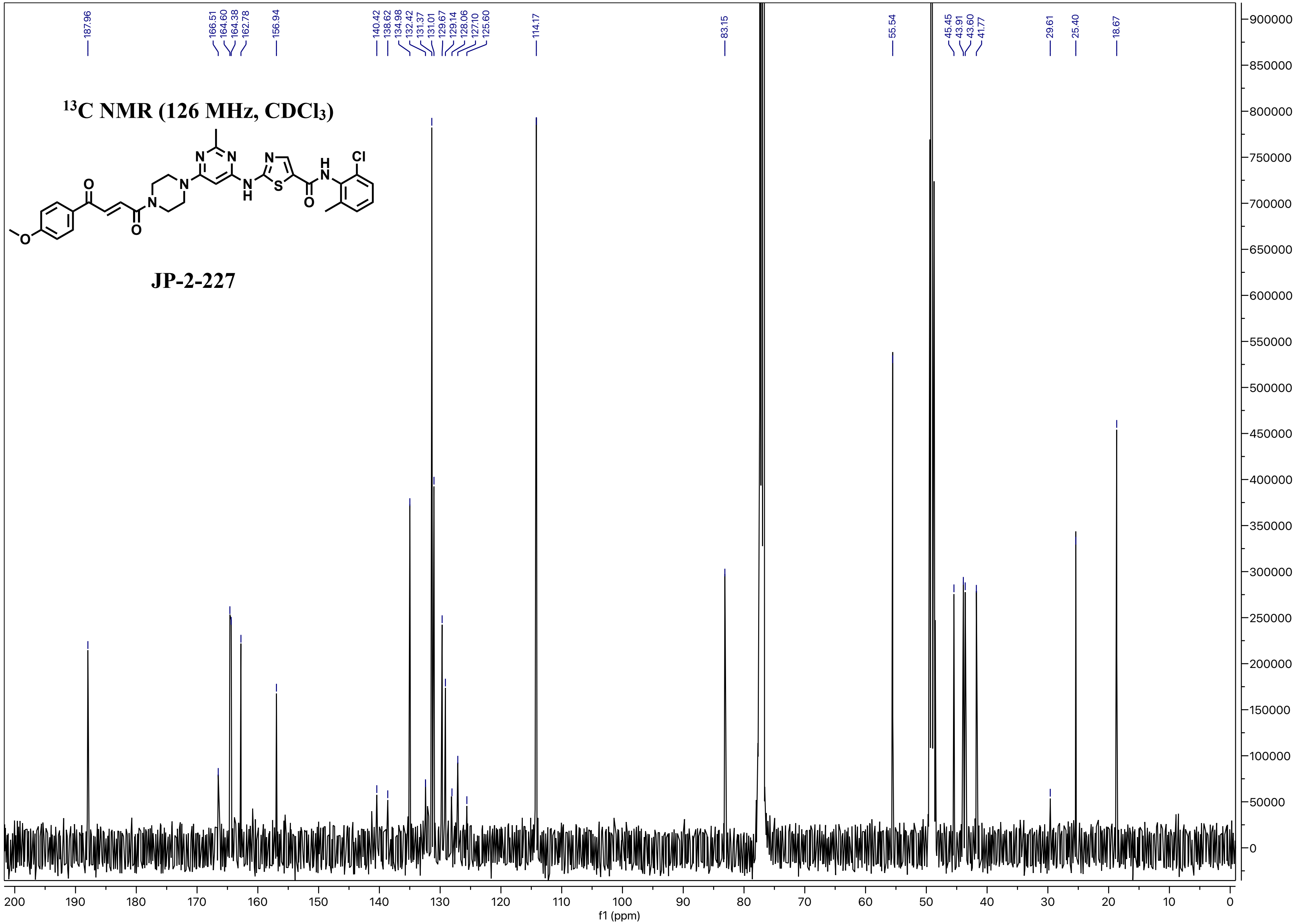
13C NMR (126 MHz, CDCl3)
JP-2-227
O
O
O
N
N
N
N
N
H
N
S
O
N
H
N
Cl
187.96
166.51
164.60
164.38
162.78
156.94
140.42
138.62
134.98
132.42
131.37
131.01
129.67
129.14
128.06
127.10
125.60
114.17
83.15
55.54
45.45
43.91
43.60
41.77
29.61
25.40
18.67
f1 (ppm)

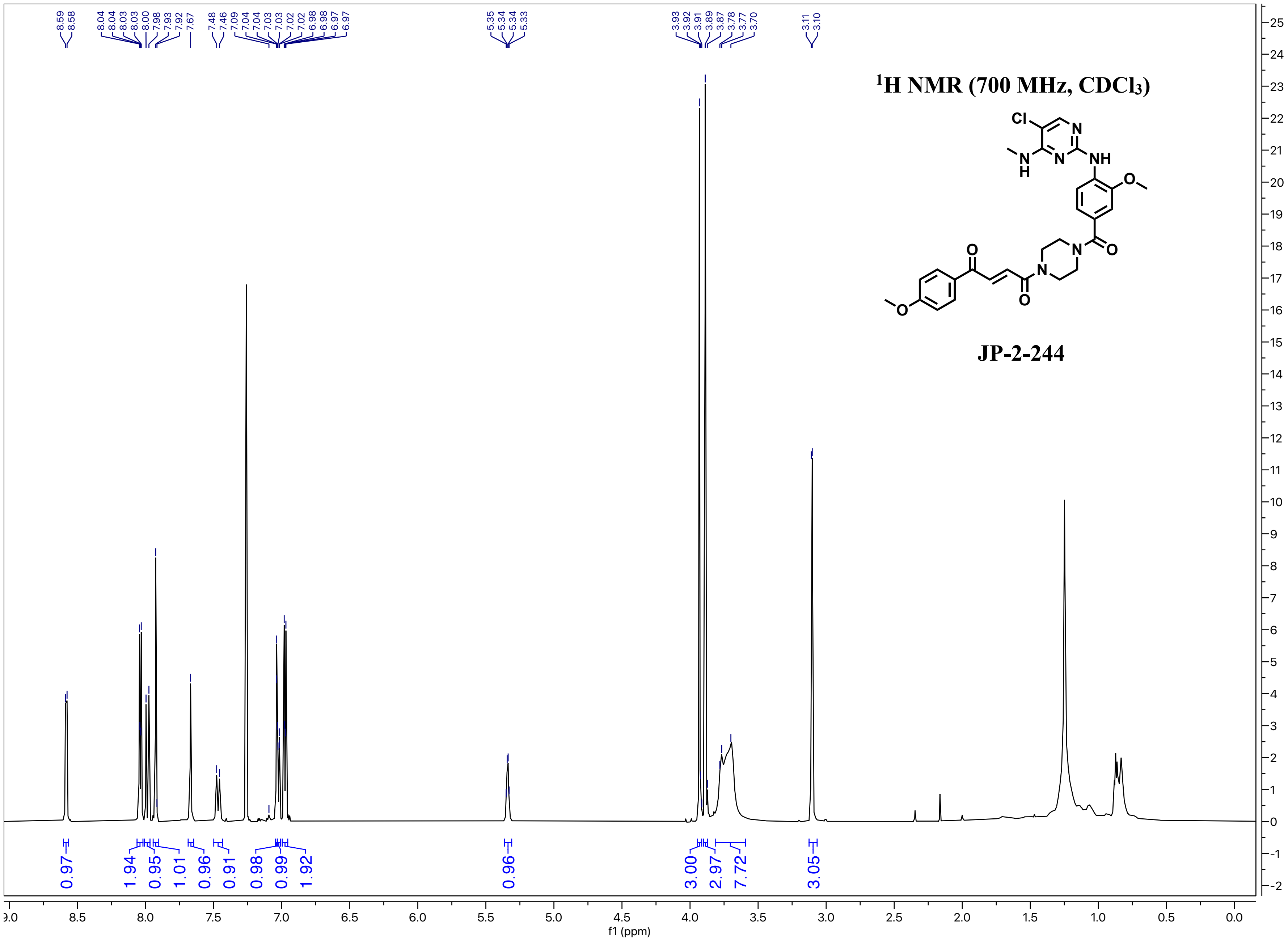

8.59
8.58
8.04
8.04
8.03
8.03
8.00
7.98
7.93
7.92
7.67
7.48
7.46
7.09
7.04
7.04
7.03
7.03
7.02
7.02
6.98
6.98
6.97
6.97
5.35
5.34
5.34
5.33
3.93
3.92
3.91
3.89
3.87
3.78
3.77
3.70
3.11
3.10
¹H NMR (700 MHz, CDCl₃)
Cl
N
NH
N
H
O
O
N
O
N
O
O
JP-2-244
0.97
1.94
0.95
1.01
0.96
0.91
0.98
0.99
1.92
0.96
3.00
2.97
7.72
3.05
f1 (ppm)

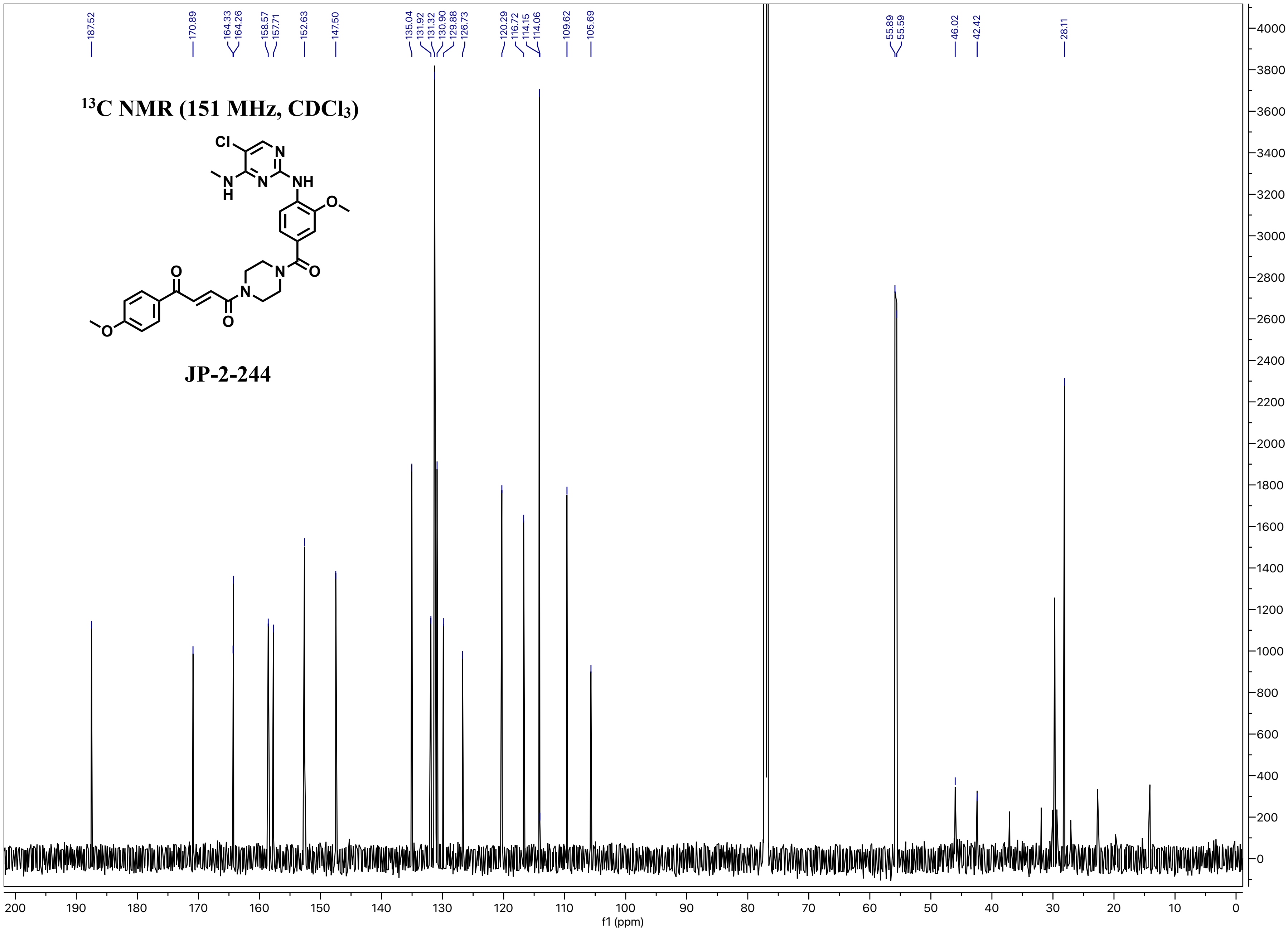

13C NMR (151 MHz, CDCl3)
JP-2-244
Cl
N
NH
N
H
O
O
N
O
N
O
O
187.52
170.89
164.33
164.26
158.57
157.71
152.63
147.50
135.04
131.92
131.32
130.90
129.88
126.73
120.29
116.72
114.15
114.06
109.62
105.69
55.89
55.59
46.02
42.42
28.11
f1 (ppm)

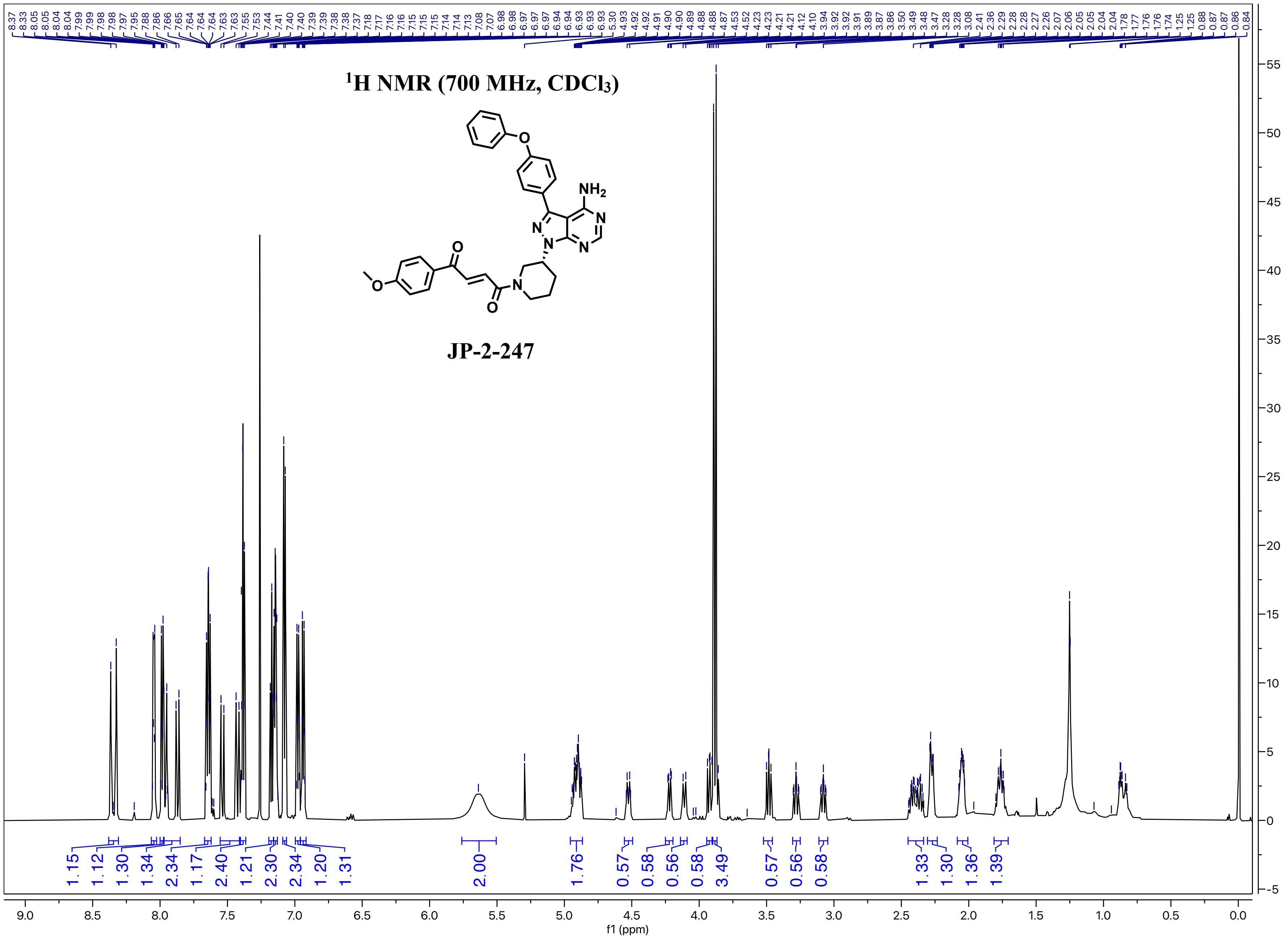

$^1$H NMR (700 MHz, CDCl$_3$)
NH$_2$
JP-2-247
f1 (ppm)

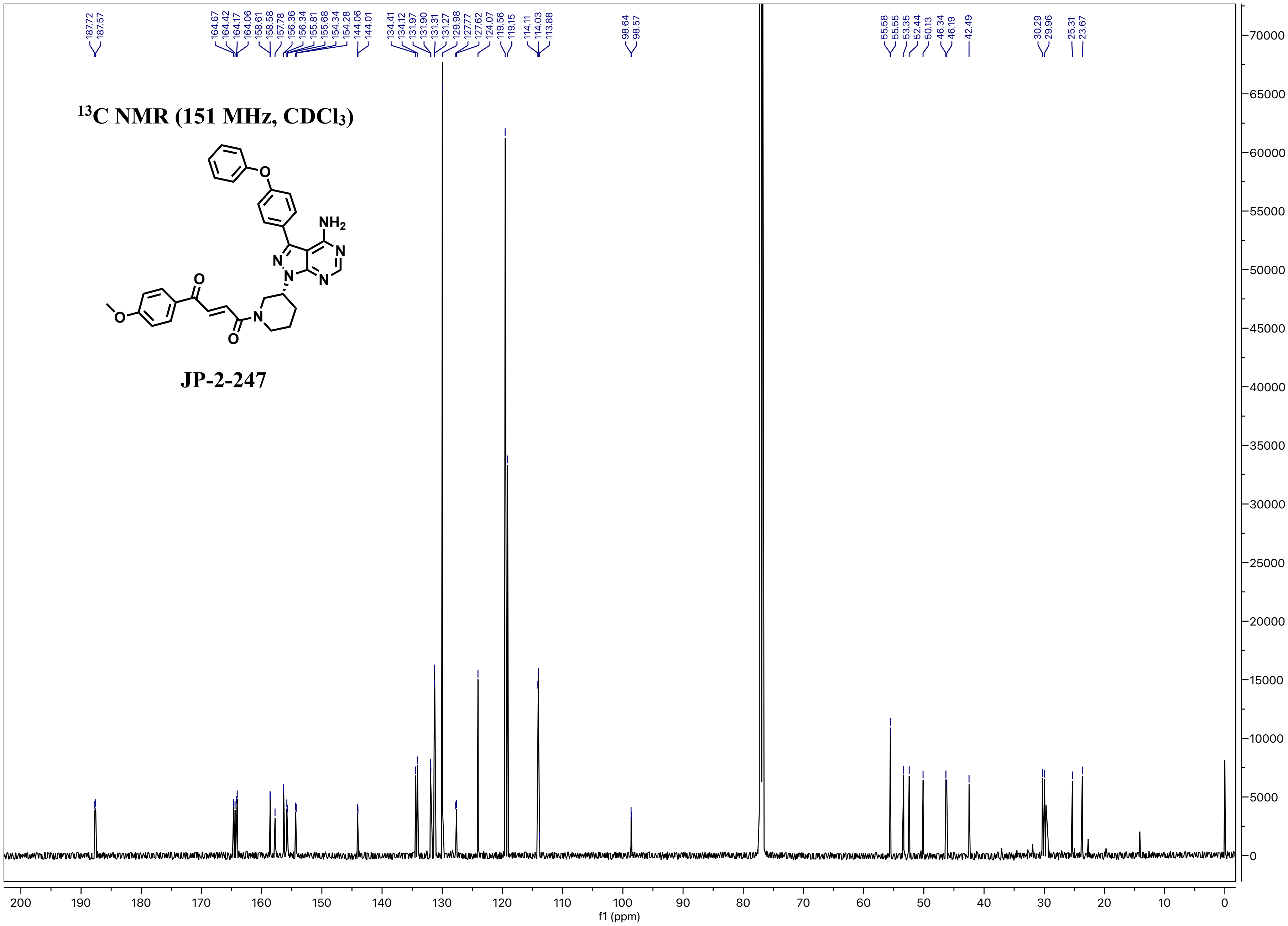

$^{13}C$ NMR (151 MHz, $CDCl_3$)
JP-2-247
NH2
N
O
187.72
187.57
164.67
164.42
164.17
164.06
158.61
158.58
157.78
156.36
156.34
155.81
155.68
154.34
154.28
144.06
144.01
134.41
134.12
131.97
131.90
131.31
131.27
129.98
127.77
127.62
124.07
119.56
119.15
114.11
114.03
113.88
98.64
98.57
55.58
55.55
53.35
52.44
50.13
46.34
46.19
42.49
30.29
29.96
25.31
23.67
f1 (ppm)

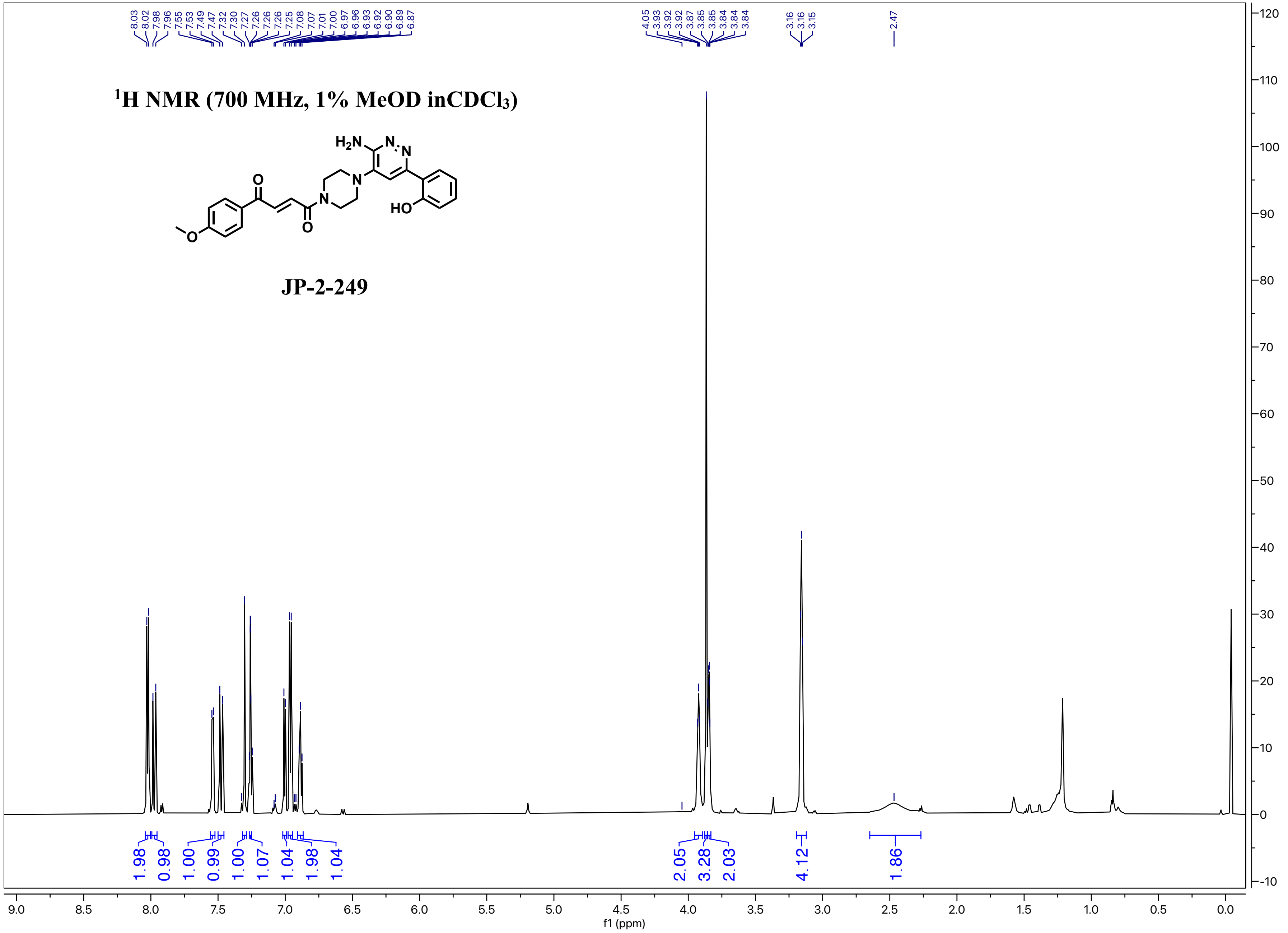

1H NMR (700 MHz, 1% MeOD inCDCl3)
H2N
N
N
O
N
N
HO
O
O
JP-2-249
8.03 8.02 7.98 7.96 7.55 7.53 7.49 7.47 7.32 7.30 7.27 7.26 7.26 7.26 7.25 7.08 7.07 7.01 7.00 6.97 6.96 6.93 6.92 6.90 6.89 6.87
4.05 3.93 3.92 3.92 3.87 3.85 3.85 3.84 3.84 3.84
3.16 3.16 3.15
2.47
1.98 0.98 1.00 0.99 1.00 1.07 1.04 1.98 1.04
2.05 3.28 2.03
4.12
1.86
f1 (ppm)

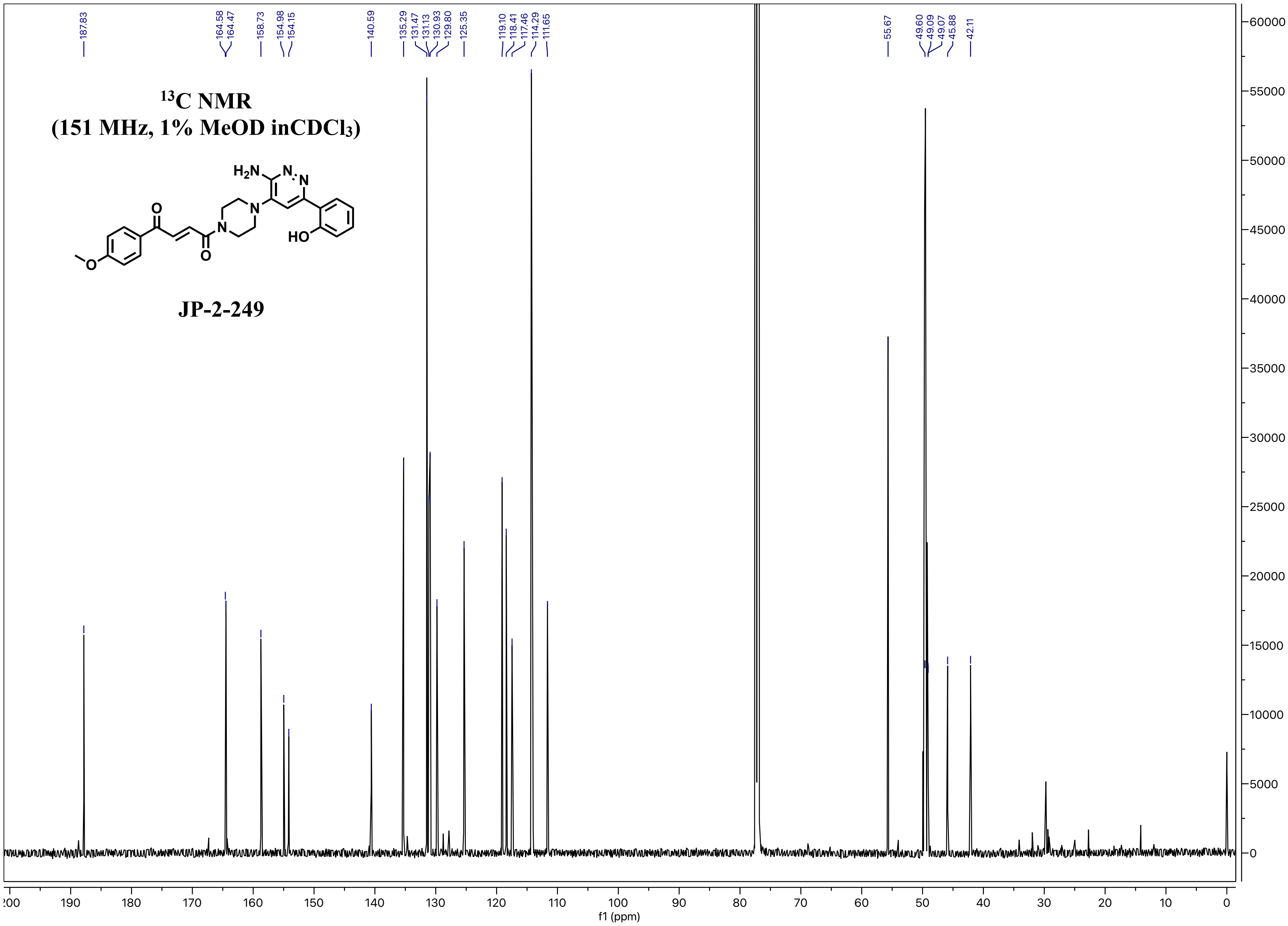

187.83
164.58
164.47
158.73
154.98
154.15
140.59
135.29
131.47
131.13
130.93
129.80
125.35
119.10
118.41
117.46
114.29
111.65
55.67
49.60
49.09
49.07
45.88
42.11
$^{13}$C NMR
(151 MHz, 1% MeOD inCDCl$_3$)
H$_2$N
N
N
N
N
O
O
O
HO
JP-2-249
f1 (ppm)
